# The cysteine-reactive covalent RNF4 ligand CCW16 induces ferroptosis in AML cells by activation of ROS signaling

**DOI:** 10.1101/2024.12.12.628083

**Authors:** Gina Gotthardt, Janik Weckesser, Georg Tascher, Sara Barros da Gama, Hannah J. Uckelmann, Martin Schwalm, Thorsten Mosler, Giulio Ferrario, Christian Münch, Stefan Knapp, Stefan Müller

## Abstract

The SUMO-targeted E3 ubiquitin ligase RNF4 plays an important role in safeguarding genome and proteome integrity. RNF4 recognizes polySUMO modified proteins and induces their proteolytic or non-proteolytic ubiquitylation. Given the key function of RNF4 in maintaining proteome and genome stability, we explored its role in cancer cell survival and as a potential cancer drug target. We found that RNF4 is overexpressed in acute myeloid leukemia and high expression levels correlate with poor survival of patients. Depletion of RNF4 exerts antiproliferative effects in AML cells and increased their sensitivity towards antileukemic drugs thus validating RNF4 as a potential vulnerability in AML. To develop PROTACs for targeted degradation of RNF4, we aimed to exploit the recently described cysteine-reactive RNF4 ligand CCW16, and synthesized a set of CCW16-derived degrader molecules using established VHL- and CRBN-E3 ligands. We validated covalent binding of CCW16 and CCW16-derived PROTACs to cysteine residues outside the RING domain of RNF4 using recombinant protein. However, none of the synthesized PROTACs were active in degrading RNF4 in a panel of different cell lines. Consistent with the high reactivity of the chosen electrophile, we detected in the cellular context covalent attachment of CCW16 to accessible cysteines in a large number of proteins including covalent linkage with the catalytic residues in members of the peroxiredoxin family. Furthermore, exposure of cells to CCW16-derived PROTACs induced upregulation of heme oxygenase 1, a ferroptosis marker and impaired cell viability in a distinct, RNF4-independent, ferroptotic cell death pathway.

## Introduction

Cells are constantly exposed to endogenous or exogenous hazards that threaten their genome or proteome integrity. Sophisticated and highly regulated DNA repair pathways safeguard genome integrity preventing the emergence and accumulation of genetic alterations that are a hallmark of tumorigenesis^1–3^. At the same time, cancer chemotherapy largely relies on the induction of genotoxic stress in order to expose DNA damage as a vulnerability of highly proliferating and mutation-prone cancer cells^4–6^. However, a hallmark of cancer cells is their ability to tolerate endogenous or therapy-induced genotoxic stress, which confers resistance to DNA damaging chemotherapies^7^. An emerging strategy to break this resistance mechanism is the targeting of DNA repair and DNA damage response factors^8^.

Accumulating evidence suggests that post-translational modifications by ubiquitin and the small ubiquitin-related modifier (SUMO) are necessary to maintain genome stability by reversibly regulating protein function and activity^9,10^. Ubiquitin is conjugated to specific lysine residues of target proteins by a multistep enzymatic cascade of E1 activating enzymes, E2 conjugating enzymes and E3 protein ligases leading to proteasomal degradation or modulation of protein activities depending on the linkage type of the ubiquitin chain^11,12^. Similar to ubiquitination, also SUMOylation occurs via a three-step cascade involving a single dimeric E1 enzyme, an E2 conjugating enzyme and E3 SUMO ligases attaching either SUMO1 or the highly related SUMO2/3 to the target. Like ubiquitin, SUMO can be conjugated as a monomer (preferably SUMO1) or as chains by SUMO-SUMO linkages via internal lysine residues (mostly SUMO2/3). In principle, SUMOylation is a non-proteolytic modification, controlling the dynamics of protein-protein interactions^13–16^. However, SUMO chains, which are typically formed in response to genotoxic or proteotoxic stress, can connect SUMO signaling to the ubiquitin system via SUMO-targeted ubiquitin ligases (StUbLs)^17^. In mammalian cells, three StUbLs, RNF4, TOPORS and RNF111 have been identified. Their common structural characteristic is a RING-type ubiquitin ligase domain and the presence of tandemly repeated SUMO interaction motifs (SIMs) enabling them to specifically target and ubiquitinate polySUMOylated substrates^18–20^.

RNF4 is a monomeric RING-type E3 ubiquitin ligase, consisting of four SIMs located in the N-terminal region. These SIMs target RNF4 to SUMOylated proteins, and a RING finger domain in the RNF4 C-terminus catalyzes the transfer of ubiquitin from an E2 enzyme to the target^21^. RNF4 plays a key role in the proteasome-dependent removal of misfolded or damaged nuclear proteins and regulates the resolution of cytoplasmic stress granules^22–24^. During DNA repair, RNF4 controls chromatin residency of DNA damage response (DDR) and repair factors and it is critical for the reversal of stalled replication forks^25–27^. In addition, RNF4 mediates the proteasomal clearance of SUMOylated post-replicative DNA protein crosslinks (DPCs), including TOP1/2- and DNMT1-DPCs which are induced by topoisomerase poisons or hypomethylating agents^20,28,29^.

Given that RNF4 maintains proteome and genome stability and that it is overexpressed in different tumor entities such as acute myeloid leukemia (AML), RNF4 may represent a vulnerability for cancer^17,23,27^. As recently shown, the depletion of RNF4 in mice expressing the oncogene *c-myc* prolongs tumor-free survival, providing a compelling rationale for targeting RNF4 as an anticancer therapy^30^.

With the advent of targeted protein degradation (TPD), a new pharmacological modality has become available that allows also the targeting of poorly druggable proteins. One class of TPD compounds are PROteolysis TArgeting Chimeras (PROTACs), consisting of two ligands, coupled by a short linker. One ligand interacts with the protein of interest (POI) while the other one recruits an E3 ubiquitin ligase, mostly CRBN (Cereblon) or VHL (Von Hippel-Lindau)^31,32^. The close proximity of the POI and the E3 ligase leads to the ubiquitination of the substrate, followed by proteasomal degradation^33^. Since RNF4 is a small protein with a predicted very long unstructured N-terminal region and only a small structured domain at the C-terminus, it is challenging to develop ligands targeting selectively RNF4^34^. However, the usage of covalent binders may offer a strategy to target exposed cysteine residues to reactive ligands^35^. Recently a cysteine-reactive compound (CCW16) has been identified by activity-based profiling^36^. Linking CCW16 to JQ1, a BET (Bromo- and extra-terminal domain) inhibitor^37^, results in the development of the RNF4-targeting BRD4 degrader CCW28-3, presumably acting through RNF4 ubiquitin ligase activity^36^. Thus, we anticipated that by utilizing the CCW16 warhead degraders of RNF4 could be designed by linking this cysteine reactive moiety to established E3 ligase ligands or to itself (homo-PROTACs)^38^.

In this study, we demonstrated that RNF4 represents a cancer vulnerability and potential drug target in AML. Depletion of RNF4 exacerbated the sensitivity of OCI-AML2 cells towards the antileukemic drug decitabine (also known as 5-AzadC, 5-Aza-2′-deoxycytidine) and the replication stress inducing agent aphidicolin. Decitabine, a cytosine analog, is incorporated into the DNA during replication leading to the formation of a DNMT1-DPCs upon DNA methylation, whereas aphidicolin inhibits the DNA polymerase α and δ blocking the cell cycle at early S-phase^39–41^. To establish a small molecule degradation system for RNF4 depletion in AML, we used the recently published RNF4 ligand CCW16^36^. To develop heterobifunctional PROTACs recruiting the E3 ubiquitin ligases VHL or CRBN to RNF4, we used the established VHL- and CRBN-based E3 ligands and different linker moieties for proteasomal degradation of RNF4. However, a large diversity of synthesized PROTACs did not result in degradation of RNF4 in different cell lines. Comprehensive *in vitro* characterization of CCW16 revealed a covalent binding to RNF4 to the unstructured region of RNF4. However, *in vivo* interaction studies demonstrated covalent binding to a large set of diverse cysteine-containing cellular proteins including proteins of the peroxiredoxin family, thus identifying CCW16 as a non-selective covalent ligand. Interestingly, treatment of cells with CCW16 and CCW16-derived PROTACs revealed an upregulation of the ferroptotic marker heme oxygenase 1 (HMOX1)^42^ and an increase in lipid peroxidation, affecting cell viability. Based on this data, we propose that treatment with CCW16 leads to the activation of an RNF4-independent, ferroptotic cell death pathway.

## Methods

### Cell culture and transfection

HeLa (female, ATCC^®^, CCL-2™) and HEK293T (female, ATCC^®^ CRL-1573™), purchased from ATCC, and OCI-AML2 (male, Leibniz-Institut DSMZ) cells were cultured in Dulbecco’s Modified Eagle Medium (DMEM) (ThermoFisher Scientific) supplemented with 10% fetal calf serum (ThermoFisher Scientific), 100 U/ml penicillin, and 100 U/ml streptomycin (ThermoFisher Scientific) at 37 °C and 5 % CO_2_. HeLa RNF4 KO and HeLa Flp in T-Rex^TM^ RNF4 WT were generated and provided by Dr. Jan Keiten-Schmitz, OCI-AML3 Cas9 and MV4-11 Cas9 were provided by Dr. Hannah Uckelmann. HEK293T BRD4-HiBiT cells were gifted by PROMEGA to Prof. Dr. Stefan Knapp. For siRNA transfection of adherent cells, the Lipofectamine RNAiMAX reagent (ThermoFisher Scientific) was used for 72 h. Suspension cells were transfected with siRNA by electroporation by means of the Neon^TM^ Transfection System (Invitrogen) for 72 h, according to the manufactureŕs manual (settings: 1350 V, 35 ms, 1 pulse). MG-132 [20 µM], MLN4942 [500 nM] and TAK-243 [1 µM] were added to cell culture media 30 min prior to PROTAC treatment and incubated for 6 h. Ferrostatin was used with a final concentration of 10 µM and was always pre-treated 1 h before PROTAC treatment. DNA damaging compounds Cytarabine [30 nM], Decitabine [1 µM] and Aphidicolin [0.075 µg/ml] were given to the cells for 72 h.

### Cellular viability measurements

siRNA transfection in OCI-AML2 was performed as above. After 72 h of knockdown in a 6 well plate, 5,000 cells per well in 100 µl media were transferred to a white 96 well plate (ThermoFisher Scientific, 136101) and treated immediately with different antileukemic and DNA damage inducing compounds for further 72 h at 37 °C and 5 % CO_2_. Finally, cell viability was measured with the CellTiter-Glo® Luminescent Cell Viability Assay (Promega, G7570). For this, the plate was equilibrated to room temperature for around 30 min, 100 µl assay reagent was added to each well, incubated for 2 min on an orbital shaker and for 10 min without shaking at room temperature. Filtered luminescence was measured on a plate reader (SYNERGY H1, microplate reader, BioTek) and data was analyzed by GraphPad Prism 8.

Furthermore, 5,000 OCI-AML2 cells per well were seeded in 100 µl of media in a white 96 well plate (ThermoFisher Scientific, 136101) and treated with ferrostatin or control for 1 h at 37 °C and 5 % CO_2_. Afterwards, different PROTACs were added to the plate and incubated for further 6 h at 37 °C and 5 % CO_2_. Cell viability measurement was performed as above.

Compound toxicity was determined by using the CellTiter-Glo® 2.0 Cell Viability Assay (PROMEGA G9241). For this, 10 µl of HEK293T cells were seeded with 10,000 cells/well into white 384-well plates (Greiner, 784075) and the cells were allowed to attach for 24 h at 37 °C and 5 % CO_2_. After incubation, the compounds were titrated using an Echo acoustic dispenser (Labcyte) and the plate was further incubated for 24 h at 37 °C and 5 % CO_2_. For the assay, 10 µl assay reagent was added and incubated for 10 min at room temperature. Filtered luminescence was measured on a PHERAstar plate reader (BMG Labtech) and data was graphed, using GraphPad Prism 10.

### CRISPR drop out assay

Lentiviruses were produced in HEK293T cells by transfection of plasmids, containing the guideRNA and an RFP gene for the readout, with Lipofectamine^TM^ 2000 transfection reagent. Supernatant, containing the virus, was harvested after 2 days of incubation at 37 °C and 5 % CO_2_. 20,000 cells in 50 µl per well of a 96 well plate (Sarstedt AG & Co. KG. Ref: 83 3924500) of OCI-AML3 and MV4-11, stably expressing Cas9, were transduced on day 0 each with 10 µl of virus after addition of polybrene [1 µg/ml]. Spin infection was performed by centrifugation of the plate for 1.5 h at 37 °C and 1363 g. Afterwards, wells were filled up with media to 200 µl and plate was incubated at 37 °C and 5 % CO_2_. Transduction efficiency (∼30-70 %) was measured 3 days after transduction by flow cytometry (BECKMAN COULTER Model: CytoFLEX S) in a 96 well plate (Corning Incorporated (Costar) Ref: 3897), followed by measuring the viability of transduced cells after 6, 9, 13 and 15 days.

### Western blot analysis

Samples were resuspended in 6x Laemmli buffer and loaded onto an SDS-PAGE for protein separation, followed by transfer to a nitrocellulose membrane with the wetblot method. Before incubation overnight at 4 °C with primary antibody (dissolved in 5 % milk in PBS-T) of the respective target, membranes were blocked for 30 min with 5 % milk in PBS-T. Membranes were washed 4 times in PBS-T, followed by addition of secondary antibody (dissolved in 5 % milk in PBS-T) for 1 h purchased from LI-COR containing a fluorophore (IRDye^®^ 800CW/680RD goat anti-mouse or goat anti-rabbit) for detection by the Odyssey^®^ DLx Imaging system. Analysis and quantification was performed by means of the Image Studio^TM^ Software from LI-COR.

### BRD4 HiBiT degradation assay

Endogenously BRD4 HiBiT-tagged HEK293T (HEK293T BRD4-HiBiT) cells were obtained as a kind gift from Promega Corp. 6.4×10^5^ cells were seeded in 1.6 ml in a 6 well plate, transfected with siRNA as described above and incubated for 72 h at 37 °C and 5 % CO_2_. To measure degradation, 10 µl of a total concentration of 2.5×10^5^ cells/ml in DMEM medium were seeded into white small volume 384 well plates (Greiner, 784075) and allowed to settle overnight. Subsequently, the PROTACs were titrated to the seeded cells, using an Echo acoustic dispenser (Labcyte) and the plate was incubated for the indicated time at 37 °C and 5 % CO_2_. After incubation, HiBiT Lytic detection reagent was prepared by dilution of LgBiT protein (1:100) and lytic substrate (1:50) in Lytic detection buffer (Promega, N3040). For detection, 10 µl of the prepared mix was added to the treated cells and incubated for 10 min at room temperature. The readout was carried out in a PheraStar FSX plate reader (BMG Labtech) using the LUM plus optical module. Degradation data were then plotted with GraphPad Prism 8 software using a normalized 3-parameter curve fit with the following equation: Y = 100/(1+10^(X-LogIC50)^).

### Streptavidin pulldown

HeLa WT cells were seeded in a 10 cm dish and harvested after 48 h in PBS, lysed in 300 µl RIPA buffer per sample (50 mM Tris-HCl pH 7.5, 0.1 % SDS, 0.5 % Sodium deoxycholate, 1mM EDTA, 150 mM NaCl, addition of protease inhibitors freshly) and incubated for 10 min on ice. Cells were sonicated at 40 % with a 1 sec pulse ON and 2 sec pulse OFF for 10 sec to fully shear chromatin and chromatin-associated proteins. To clear the lysate, cells were centrifuged for 10 min at 16,000 g at 4 °C, followed by performing a Lowry protein assay of the supernatant for determination of the protein concentration. 50 µg of protein lysis was used for the Input control and 1 mg was used for the pulldown, where biotin-CCW16 and biotin, respectively were added in a final concentration of 10 µM and incubated for 2 h while rotating at 4 °C. Afterwards, lysates were incubated with Streptavidin beads (ThermoFisher Scientific) overnight at 4 °C. After washing once with RIPA buffer, 3 times with 1 % (v/v) SDS, 8 M Urea in PBS, once with 1 % (v/v) SDS in PBS and 2 times with PBS, pulled down proteins were either eluted by 2x Laemmli, including 100 µM biotin, for western blot analysis or on-bead digested for mass spectrometry evaluation (see mass spectrometry methods).

### Mass spectrometry

Whole-cell proteome (WCP) was performed in HeLa Flp in T-Rex^TM^ RNF4 WT cells, pre-treated with doxycycline (1 µg/ml) for induction of RNF4 expression for 24 h before addition of PROTACs with a final concentration of 5 µM. After 6 h of treatment, cells were washed 3 times with warm PBS and scraped in lysis buffer (2 % SDS, 50 mM Tris/HCl pH 8.5, 10 mM TCEP, 40 mM CAA, 1 mM PMSF, 1x cOmplete™, Mini, EDTA-free Protease Inhibitor Cocktail), followed by boiling at 95 °C for 10 min, sonication for 5 min and again boiling for 5 min at 95 °C. All following steps were performed in low-binding tubes (Eppendorf). To precipitate the proteins, the methanol/chloroform method was used. Initially, 4 sample volumes of ice-cold methanol, 1 sample volume of chloroform and 3 sample volumes of water were added and mixed vigorously between every step. Afterwards, the samples were centrifuged (20,000 g, 15 min, 4 °C) and the top layer was removed without disturbing the interphase containing the proteins. After addition of 3 sample volumes of ice-cold methanol, mixing and centrifugation (20,000 g, 10 min, 4 °C), the supernatant was removed and the pellet washed with 3 sample volumes of ice-cold methanol, mixed and centrifuged (20,000 g, 10 min, 4 °C), followed by air-drying for around 10 min. The pellet was dissolved in 100 µl digestion buffer (8 M urea, 50 mM Tris/HCl, pH 8.5), mixed vigorously and incubated for 10 min at room temperature to determine the protein concentration using the BCA assay (ThermoFisher Scientific). After dilution of the samples to a 2 M Urea concentration, the assay was performed according to the manufactureŕs manual. 50 µg of protein was digested overnight at 37 °C using Lys-C (10 µl/1 mg protein) and trypsin (20 µl/1 mg protein) and stopped on the next day by addition of 1 % (v/v) trifluoroacetic acid (TFA). For desalting the samples, the tC18 Sep-Pak SPE cartridges (Waters) were used. After drying the samples by speed-vac at 60 °C, peptides were resuspended in 100 mM EPPS, 10 % acetonitrile (ACN), pH 8.2. MicroBCA assay (ThermoFisher Scientific) was performed according to the manufactureŕs manual to determine peptide concentrations. 10 µg of peptides were labelled with the respective TMTpro^TM^ reagent (ThermoFisher Scientific) in 10 % ACN for 1 h at room temperature. Before pooling the samples, a labeling incorporation test (1/20^th^ of each sample, label efficiency >98 %) was performed to determine equal labelling. Addition of hydroxylamine (final concentration 0.5 %) and incubating for 15 min at room temperature quenched the labelling. According to the determined labelling intensities, labelled peptides were pooled to achieve the same labelling intensity for every sample. Using the tC18 Sep-Pak SPE cartridges (Waters), samples were desalted and were removed from remaining free TMTpro^TM^ reagent. Peptide fractionation was performed by high-pH liquid-chromatography on a micro-flow HPLC (Dionex U3000 RSLC, ThermoFisher Scientific). Tryptic peptides were analyzed on an Orbitrap Lumos coupled to an easy nLC 1200 (ThermoFisher Scientific) using a 35 cm long, 75µm ID fused-silica column packed in house with 1.9 µm C18 particles (Reprosil pur, Dr. Maisch), and kept at 50°C using an integrated column oven (Sonation). HPLC solvents consisted of 0.1 % Formic acid in water (Buffer A) and 0.1 % Formic acid, 80% acetonitrile in water (Buffer B). Assuming equal amounts in each fraction, 400 ng of peptides were eluted by a non-linear gradient from 7 to 40% B over 90 minutes followed by a step-wise increase to 90 % B in 6 minutes which was held for another 9 minutes. A synchronous precursor selection (SPS) multi-notch MS3 method was used in order to minimize ratio compression as previously described ^43^. Full scan MS spectra (350-1400 m/z) were acquired with a resolution of 120,000 at m/z 200, maximum injection time of 100 ms and AGC target value of 4 x 10^5^. The most intense precursors with a charge state between 2 and 6 per full scan were selected for fragmentation (“Top Speed” with a cycle time of 1.5 sec) and isolated with a quadrupole isolation window of 0.7 Th. MS2 scans were performed in the Ion trap (Turbo) using a maximum injection time of 50ms, AGC target value of 1.5 x 10^4^ and fragmented using CID with a normalized collision energy (NCE) of 35%. SPS-MS3 scans for quantification were performed on the 10 most intense MS2 fragment ions with an isolation window of 0.7 Th (MS) and 2 m/z (MS2). Ions were fragmented using HCD with an NCE of 50% and analyzed in the Orbitrap with a resolution of 50,000 at m/z 200, scan range of 100-500 m/z, AGC target value of 1.5 x10^5^ and a maximum injection time of 86ms. Repeated sequencing of already acquired precursors was limited by setting a dynamic exclusion of 60 seconds and 7 ppm and advanced peak determination was deactivated. All spectra were acquired in centroid mode.

Raw data was analyzed with Proteome Discoverer 2.4 (ThermoFisher Scientific). Acquired MS2-spectra were searched against the human reference proteome (Taxonomy ID 9606) downloaded from UniProt (12-March-2020; “One Sequence Per Gene”, 20531 sequences) and a collection of common contaminants (253 entries) using SequestHT, allowing a precursor mass tolerance of 7 ppm and a fragment mass tolerance of 0.5 Da after recalibration of mass errors using the Spectra RC-node applying default settings. In addition to standard dynamic (Oxidation on methionines and Met-loss at protein N-termini) and static (Carbamidomethylation on cysteines) modifications, TMT-labelling of N-termini and lysines were set as static modifications. False discovery rates were controlled using Percolator (< 1 % FDR on PSM level). Only PSMs with a signal-to-noise above 10 and a co-isolation below 50 % were used for protein quantification after total intensity normalization. Only high confident proteins (combined q-value <0.01) were used for downstream analyses. MS data were deposited on PRIDE (Whole cell proteome of HeLa Flp in T-Rex^TM^ RNF4 WT treated with either CCW16, 1a or 2c, PXD056951).

Biotin-CCW16 interactors were enriched by Streptavidin pulldown as described above. Each pulldown was performed in triplicates and in low-binding tubes (Eppendorf) and 1 mg of protein was used per pulldown. For on-bead digestion, beads were again washed twice with 50 mM Tris/HCl, pH 8.5 and resuspended in sodium deoxycholate (SDC) buffer (2 % SDC, 4 mM CAA, 1 mM TCEP in 50 mM Tris/HCl, pH 8.5) and incubated for 10 min at 95 °C for alkylation and reduction of the proteins. After cooling down to room temperature, digestion buffer was added (Tris/HCl pH 8.5) including 0.5 µg trypsin per sample and incubated overnight at 37 °C. The digestion was stopped by adding 1 % (v/v) TFA in isopropanol and peptides were loaded onto styrene-divinyl benzene reverse phasesulfonate (SDB-RPS) polymer sorbent solid phase extraction STAGE tips for clean-up^44^. Peptides were washed once with 1 % TFA (v/v) in isopropanol, followed by washing with 2 % ACN and 0.2 % TFA and elution in 80 % ACN/1.25 % ammonia. Peptides were dried by speed-vac at 60 °C and resuspended in 2 % ACN and 0.1 % TFA. Samples were analyzed on a Q Exactive HF coupled to an easy nLC 1200 (ThermoFisher Scientific) using a 35 cm long, 75µm ID fused-silica column packed in house with 1.9 µm C18 particles (Reprosil pur, Dr. Maisch), and kept at 50°C using an integrated column oven (Sonation). Peptides were eluted by a non-linear gradient from 4-28% acetonitrile over 45 minutes and directly sprayed into the mass-spectrometer equipped with a nanoFlex ion source (ThermoFisher Scientific). Full scan MS spectra (300-1650 m/z) were acquired in Profile mode at a resolution of 60,000 at m/z 200, a maximum injection time of 20 ms and an AGC target value of 3 x 10^6^ charges. Up to 10 most intense peptides per full scan were isolated using a 1.4 Th window and fragmented using higher energy collisional dissociation (normalized collision energy of 27). MS/MS spectra were acquired in centroid mode with a resolution of 30,000, a maximum injection time of 54 ms and an AGC target value of 1 x 10^5^. Single charged ions, ions with a charge state above 5 and ions with unassigned charge states were not considered for fragmentation and dynamic exclusion was set to 20s.

Raw data was analyzed with Proteome Discoverer 2.4 (ThermoFisher Scientific). Acquired MS2-spectra were searched against the human reference proteome (Taxonomy ID 9606) downloaded from UniProt (17-April-2022; “HoSP_OSPG_20220417.fasta”, 20509 sequences) and a collection of common contaminants (253 entries) using SequestHT, allowing a precursor mass tolerance of 10 ppm and a fragment mass tolerance of 0.02 Da after recalibration of mass errors using the Spectra RC-node applying default settings. Standard dynamic (Oxidation on methionines and acetylation at protein N-termini) and static (Carbamidomethylation on cysteines) modifications were set as modifications. False discovery rates were controlled using Percolator (< 1% FDR on PSM level). Only high confident proteins (combined q-value <0.01) were used for downstream analyses. MS data were deposited on PRIDE (Streptavidin biotin-CCW16 pulldown to determine CCW16-modified proteins in HeLa lysate, PXD056954).

For identification of CCW16-bound cysteine residues on RNF4, an *in vitro* interaction assay was performed as described below in triplicates. Before separating the sample using SDS-PAGE, 40 mM CAA was added to the SDS-sample buffer and boiled for 10 min to reduce and alkylate proteins. After separation by SDS-PAGE, samples were stained with InstantBlue and bands were cut into small pieces (1-2 mm³). Gel pieces were transferred to low-binding tubes (Eppendorf) and washed 3 times for 10 min each with 50 mM ammonium bicarbonate (ABC)/40 % ACN. After dehydrating gel pieces for 10 min at 37 °C with 100 % ACN and drying for 30 min, proteins were digested with 1 µg trypsin (in 50 mM ABC) per sample for 30 min at 4 °C. Remaining trypsin was removed and 50 mM ABC was added and incubated on a shaker at 500 rpm over night at 37 °C. For peptide extraction the samples were cooled down to room temperature and 50 % ACN/0.5 % TFA was added and mixed for 30 min. Afterwards, the supernatant was transferred to a new low-binding tube (Eppendorf). 50 % isopropanol/0.5 % TFA was added to the gel pieces and mixed again for 30 min. The supernatant was pooled with the supernatant of the step before. Finally, 1 % TFA in isopropanol was incubated for 10 min with the gel pieces and supernatant was transferred to the remaining supernatant. Peptides were washed once with 1 % TFA (v/v) in isopropanol, followed by washing with 2 % ACN and 0.2 % TFA and elution in 80 % ACN/1.25 % ammonia. Peptides were loaded onto styrene-divinyl benzene reverse phasesulfonate (SDB-RPS) polymer sorbent solid phase extraction STAGE tips for clean-up^44^. Peptides were dried by speed-vac at 60 °C and resuspended in 2 % ACN, 0.1 % TFA. Samples were analyzed on a Q Exactive HF coupled to an easy nLC 1200 (ThermoFisher Scientific) using a 35 cm long, 75µm ID fused-silica column packed in house with 1.9 µm C18 particles (Reprosil pur, Dr. Maisch), and kept at 50 °C using an integrated column oven (Sonation). Peptides were eluted by a non-linear gradient from 4-28% acetonitrile over 45 minutes and directly sprayed into the mass-spectrometer equipped with a nanoFlex ion source (ThermoFisher Scientific). Full scan MS spectra (300-1650 m/z) were acquired in Profile mode at a resolution of 60,000 at m/z 200, a maximum injection time of 20 ms and an AGC target value of 3 x 10^6^ charges. Up to 10 most intense peptides per full scan were isolated using a 1.4 Th window and fragmented using higher energy collisional dissociation (normalized collision energy of 27). MS/MS spectra were acquired in centroid mode with a resolution of 30,000, a maximum injection time of 54 ms and an AGC target value of 1 x 10^5^. Single charged ions, ions with a charge state above 5 and ions with unassigned charge states were not considered for fragmentation and dynamic exclusion was set to 20s.

MS raw data processing was performed with MaxQuant v1.6.17.0^45^. Acquired spectra were searched against the sequence of RNF4, the *E.coli* (strain K12) reference proteome (Taxonomy ID 83333) downloaded from UniProt (17-04-2022; 4401 sequences) and a collection of common contaminants (244 entries) using the Andromeda search engine integrated in MaxQuant ^46^. Oxidation on methionines and acetylation at protein N-termini as well as carbamidomethylation, CCW16 (+345.13649 Da), and 212 (+928.41933 Da) on cysteines were considered as variable modifications during the search. Identifications were filtered to obtain false discovery rates (FDR) below 1% for modification sites, peptide spectrum matches (PSM; minimum length of 7 amino acids) and proteins using a target-decoy strategy^47^.

### Lipid peroxidation assay

750,000 OCI-AML2 cells were seeded one day before treatment in a 6 well plate. On the next day, the PROTAC [2.5 µM] or the positive control cumene hydroperoxide [100 µM] was added to the cell culture media for 2.5 h and 2 h, respectively. 30 min before imaging, the cells were incubated with the Image-iT^®^ lipid sensor C11-BODIPY^581/591^ [10 µM] and washed afterwards 3 times in ice-cold PBS to visualize lipid peroxidation by flow cytometry. The sensor can incorporate into the membrane of living cells and shifts its fluorescence from red to green during oxidation of lipids through lipid hydroperoxides. By calculating the ratio of the reduced lipids (mean red fluorescence intensity) by the oxidized lipids (mean green fluorescence intensity) the oxidation of lipids can be determined. For imaging the BD FACSymphony^TM^ A5 Cell Analyzer (BD Biosciences) was used. The mean fluorescence intensities were calculated using the FlowJo^TM^ (FlowJo^TM^, LLC) Software for flow cytometry analysis.

### Protein expression and purification from Rosetta *E.coli*

pGEX4-T1 expression plasmids were transformed in Rosetta *E. coli* cells and GST-tagged proteins were expressed by addition of IPTG (0.5 mM) overnight at 16 °C. Cells were centrifuged and cell pellets were resuspended in lysis buffer (PBS, 1 % (v/v) Triton X-100, 1 mM DTT, 1 mM PMSF, 25 ml lysis buffer per 500 ml bacteria culture). For complete cell lysis, cells were frozen in liquid nitrogen and thawed at 37 °C three times, followed by sonification. Afterwards, cell lyses was cleared from cell debris by centrifugation (20,000 g for 30 min at 4 °C). The supernatant was incubated with Protino® Glutathione Agarose 4B beads for 2 h at 4 °C while rotating. Afterwards, the beads were washed three times with ice-cold washing buffer (=lysis buffer) and used directly for *in vitro* interaction studies or stored at -20 °C with 20 % of glycerol.

### *in vitro* interaction assay

GST-based purified proteins bound to GSH beads were incubated either with biotin-CCW16 [5 µM] (Western blot analysis) or CCW16/**2a** [10 µM] (Mass spectrometry analysis) overnight at 4 °C while rotating. Afterwards, beads were washed four times with the TPA washing buffer (30 mM TRIS pH 7.5, 100 mM NaCl, 5 mM MgCl_2_, 2 mM DTT, 0.1 mg/ml BSA, 10 % glycerol, 0.01 % (v/v) NP-40) and beads were either resuspended in 30 µl of 2x Laemmli for Western blot analysis or further proceed for mass spectrometry analysis (see mass spectrometry methods).

### NanoBRET^TM^ Target engagement assay

The assay was performed as described previously^48^. In brief, full-length VHL and CRBN were obtained as plasmids cloned in frame with an N-terminal NanoLuc-fusion (kind gift from Promega). For CRBN, DDB1 was co-expressed as additional untagged protein. The plasmids were transfected into HEK293T cells using FuGENE 4K (Promega, E5911), and proteins were allowed to express for 20 h. Serially diluted inhibitor and NanoBRET^TM^ VHL and CRBN Tracer (Promega, TracerDB IDs: T000018 (CRBN) and T000019 (VHL)) at the Tracer *K*_D_ concentration taken from TracerDB (tracerdb.org)^49^ were pipetted into white 384-well plates (Greiner 781207) using an Echo acoustic dispenser (Labcyte). The corresponding protein-transfected cells were added and reseeded at a density of 2×10^5^ cells/mL after trypsinization and resuspending in Opti-MEM without phenol red (Life Technologies). The system was allowed to equilibrate for 3 h at 37 °C and 5 % CO_2_ prior to bioluminescence resonance energy transfer (BRET) measurements. To measure BRET, NanoBRET^TM^ NanoGlo Substrate + extracellular NanoLuc Inhibitor (Promega, N2540) was added as described in the manufacturer’s protocol, and filtered luminescence was measured on a PHERAstar plate reader (BMG Labtech) equipped with a luminescence filter pair (450 nm BP filter (donor) and 610 nm LP filter (acceptor)). Competitive displacement data were then graphed using GraphPad Prism 10 software using a normalized 3-parameter curve fit with the following equation: Y = 100/(1 + 10^(X-LogIC50)^).

### Quantification and statistical analysis

Western blots were quantified by the manufactureŕs instructions with the LI-COR Image Studio^TM^ software. All quantified Western blot experiments were performed in three independent experiments and a representative Western blot is shown here. All FACS experiments were analyzed by FlowJo^TM^ (FlowJo^TM^, LLC) and plotted in GraphPad Prism (version 8.4.2). For statistical analysis the unpaired Student’s t-test was applied for Western blot quantification, viability assays and FACS experiments.

Mass spectrometry data (whole cell proteome, Streptavidin pulldown) were statistically analyzed with the Perseus software (version 1.6.15.0). For analysis of the whole cell proteome, contaminants, low confidences and reverse entries were removed. Afterwards, all four replicates were grouped, followed by filtering for minimal 4 valid values in each row in at least one group. In addition, log_2_(x) transformation was performed for each normalized abundances and Student’s two sample t-test was applied with a Benjamini-Hochberg FDR of 0.05. For analysis of the Streptavidin pulldown, contaminants, low confidences and reverse entries were removed, followed by log_2_(x) transformation of the LFQ intensities. Afterwards, samples were grouped into 3 replicates and every row was filtered for minimal valid values in at least one group. In addition, imputation of missing values was performed by replacing missing values based on normal distribution using the default settings of Perseus. For statistical analysis, Student’s two sample t-test was applied with a Benjamini-Hochberg FDR of 0.05. To calculate significant hits, Microsoft Excel was used by using the following criteria: log_2_ ratio ≥ 0.58 & –log_10_ p value ≥ 1.3 (proteome) and log_2_ ratio ≥ 1 & –log_10_ p value ≥ 1.3 (Streptavidin pulldown).

### Bioinformatics tools

For GO Biological Process and KEGG pathway analysis of the proteome and interaction MS studies the public available ShinyGO tool (version 0.80) was used, developed by a team at South Dakota State University (SDSU) (http://bioinformatics.sdstate.edu/go/). The following settings were used: FDR cutoff (0.05), Number pathways to show (10), Pathway size: Minimum (2) and Maximum (5000). The protein coding genes of the human genome were used as background. The significant enriched pathways were selected by FDR and ordered by fold enrichment.

### PROTAC and compound synthesis

All commercial chemicals and solvents were used without further purification. All reactions were performed in an inert atmosphere (Ar). Product purification was performed on a PuriFlash Flash Column Chromatography System from Interchim using prepacked silica or RP C18 columns.

The synthesized compounds were characterized by ^1^H NMR, ^13^C NMR, and mass spectrometry (ESI). NMR spectra were measured in DMSO*-d_6_* or CD_2_Cl_2_ on a Bruker AV300, AV500 or DPX600 spectrometer. Chemical shifts δ are reported in parts per million (ppm). Determination of the compound purity and mass by HPLC was carried out on an Agilent 1260 Infinity II device with a 1260 DAD HS detector (G7117C; 254 nm, 280 nm, 320 nm) and a LC/MSD device (G6125B). The compounds were analyzed on a Poroshell 120 EC-C18 (Agilent, 3 x 150 mm, 2.7 µm) reversed phase column using 0.1% formic acid in water (A) and 0.1% formic acid in acetonitrile (B) as a mobile phase. The following gradient was used:

Method A (ESI pos. 100-1000): 0 min. 5 % B - 2 min. 5 % B - 7 min. 98 % B (flow rate of 0.5 mL/min.). Method B (ESI pos. 100-1300): 0 min. 5 % B - 2 min. 5 % B - 7 min. 98 % B (flow rate of 0.5 ml/min.).

UV-detection was performed at 254, 320 nm and all compounds used for further biological characterizations showed >95% purity.

High resolution (Orbitrap) measurements were executed on a Exploris 480 Thermo (Bremen, Germany) mass spectrometer equipped with a heated electrospray source (HESI) and coupled to a liquid chromatography system Vanquish VF-P10-A binary pump, VF-A10-A auto sampler which was set to 10 °C and which was equipped with a 25 µl injection syringe and a 100 µl sample loop. Instead of a column a 0.18 mm, 600 mm length capillary was installed within the column compartment VH-C10-A. For automated direct infusion 2.0 µL sample was injected using the flow gradient described in the Table below using 90% pure acetonitrile and 10% water with 0.1% formic acid. The flow was switched according to the Table below from waste to the MS and back to the waste, to prevent source contamination. For monitoring two full scan modes were selected with the following parameters. Polarity: positive; scan range: 100 to 1500 *m*/*z*; resolution: 480,000; AGC target: “Standard”; maximum IT: “Auto”. General settings: sheath gas flow rate: 20; auxiliary gas flow rate 5; sweep gas flow rate: 1; spray voltage: 3.5 kV; capillary temperature: 325 °C; S-lens RF level: 50; auxiliary gas heater temperature: 125 °C. For negative mode, all values were kept instead of the spray voltage which was set to 2.5 kV.

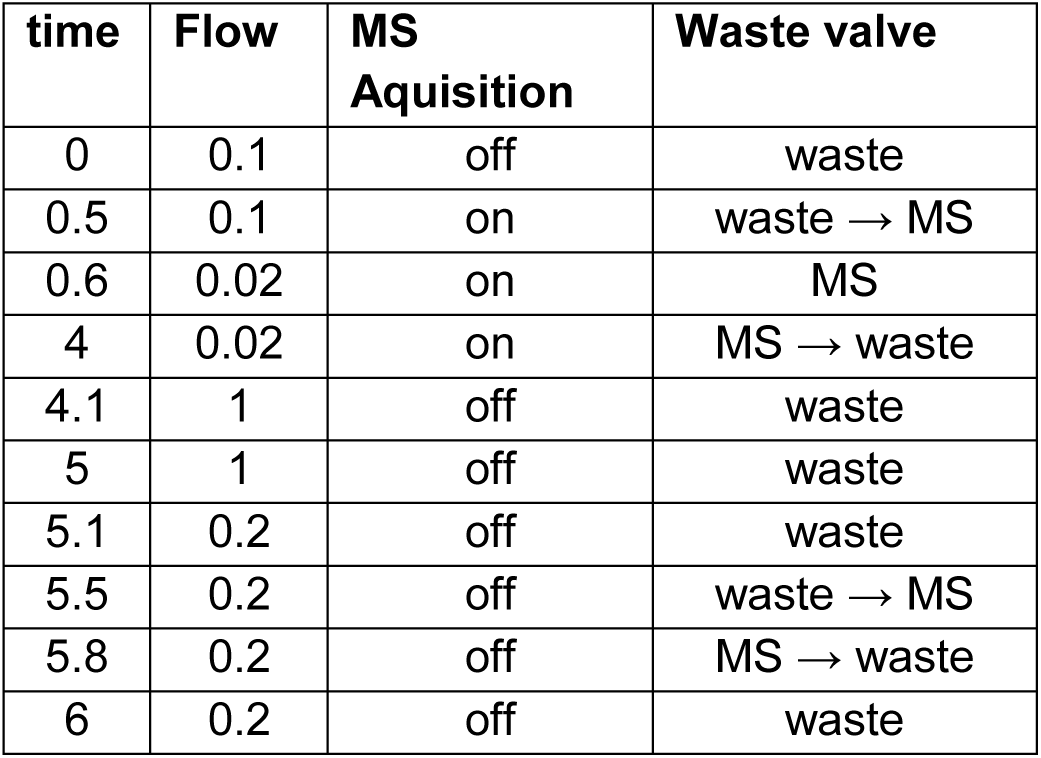

Synthetic procedures and analytical data of all synthesized compounds are provided in the supplemental data file (**Supplementary data 1**).

## Results

### RNF4 represents a vulnerability in AML cells

SUMO signaling is upregulated in many tumor entities, including hematological malignancies^50^. Mining of TGCA expression datasets revealed an upregulation of RNF4 in leukemic cells of AML patients (**Figure 1A**). Intriguingly, high expression of RNF4 correlated with poor survival of AML patients, thus defining RNF4 as a potential drug target (**Figure 1B**). To test this hypothesis, we investigated the dependence of AML cell lines on RNF4 using various cell-based phenotypic assays.

**Figure 1:**
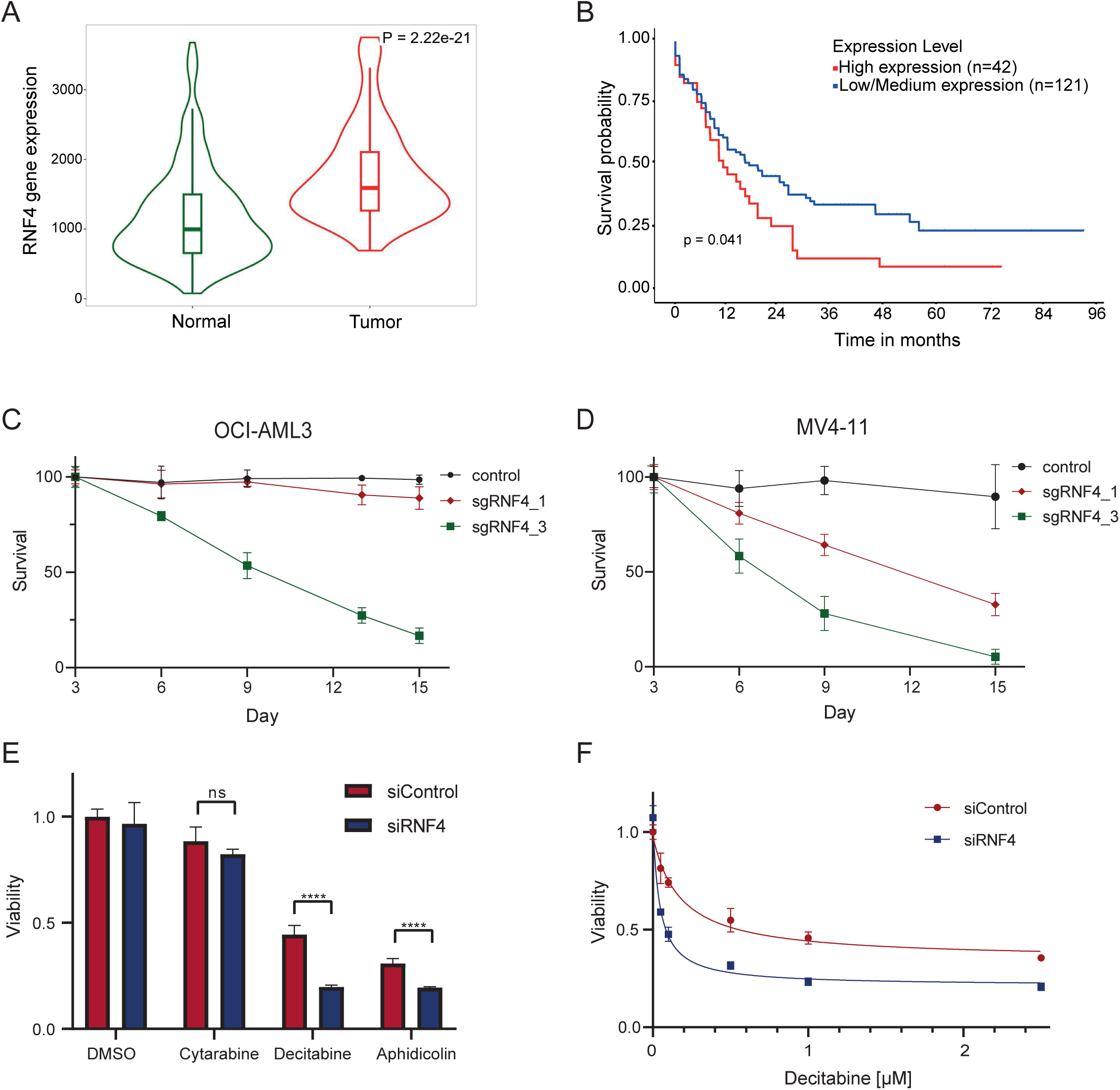
RNF4 as a vulnerability of AML cells. (A) AML cells exhibit an elevated expression of RNF4 (derived from TNMplot.com; based on TGCA data). (B) Effect of RNF4 expression level on AML patient survival. High expression of RNF4 is associated with a poor prognosis (analyzed using UALCAN). (C)/(D) RNF4 CRISPR drop out experiment in Cas9 expressing OCI-AML3 (C) or MV4-11 (D) cells. Transduction efficiency of PE positive cells was around 50 %. Survival rate was measured by flow cytometry and normalized to empty vector control on day 3. (E) Cell viability assay of OCI-AML2 cells 6 days after RNF4 KD with siRNA in combination with cytarabine [50 nM], decitabine (5-azadC) [1μM] or aphidicolin [0.075 μg/ml] (F) Viability assay of OCI-AML2 cells after RNF4 KD with siRNA in combination with different decitabine concentrations to measure dose-response dependency. IC_50_ value was determined by nonlinear dose-response – inhibition fit.

First, we performed a CRISPR/Cas9 dropout experiment in Cas9 expressing OCI-AML3 (**Figure 1C**) and MV4-11 (**Figure 1D**) cell lines and measured the cell survival over 15 days after lentiviral transduction with 3 different sgRNAs targeting RNF4 or a non-targeting control. We observed a drastic decrease in cell viability especially after transduction with sgRNF4_3 in both cell lines, whereas transduction with sgRNF4_1 affected viability only in MV4-11 cells (**Figure 1C, D**). In contrast, sgRNF4_2 did not have an effect on the viability of both cell lines tested (data not shown). Immunoblot analysis of RNF4 levels in OCI-AML3 Cas9 and MV4-11 Cas9 demonstrated that the reduced viability correlated with knockout (KO) efficiency. Transduction with sgRNF4_1 and sgRNF4_3 led to a reduction in RNF4 levels and the detection of a truncated form of RNF4 (**Supplementary Figure S1A, B**). This truncated form was not expected to be functional since RNF4 activity relies on the C-terminal catalytic RING domain^51^. However, sgRNF4_2 did not significantly reduce RNF4 levels, explaining why it did not affect viability in the CRISPR drop out experiments (**Supplementary Figure S1A, B**).

### RNF4 exerts key functions in genome maintenance by mediating tolerance to replication stress or clearance of DNA-protein crosslinks

To investigate if RNF4 depletion sensitizes AML cells to genotoxic stress, we performed an siRNA-mediated knockdown (KD) of RNF4 in OCI-AML2 cells (**Supplementary Figure 1C**), followed by treatment with the DNA damage compounds cytarabine, decitabine and aphidicolin (**Figure 1E**). We used the CellTiter-Glo® Luminescent Cell Viability Assay (CTG) to measure cell viability. In contrast to the CRISPR drop out experiment, RNF4 KD alone or in combination with cytarabine did not significantly affect cell viability. This observation can either be explained by the shorter time-course (3 days), transient transfection of siRNA or less efficient RNF4 depletion by siRNA-mediated KD when compared to CRISPR/Cas9-mediated KO. However, RNF4 KD in combination with the DNA damage compounds decitabine and aphidicolin showed synthetic lethality (**Figure 1E**). The increased sensitivity of RNF4 depleted cells to aphidicolin, which induces replication stress, is consistent with the established function of RNF4 in counteracting replicative stress. Since decitabine treatment showed a strong decrease in cell viability after RNF4 KD compared to RNF4 KD alone, we treated OCI-AML2 with different concentrations of decitabine in combination with RNF4 KD. We detected a dose-response dependency with an EC_50_ value of 0.2 µM for decitabine treatment alone and an EC_50_ value of 0.04 µM for combination with RNF4 KD (**Figure 1F**) demonstrating that targeting RNF4 sensitizes cells to the antileukemic drug decitabine. Decitabine induces DNMT1-DPCs, leading to DNMT1 degradation and DNA hypomethylation. Since RNF4 is engaged in clearance of DNMT1-DPCs, its depletion likely explains the sensitization of the cells to decitabine^52^.

### Design, synthesis and validation of CCW16-based RNF4 degraders

Based on the above findings, we aimed to develop a PROTAC targeting RNF4 for degradation. PROTACs are heterobifunctional molecules that recruit a protein of interest to the ubiquitin system. The Nomura group recently identified a covalent RNF4 binder, CCW16, which was used to design PROTACs in which RNF4 acted as the recruited E3 ligase^36^. We hypothesized that CCW16 could also be used as a ligand to design PROTACs that lead to RNF4 degradation by targeting E3 ligases such as CRBN and VHL that have been established for the design of PROTACs. We therefore functionalized CCW16 as a POI recruitment ligand and designed a series of PROTACs, where CCW16 was linked to VHL032- and 4-hydroxy thalidomide-derived-E3 ligands, recruiting the E3 ligases VHL and CRBN, respectively. We synthesized four different CRBN-based (**1a-d**) and three different VHL-based PROTACs (**2a-c**) using a variety of polyethylene glycol (PEG) and alkyl linkers (**Figure 2A**). The synthetic procedures and chemical characterization of all synthesized PROTACs are included in the supplemental information (**Supplementary data 1**).

**Figure 2:**
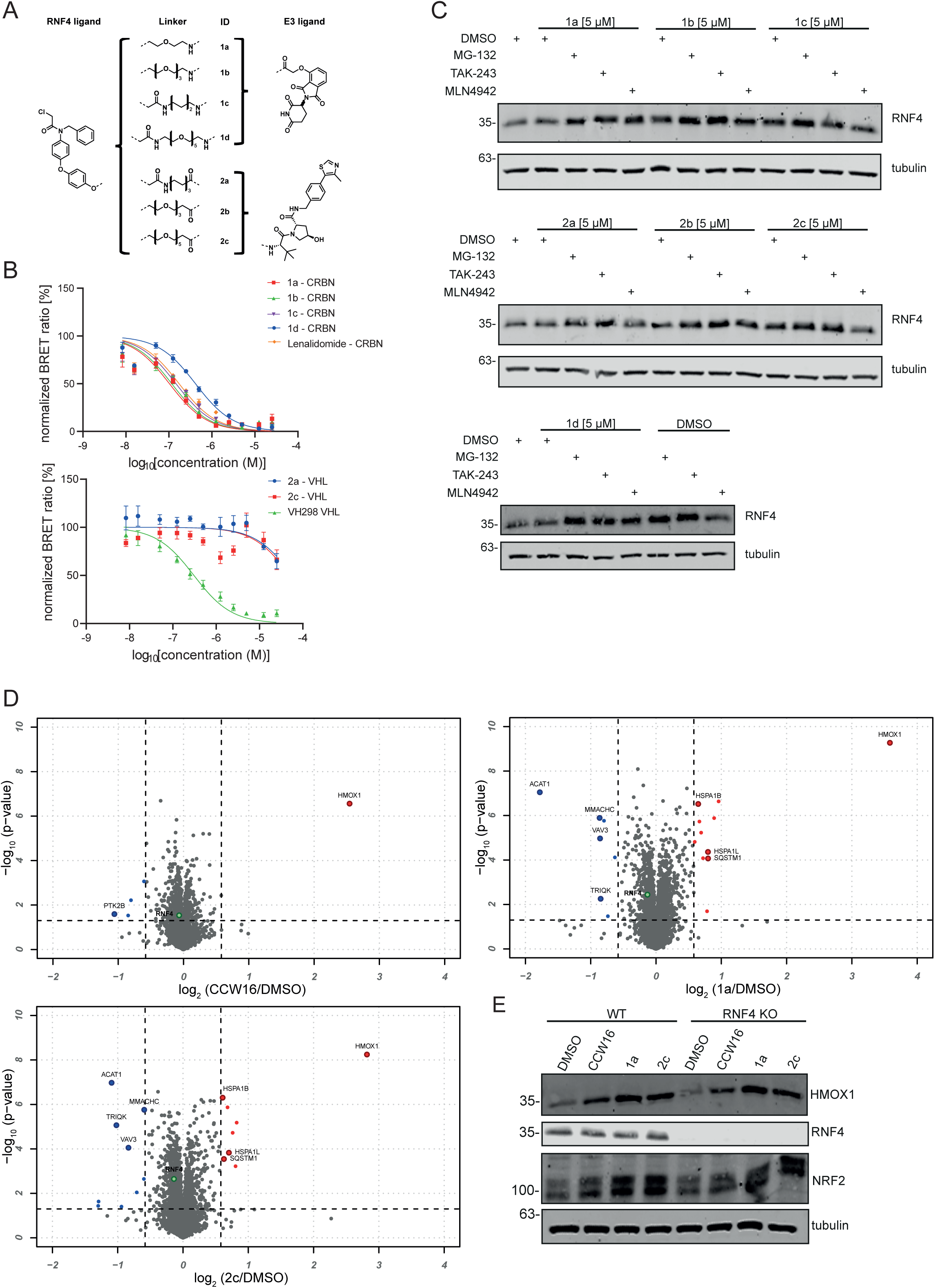
Design and evaluation of RNF4 targeting PROTACs. (A) The RNF4 binder CCW16 (left panel) was utilized for PROTAC development using diverse linkers (central panel) as well as VHL and CRBN E3 ligands (right panel). (B) NanoBRET assay for CRBN and VHL E3 ligand based RNF4 PROTACs to determine cell membrane permeability. NanoBRET assays in intact cells transiently expressing CRBN (upper panel) or VHL (lower panel) with an N-terminally tagged NanoLuc to investigate cell membrane permeability of PROTACs. (C) Treatment of HeLa WT cells with different RNF4 targeting PROTACs and evaluation of RNF4 degradation level by immunoblotting. Cells were treated 30 min before PROTAC treatment [5 µM] with MG-132 [20 µM], TAK-243 [1 µM] or MLN-4942 [500 nM] and harvested after 6 hours. Control cells were treated with DMSO. Tubulin was used as loading control. (D) Whole cell proteome analysis by mass-spectrometry. HeLa cells expressing RNF4 from a doxycycline-inducible promoter were treated either with CCW16 [5 µM] alone (left upper panel), PROTAC 1a [5 µM] (right upper panel) or 2c [5 µM] (lower left panel) for 6 hours. Results of the TMT-based MS analysis are visualized in a volcano plot comparing PROTAC treatment vs. DMSO control. Hits considered as significant are highlighted in red (log2 ratio ≥ 0.58, –log10 p value ≥ 1.3). The identification of those candidates was based on two-sided Student’s t-test analysis comparing the normalized TMT abundances of CCW16/1a/2c treatment with DMSO control treatment. Experiments were performed with four replicates. (E) Validation of proteomic results by immunoblotting in HeLa WT and HeLa RNF4 KO cells. Same treatment procedure as in (D). Tubulin was used as loading control.

To ensure cell penetration of the synthesized PROTACs, we performed NanoBRET^TM^ (Bioluminescence Resonance Energy Transfer) assays (**Figure 2B**). NanoLuciferase was ectopically expressed in HEK293T cells in frame with CRBN and VHL, respectively. As expected, the synthesized compounds demonstrated target engagement with CRBN and VHL in intact cells with EC_50_ values in the nanomolar and micromolar range, respectively. The measured EC_50_ were similar to the positive control lenalidomide for CRBN-based PROTACs demonstrating excellent cell penetration. However, EC_50_ values were in the micromolar range for VHL-based PROTACs which was significantly weaker compared with the VHL inhibitor VH298 possibly due to less efficient cell penetration of these PROTACs (**Supplementary Figure 2A**).

To evaluate the potency and degradation efficiency of the candidate PROTACs, we performed Western blot analysis detecting endogenous RNF4. Initially, HeLa cells were treated for 6 h at a PROTAC concentration of 5 µM. To validate the dependence of possible degradation on the ubiquitin-proteasome system (UPS), control samples were pre-treated with the proteasomal inhibitor MG-132, the ubiquitin activating enzyme (UAE) inhibitor TAK-243 or with the neddylation inhibitor MLN4942 (blocking CRBN or VHL activity). While inhibition of the UPS machinery stabilized RNF4, an efficient and reproducible degradation of RNF4 was not detected for any of the CRBN- or VHL-based PROTACs (**Figure 2C)**. To rule out that the used antibody did not specifically monitor RNF4, we switched to a HeLa cell line expressing an endogenously FLAG tagged RNF4. Cells were treated for 5, 8 and 24 h with the synthesized PROTACs at concentrations between 1 µM and 5 µM (**Supplementary Figure 2B**), but no significant reduction of RNF4 was observed under any of these conditions. Since the efficiency of PROTACs can be cell-type specific, we validated our compounds in two AML cell lines, NB-4 and OCI-AML2, but again we were unable to detect any degradation of RNF4 upon exposure to the CCW16-based degraders at different concentrations and time points (**Supplementary Figure 2C, D**). Since RNF4 is an E3 ubiquitin ligase possibly ubiquitinating VHL or CRBN after PROTAC addition, we also tested if the synthesized PROTACs induced CRBN or VHL degradation. However, a reduction in CRBN or VHL signal was not detected, demonstrating that the PROTACs did not function as RNF4 recruiting molecules (**Supplementary Figure 2E**). Altogether, these data showed that both the VHL-type and the CRBN-type CCW16-based PROTACS failed to efficiently degrade RNF4.

### CCW16 and CCW16-based PROTACs induce HMOX1 expression

Based on the lack of CCW16-based protein degradation in our initial PROTAC design, we aimed to comprehensively characterize the cellular properties of CCW16 before designing the next generation of CCW16-based RNF4 degrader molecules. First, we used quantitative mass spectrometry to detect subtle differences in protein levels which may not be quantifiable by Western blotting. We performed TMT-based quantitative proteomics in cells expressing FLAG-tagged RNF4 from a doxycycline inducible promoter. Cells were either treated with the CRBN-based PROTAC **1a**, with the VHL-based PROTAC **2c** or with the RNF4 binder CCW16 alone. In accordance with our immunoblot experiments, no significant reduction of RNF4 level was detectable upon treatment of cells with the PROTACs. The most significantly downregulated protein following treatment of cells with either PROTAC 1a or 2a was mitochondrial Acetyl-CoA acetyltransferase (ACAT1), which catalyzes the last step of the mitochondrial beta-oxidation pathway. Strikingly, however, the proteomic analysis also revealed a drastic induction in heme oxygenase 1 (HMOX1) levels, which has an important function in protecting cells from oxidative damage^53^, in cells treated with the CRBN-based PROTAC **1a** or VHL-based PROTAC **2c** as well as in cells treated with CCW16 only (**Figure 2D, Supplementary Table I).** This finding was confirmed by immunoblotting using a specific HMOX1 antibody. Furthermore, anti-HMOX1 immunoblotting in RNF4 KO HeLa cells revealed that the upregulation of HMOX1 upon exposure of cells to CCW16 or the CCW16-based PROTACs was independent of RNF4 (**Figure 2E**). Interestingly, MS-data also revealed that proteins of the heat shock protein family A (Hsp70) were upregulated after treatment with **1a** and **2c** (**Figure 2D**), indicating a more general stress response after exposure with CCW16 and its PROTAC derivatives. In line with our observation of HMOX1 and Hsp70 upregulation, the transcription factor NRF2 (Nuclear Factor Erythroid-Derived 2-Related Factor 2), which controls expression of HMOX1 and other stress-responsive targets^54^, was increased after CCW16 and PROTAC treatment (**Figure 2E**). Altogether our proteomic data confirmed that CCW16 and CCW16-based PROTACs did not degrade RNF4, but induced a general RNF4 independent oxidative stress response leading to upregulation of HMOX1 and other heat shock proteins.

### CCW16 targets cysteine residues 51 and 91 of RNF4 *in vitro*

Next, we aimed to confirm binding of CCW16 to RNF4 *in vitro* using purified recombinant protein. CCW16 has been described to covalently target cysteine residues 132 and 135 of RNF4 residing within its RING domain^36^. To confirm this interaction, we first expressed GST-RNF4 WT and a GST-RNF4 C132/135S mutant in *E. coli* and purified the respective proteins to homogeneity using GSH-beads (**Figure 3A**). The purified GST-RNF4 was incubated with biotin-linked CCW16 (biotin-CCW16) using the established CCW16 exit vector for coupling the biotin moiety (**Supplementary data 1**). After stringent washing, samples were separated by SDS-PAGE and biotin-CCW16-RNF4 conjugates were detected using Streptavidin linked to a fluorophore (**Figure 3A, B**). Under these conditions, biotinylation was detected in both wild-type RNF4 and the RNF4 C132/135S variant. Importantly, binding of biotin-CCW16 to GST alone was not detected indicating that *in vitro* CCW16 bound to RNF4, but not via the proposed cysteine residues C132 and C135.

**Figure 3:**
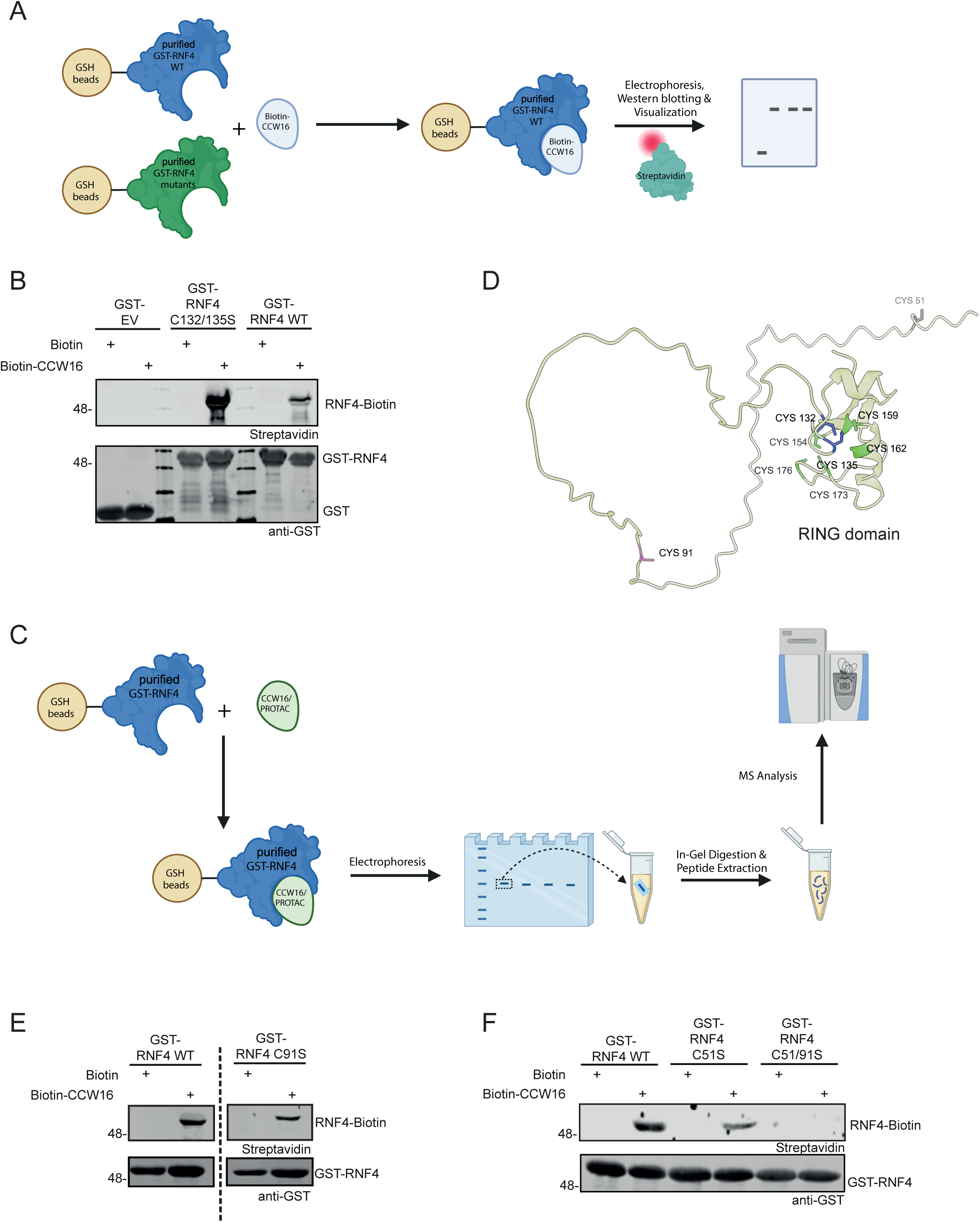
Further evaluation of RNF4-binder CCW16 *in vitro*. (A) Scheme outlining the *in vitro* interaction studies of biotin-CCW16 with GST-RNF4. Biotin-CCW16 modification of RNF4 was visualized by immunoblotting with a Streptavidin protein linked to a fluorophore. (B) Biotin or biotinylated CCW16 was incubated with wild-type GST-RNF4 or mutants with cysteine residues changed to serines as indicated. Covalent RNF4-CCW16 conjugates were detected by Streptavidin immunoblotting. (C) Experimental procedure of *in vitro* interaction studies of CCW16 or PROTAC 2a with GST-RNF4 WT to identify modified cysteine residues in RNF4 by mass spectrometry. (D) AlphaFold2 model of human RNF4, highlighting the CCW16 modified cysteine residue identified in the mass spectrometry experiment in (C) (pink), the two catalytic, but unmodified cysteine residues are shown in blue, the not detectable cysteine residue 51 is shown in grey and all remaining cysteine residues are highlighted in green. (E)/(F) Same procedure as in (B). GST-RNF4 cysteine to serine mutations are indicated. Covalent RNF4-CCW16 conjugates were detected by Streptavidin immunoblotting.

To identify all cysteine residues in RNF4 modified by CCW16, we performed a mass spectrometry mapping experiment. To this end, we incubated purified GST-RNF4 with CCW16 or the CCW16-based PROTAC **2a** and analyzed the resulting peptides by tandem mass spectrometry after trypsin digestion (**Figure 3C, Supplementary Table II**). Importantly, we identified by MS/MS a peptide in which CCW16 was conjugated to cysteine 91, as indicated by a mass shift in the peptide containing residue (**Figure 3D (pink), Supplementary Figure 3A**), but we were unable to detect conjugation to cysteine residues C132/135 or any other cysteine residue of the RING domain (**Figure 3D, blue and green, Supplementary Figure 3B**). Notably, when RNF4 was incubated with the CCW16-based PROTAC **2a**, we also observed the formation of a covalent bond between **2a** and RNF4 at cysteine C91 (**Figure 3D (pink), Supplementary Figure 3C**). While we did find the cysteines 132/135 to be modified by PROTAC **2a**, we only detected the modified peptide in one replicate with very low intensity and could not pinpoint the modification to one residue, but a doubly modified peptide was not found (**Supplementary Figure 3B**).

To validate these results, we applied the same approach as described in **Figure 3A**, but used an RNF4 variant, where C91 was mutated to a serine residue. When compared to wild-type RNF4 this variant exhibited reduced binding to CCW16, but still exhibited residual binding which suggested that CCW16 bound to additional residues (**Figure 3E**). Our mass-spectrometry analysis did not cover a long peptide within the unstructured N-terminus of RNF4, which harbors C51 (**Figure 3D, grey**). To validate whether C51 served as an additional CCW16 attachment site, we generated an RNF4 variant, in which C51 was mutated to serine as well as a double C51/91S mutant. Gratifyingly, no biotinylation was observed using the double mutant demonstrating that biotin-CCW16 bound to C51 and C91 located in the unstructured region of RNF4 (**Figure 3D, 3F**).

### BRD4 degradation via CCW28-3 does not depend on RNF4

Because of the high reactivity of CCW16 required to react with two cysteines in the unstructured protein region of RNF4, we hypothesized that the RNF4-based BRD4 degrader CCW16-JQ1 (CCW28-3) may not degrade BRD4 via an RNF4 dependent mechanism^36^. To investigate this, we compared CCW28-3 degradation in HeLa WT with degradation in HeLa RNF4 KO cells. The established CRBN ligand based PROTAC dBET6 served as a positive control. As reported^36^, BRD4 immunoblotting revealed significant degradation of the short and the long isoform of BRD4 after CCW28-3 or dBET6 treatment in wild type cells (**Figure 4A, B**). However, similar levels of degradation were also observed in RNF4 KO cells using CCW28-3 (**Figure 4A, B**), suggesting RNF4-independent degradation of BRD4 by CCW28-3. Recovery of BRD4 levels was observed by co-treatment with the proteasome inhibitor MG-132, confirming a proteasome dependent BRD4 degradation. Interestingly, however, treatment of cells with the neddylation inhibitor MLN4942, also restored BRD4 expression (**Supplementary Figure 4A**). Since MLN4942 only affects cullin-dependent RING E3 ubiquitin ligases^55^, but not RNF4, this data confirmed that CCW28-3 induced RNF4-independent degradation via a different but so far unidentified E3 ligase.

**Figure 4:**
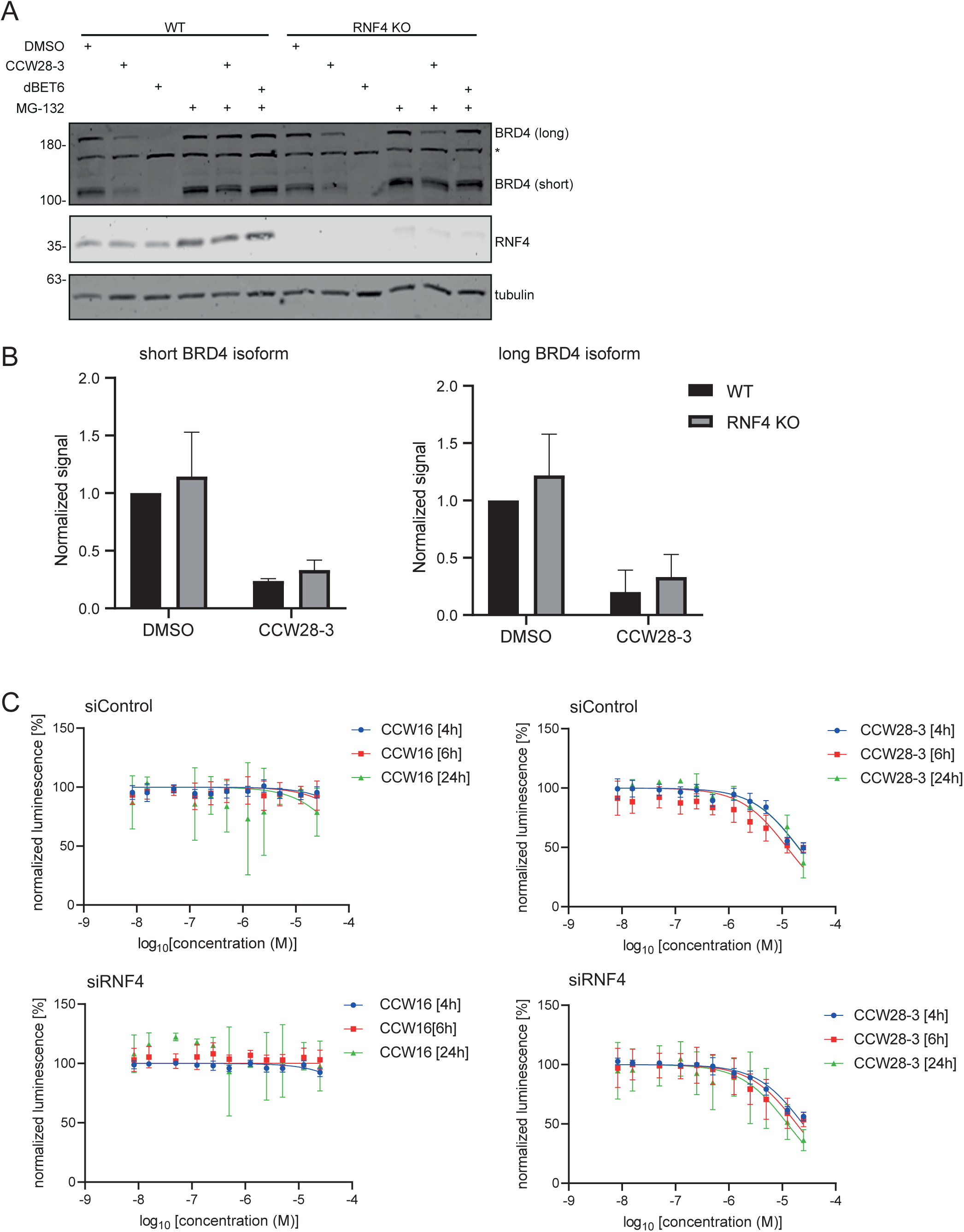
Investigation of the RNF4-dependent BRD4 degrader JQ1-CCW16 (CCW28-3). (A) Pre-treatment of HeLa WT or HeLa RNF4 KO cells with MG-132 [20 µM] 30 min before treatment with CCW28-3 [10 µM] or dBET6 [500 nM] and incubation for 6 hours. *Unspecific band (B) Quantification of BRD4 levels (short and long isoform) after CCW28-3 treatment in HeLa WT and HeLa RNF4 KO cells. Experiment was performed in triplicates. (C) Measurement of BRD4 levels based on luciferase activity. Treatment of HEK BRD4-HiBiT cells with different concentrations of CCW16 (left panel) or CCW28-3 (right panel) for 6 h. Performing of control KD (upper panel) or RNF4 KD (lower panel) 3 days before treatment. Luciferase activity was measured by addition of the large luciferase fragment (largeBiT) and substrate.

To further validate the RNF4-independent activity of CCW28-3, we performed an RNF4 KD in HEK293T cells expressing an endogenously HiBiT tagged BRD4 and treated cells with CCW28-3 or CCW16 alone for 4 h, 6 h and 24 h (**Figure 4C, D**). While CCW16 alone did not affect BRD4 levels, CCW28-3 treatment led to a decrease of HiBiT-tagged BRD4 (**Figure 4C**). However, the BRD4-HiBiT signal was not restored after siRNA-mediated depletion of RNF4 confirming RNF4 independent BRD4 degradation (**Figure 4D**). Efficient KD was confirmed by anti-RNF4 immunoblotting (**Supplementary Figure 4B**) and viability assays showed that the viability of HEK293T BRD4-HiBiT cells treated with CCW28-3 was only affected at concentrations in the high micromolar range and at treatment times longer than 9 h (**Supplementary Figure 4C**).

### CCW16 covalently targets a broad spectrum of cellular proteins

To investigate the selectivity of CCW16 in the cellular environment, cell lysates from HeLa cells were incubated with biotin-CCW16 or with biotin alone, followed by Streptavidin pulldowns and mass spectrometry analysis (**Figure 5A**). In the biotin-CCW16 pulldown around 2000 proteins were significantly enriched compared to biotin control pulldown (**Supplementary Figure 5A, Supplementary Table III**). Analysis of the mass spectrometry data and the modified peptides revealed 30 proteins, in which we identified specific cysteine residues which were modified by covalent biotin-CCW16 adducts (**Figure 5B**).

**Figure 5:**
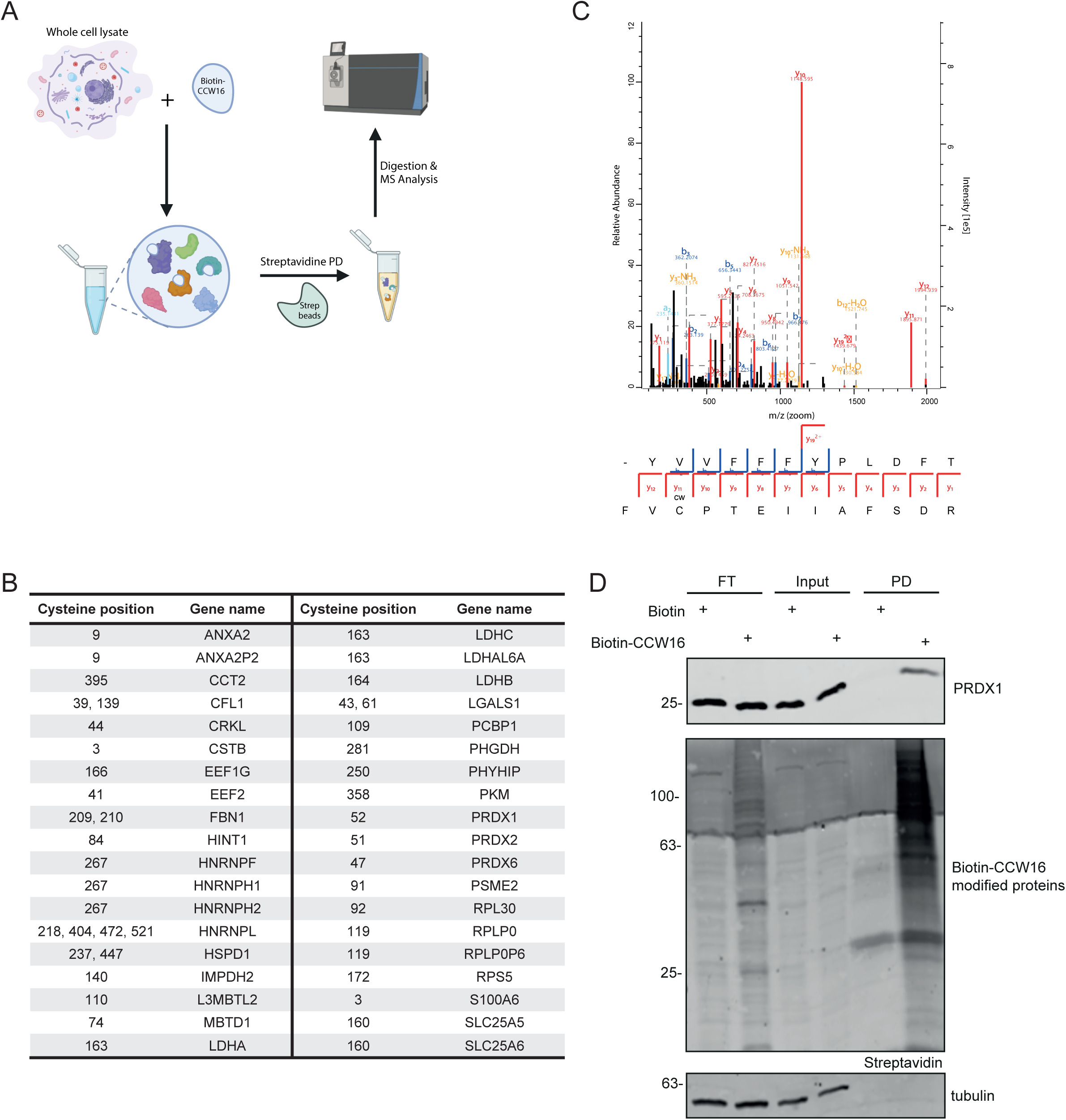
Identification of CCW16 targets *in vivo*. (A) Experimental scheme to identify *in vivo* targets of CCW16. Whole HeLa WT cell lysate was incubated with biotin [10 µM] or biotin-CCW16 [10 µM], followed by Streptavidin pulldown to enrich for biotin-CCW16 modified proteins, on bead-digestion and quantitative MS analysis. (B) All detectable proteins from the mass spectrometry experiment in (A) which were modified by biotin-CCW16. Gene names and respective modified cysteine residue are indicated. CCW16 target proteins were defined by two-sided Student’s t-test analysis comparing LFQ intensities of biotin-CCW16 pull down with the respective biotin control pulldown. Experiments were performed as triplicates. (C) MS Spectrum of biotin-CCW16 modified PRDX1. (D) Validation of Streptavidin pulldown results by immunoblotting in HeLa WT cells. Same treatment procedure as in (A). Tubulin was used as loading control.

### CCW16 covalently targets peroxiredoxins

Peroxiredoxins (PRDXs) are important for the clearance of peroxides by oxidizing a thiol group of a cysteine residue in PRDX resulting in the reduction of peroxides^56,57^. Six different PRDXs (PRDX1-6) are expressed in mammalians. PRDX1, 2 and 6 are mainly localized in the cytosol, while PRDX3 is present in the mitochondria and PRDX4 is found in the endoplasmic reticulum and is also localized in the extracellular space. PRDX5 is distributed to the cytosol, the mitochondria, the nucleus and the peroxisomes^57^. Intriguingly, three paralogs, PRDX1, PRDX2 and PRDX6 showed covalent binding to biotin-CCW16 on their catalytically active cysteine residues (**Figure 5B, C, Supplementary Figure 5A**). GOBP analysis of all biotin-CCW16 modified proteins revealed a significant enrichment of proteins involved in cell redox homeostasis (**Supplementary Figure 5B**). These data suggested that the binding of CCW16 caused inhibition of PRDX activity resulting in oxidative stress which was also supported by the observed upregulation of HMOX1 upon treatment of cells with CCW16 or CCW16-based PROTACs (**Figure 2D**). In agreement with mass spectrometry data, immunoblotting data confirmed the association of a large amount of biotin-CCW16 modified proteins compared to the biotin control after performing a biotin-CCW16 pulldown (**Figure 5D**). Furthermore, we also confirmed PRDX1 binding to biotin-CCW16 by immunoblotting, validating also our mass spectrometry results (**Figure 5D**). We also observed a very weak band for RNF4 in the pulldown fraction, questioning that RNF4 is a specific CCW16 target in cell lysates (**Supplementary Figure 5C**). This data confirms our suggestion that CCW16 binds in a non-selective manner to reactive cysteine residues via the highly reactive α-chloroamide warhead, thereby inducing redox imbalance and oxidative stress.

### CCW16 treatment induces lipid peroxidation followed by ferroptosis in AML cells

Upregulation of HMOX1 is a hallmark of the ferroptosis cell death pathway^42^. Furthermore, Kyoto Encyclopedia of Genes and Genomes (KEGG) analysis of our proteomic data revealed a significant enrichment of proteins associated with the ferroptosis pathway (**Figure 2D, Supplementary Figure 5D**). Since ferroptosis is driven by lipid peroxidation^58–61^, we investigated if our compounds led to lipid peroxidation in AML cells. We treated OCI-AML2 cells either with CCW16 alone, with the PROTAC **2c** or with cumene hydroperoxide (HP) as positive control for 2.5 h and 2 h, respectively. To analyze lipid peroxidation, cells were treated for the last 30 minutes with the fluorescent conjugated fatty acid dye C11-BODIPY581/591 which changes from red to green fluorescence upon oxidation after accumulation in membranes, and flow cytometry was performed^62^. We detected a highly significant induction of lipid peroxidation after PROTAC **2c** treatment as well as after treatment with CCW16 alone, which was comparable with peroxidation levels after HP treatment, suggesting that CCW16 and PROTAC **2c** were involved in the activation of lipid peroxidation (**Figure 6A**).

**Figure 6:**
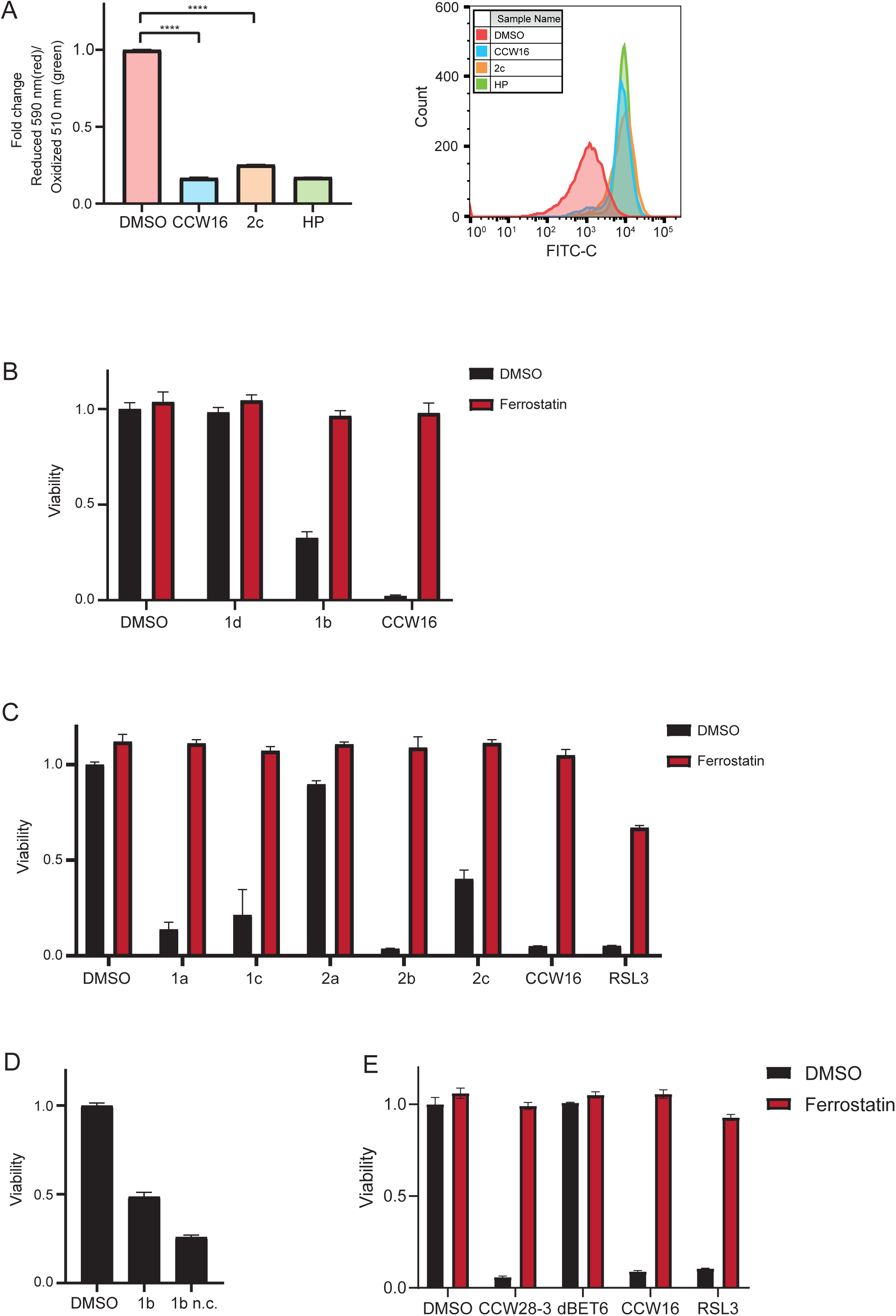
Analysis of the treatment effects of RNF4 binder CCW16 *in vivo*. (A) Treatment of OCI-AML2 cells for 2.5 h with CCW16/2c [each 2.5 µM] or for 2 h with cumene hydroperoxide [100 µM] (HP, positive control) followed by labelling with BODIPY® 581⁄591 C11 fluorophor to measure lipid peroxidation. Upon oxidation in living cells, fluorescence shifts from red (reduced) to green (oxidized). The fluorescence signal was measured by flow cytometry (events: 10,000 cells). Experiment was performed in duplicates. Ratio of reduced vs. oxidized signal is indicated. (B)/(C) Pre-treatment of OCI-AML2 cells with 10 µM of ferrostatin 1 h before treatment with either 1 µM of 1b, 1d or CCW16 (B) and 1a, 1c, 2a, 2b, 2c, CCW16 or RSL3 (positive control) (C) for 6 h and measurement of viability with the CellTiterGlo assay. (D) Treatment of OCI-AML2 with either 1 µM of PROTAC1b or the inactive counterpart 1b n.c.. Measurement of viability by CellTiterGlo. (E) Pre-treatment of OCI-AML2 with ferrostatin for 1 h [10 µM], followed by treatment with CCW28-3 [1 µM], dBET6 [500 nM], CCW16 [1 µM] or RSL3 [10 µM] for 6 h and measurement of cell viability by CellTiterGlo.

To evaluate if the lipid peroxidation resulted in the induction of ferroptosis, we pre-treated the OCI-AML2 cells with ferrostatin, a ferroptosis inhibitor^63^, followed by treatment with CCW16 alone or with all our synthesized CRBN- and VHL-based PROTACs and measured the effect on cell viability via a CTG assay (**Figure 6B, C**). The GPX4 inhibitor RSL3, a well-known ferroptosis inducer leading to a significant reactive oxygen species (ROS) accumulation, was used as positive control^64^. Analysis revealed that most of the PROTACs led to a strong reduction in cell viability, which was rescued by ferroptosis inhibition (**Figure 6B, C**). Within the set of tested PROTACs, a trend towards reduced effects on cell viability was observed for PROTACs carrying an amide group connecting the CCW16 ligand and the chemical linker moiety, which was introduced for synthetic reasons, compared to the derivatives and CCW16 lacking an amide bond at this position. Furthermore, cell viability correlated with cellular permeability of the compounds as estimated by the NanoBRET^TM^. The RNF4 binder alone also led to a significant decrease in cell viability (**Figure 6B, C**), indicating that CCW16 is the chemical moiety triggering the induction of lipid peroxidation followed by ferroptosis.

To exclude that interaction with CRBN was involved in ferroptosis induction, we synthesized a negative control of PROTAC **1b** (**1b n.c**.). **1b n.c.** was identical in its chemical structure to **1b** but contained an N-methyl thalidomide instead of thalidomide, preventing CRBN binding. However, also **1b n.c.** strongly decreased viability, confirming that ferroptosis was induced by the electrophile and not through CRBN interaction of this PROTAC series (**Figure 6D**). Furthermore, we investigated the viability of OCI-AML2 cells after treatment with the RNF4-based BRD4 degrader CCW28-3 and compared it with the treatment of the CRBN-based BRD4 degrader dBET6. CCW28-3 treatment induced a significant decrease in cell viability, which was rescued by ferroptosis inhibition. By contrast, dBET6 did not influence the survival of OCI-AML2 cells (**Figure 6E**). In conclusion, our data revealed that CCW16 covalently bound to a large variety of different proteins thereby inducing a cell death mechanism based on oxidative stress and ferroptosis in an RNF4 independent manner.

## Discussion

The resilience to endogenous or therapy-induced genotoxic stress poses a severe clinical problem. The hypomethylating agent decitabine is a mainstay treatment for myeloid malignancies, but more than 50% of patients do not respond to decitabine or develop resistance in the course of treatment. Decitabine primarily acts by inducing the formation of DNMT1-DPCs, which are cleared via proteasomal degradation through the SUMO-targeted ubiquitylation (StUbL) pathway. Our data show that depletion of the StUbL RNF4 sensitizes cells to decitabine most likely through impairment of DPC clearance. In line with our data, very recent findings demonstrated that the inhibition of the E3 ubiquitin ligase TOPORS, which acts in parallel with RNF4 in proteasomal clearance of DNMT1-DPCs^20,65^, also augments the efficacy of hypomethylating agents^66,67^. Furthermore, our data and published data indicate that upregulation of TOPORS or RNF4 mediates resistance to hypomethylating agents in AML. Targeting of RNF4 or TOPORS can therefore open a therapeutic window for rational combination therapies with decitabine. Given that depletion of RNF4 also sensitizes AML cells to replicative stress, targeting of RNF4 could be generally beneficial in tumors with a high replication stress. Consistent with this idea, inactivation of RNF4 in mice delays tumor formation of *myc*-driven B cell lymphoma, which exhibit a high replicative stress^30^.

RNF4 does not contain a druggable domain, making it a challenging target for the development of conventional small molecule inhibitors. However, covalent inhibitors have been developed targeting cysteine residues in RNF4 for covalent bond formation. Here, we aimed to exploit developed ligands for the design of PROTACs using established CRBN- or VHL-ligands and the irreversible RNF4 ligand CCW16 targeting RNF4 for degradation. However, after synthesizing and testing a set of PROTACs this strategy was not successful. Detailed analysis using recombinant full-length RNF4 as well as cellular extracts revealed covalent binding of CCW16 to two cysteines in RNF4 (C51 and C91), but also covalent attachment to many non-specific covalent interactions in cell extracts including also RNF4. In our proteomic analysis, more than 2000 proteins were captured by biotinylated CCW16 in a streptavidin pulldown experiment. NMR spectra of RNF4 as well as AlphaFold 2 models revealed that RNF4 is largely unstructured except for its RING domain. Therefore, cysteines located in the N-terminus of RNF4 are likely to be solvent-exposed and thus easily targeted by covalent inhibitors. However, due to the unstructured nature of RNF4, the electrophiles used must be highly reactive, as non-covalent interactions with an unstructured binding surface are likely to be weak and cannot enhance the local concentration of covalent compounds. Thus, the electrophile used in CCW16, a chloro-N-phenyl-acetamide, does not appear to be sufficiently selective for its designated target, RNF4. However, this electrophile has been frequently used in the development of covalent ligands. For example, chloro-N-acetamide is the covalent warhead in the DDB1 ligand MM-02-57^68^, the E3 ligands targeting RNF114 such as EN219^69^, a minimal recruitment moiety, targeting DCAF16^70^ as well as covalent FEM1B ligands^71^ and many others. Interestingly, similar to what we observed with our CCW16-based PROTACs, the DCAF16 ligand KB02 reduces cellular ACAT1 levels presumably by targeting a highly reactive cysteine (C126) in ACAT1^70^. These data suggest that ACAT1 may represent a common off-target of chloro-N-acetamide based PROTACs.

Highly reactive ligands, such as the approved drugs used for the treatment of multiple sclerosis have been described to covalently bind to cysteines within the BTB domain of the E3 ligase Keap1^72^. However, these drugs also induce oxidative stress in cells, resulting in the induction of the expression of antioxidative transcription factors such as NRF2. Here, we also observed induction of oxidative stress and the cell death program ferroptosis in cells treated with chloro-N-phenyl-acetamide containing PROTACs. This observation prompted us to reexamine published CCW28–3 based degraders targeting BRD4 which have been developed as a proof of concept that covalent RNF4 ligands can be used for the development of target specific degraders^36^. However, our data suggest that RNF4-based BRD4 degrader CCW28–3 does not function through RNF4 recruitment as observed BRD4 degradation was independent of RNF4. The partial rescue of BRD4 degradation by neddylation inhibition suggested that BRD4 is degraded by a yet to be identified cullin-based E3 ligase.

We propose that the high reactivity of the chloro-N-phenyl-acetamide results in a stress signature observed by our proteomic study which revealed a dramatic up-regulation of HMOX1 together with enhanced expression of NRF2, a key transcription factor in redox signaling and inducer of cytoprotective key enzymes, including HMOX1. HMOX1 is important for the protection of cells from oxidative damage during stress by degrading heme^73^. Further, HMOX1 is a typical marker of ferroptosis, a specific type of programmed cell death characterized by lipid peroxidation through enhanced ROS production and iron overload^74^. We observed a strong ferroptotic potential of CCW16 and the CCW16-based PROTAC molecules in AML cells, which are known to be very sensitive towards this pathway^75^. Importantly, our data demonstrate that the ferroptotic activity of CCW16 is independent of RNF4 or the linkage of CCW16 to a CRBN- or VHL-recruiter, further supporting our interpretation that CCW-16 has an intrinsic ferroptotic potential, which can be abrogated with the ferroptosis inhibitor ferrostatin. Lipid peroxidation is primarily detoxified by the action of glutathione peroxidase 4 (GPX4) or peroxiredoxins, such as PRDX6^64,76,77^. Interestingly, we identify the active cysteine residues of PRDX1 (C52), PRDX2 (C51) and PRDX6 (C47) as direct covalent targets of CCW16, suggesting that inhibition of these PRDX proteins by CCW16 contributes to the ferroptosis phenotype. High expression of PRDX1 correlates with an unfavorable prognosis in AML patients indicating that targeting of PRDX1 by CCW16 might be a therapeutic option in a subset of AML patients^78^. Notably, by using the CCW16 derivative that contains an alkyne handle, the Nomura group demonstrated targeting of HMOX2 and GPX4 by CCW16 suggesting that CCW16 preferential reacts with highly reactive cysteines of proteins involved in redox signaling^36^. The ferroptotic potential of CCW16 is reminiscent to the small molecule RSL3 that acts as a covalent inhibitor of GPX4^79^. Like CCW16, RSL3 contains an activated chloro-N-acetamide and shows low selectivity^80^. Notably, the covalent E3 ligand EN219 targeting RNF114 that also contains a chloro-N-acetamide binds as well GPX4 suggesting that covalent E3 ligands with this electrophile may potentially be at risk to exert a more general off-target toxicity ^69^. Altogether, our data demonstrated that RNF4 is a clinically relevant drug target in AML. However, CCW16 is unlikely to represent a suitable lead compound for the development of RNF4-based degraders but may have potential as an experimental inducer of ferroptosis.

## Supporting information

Supplemental_Figure_Synthesis

Supplemental_Methods_Synthesis

Supplemental_Table_III

## Acknowledgements

This work was supported by Deutsche Forschungsgemeinschaft (DFG grants MU-1764/6-Project ID-465470262, MU-1764/7-Project ID- 494535244 and CRC387), the BMBF Project: “PROXIDRUGS” funded by the German Federal Ministry of Education and Research to S.M. and S.K. We thank the Quantitative Proteomics Unit at IBC2 (Goethe University, Frankfurt) for supporting MS and the DFG for funding the LC-MS systems used in this study: Orbitrap Fusion LUMOS: FuGG Project-ID: 403765277; QExactive HF: Project-ID: 259130777, SFB1177.

## Notes

### Competing Interest Statement

The authors have declared no competing interest.

### Summary of Updates

A full author list has been provided.

