## Supplemental_Figures_Legends for "The cysteine-reactive covalent RNF4 ligand CCW16 induces ferroptosis in AML cells by activation of ROS signaling"

### Supplements

**Figure S1: RNF4 as a vulnerability of AML cells.**

(A)/(B) Confirmation of functional RNF4 KO in OCI-AML3 (A) and MV4-11 Cas9 expressing cells after transduction with three different guideRNAs by immunoblotting. Cells were transduced and medium was exchanged after one day and supplemented with 2.5 µg/ml puromycin. After 3 days (OCI-AML3) and 4 days (MV4-11) cells were harvested. (C) Validation of functional RNF4 KD corresponding to Figure 1E and 1F by immunoblotting. Cells were harvested 3 days after performance of KD.

**Figure S2: Evaluation of RNF4 targeting PROTACs.**

(A) NanoBRET assay for CRBN and VHL E3 ligand based RNF4 PROTACs to determine cell membrane permeability. $IC_{50}$ values are indicated. (B) Treatment of HeLa FLAG-RNF4 (endog.) cells with different RNF4 targeting PROTACs and evaluation of RNF4 degradation level by immunoblotting. Different used concentrations and time points are indicated. Control cells were treated with DMSO. Tubulin was used as loading control. (C) Evaluation of RNF4 degradation level by immunoblotting in NB-4 cells after treatment with different RNF4 targeting PROTACs. Control cells were treated with DMSO. Vinculin was used as loading control. (D) Treatment of OCI-AML2 cells with different RNF4 targeting PROTACs and evaluation of RNF4 degradation level by immunoblotting. Cells were pre-treated with MG-132 [20 µM], TAK-243 [1 µM] or MLN-4942 [500 nM] 30 min before PROTAC treatment [5 µM] and harvested after 6 hours. Control cells were treated with DMSO. Tubulin was used as loading control. (E) Evaluation of CRBN and VHL levels in HeLa WT cells by immunoblotting after pre-treating cells with MG-132 [20 µM], TAK-243 [1 µM] or MLN-4942 [500 nM] 30 min before PROTAC treatment [5 µM]. DMSO was used as control treatment and tubulin as loading control.

**Figure S3: Further evaluation of RNF4-binder CCW16 *in vitro*.**

(A) MS Spectrum of CCW16 modified GST-RNF4 on cysteine residue 91. (B) Detected RNF4 peptides including the respective modifications on different cysteine residues (as indicated). (C) MS Spectrum of 2a modified GST-RNF4 on cysteine residue 91.

**Figure S4: Further evaluation of RNF4-binder CCW16 *in vitro*.**

(A) Pre-treatment of HeLa WT cells with MG-132 [20 µM] or MLN-4942 [500 nM] 30 min followed by treatment with CCW28-3 [10 µM] or dBET6 [500 nM] for 6 hours and evaluation by immunoblotting. *Unspecific band (B) Confirmation of KD efficiency 3 days after performance of KD of Figure 4C and 4D by immunoblotting. (C) Evaluation of cell viability after CCW28-3 treatment in HEK BRD4-HiBiT cell lines by CellTiterGlo assay. Used concentrations and time points are indicated.

**Figure S5: Identification of CCW16 targets *in vivo*.**

(A) Volcano plot of quantitative MS analysis after biotin-CCW16 pull down of HeLa WT cell lysates. Significantly enriched interactors are shown in red ($\log_2$ ratio ≥ 1, $-\log_{10}$ p value ≥ 1.3). Identification of candidates is based on two-sided Student's t-test analysis comparing LFQ intensities of biotin-CCW16 pulldown and biotin control pulldown. Experiment was performed in triplicates. Proteins involved in the reduction of peroxides are additionally highlighted. (B) Gene Ontology term enrichment analysis of biological processes (GOBP) of the 30 biotin-CCW16 modified proteins (from Figure 5B) identified by MS (log2 ratio ≥ 1, –log10 p value ≥ 1.3). Shown here are the top 10 enriched biological processes. The enrichment analysis was done using the ShinyGO tool, applying an FDR cutoff of 0.05. (C) RNF4 immunoblotting of Streptavidin pulldown in HeLa WT cells. Same treatment procedure as in Figure 5A. Tubulin was used as loading control. (D) Kyoto Encyclopedia of Genes and Genomes (KEGG) pathway analysis of the 38 biotin-CCW16 modified proteins (from Figure 5B) identified by MS ($\log_2$ ratio ≥ 1, $-\log_{10}$ p value ≥ 1.3). Shown here are the top 3 enriched biological processes. The enrichment analysis was done using the ShinyGO tool, applying an FDR cutoff of 0.05.

Supplements
Figure 1

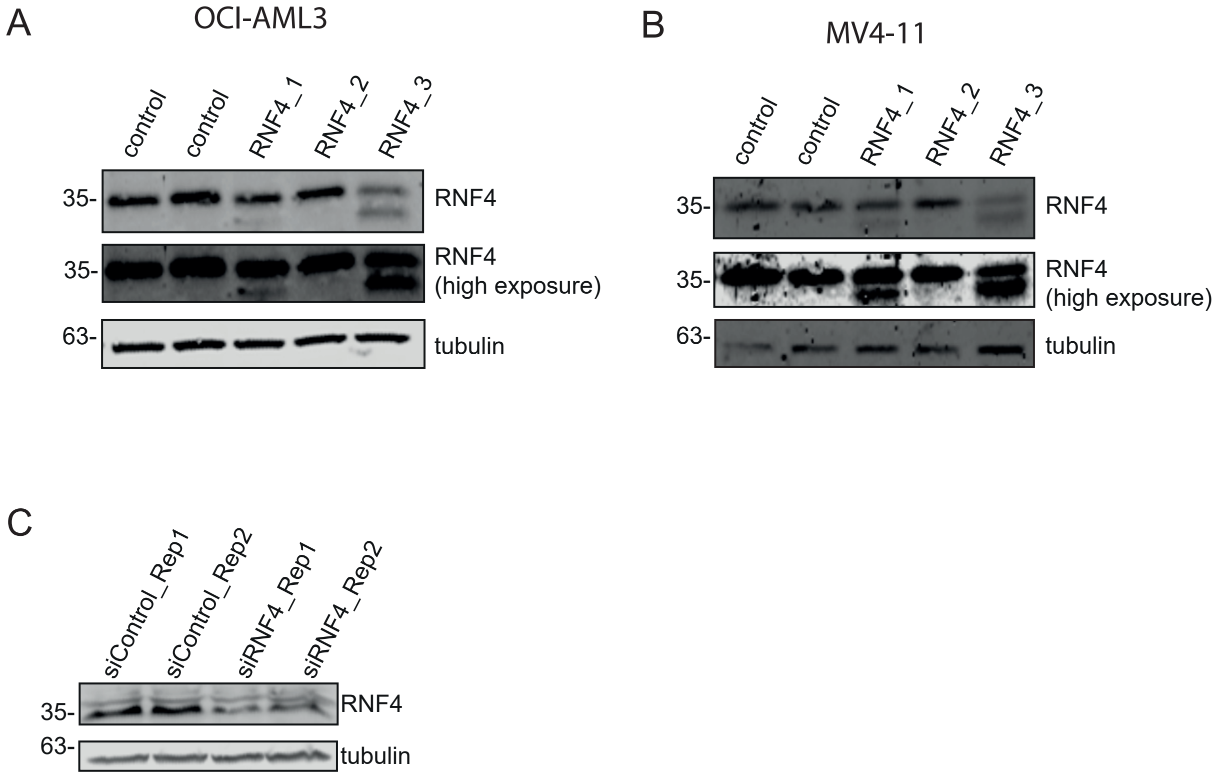

Supplements
Figure 2

A

| | 1a - CRBN | 1b - CRBN | 1c - CRBN | 1d - CRBN | Len. - CRBN | 2a - VHL | 2c - VHL | VH298 VHL |
|---|---|---|---|---|---|---|---|---|
| $LogIC_{50}$ | -6.381 | -6.987 | -6.934 | -6.835 | -6.763 | -4.24 | -4.293 | -6.513 |
| $IC_{50}$ | 4.16E-07 | 1.03E-07 | 1.17E-07 | 1.46E-07 | 1.73E-07 | 5.76E-05 | 5.09E-05 | 3.07E-07 |

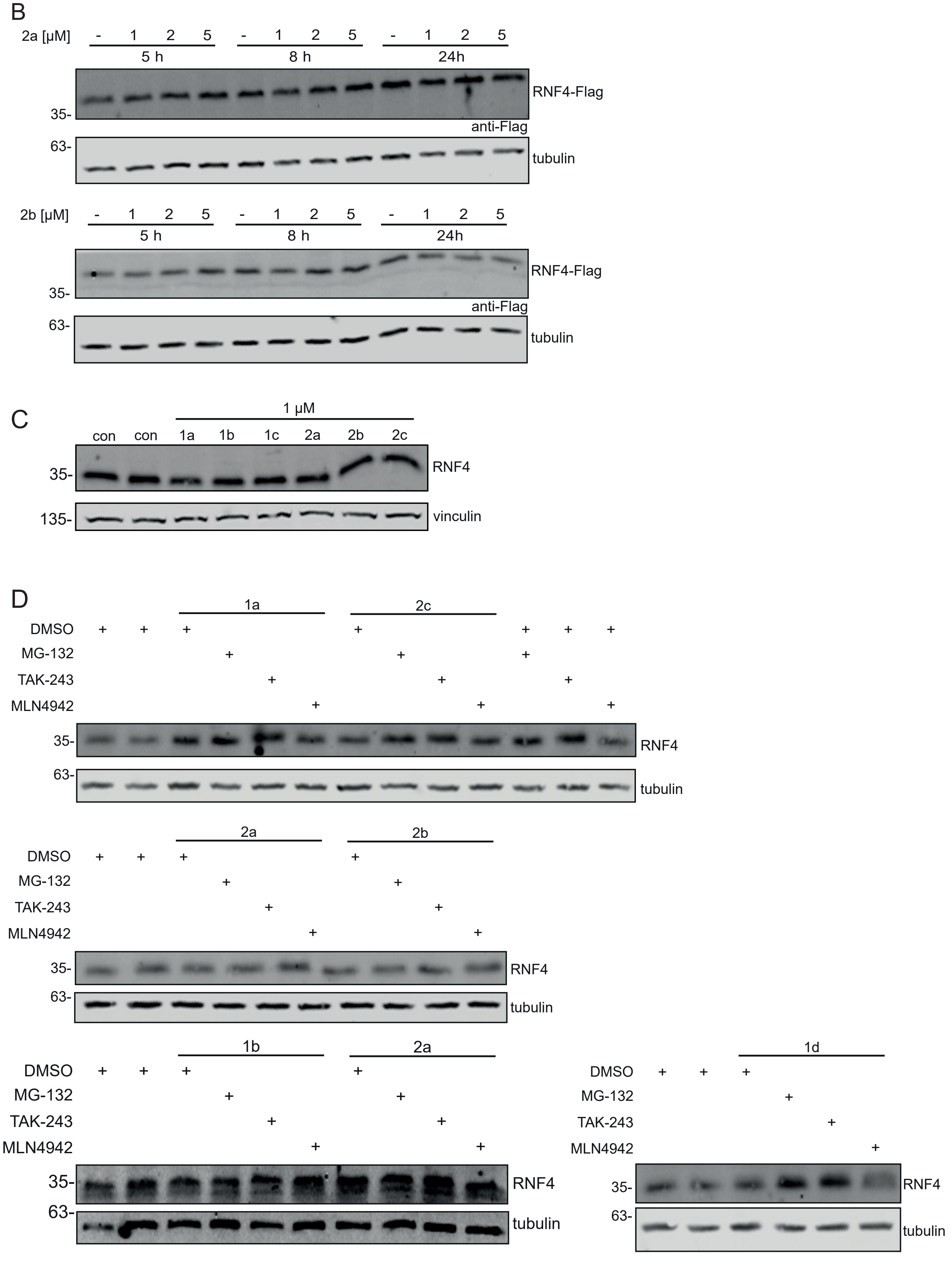

Supplements
Figure 2

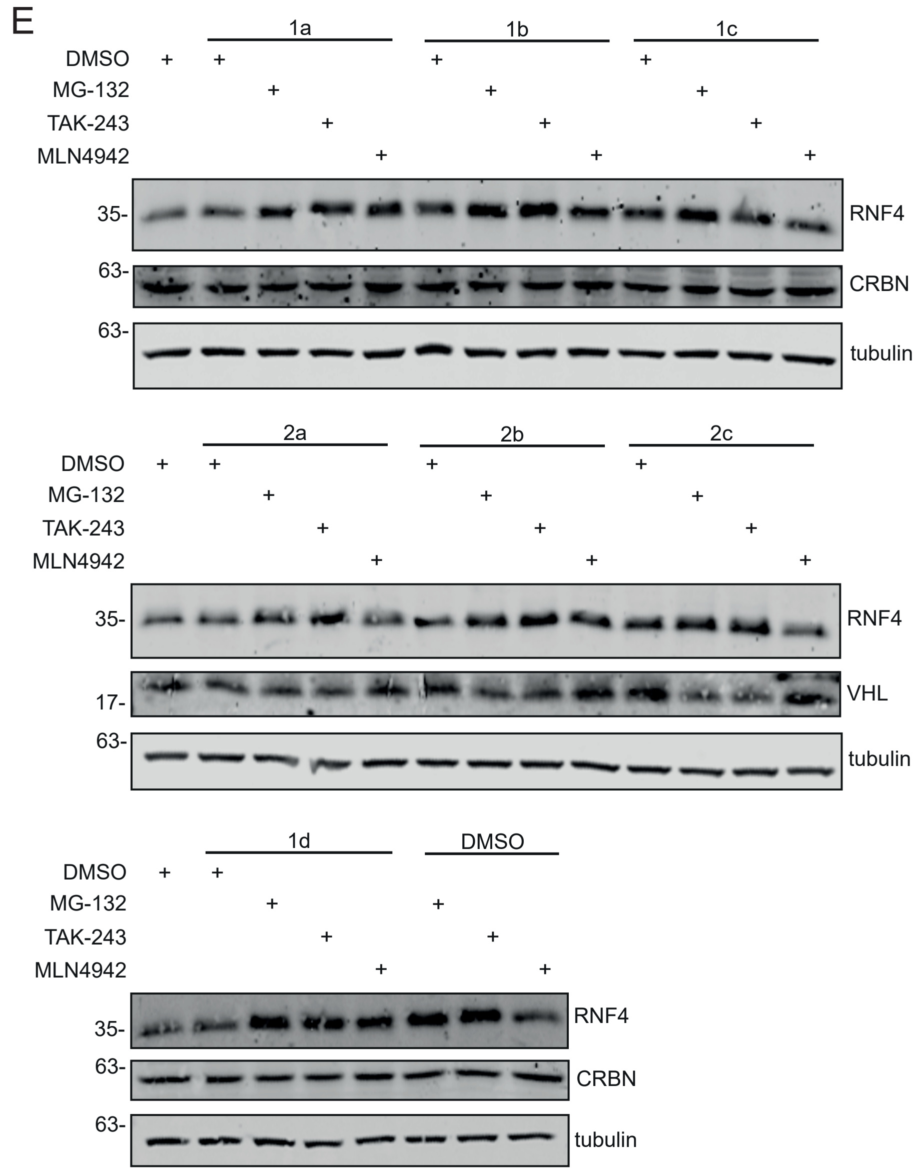

Supplements
Figure 3

A

CCW16 C91

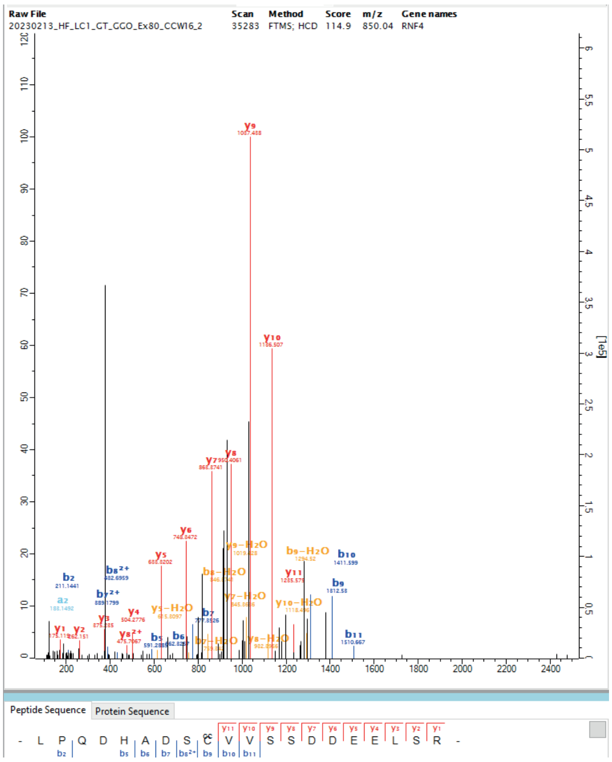


C

2a C91

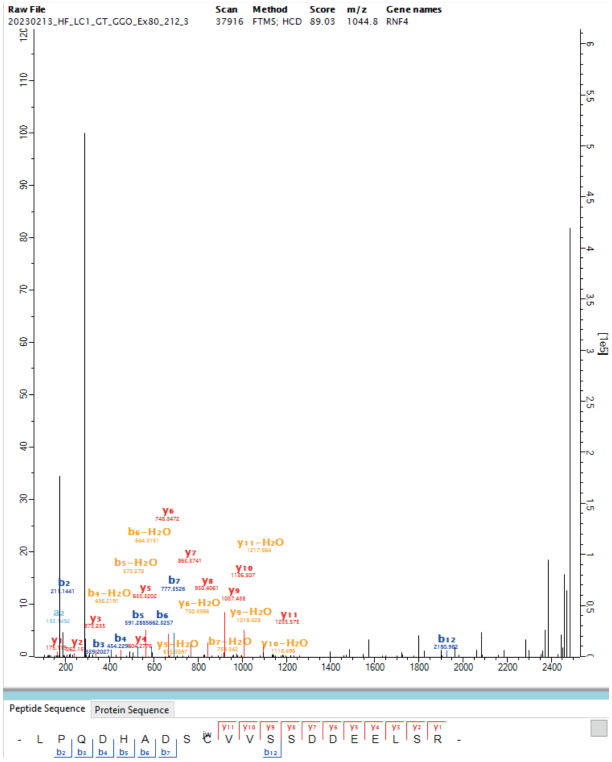


B

| Sequence | Modifications | Cysteine residues |
|---|---|---|
| DEGATGLR | Unmodified | - |
| DEGATGLRPSGTVSCPICMDGYSEIVQNGR | Carbamidomethyl (C) | C132, C135 |
| DEGATGLRPSGTVSCPICMDGYSEIVQNGR | 2 Carbamidomethyl (C);Oxidation (M) | C132, C135 |
| DEGATGLRPSGTVSCPICMDGYSEIVQNGR | Oxidation (M) | C132, C135 |
| DRDVYVTTHTPR | Unmodified | - |
| DVYVTTHTPR | Unmodified | - |
| GGAINSR | Unmodified | - |
| LIVSTECGHVFCSQCLR | 3 Carbamidomethyl (C) | C154, C159, C162 |
| LPQDHADSCVVSSDDEELSR | Unmodified | - |
| LPQDHADSCVVSSDDEELSR | Carbamidomethyl (C) | C91 |
| LPQDHADSCVVSSDDEELSR | 2a (C) | C91 |
| LPQDHADSCVVSSDDEELSR | CCW16 (C) | C91 |
| NANTCPTCR | 2 Carbamidomethyl (C) | C173, C176 |
| PSGTVSCPICMDGYSEIVQNGR | 2a (C) | C132, C135 |
| RLPQDHADSCVVSSDDEELSR | Carbamidomethyl (C) | C91 |
| RLPQDHADSCVVSSDDEELSR | Unmodified | C91 |
| RLPQDHADSCVVSSDDEELSR | CCW16 (C) | C91 |
| RYHPIYI | Unmodified | - |

Supplements
Figure 4

A

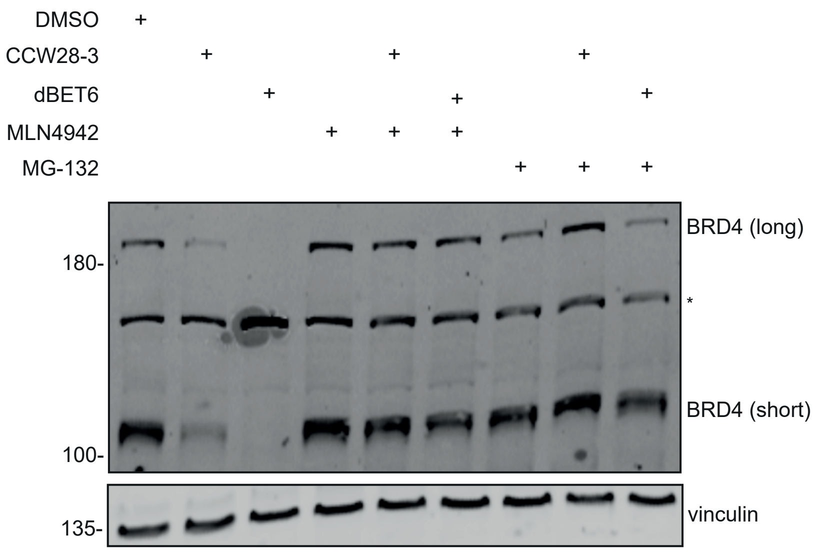


B

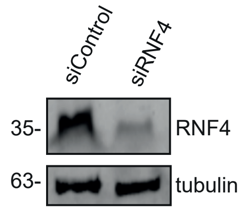


C

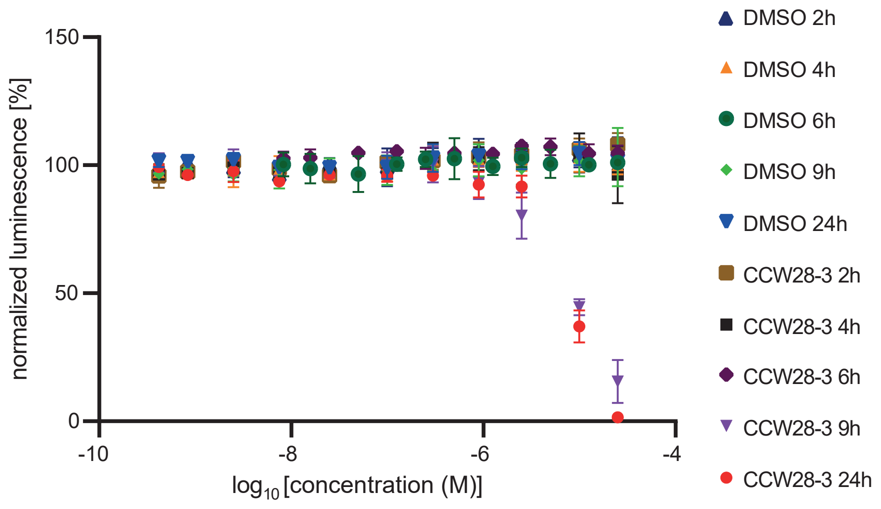

Supplements

Figure 5

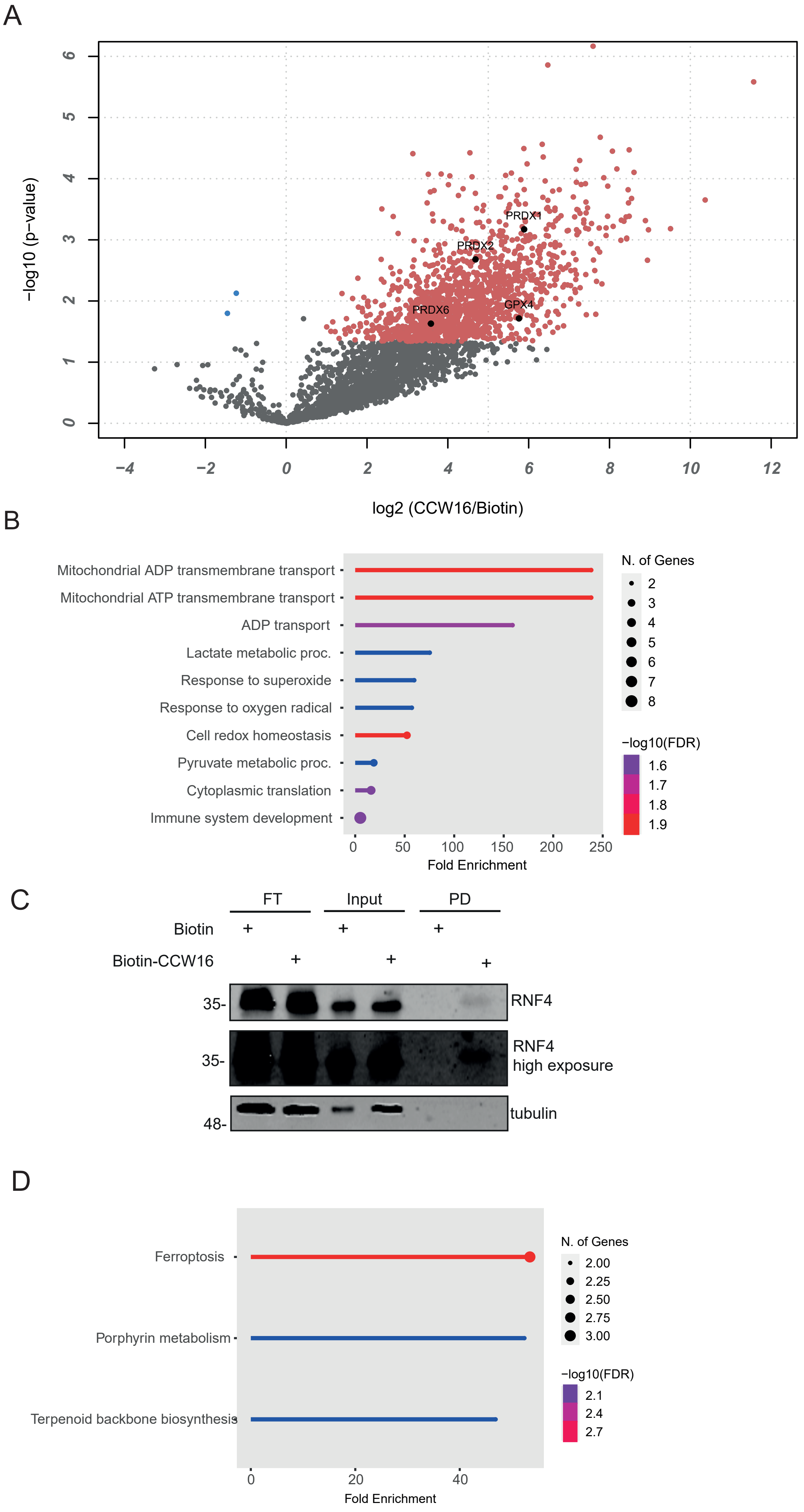
