## Supplemental_Figure_Synthesis for "The cysteine-reactive covalent RNF4 ligand CCW16 induces ferroptosis in AML cells by activation of ROS signaling"

**^1^H, ^13^C NMR Spectra, Mass spectra and Chromatograms, HRMS**

2-(2,6-dioxopiperidin-3-yl)-4-hydroxyisoindoline-1,3-dione **S3**

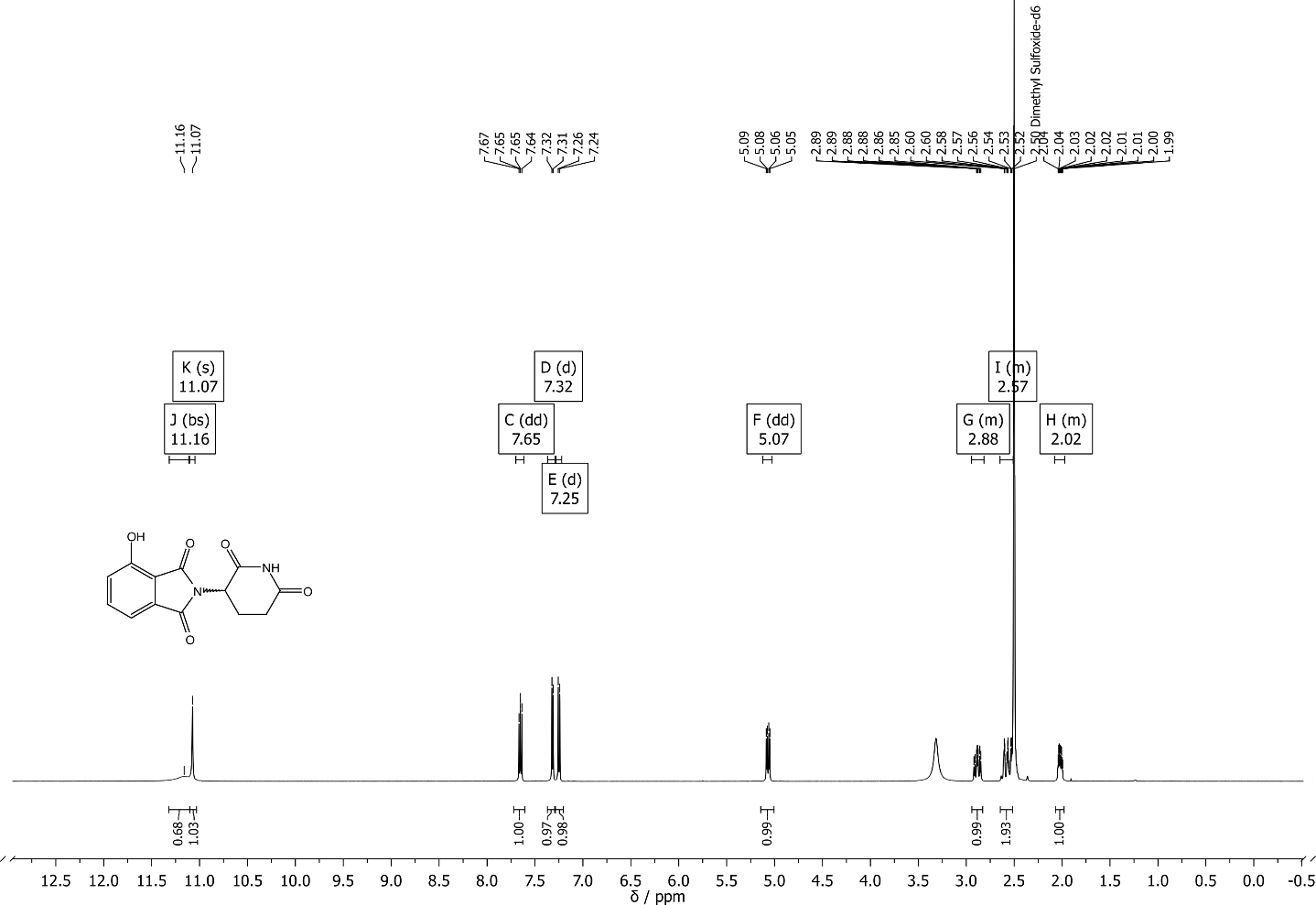

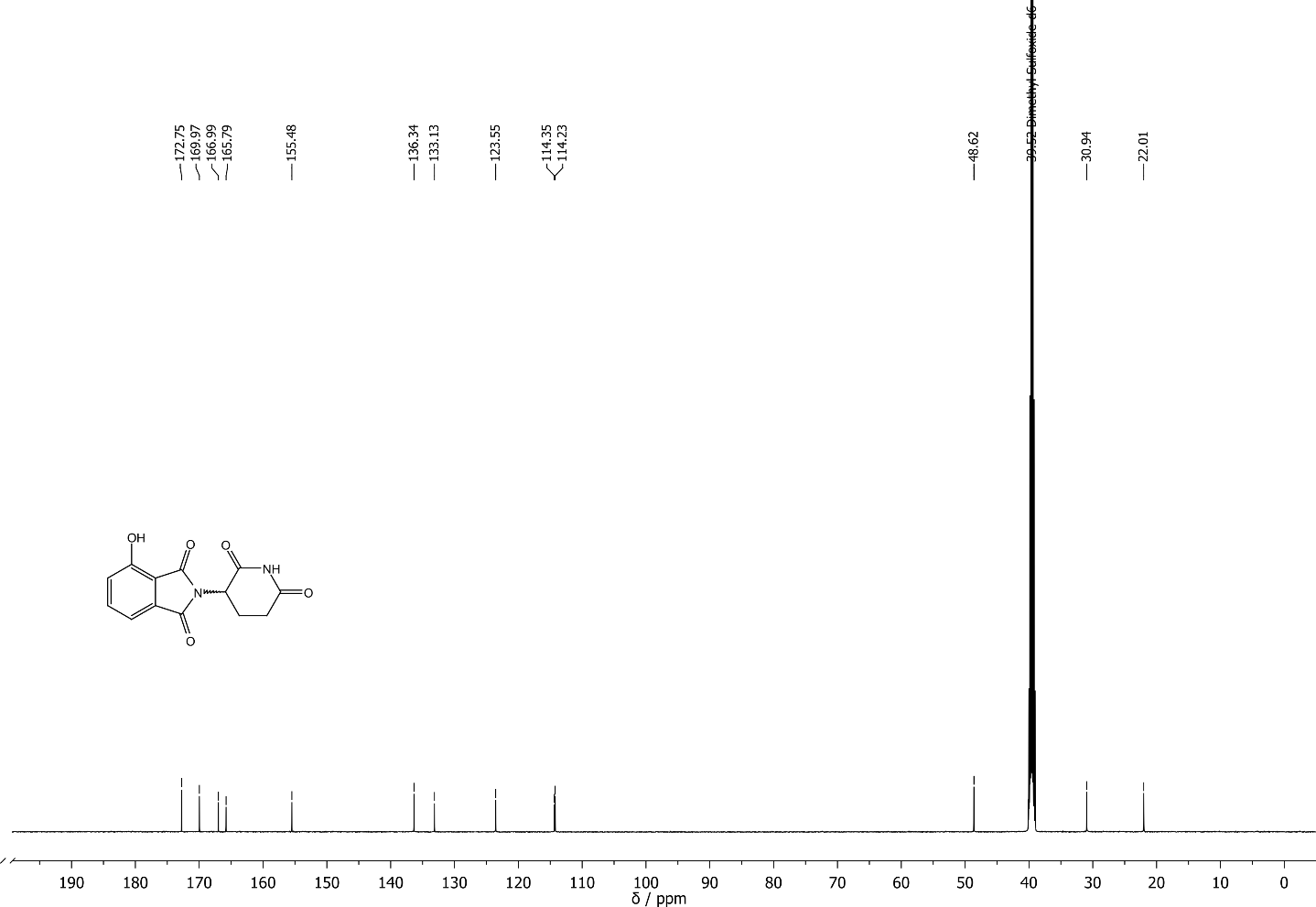

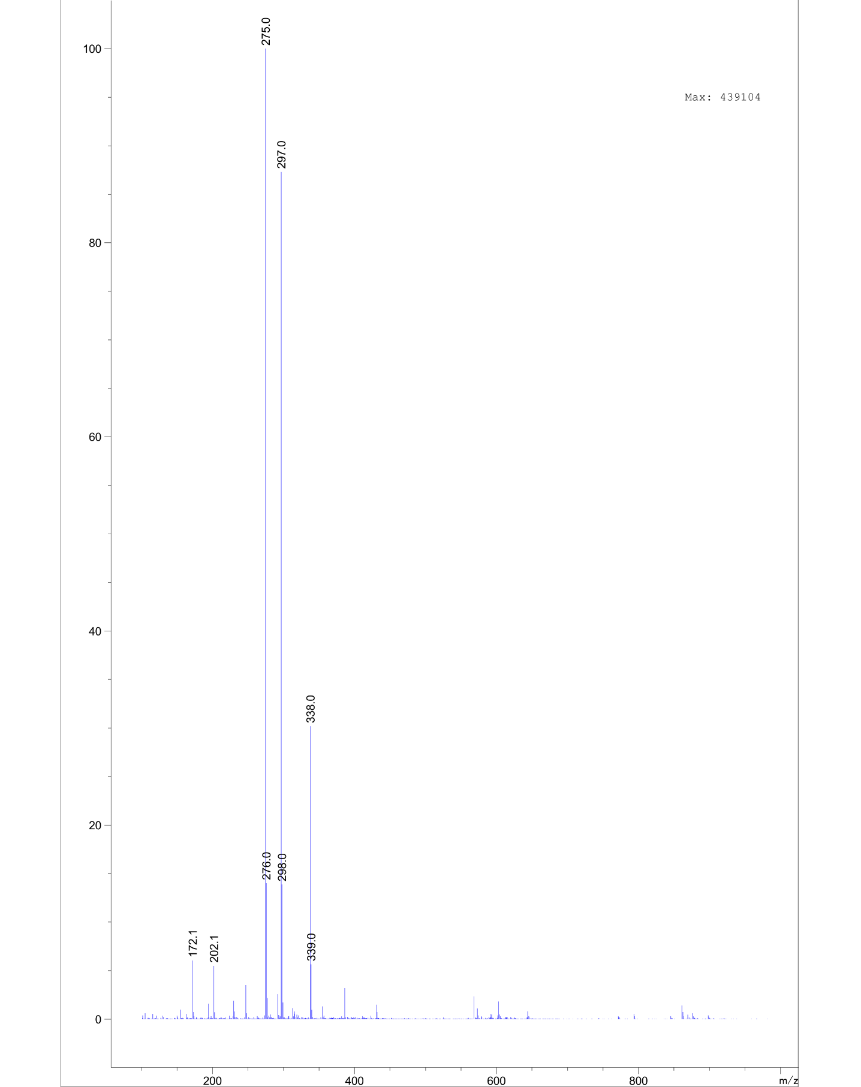

*tert*-butyl 2-((2-(2,6-dioxopiperidin-3-yl)-1,3-dioxoisoindolin-4-yl)oxy)acetate **S4**

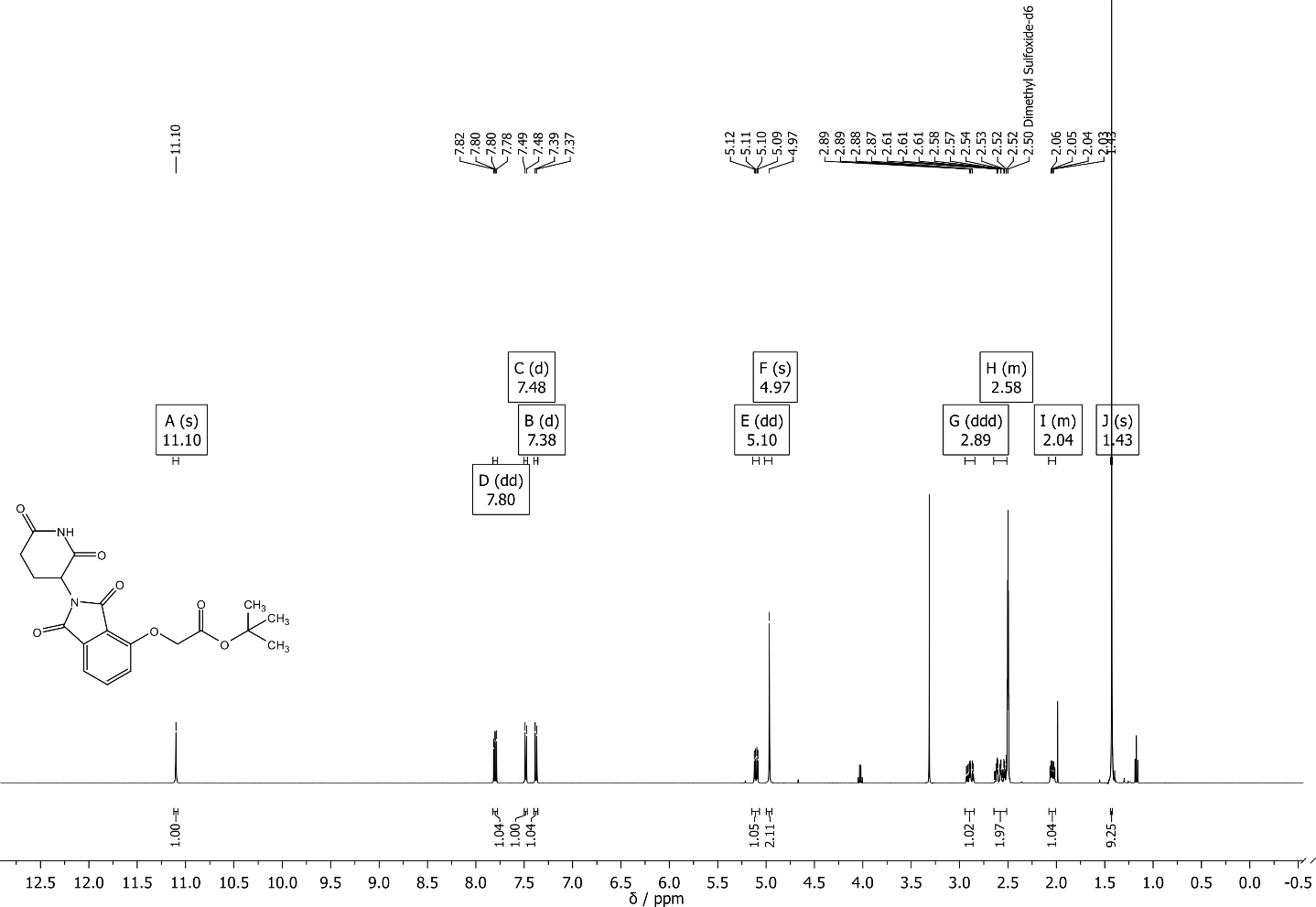

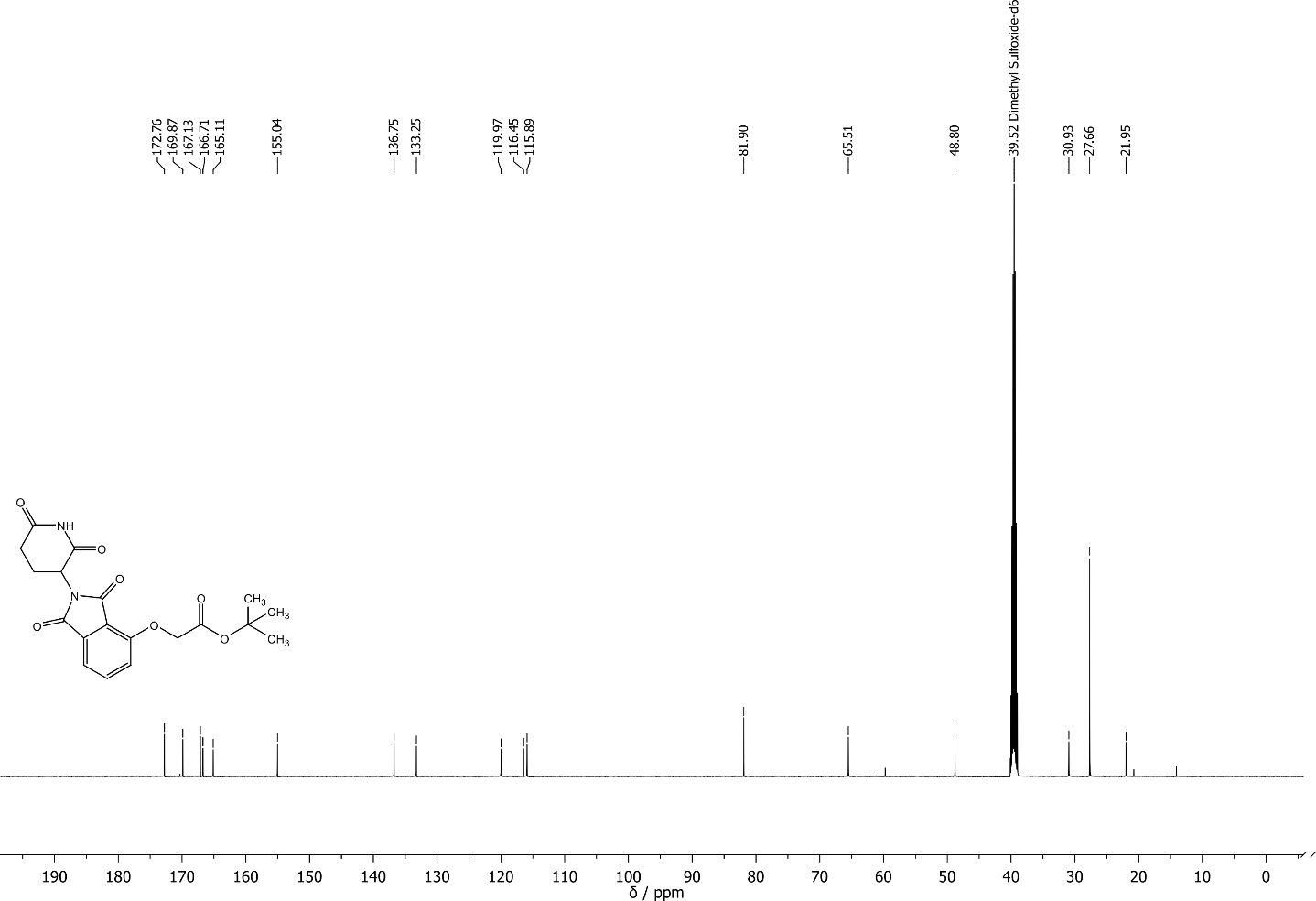

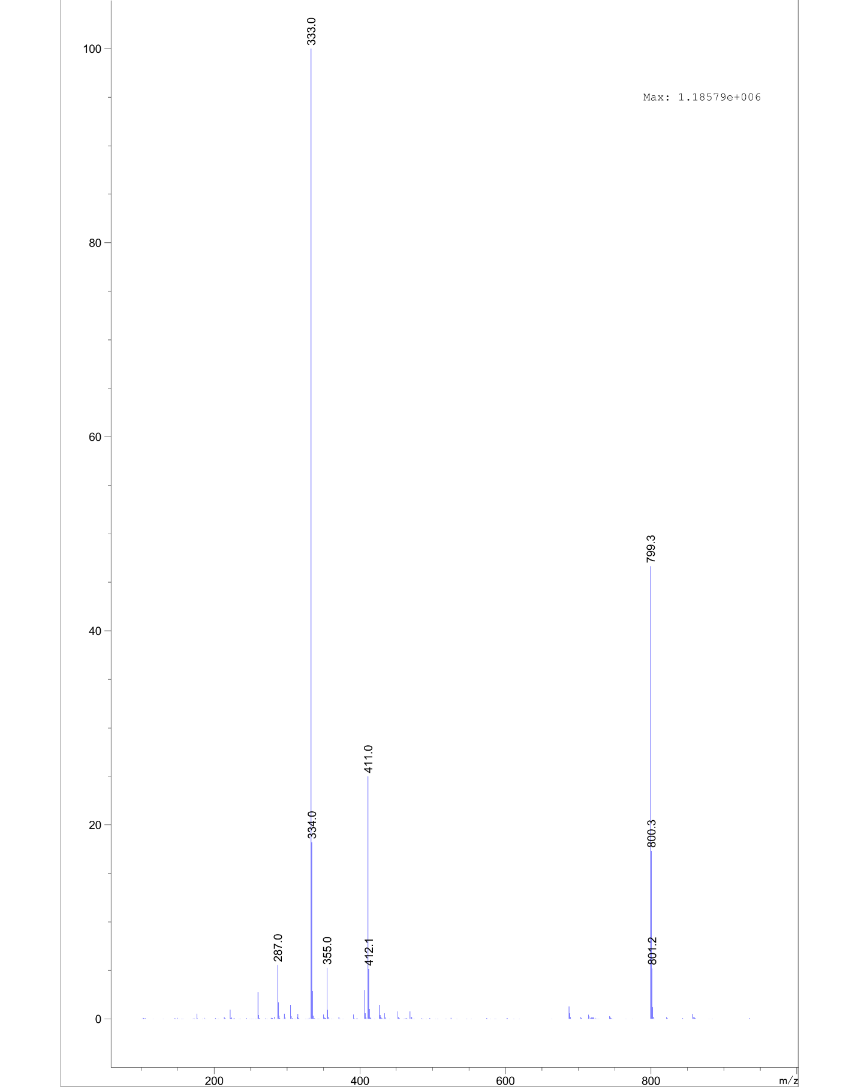

tert-butyl 2-((2-(1-methyl-2,6-dioxopiperidin-3-yl)-1,3-dioxoisoindolin-4-yl)oxy)acetate **S5**

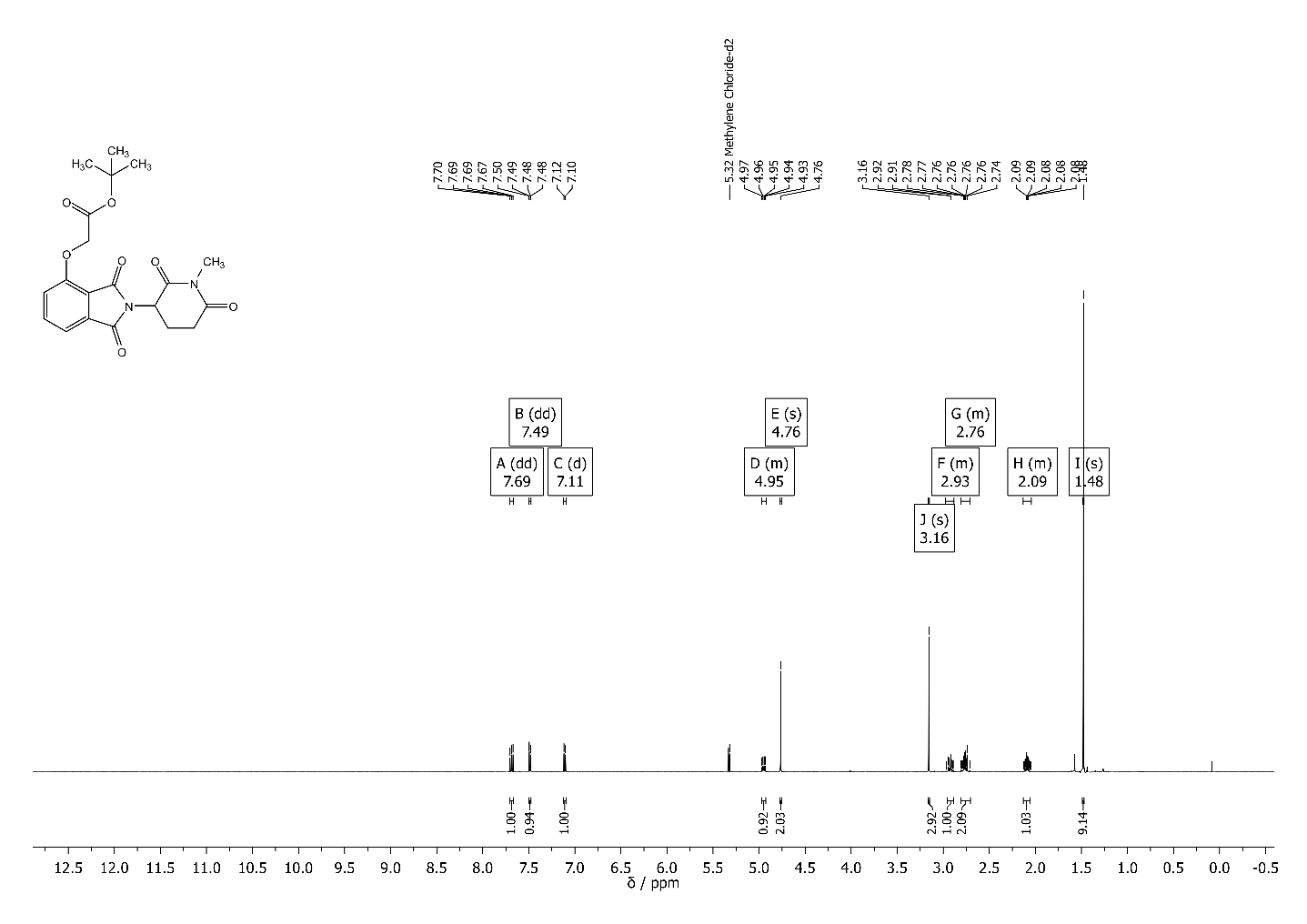

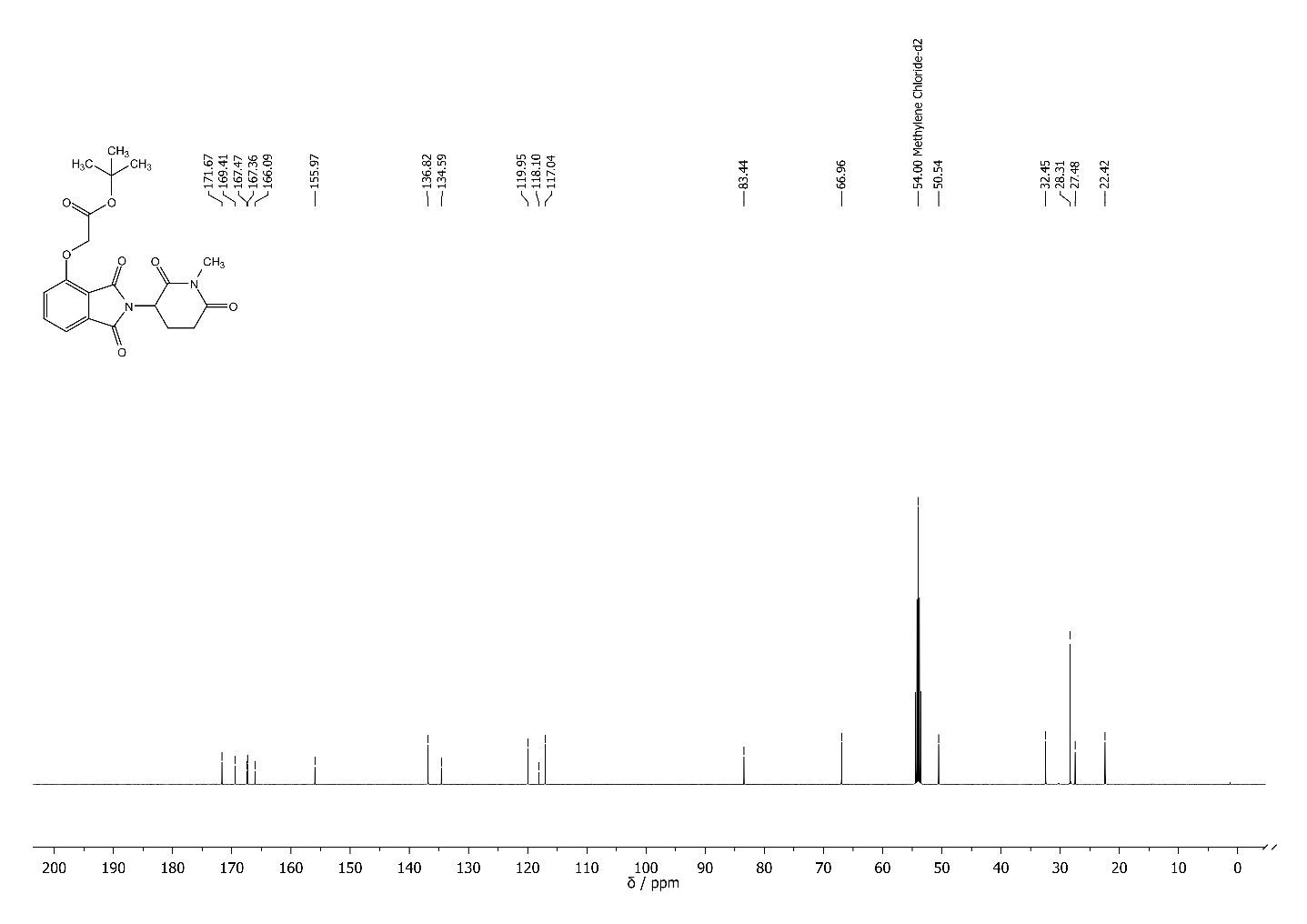

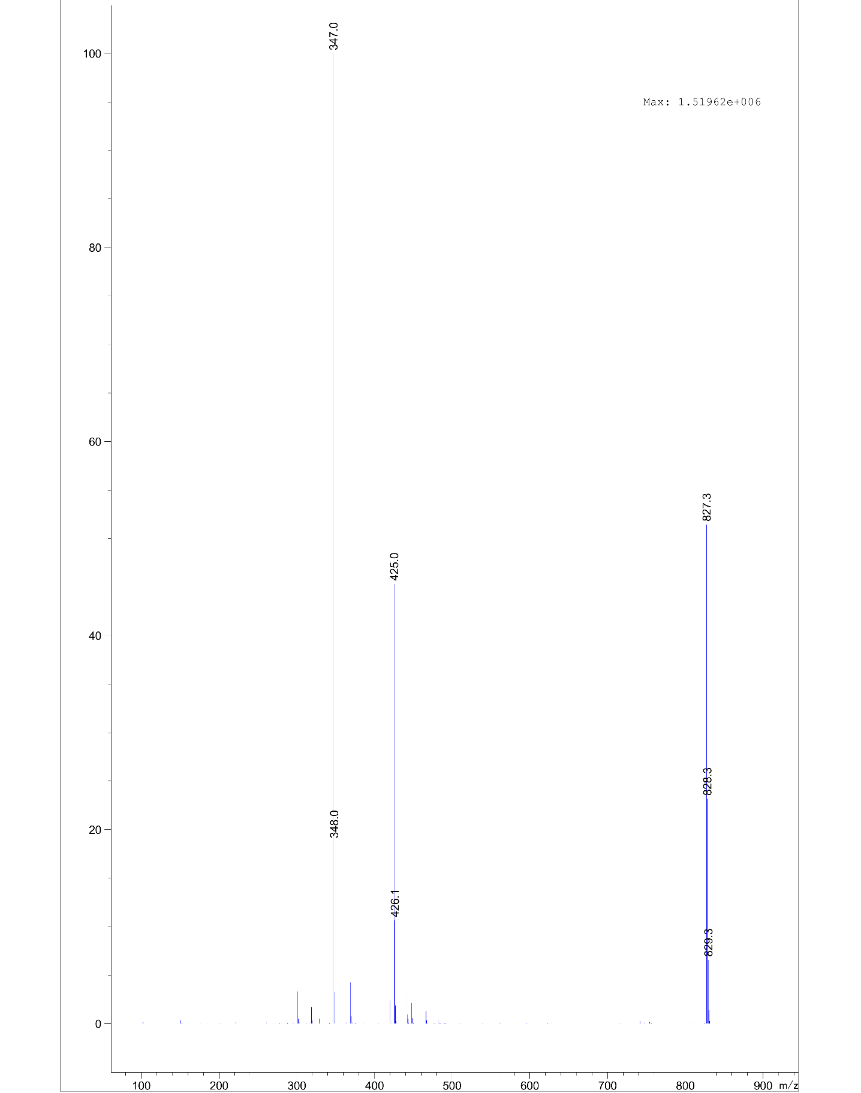

4-(4-(benzylamino)phenoxy)phenol **S7** via 4-(4-aminophenoxy)phenol

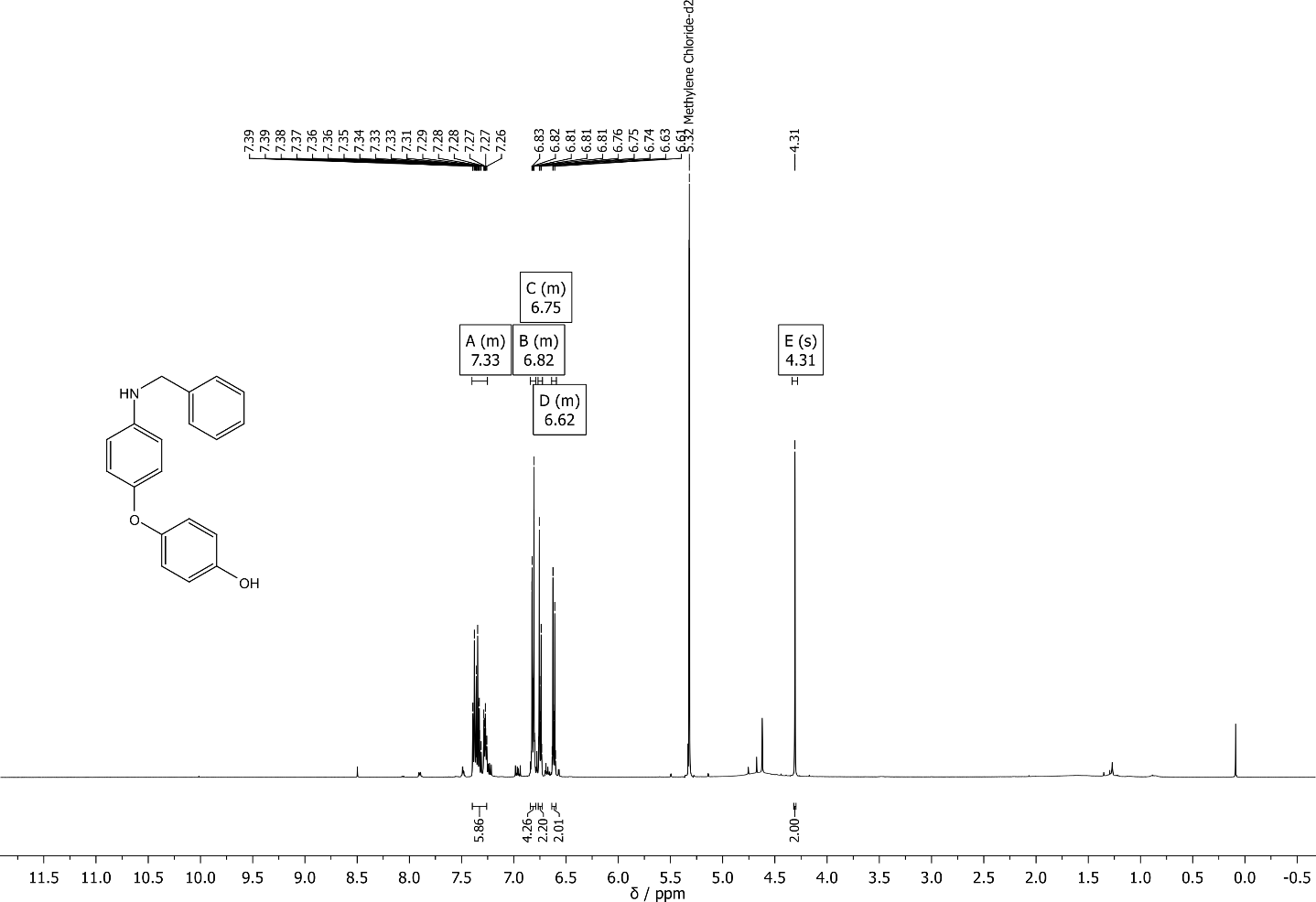

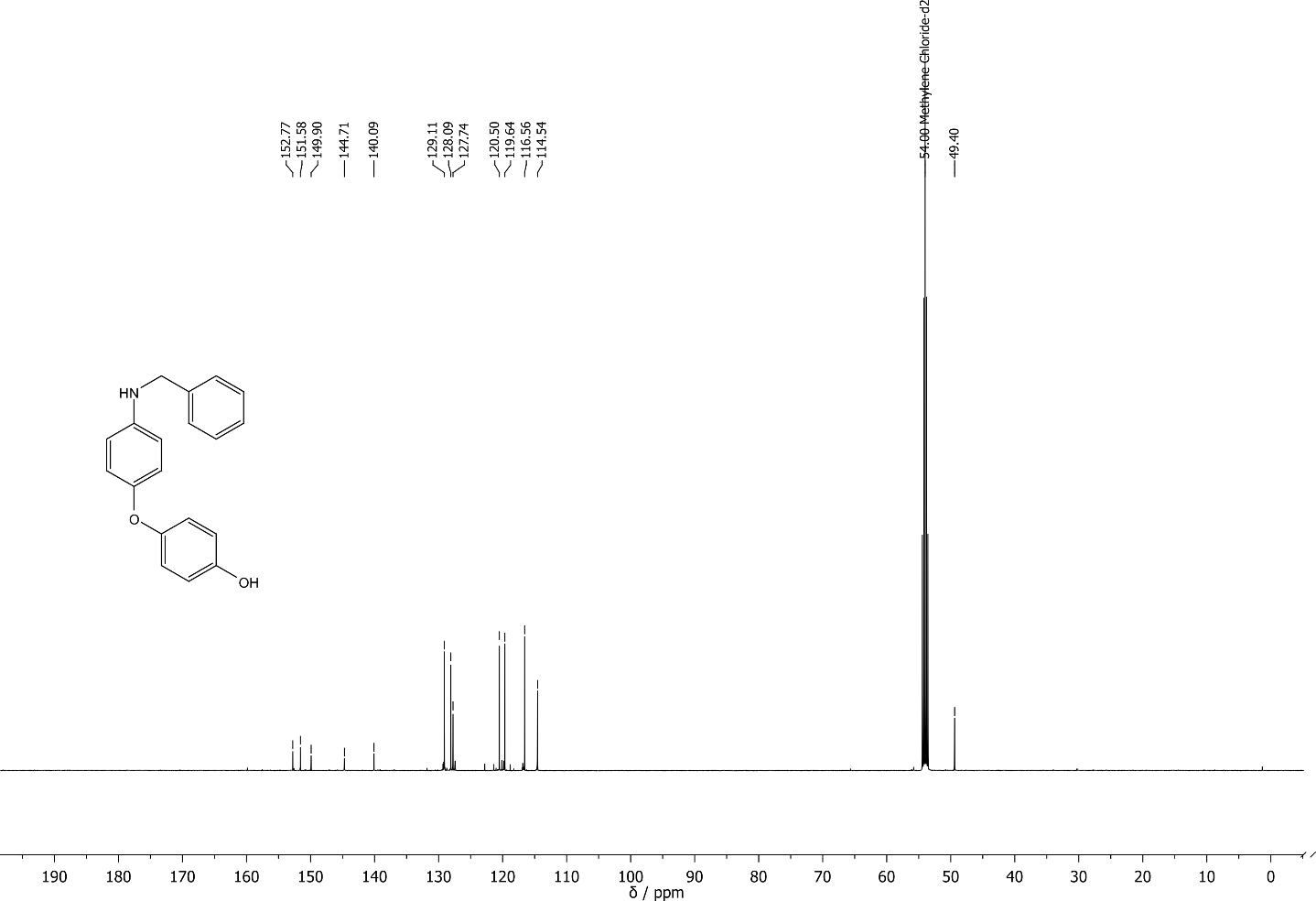

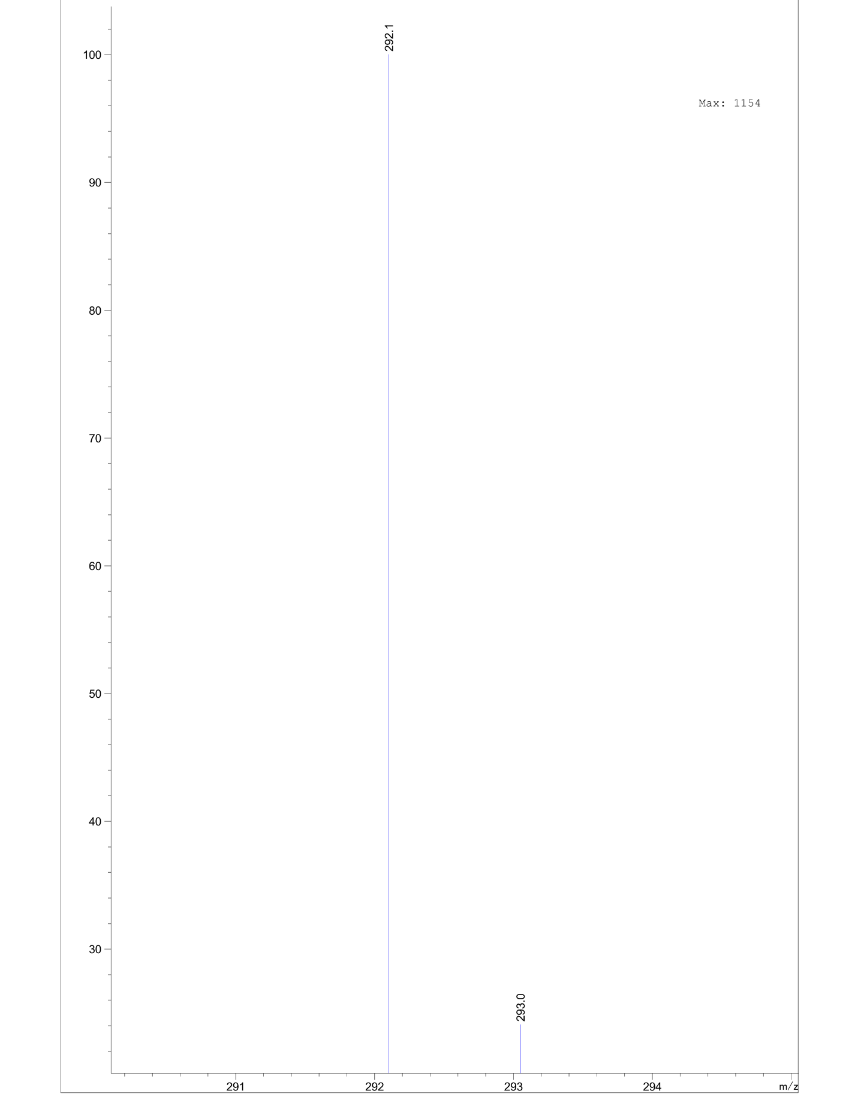

tert-butyl 2-(4-(4-(benzylamino)phenoxy)phenoxy)acetate **S8**

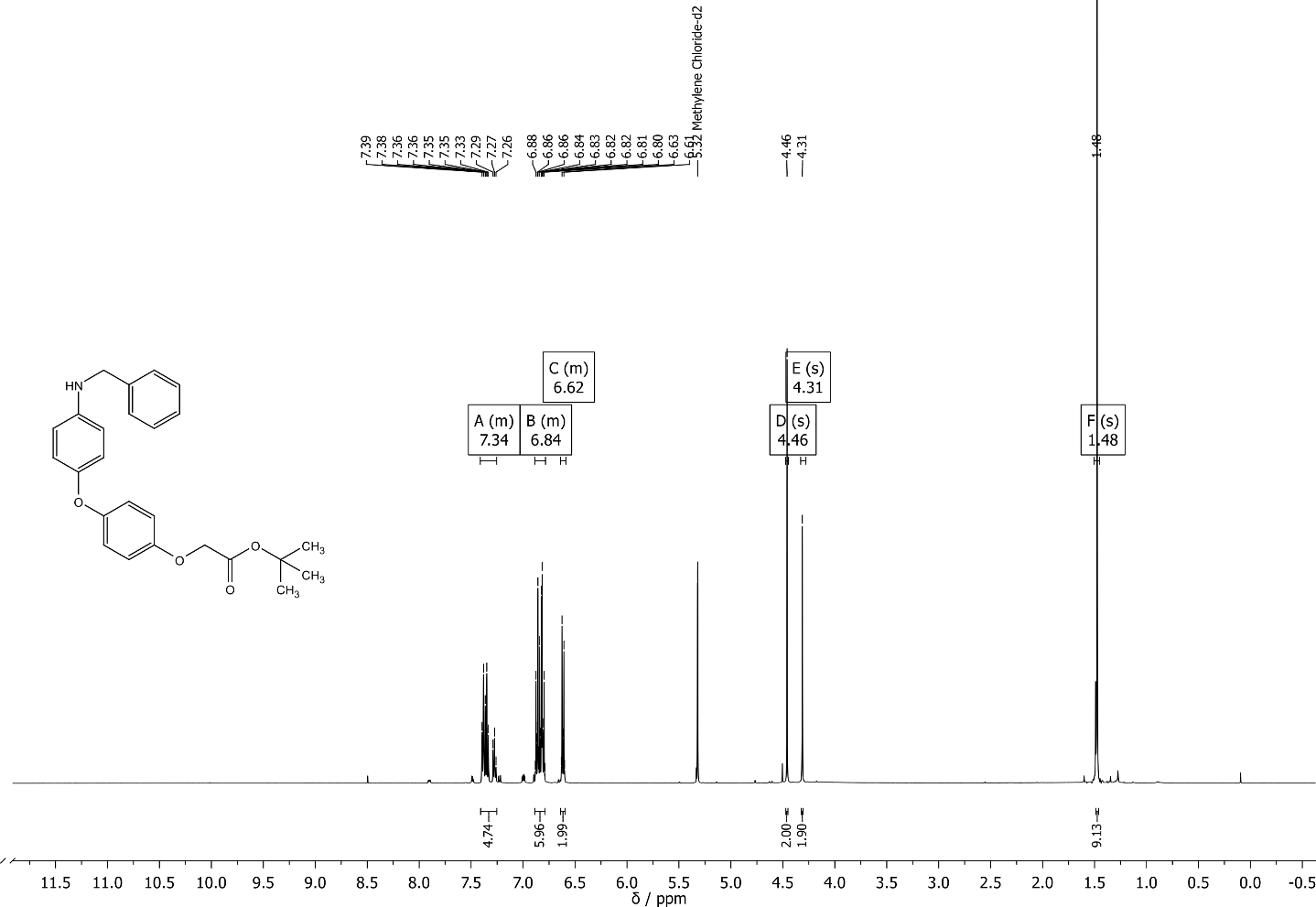

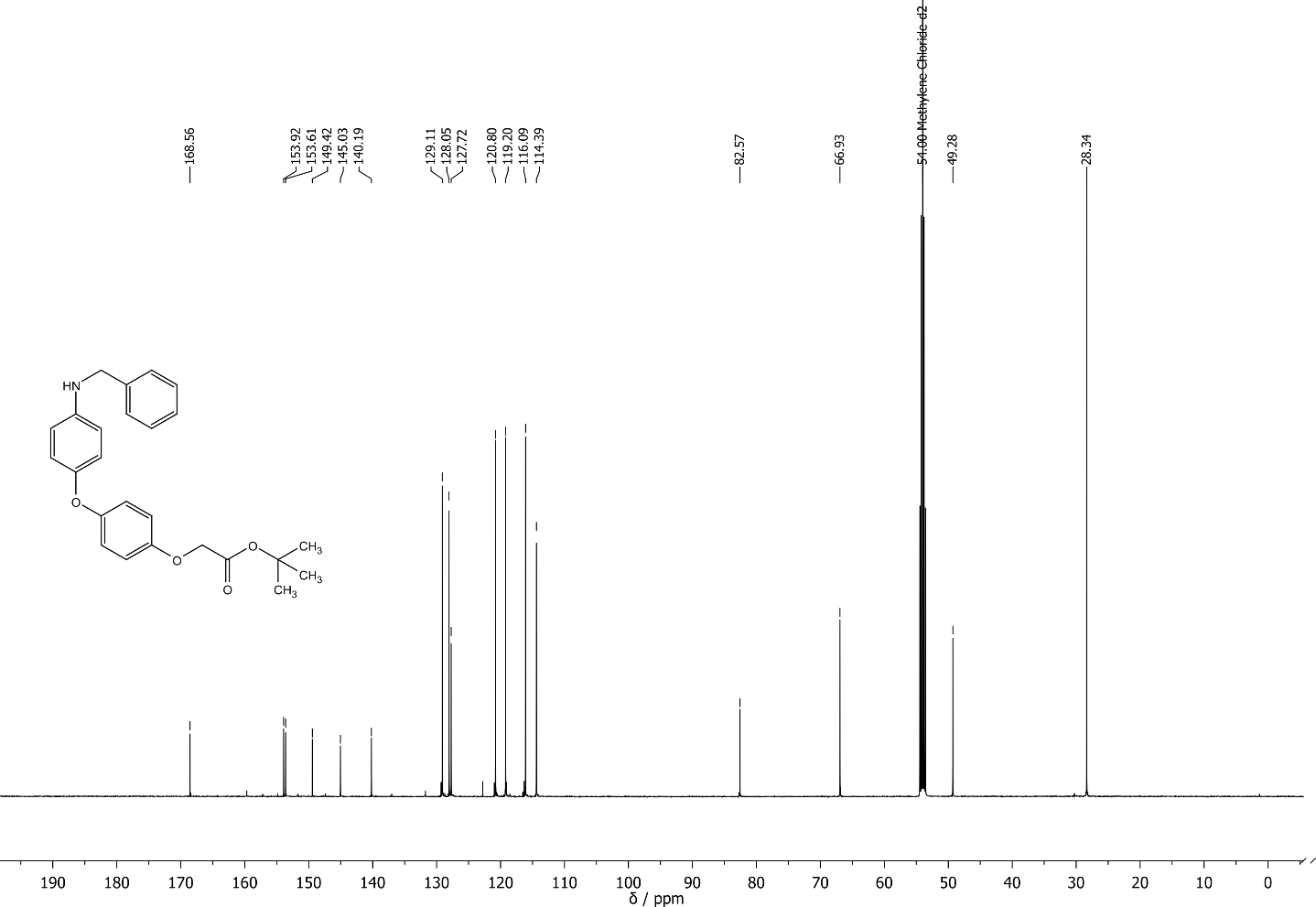

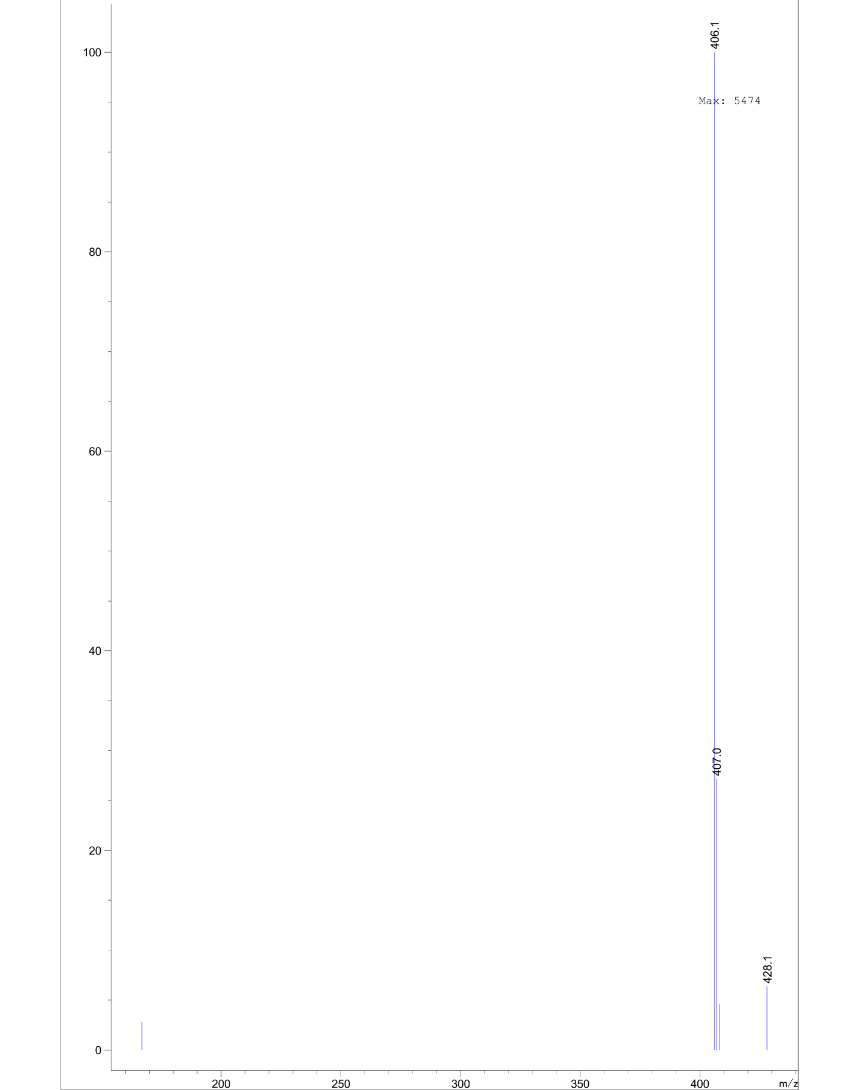

N-benzyl-2-chloro-N-(4-(4-methoxyphenoxy)phenyl)acetamide **CCW16**

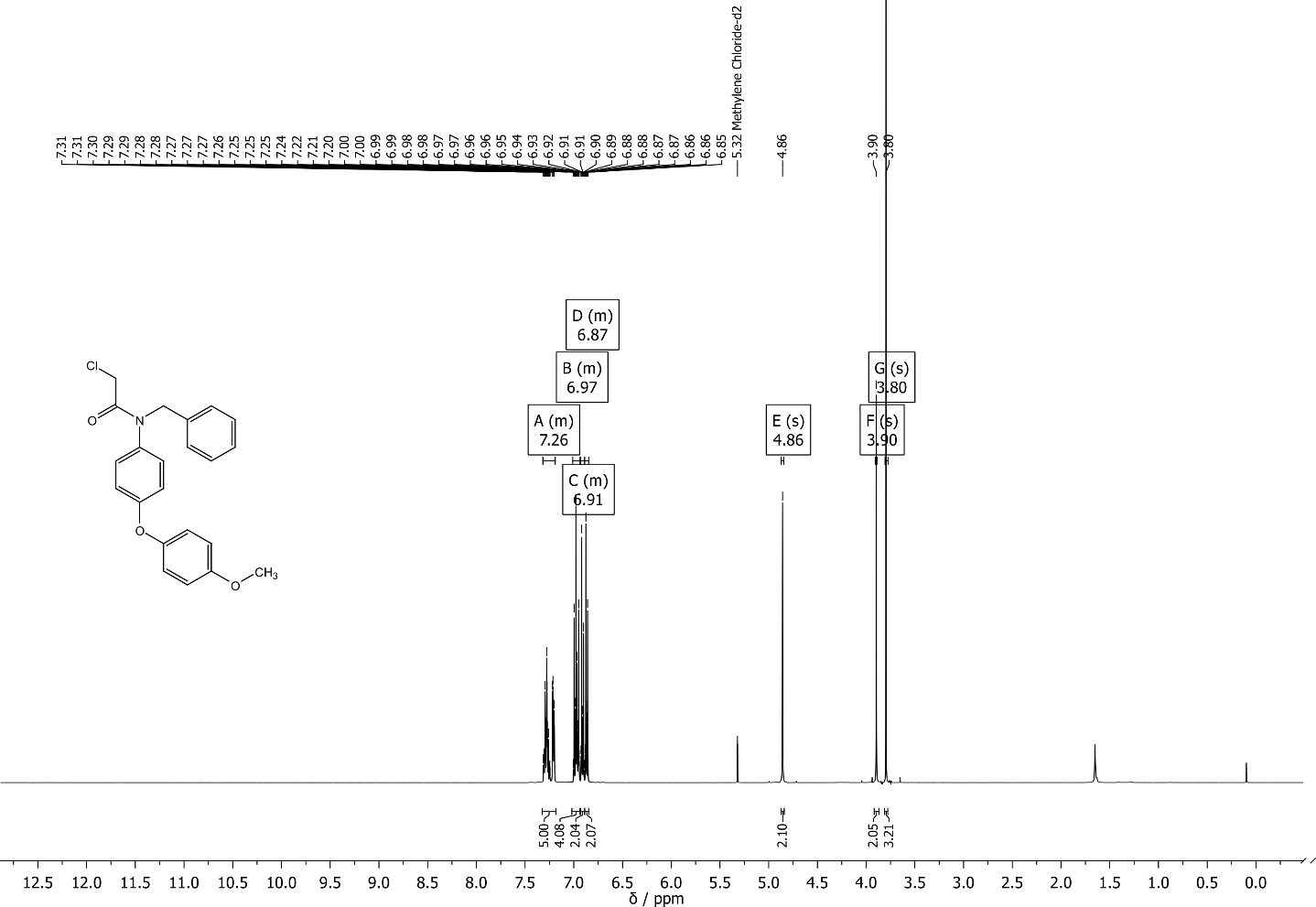

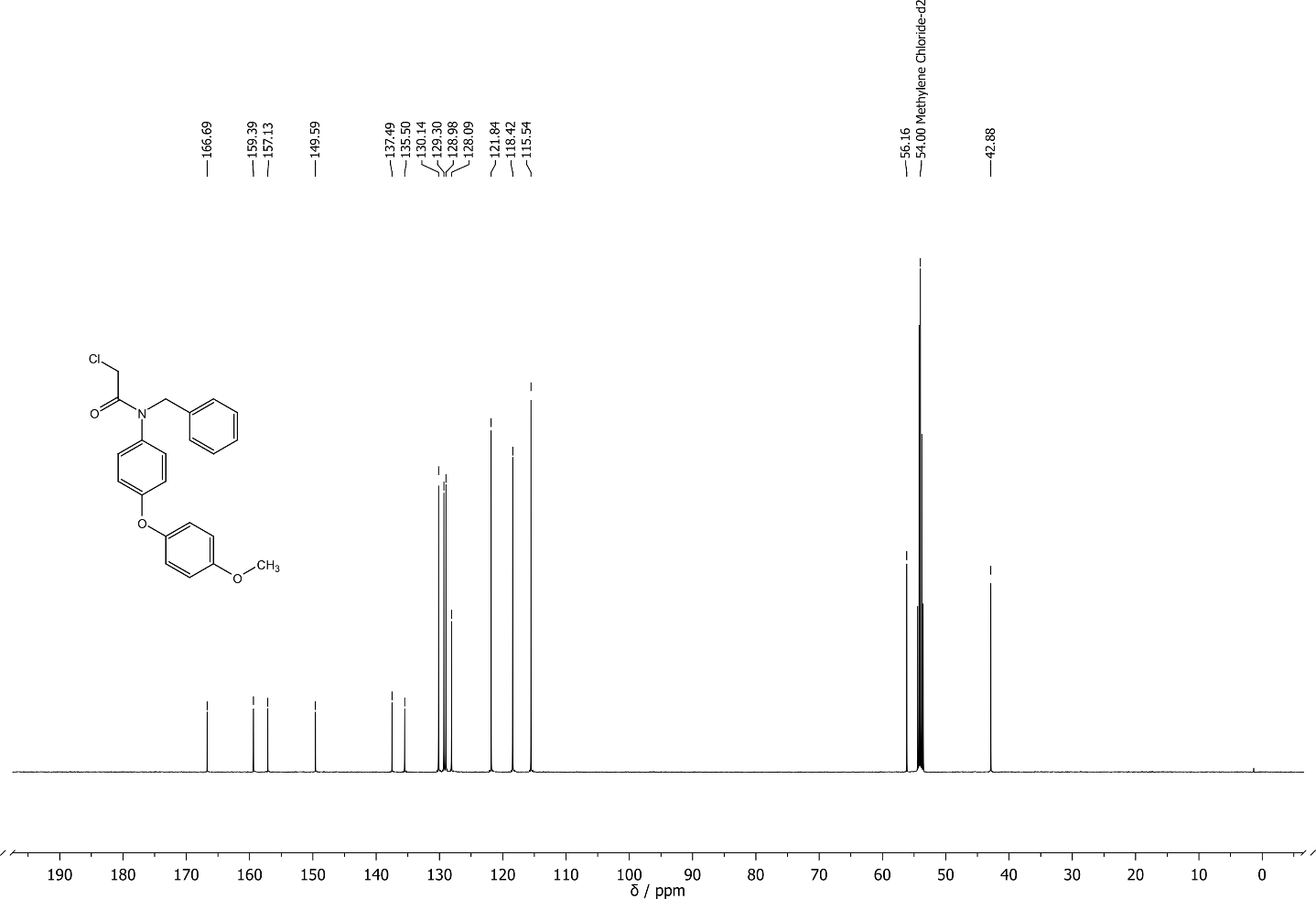

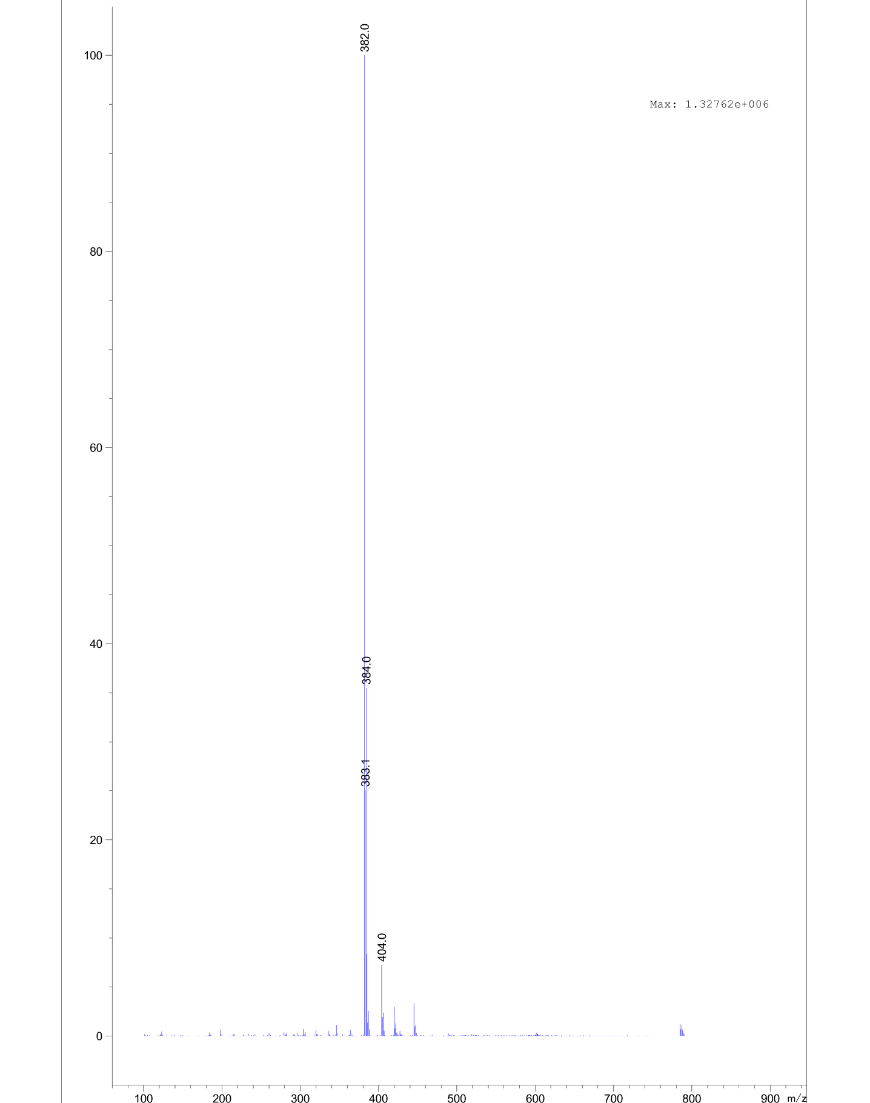

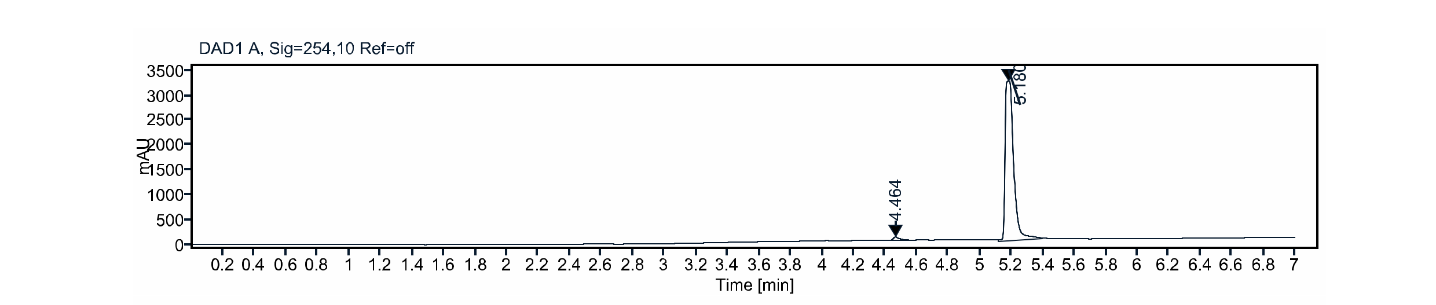

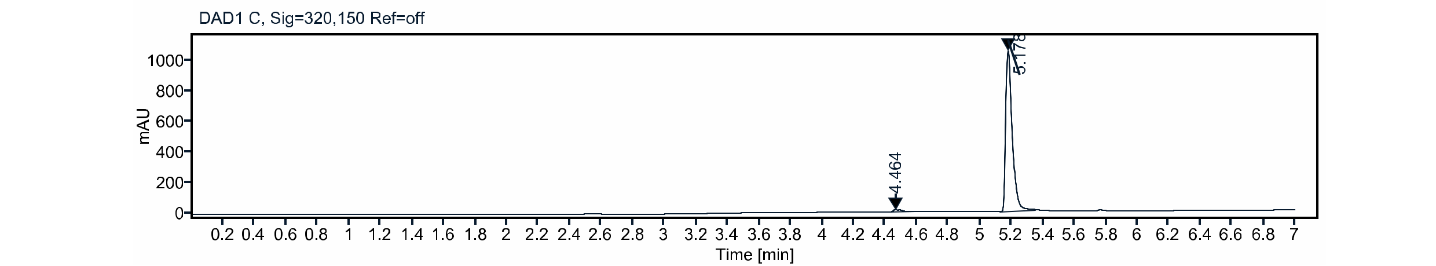

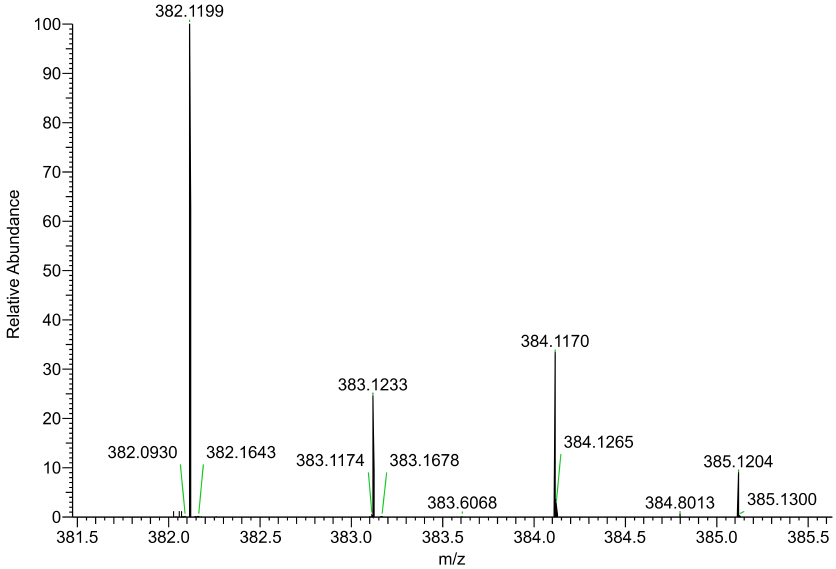

tert-butyl (2-(2-(4-(4-(benzylamino)phenoxy)phenoxy)ethoxy)ethyl)carbamate **S11**

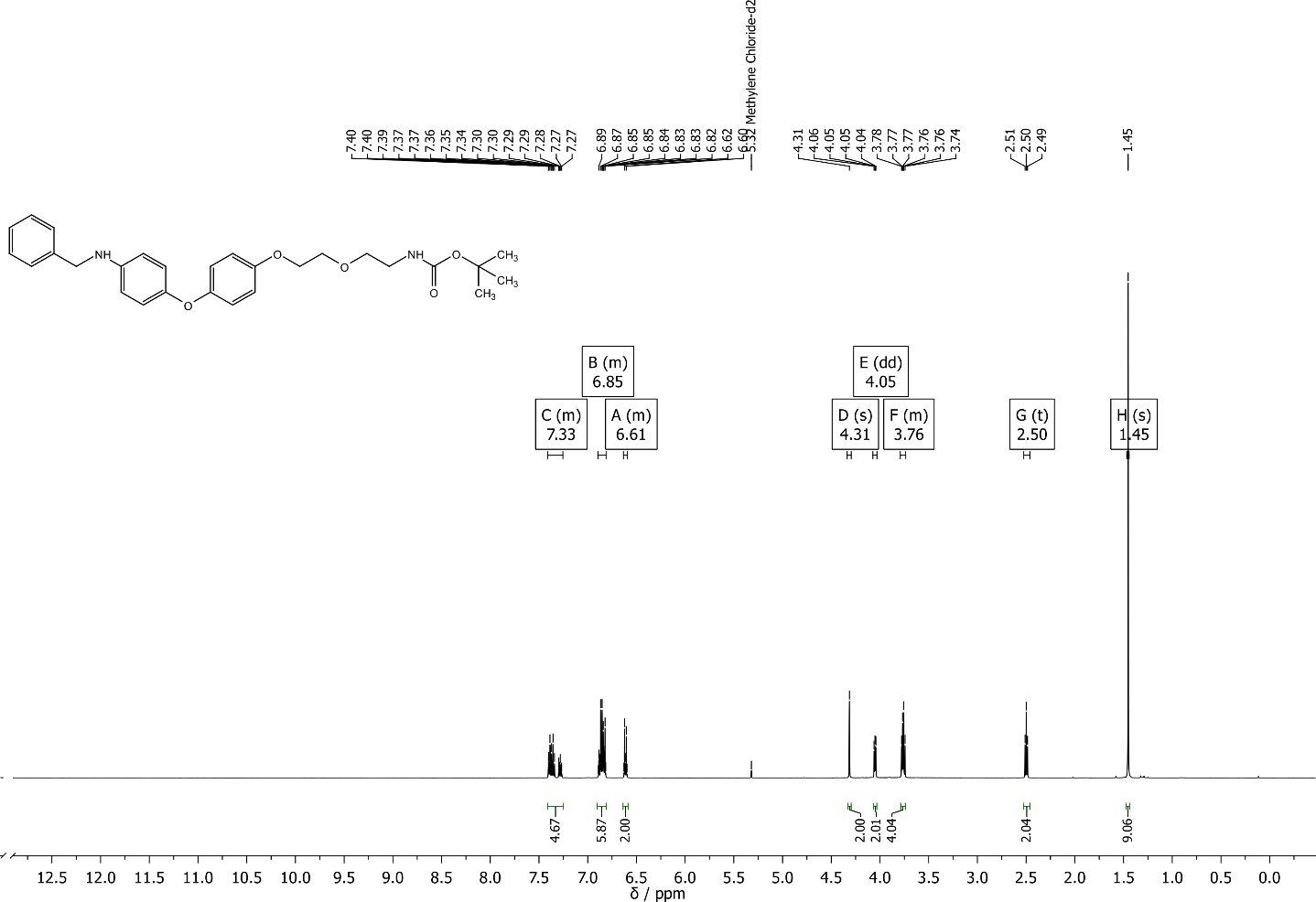

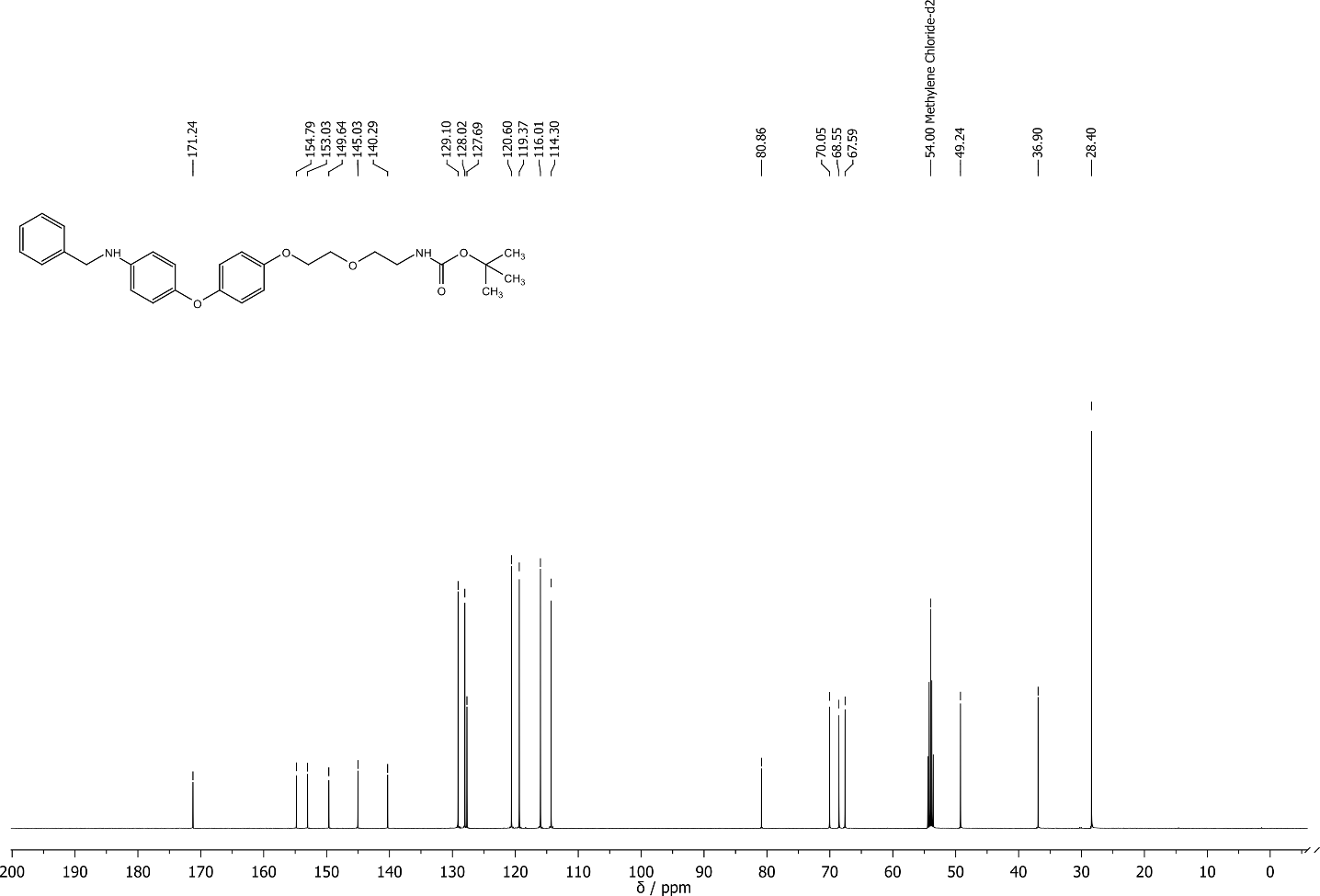

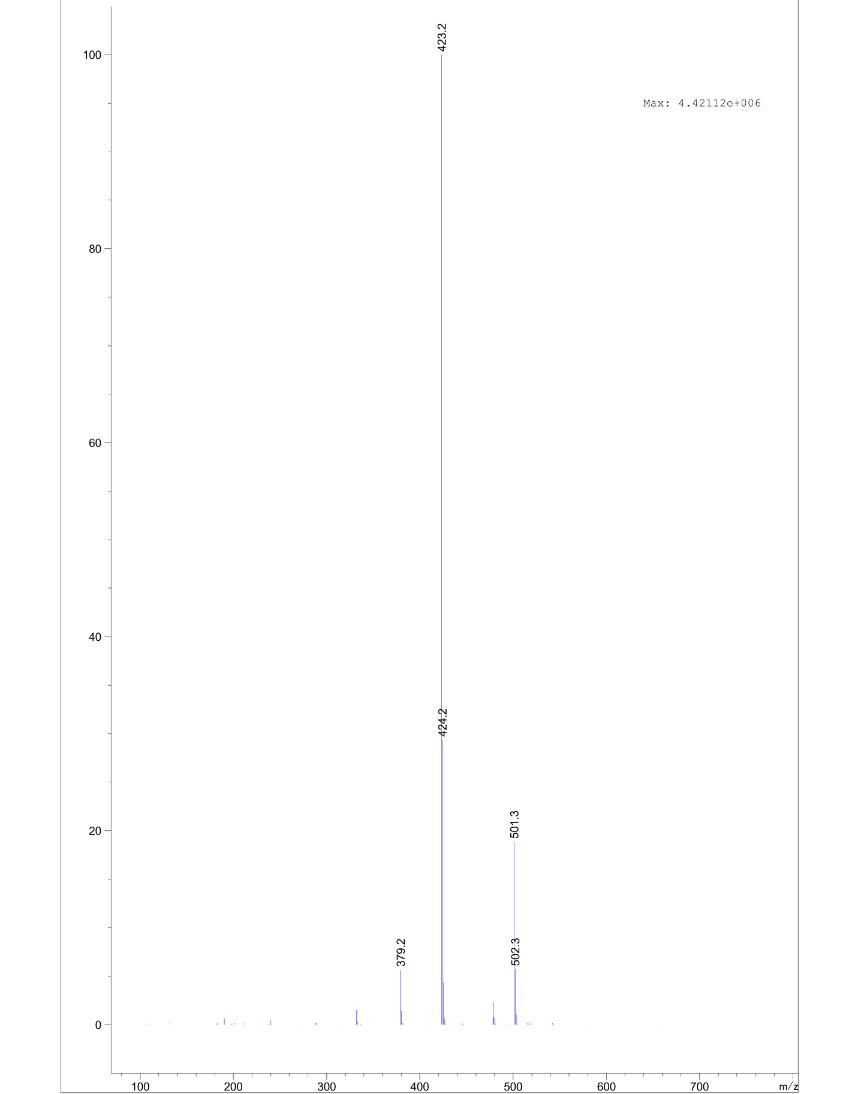

tert-butyl (2-(2-(2-(2-(4-(4-(benzylamino)phenoxy)phenoxy)ethoxy)ethoxy)ethoxy)ethyl)carbamate **S12**

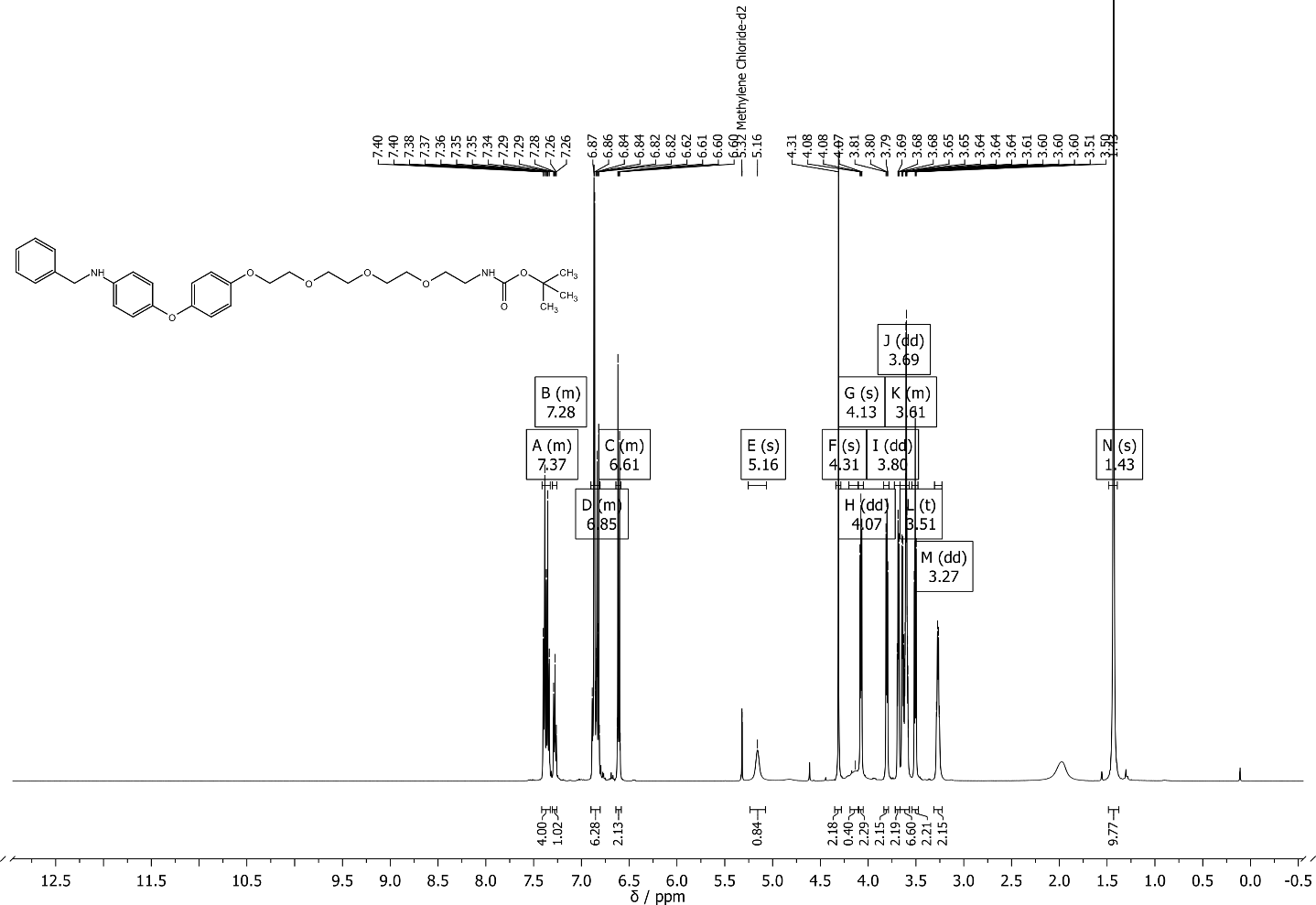

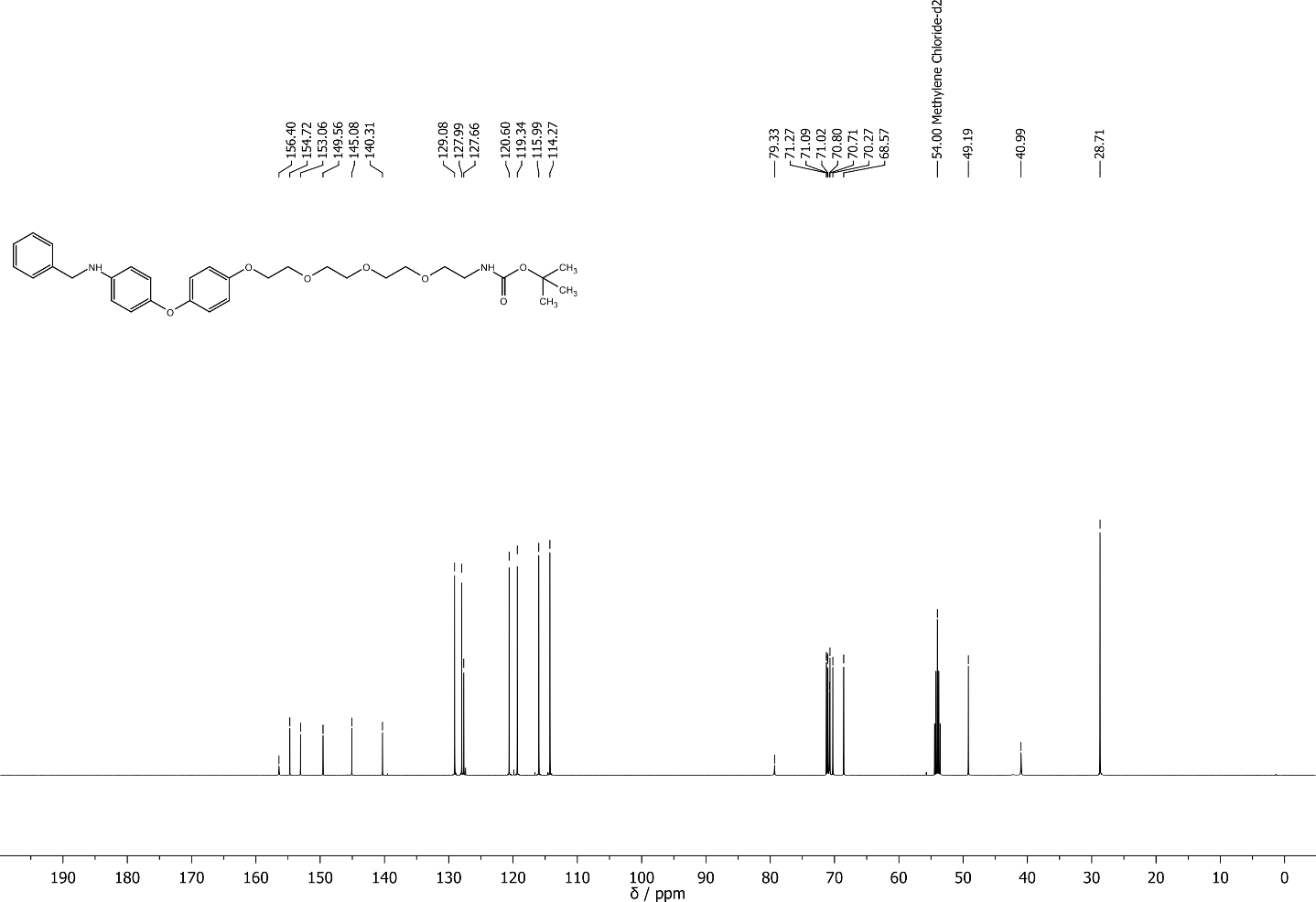

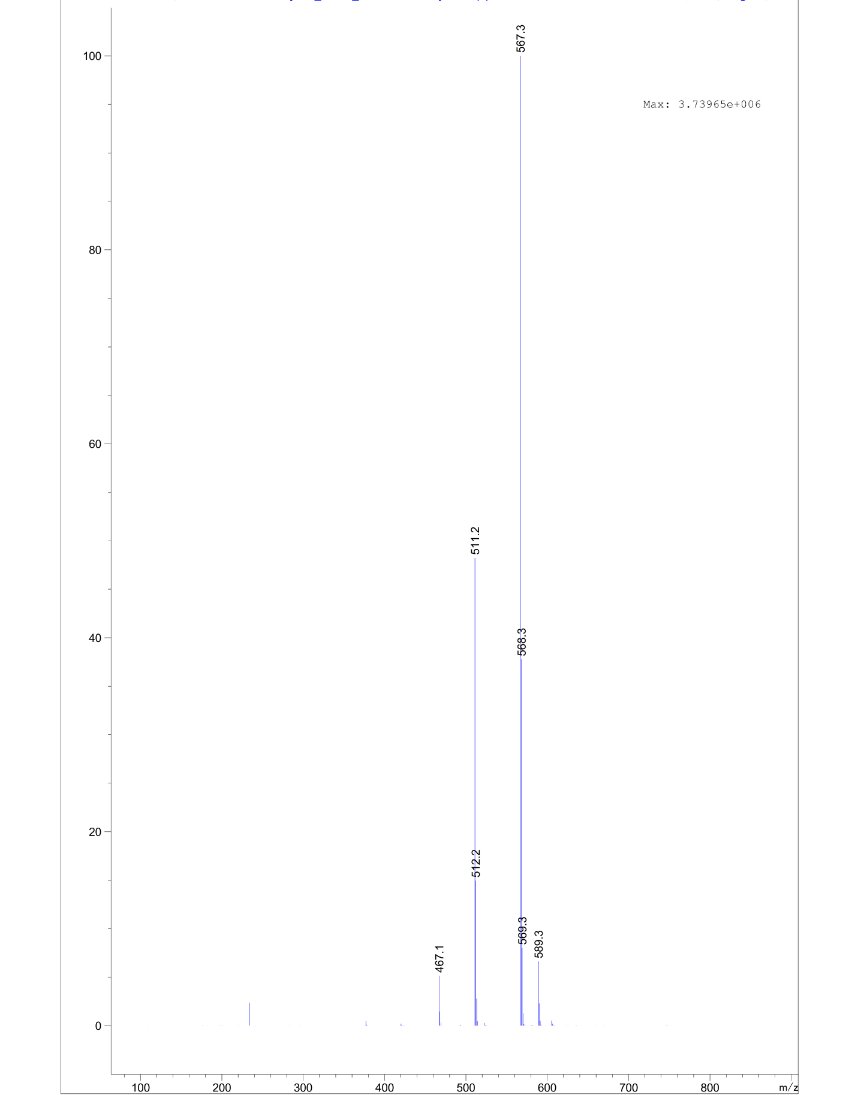

N-(2-(2-(4-(4-(benzylamino)phenoxy)phenoxy)ethoxy)ethyl)-2-((2-(2,6-dioxopiperidin-3-yl)-1,3-dioxoisoindolin-4-yl)oxy)acetamide **S13**

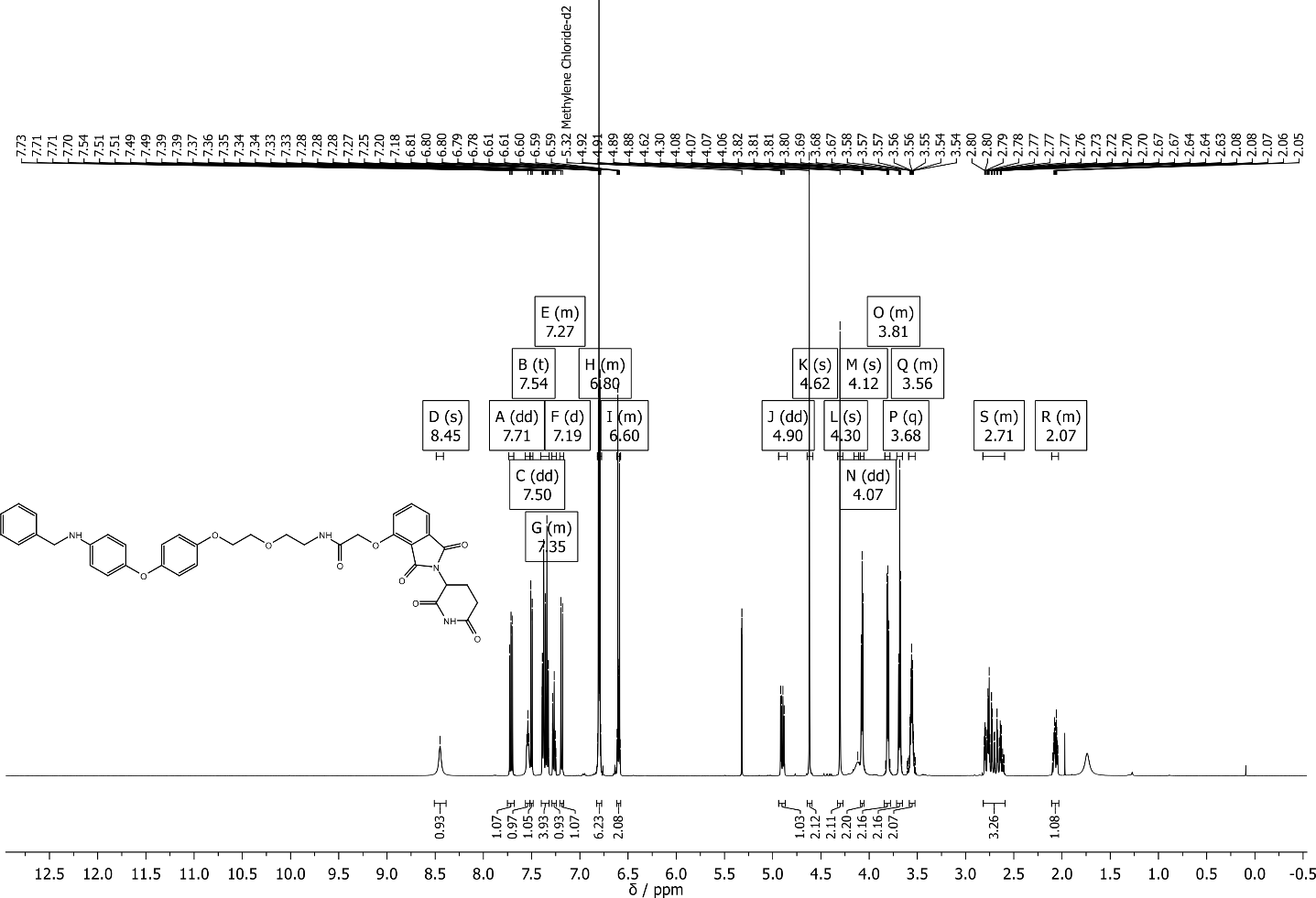

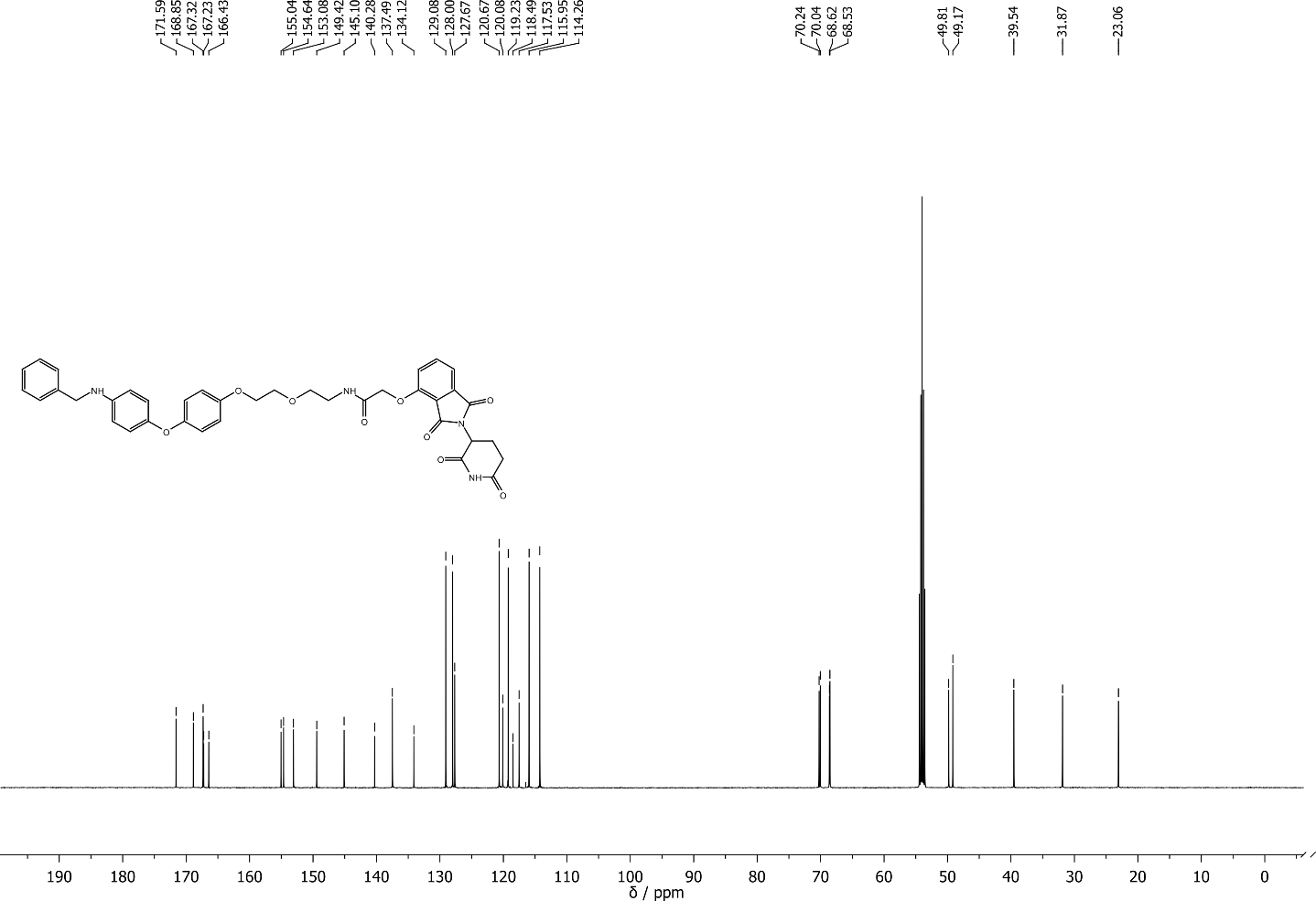

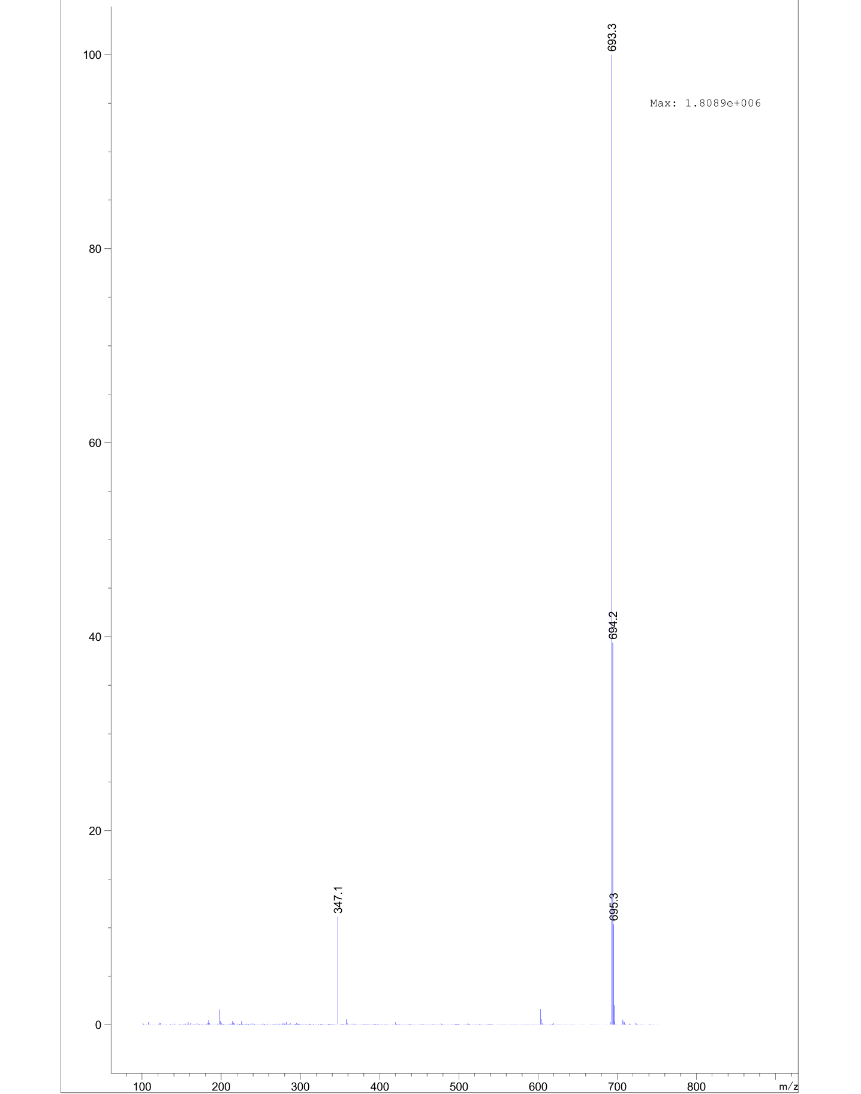

N-(2-(2-(2-(2-(4-(4-(benzylamino)phenoxy)phenoxy)ethoxy)ethoxy)ethoxy)ethyl)-2-((2-(2,6-dioxopiperidin-3-yl)-1,3-dioxoisoindolin-4-yl)oxy)acetamide **S14**

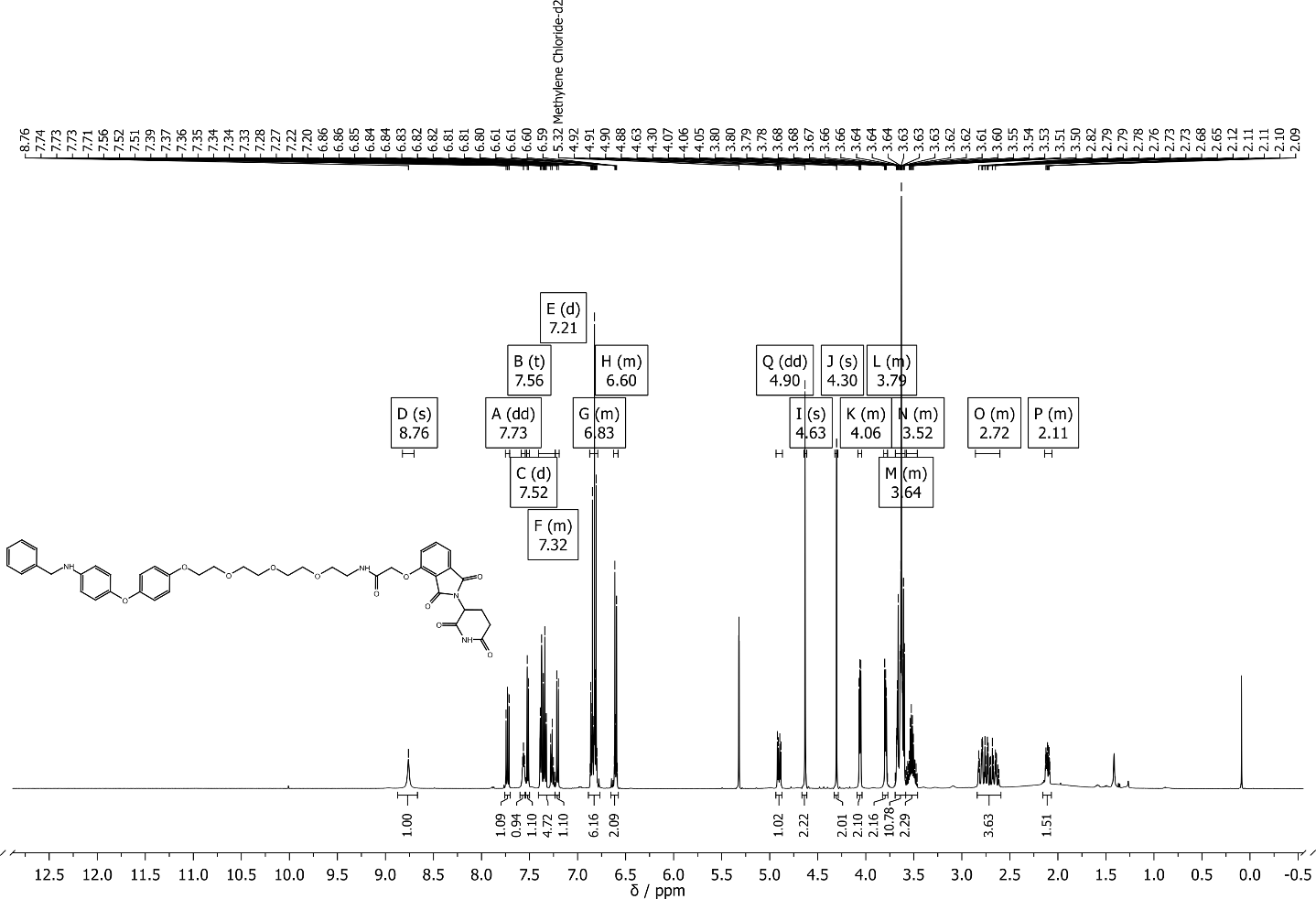

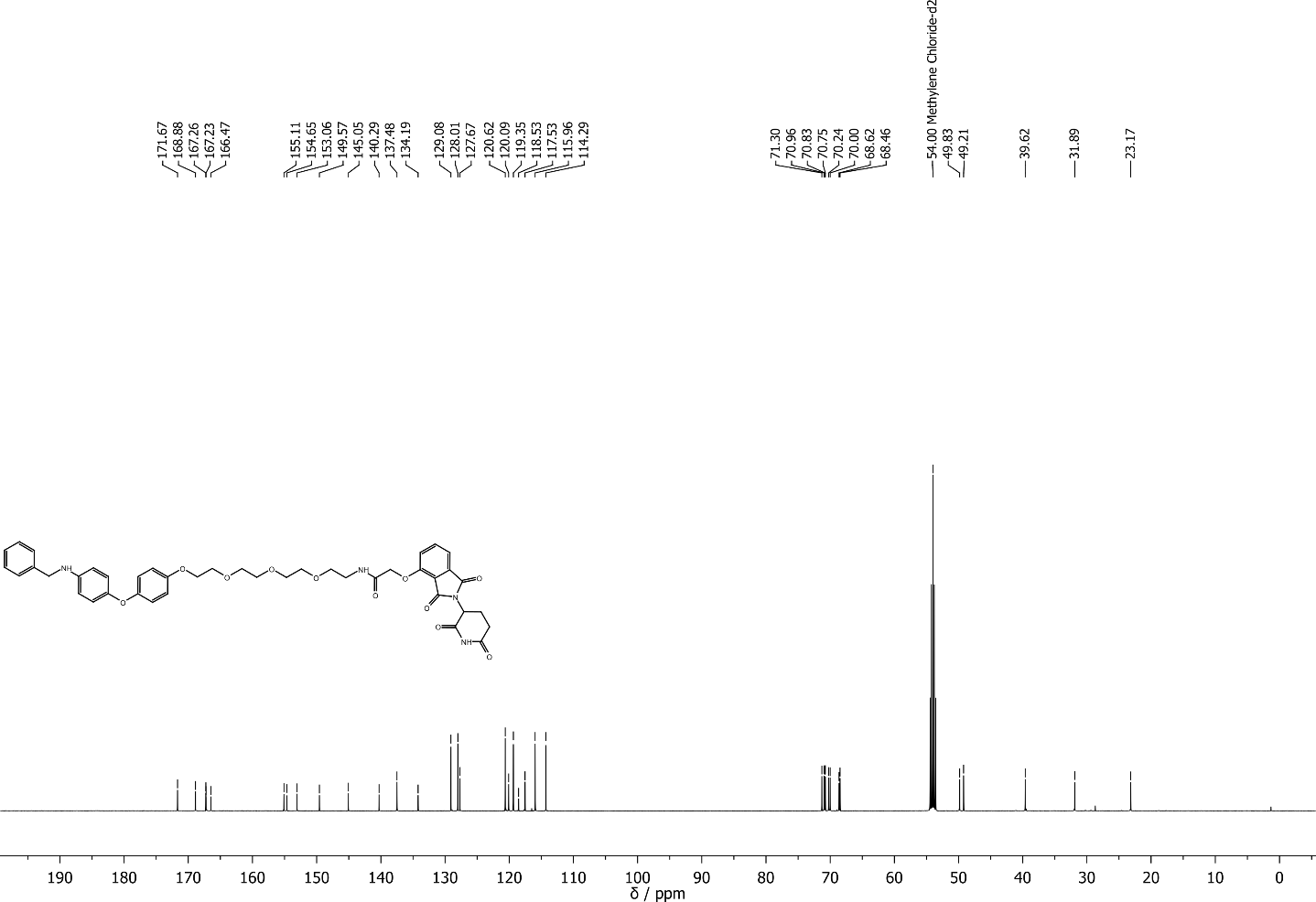

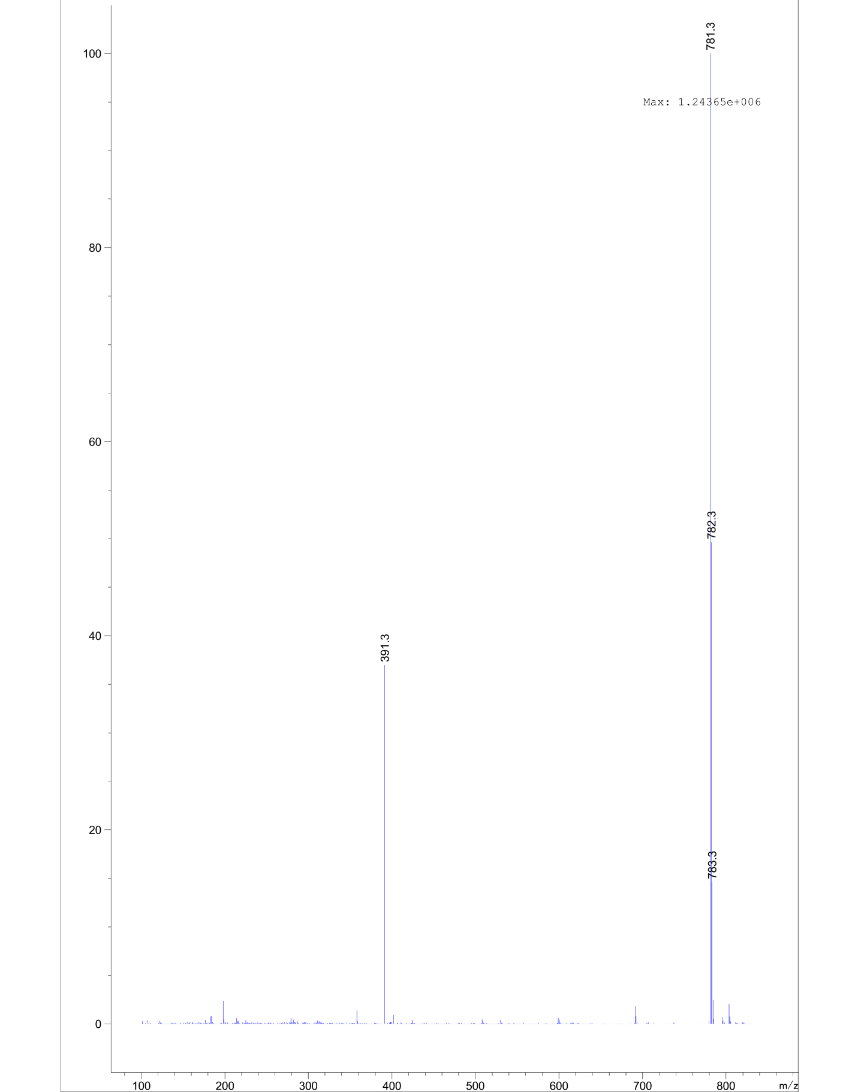

N-(2-(2-(2-(2-(4-(4-(benzylamino)phenoxy)phenoxy)ethoxy)ethoxy)ethoxy)ethyl)-2-((2-(1-methyl-2,6-dioxopiperidin-3-yl)-1,3-dioxoisoindolin-4-yl)oxy)acetamide **S15**

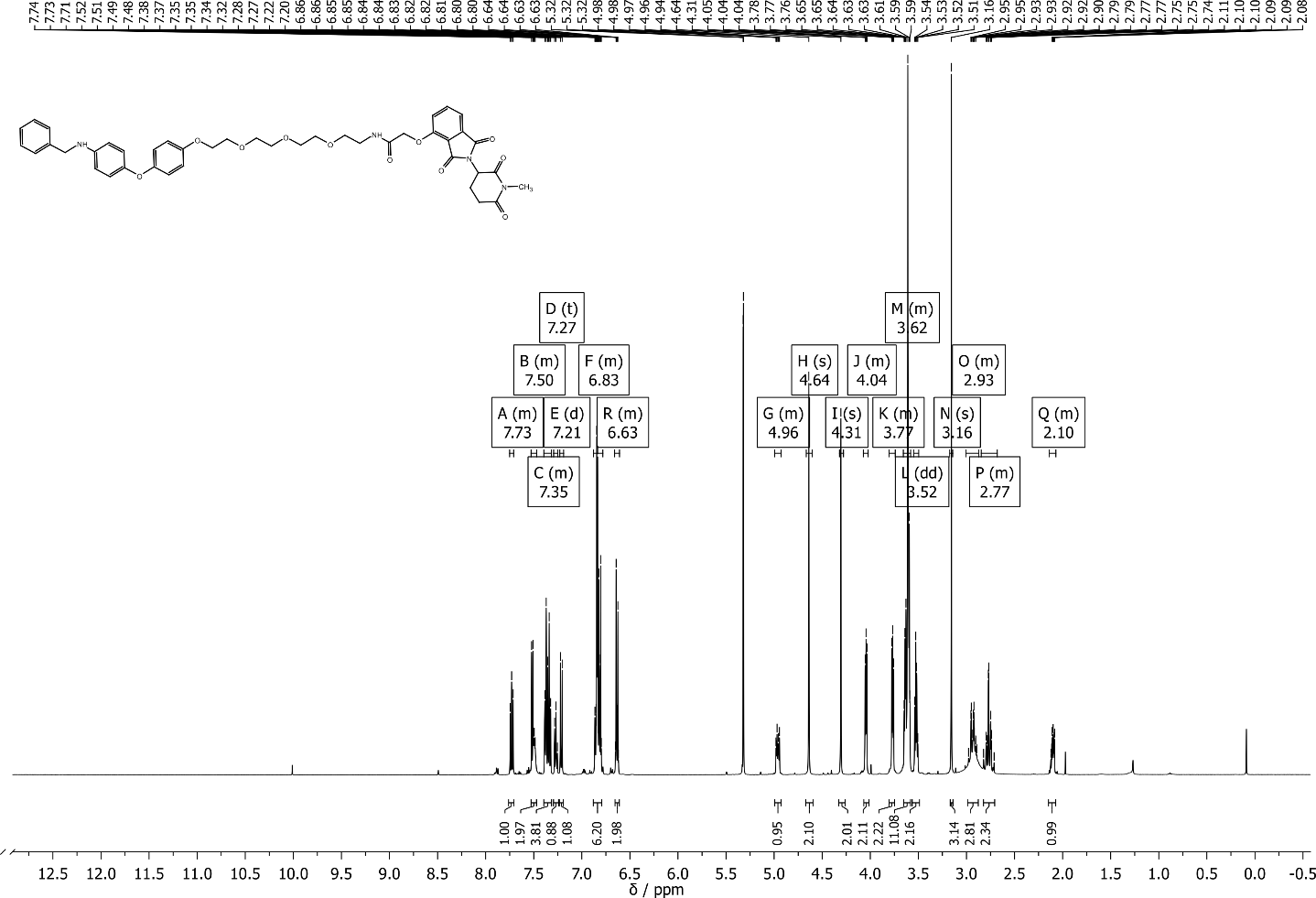

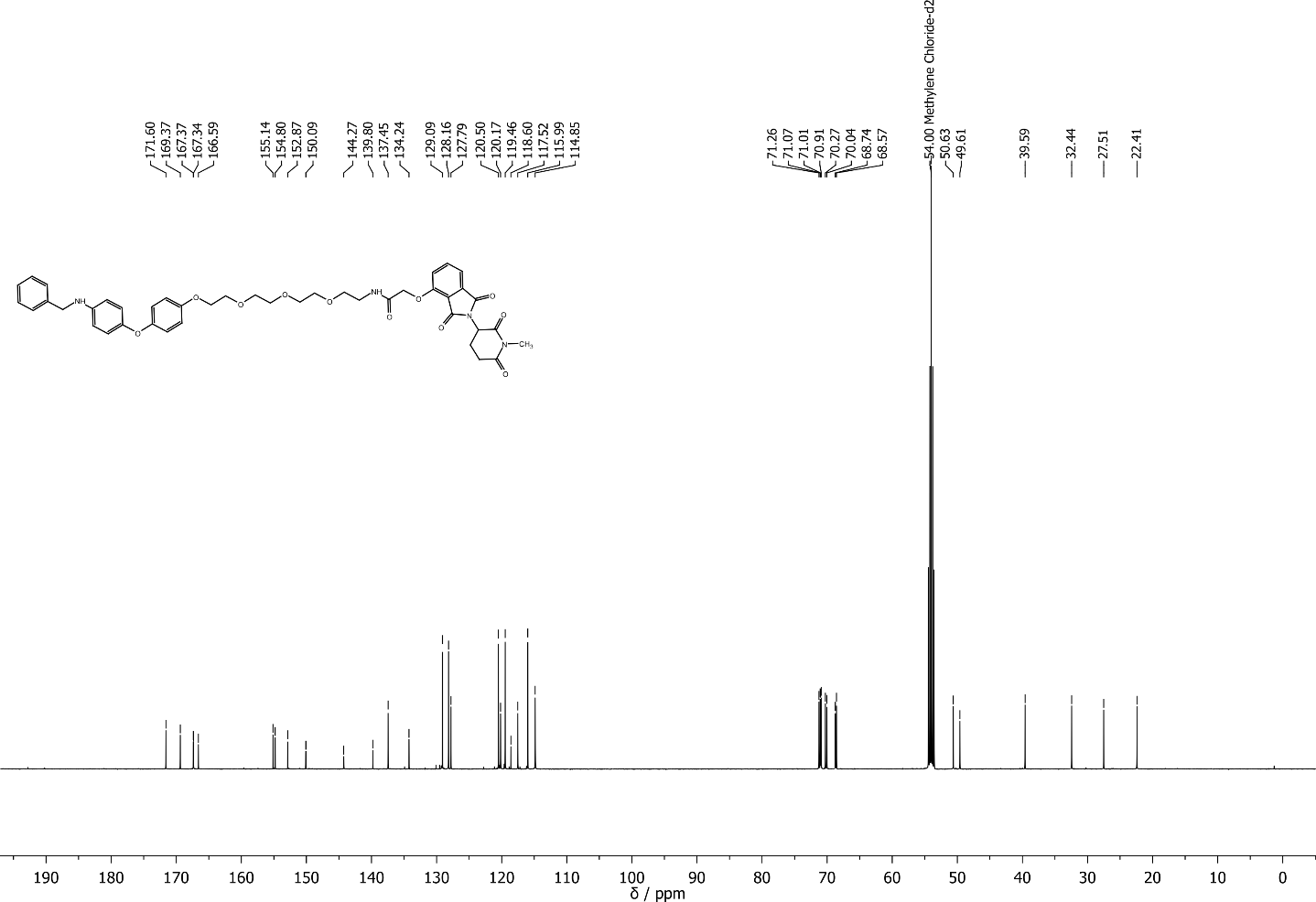

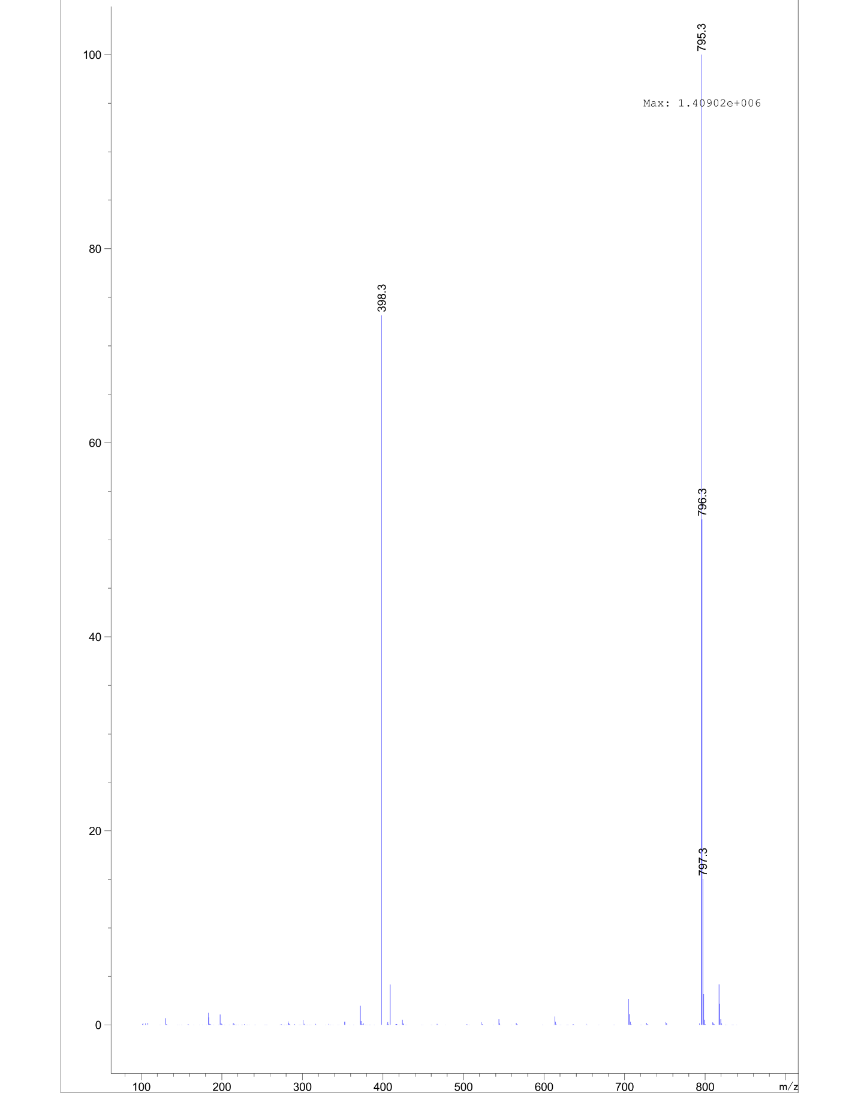

N-benzyl-2-chloro-N-(4-(4-(2-(2-(2-((2-(2,6-dioxopiperidin-3-yl)-1,3-dioxoisoindolin-4-yl)oxy)acetamido)ethoxy)ethoxy)phenoxy)phenyl)acetamide **1a**

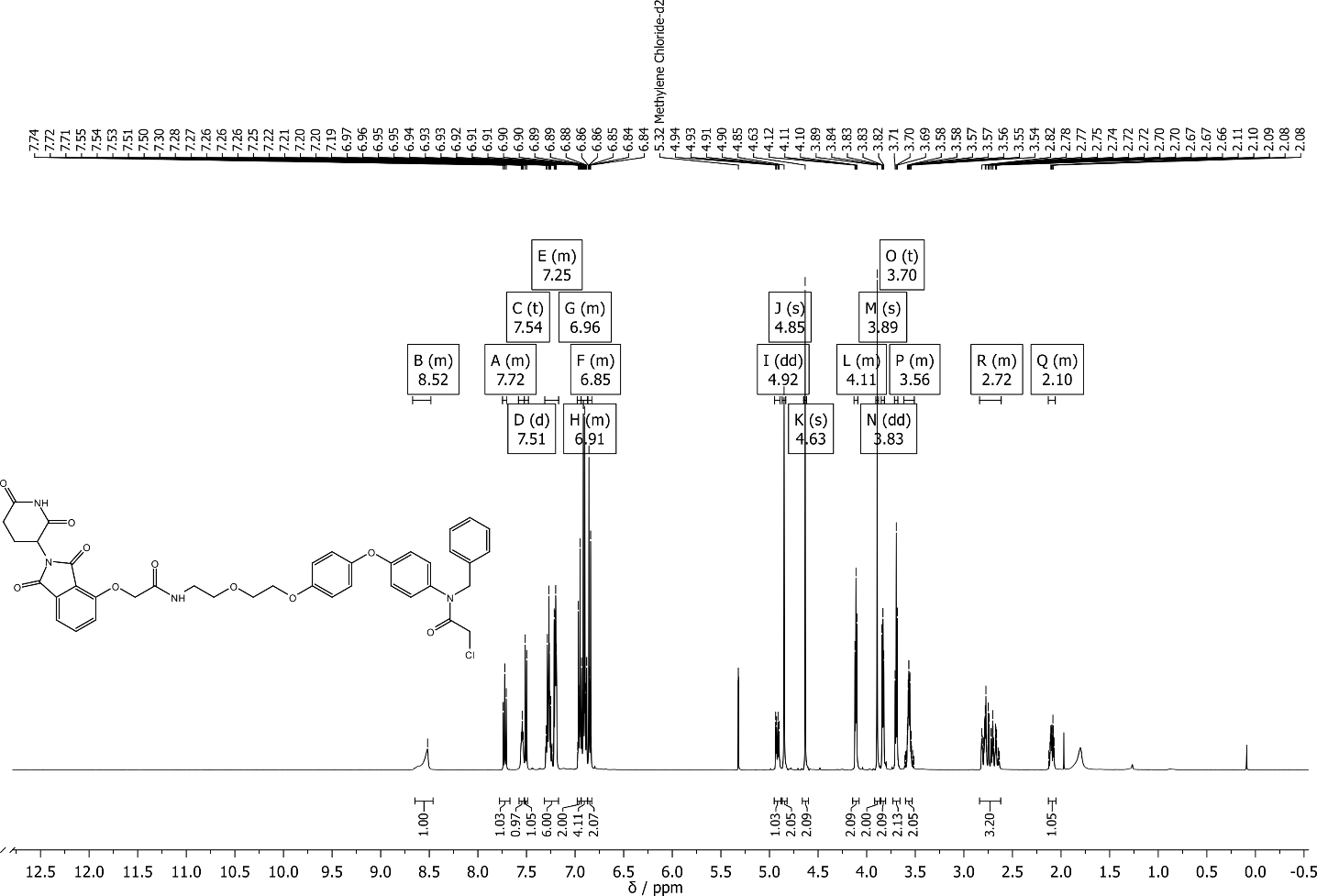

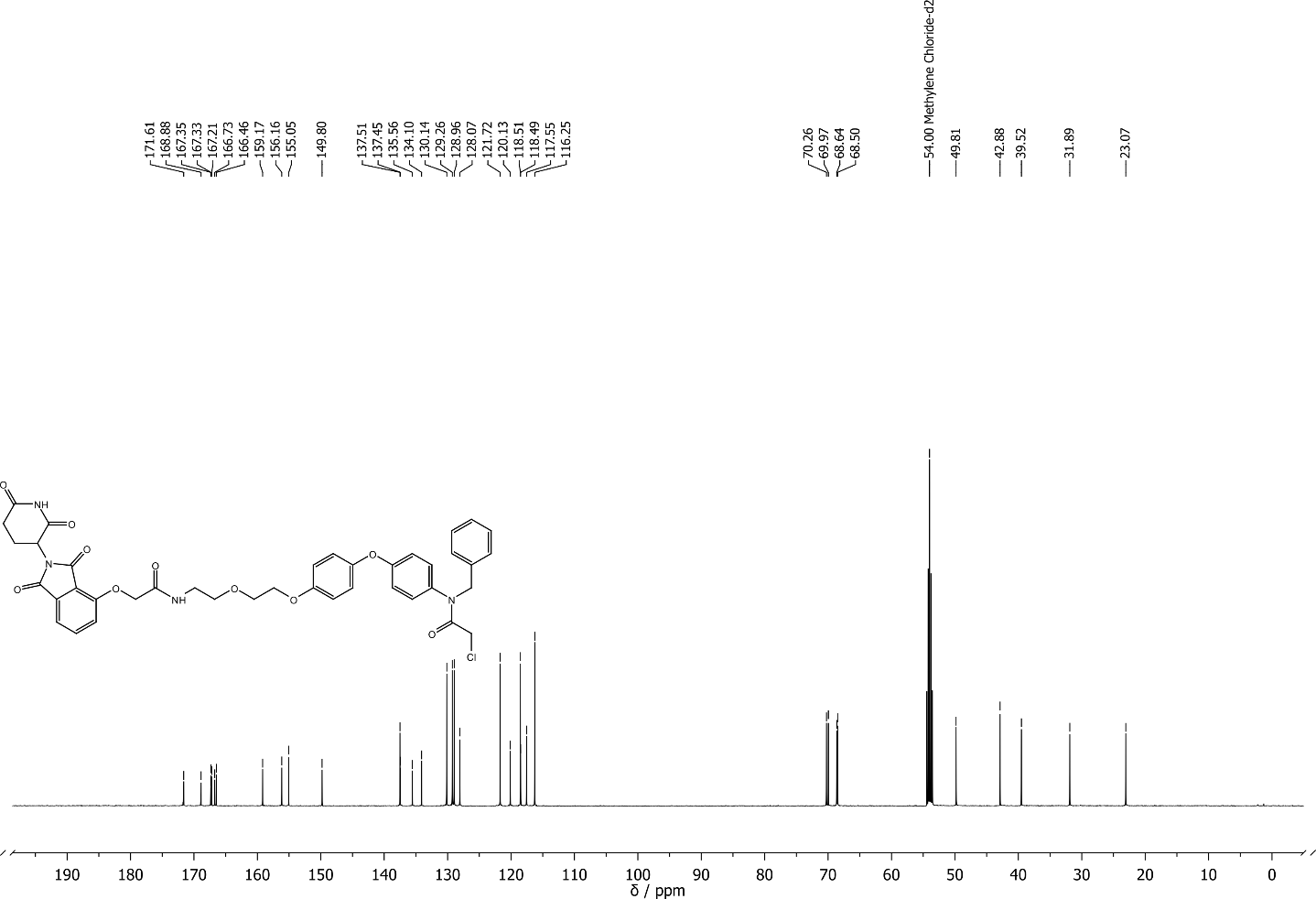

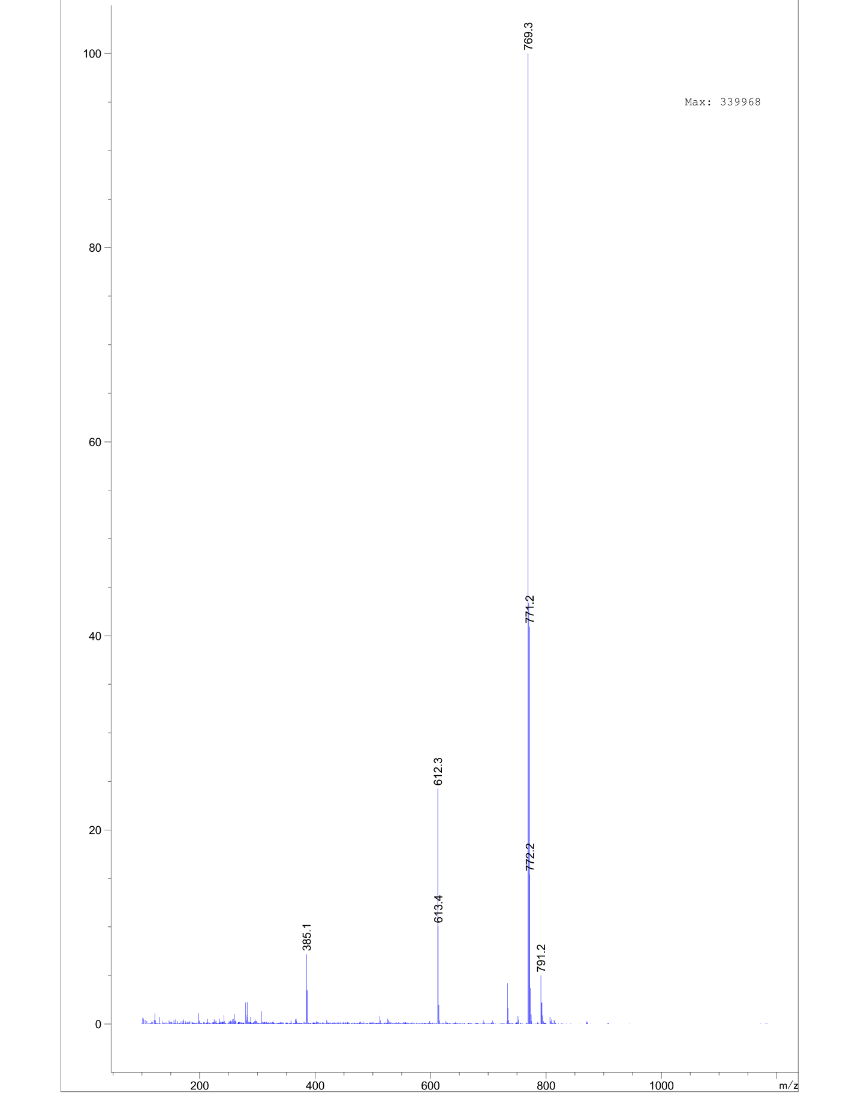

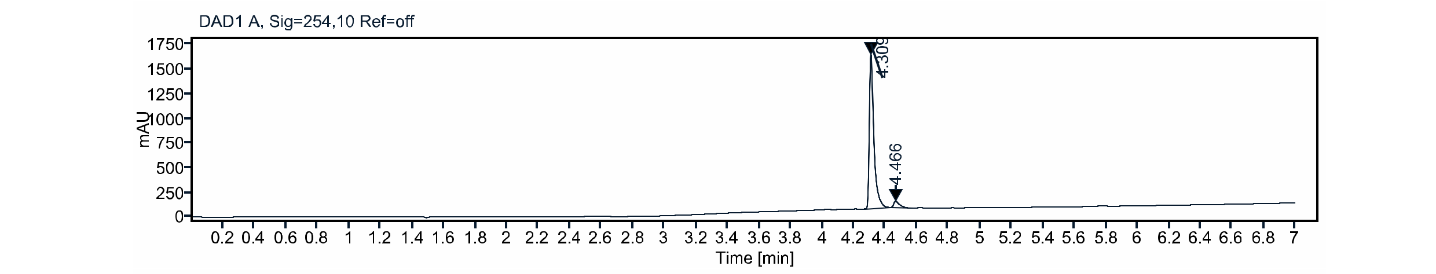

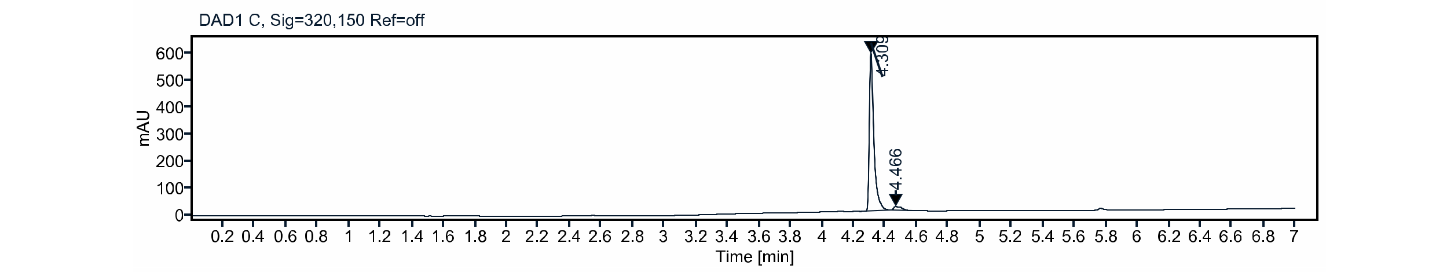

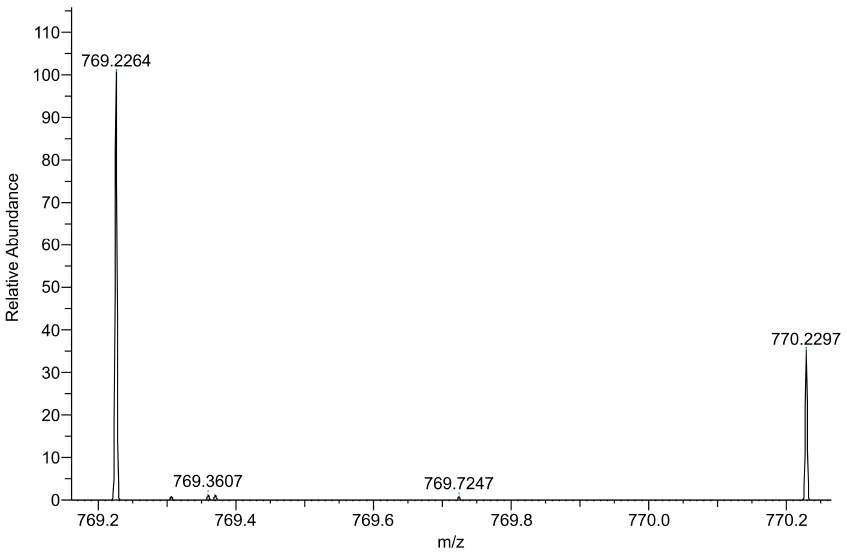

N-benzyl-2-chloro-N-(4-(4-((1-((2-(2,6-dioxopiperidin-3-yl)-1,3-dioxoisoindolin-4-yl)oxy)-2-oxo-6,9,12-trioxa-3-azatetradecan-14-yl)oxy)phenoxy)phenyl)acetamide **1b**

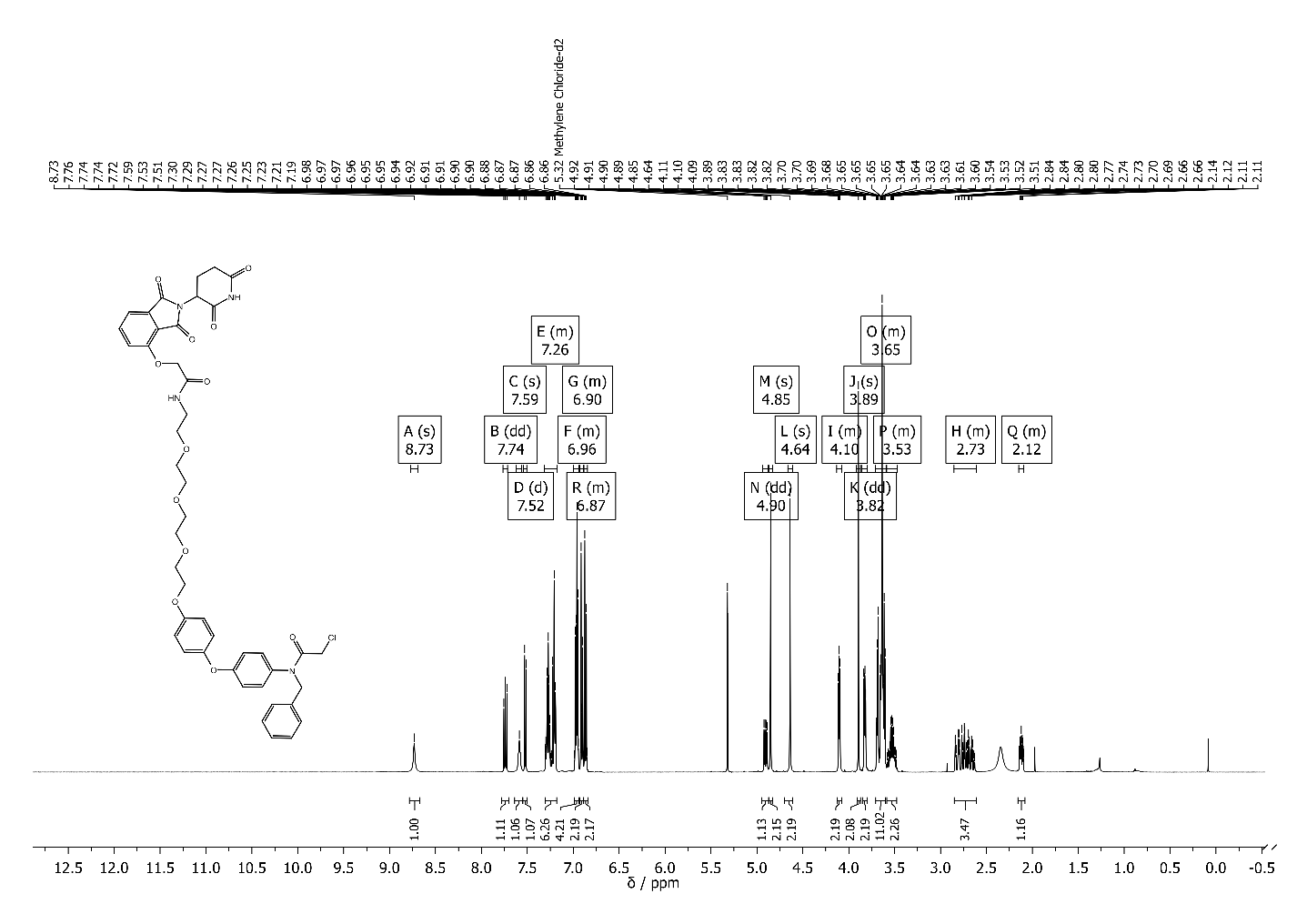

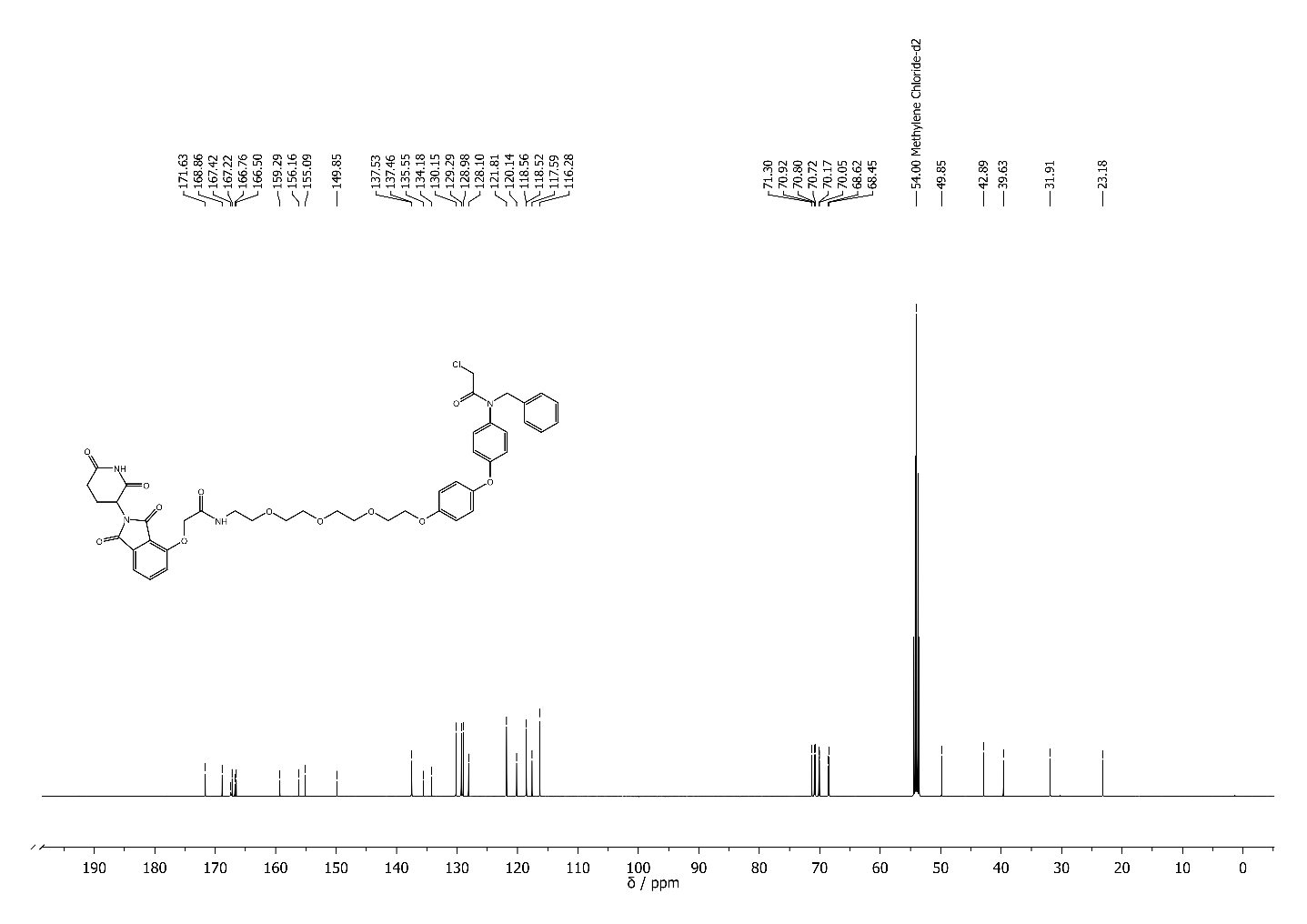

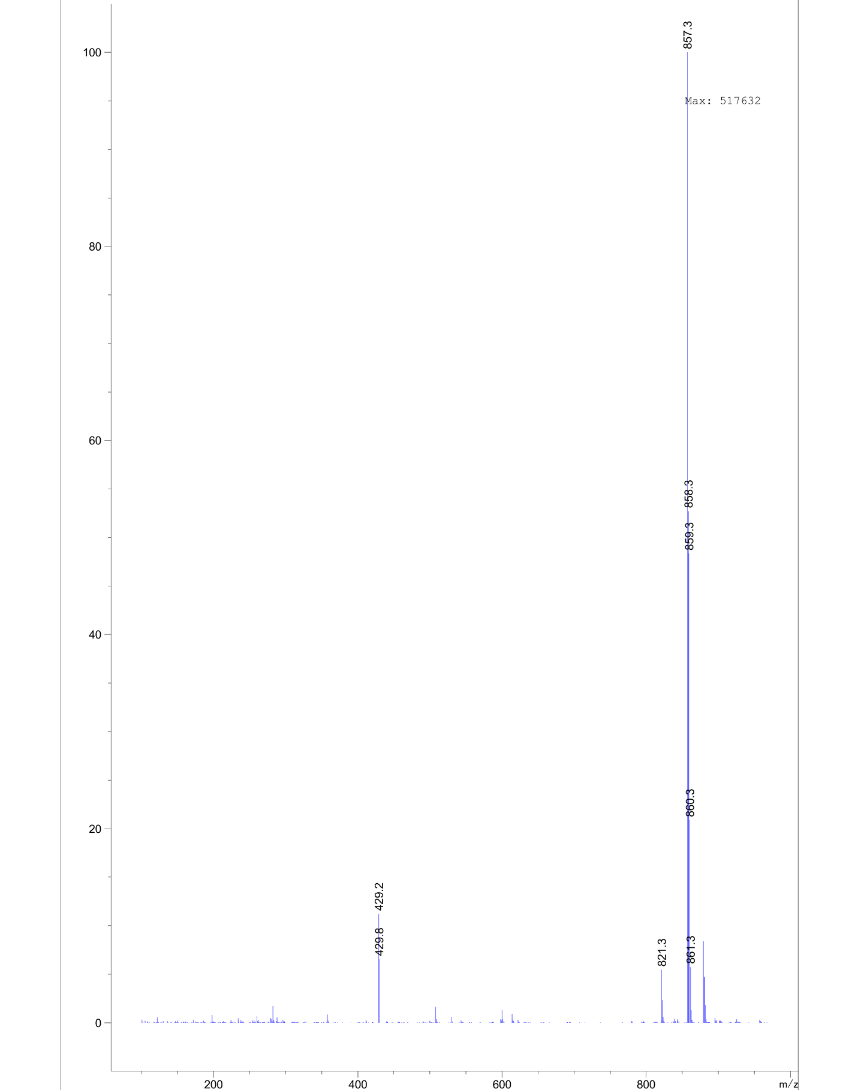

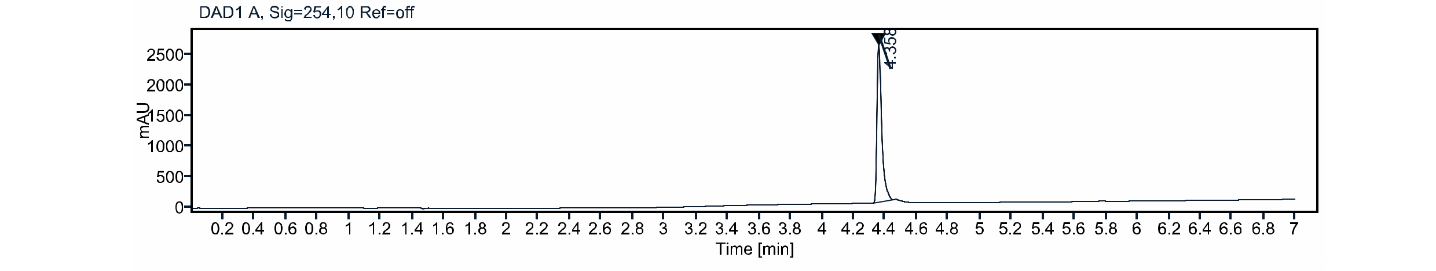

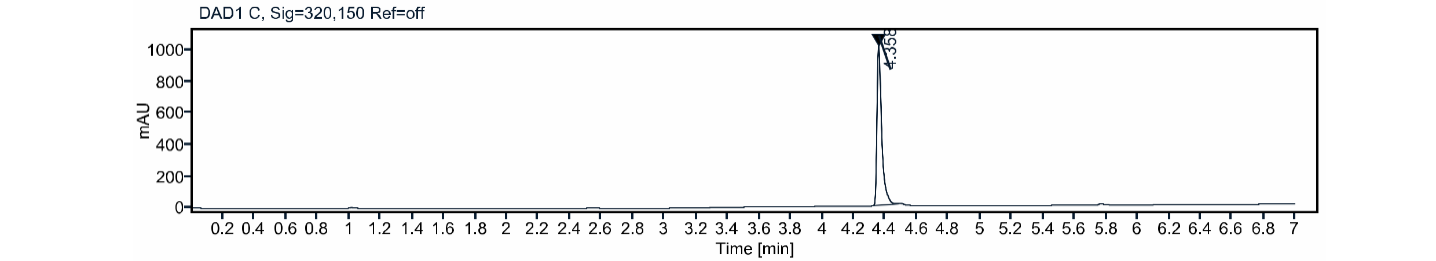

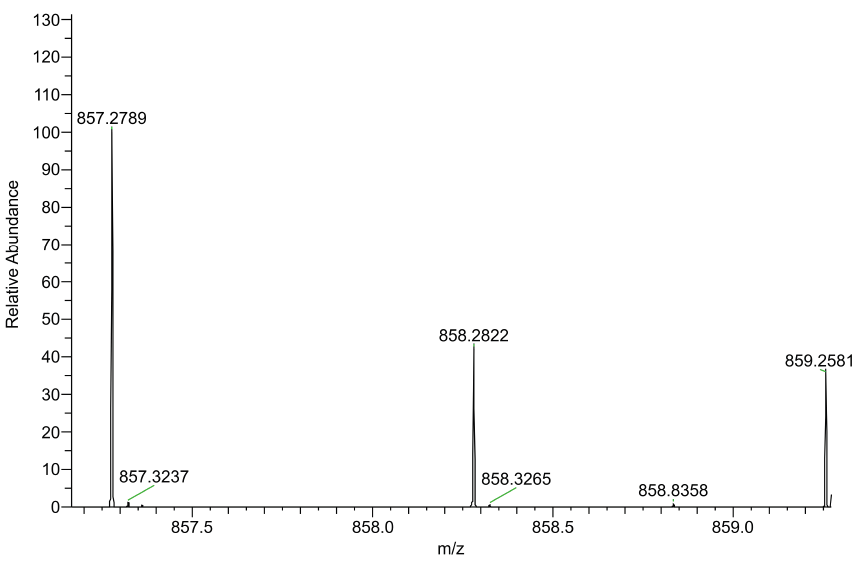

N-benzyl-2-chloro-N-(4-(4-((1-((2-(1-methyl-2,6-dioxopiperidin-3-yl)-1,3-dioxoisoindolin-4-yl)oxy)-2-oxo-6,9,12-trioxa-3-azatetradecan-14-yl)oxy)phenoxy)phenyl)acetamide **1b n.c.**

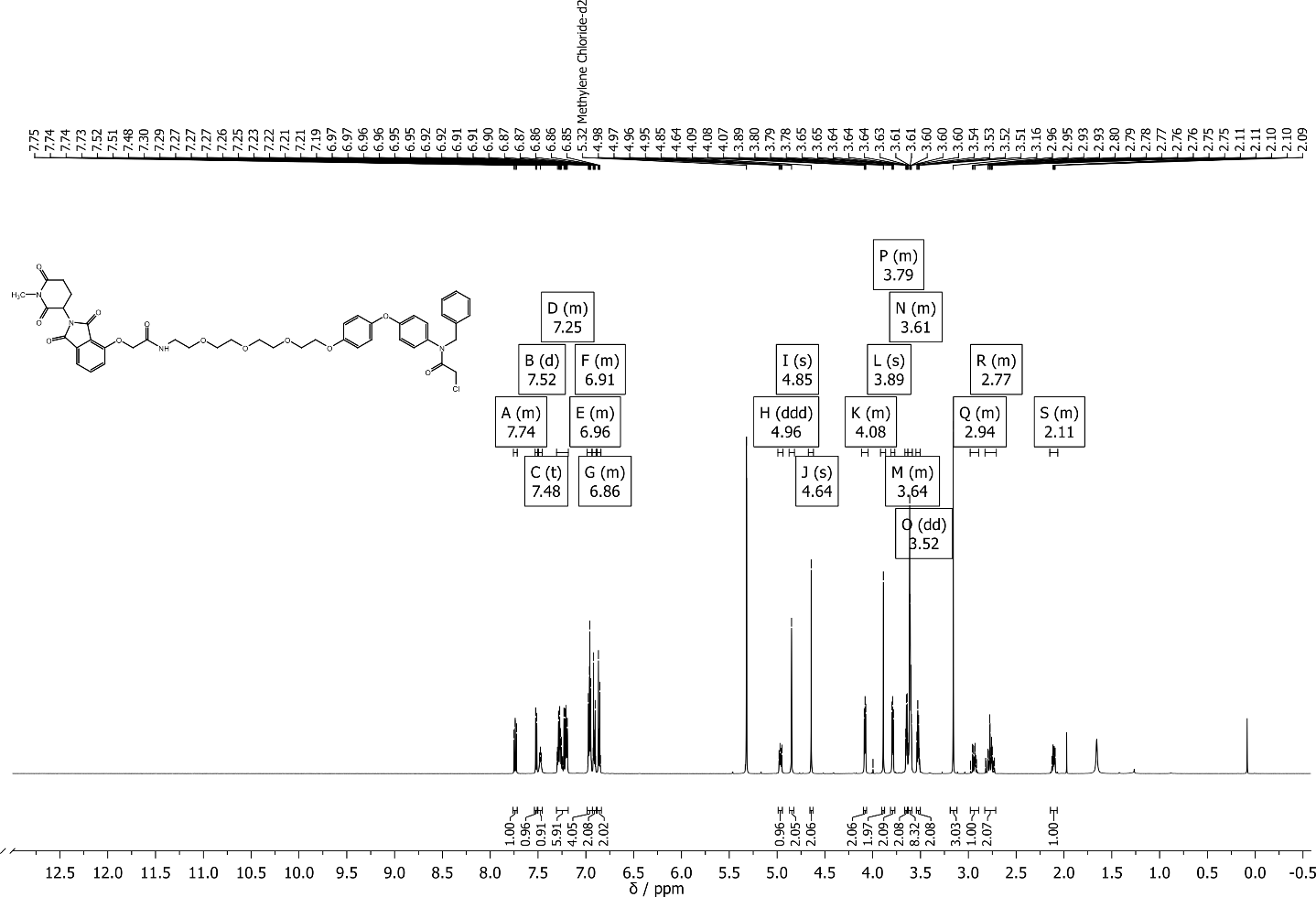

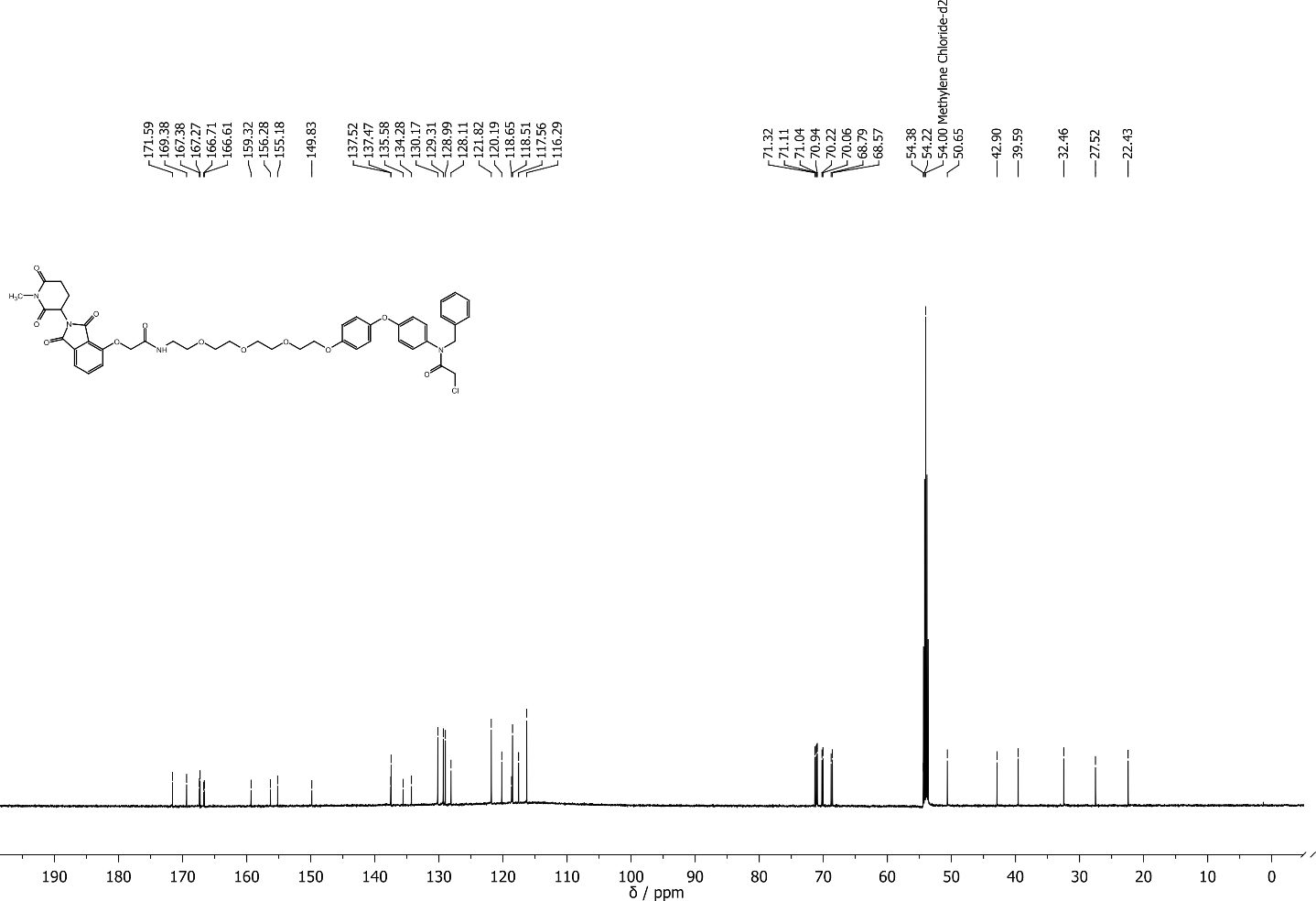

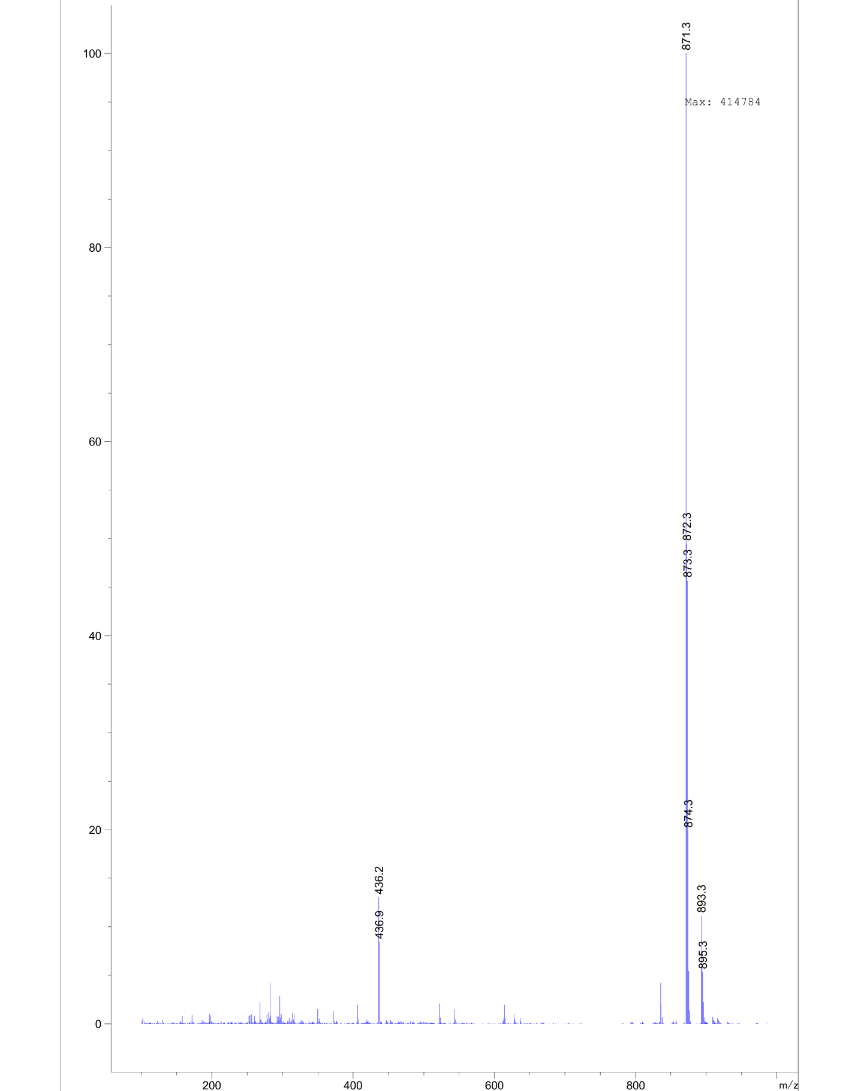

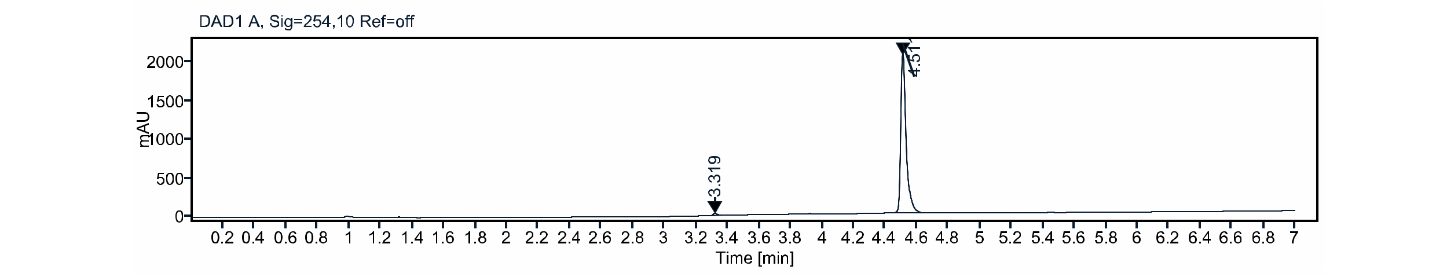

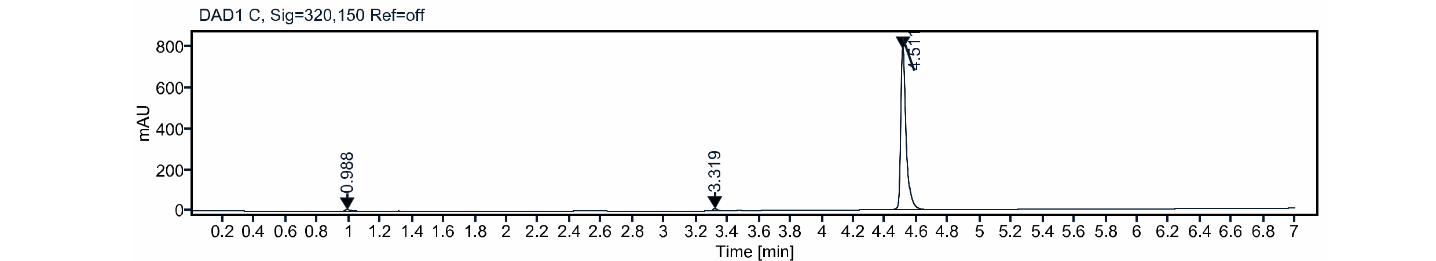

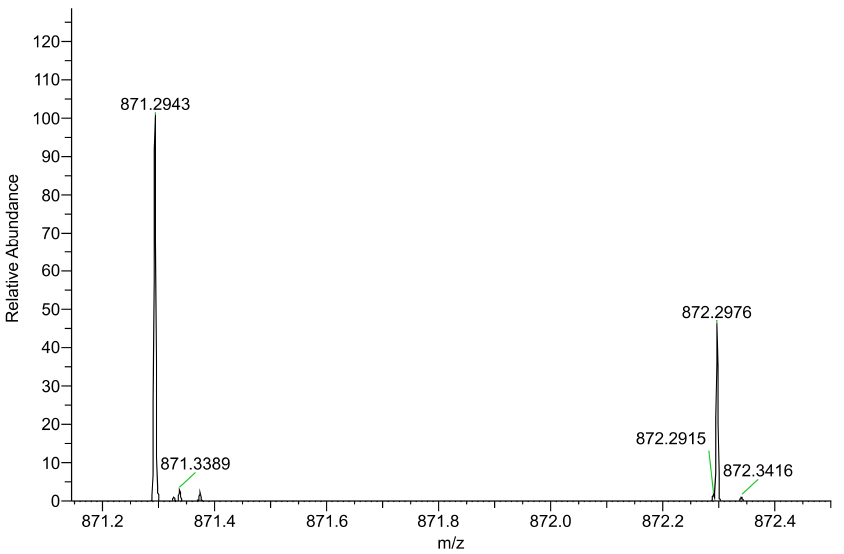

tert-butyl (5-(2-((2-(2,6-dioxopiperidin-3-yl)-1,3-dioxoisoindolin-4-yl)oxy)acetamido)pentyl)carbamate **S16**

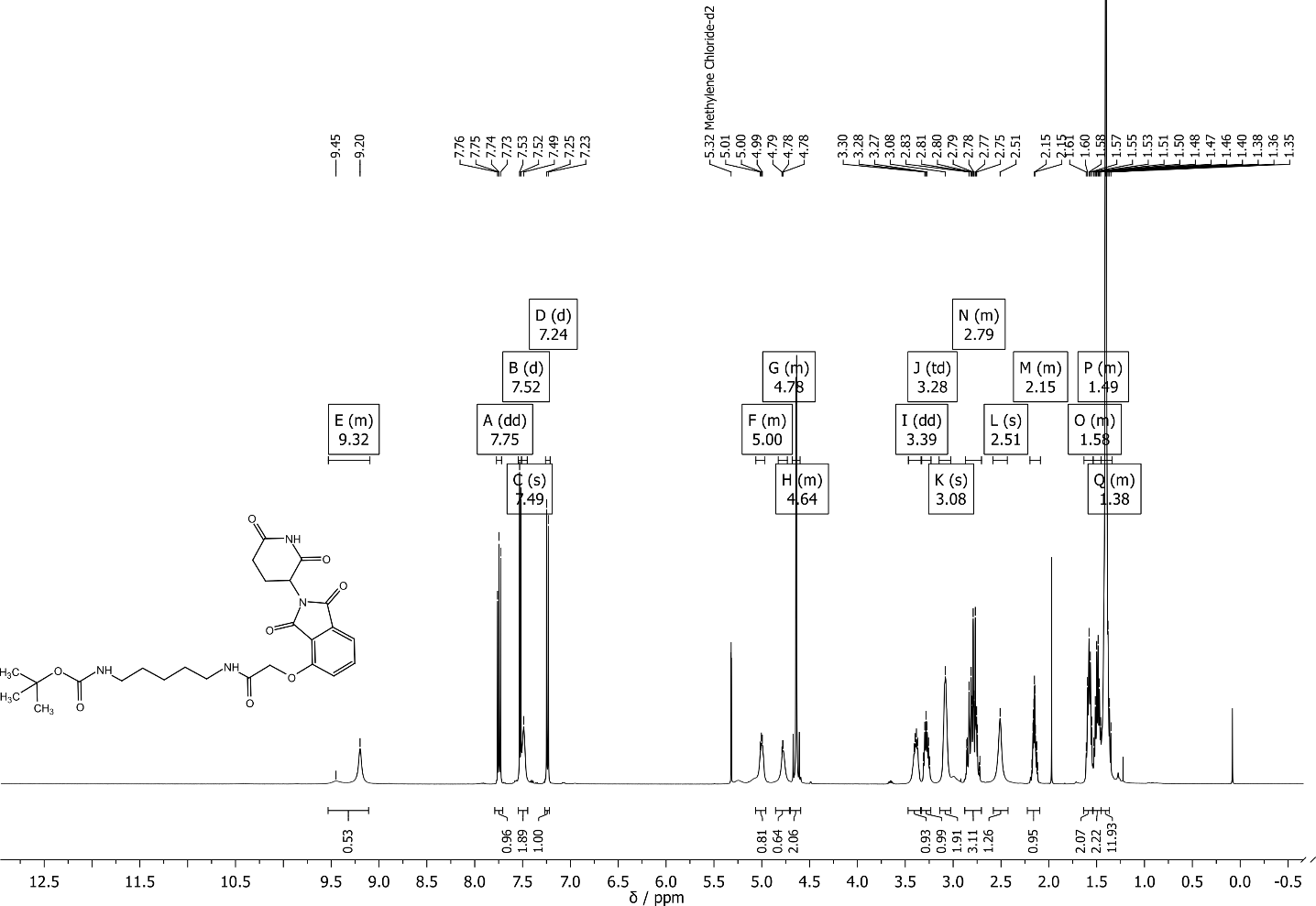

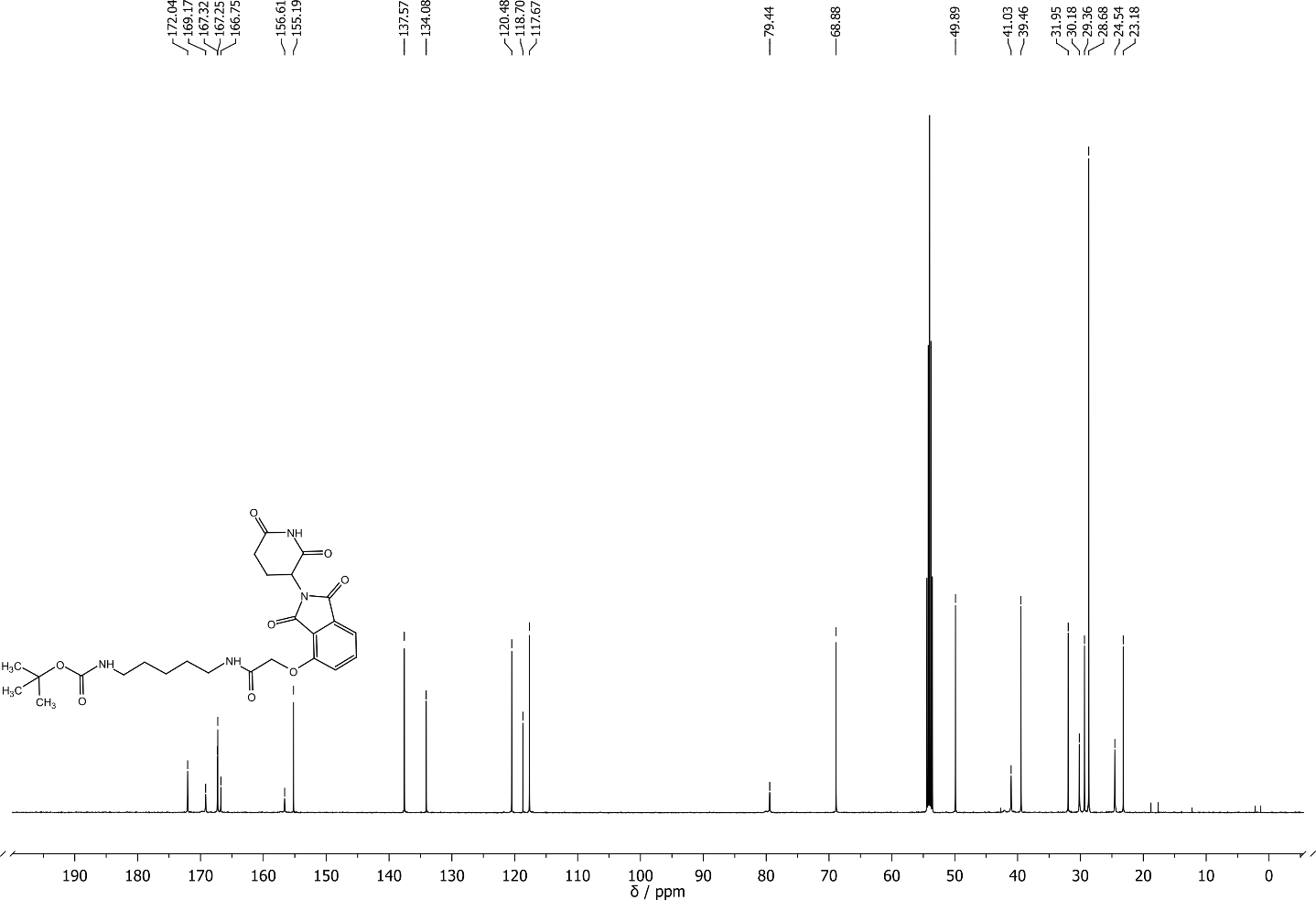

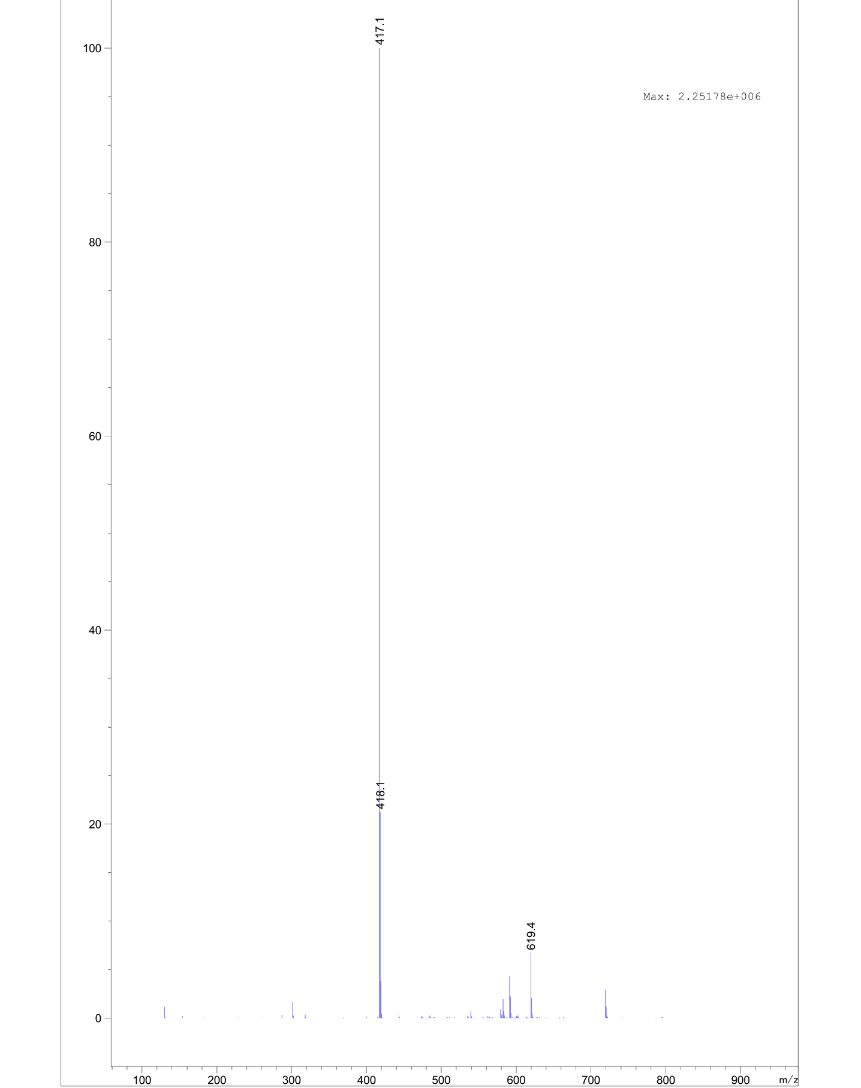

tert-butyl (1-((2-(2,6-dioxopiperidin-3-yl)-1,3-dioxoisoindolin-4-yl)oxy)-2-oxo-6,9,12,15,18-pentaoxa-3-azaicosan-20-yl)carbamate **S17**

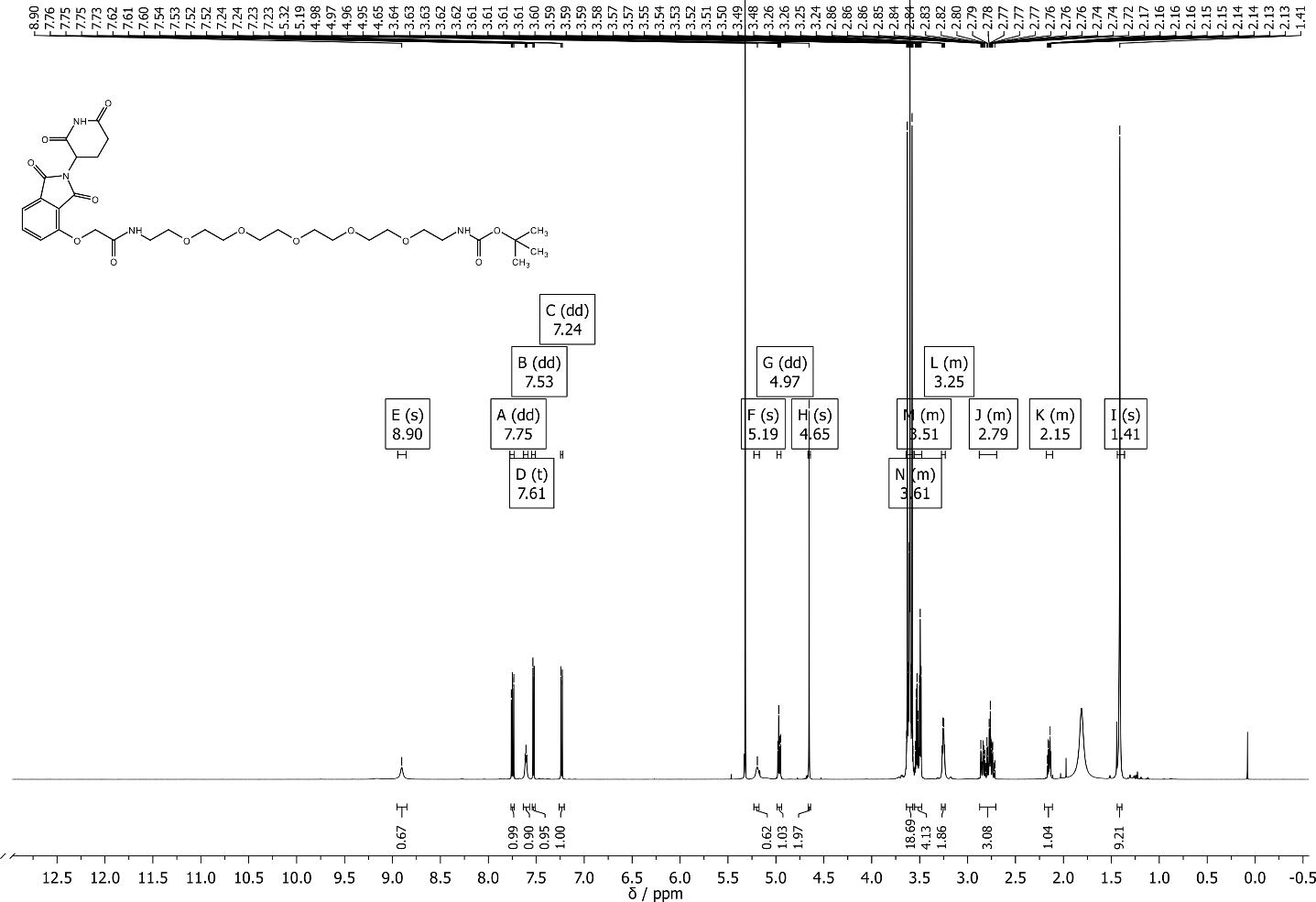

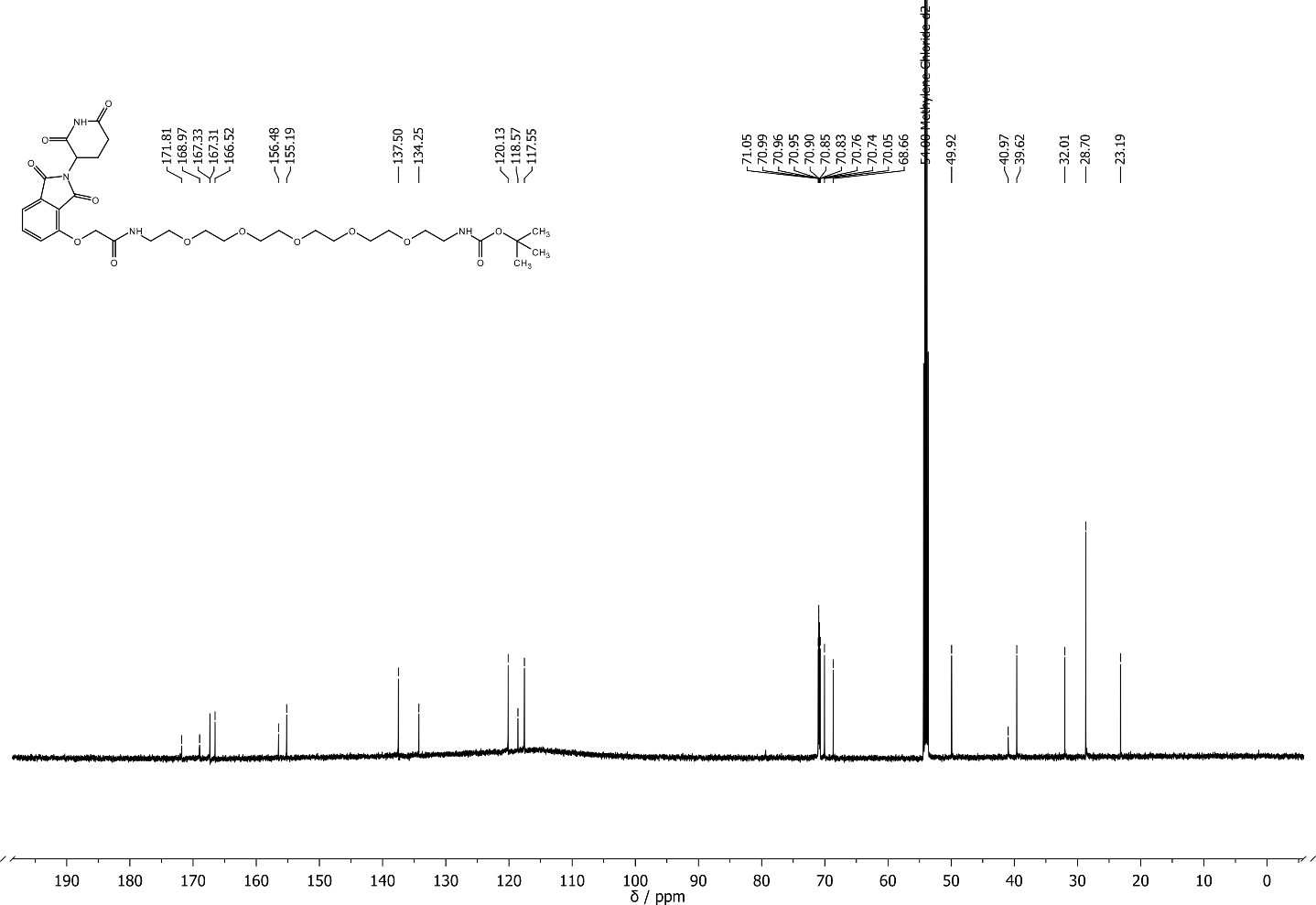

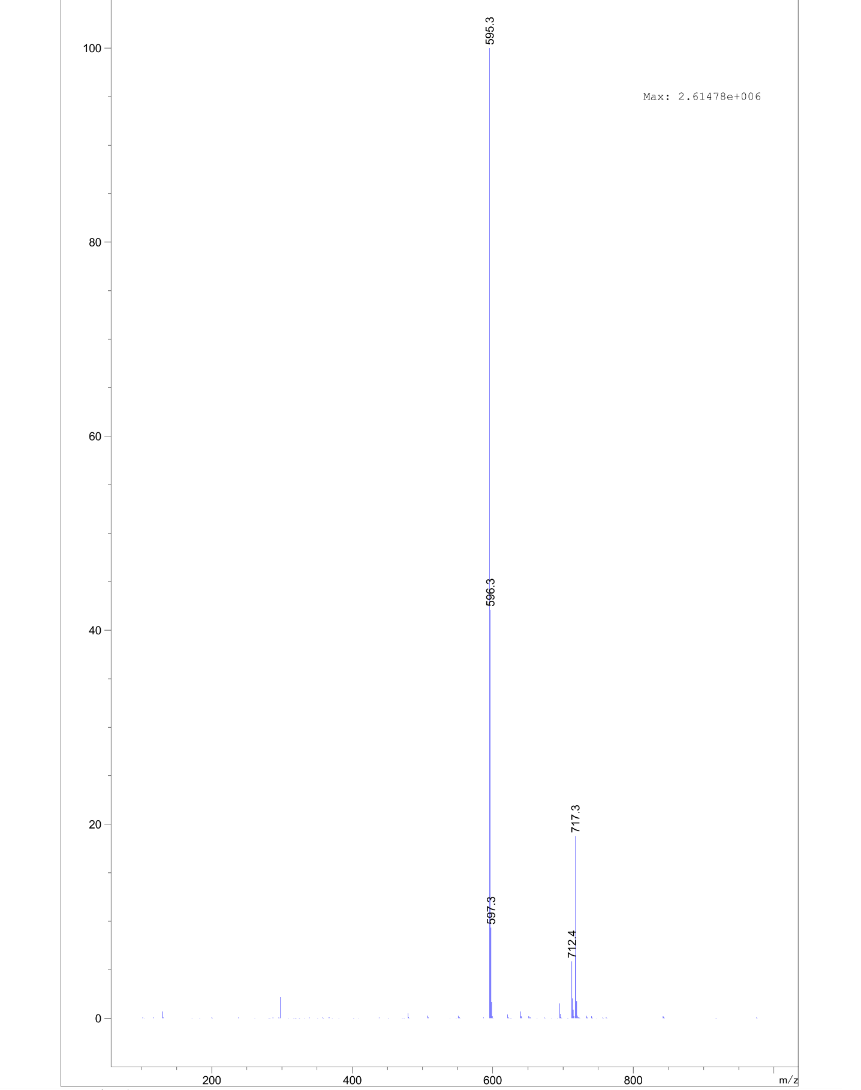

2-(4-(4-(benzylamino)phenoxy)phenoxy)-N-(5-(2-((2-(2,6-dioxopiperidin-3-yl)-1,3-dioxoisoindolin-4-yl)oxy)acetamido)pentyl)acetamide **S18**

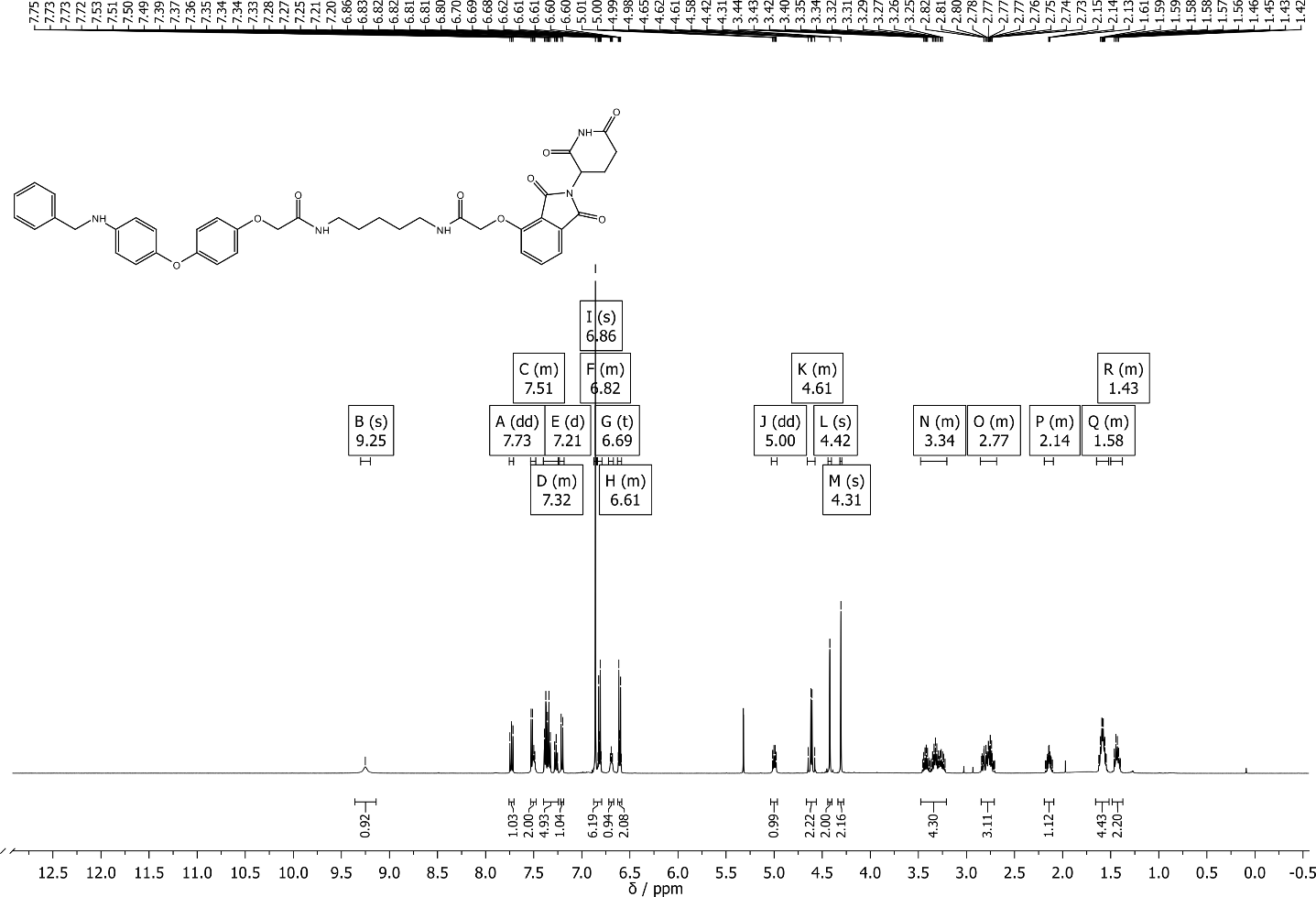

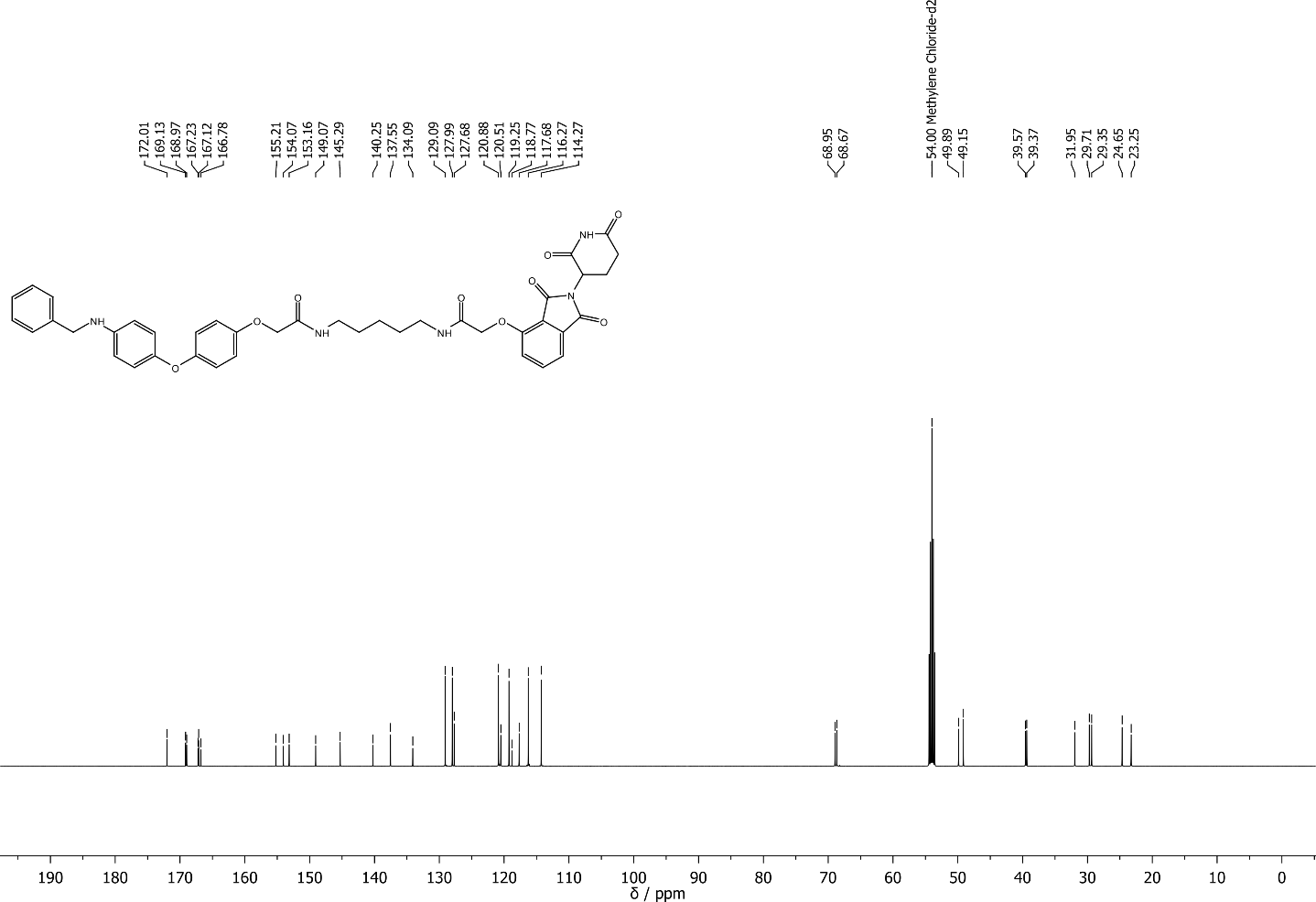

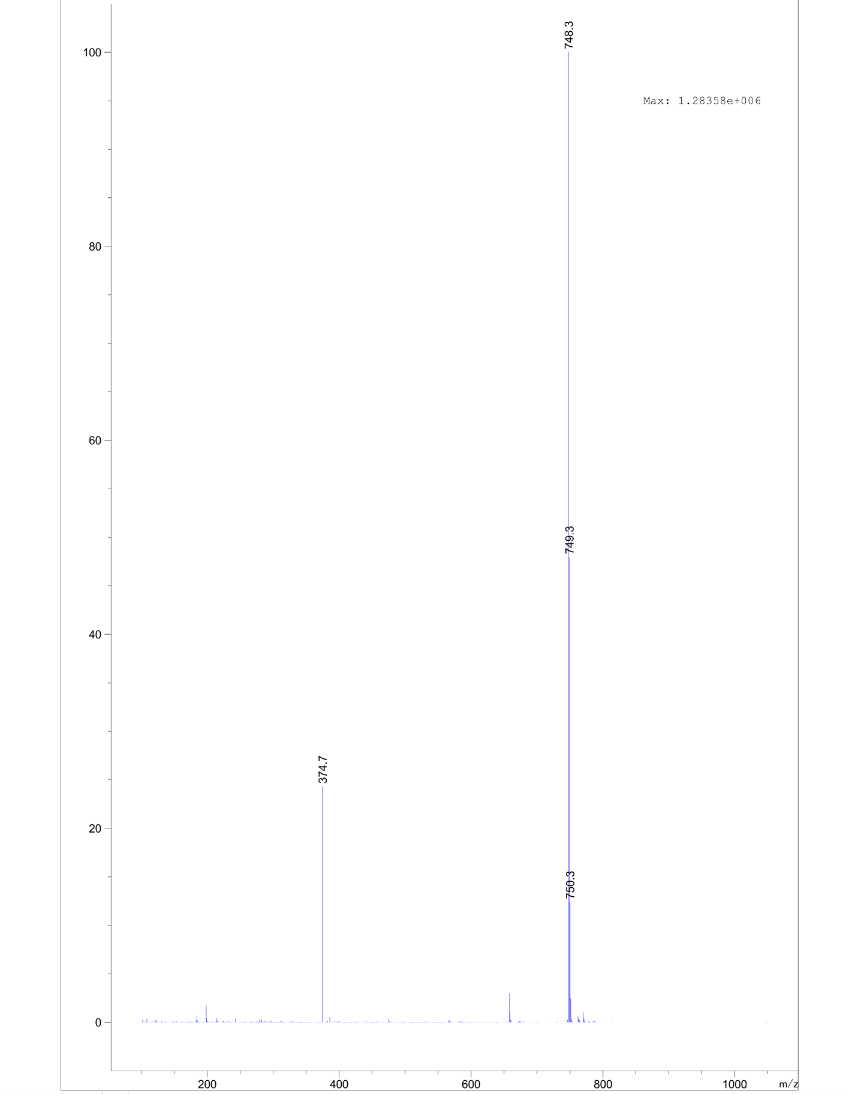

2-(4-(4-(benzylamino)phenoxy)phenoxy)-N-(1-((2-(2,6-dioxopiperidin-3-yl)-1,3-dioxoisoindolin-4-yl)oxy)-2-oxo-6,9,12,15,18-pentaoxa-3-azaicosan-20-yl)acetamide **S19**

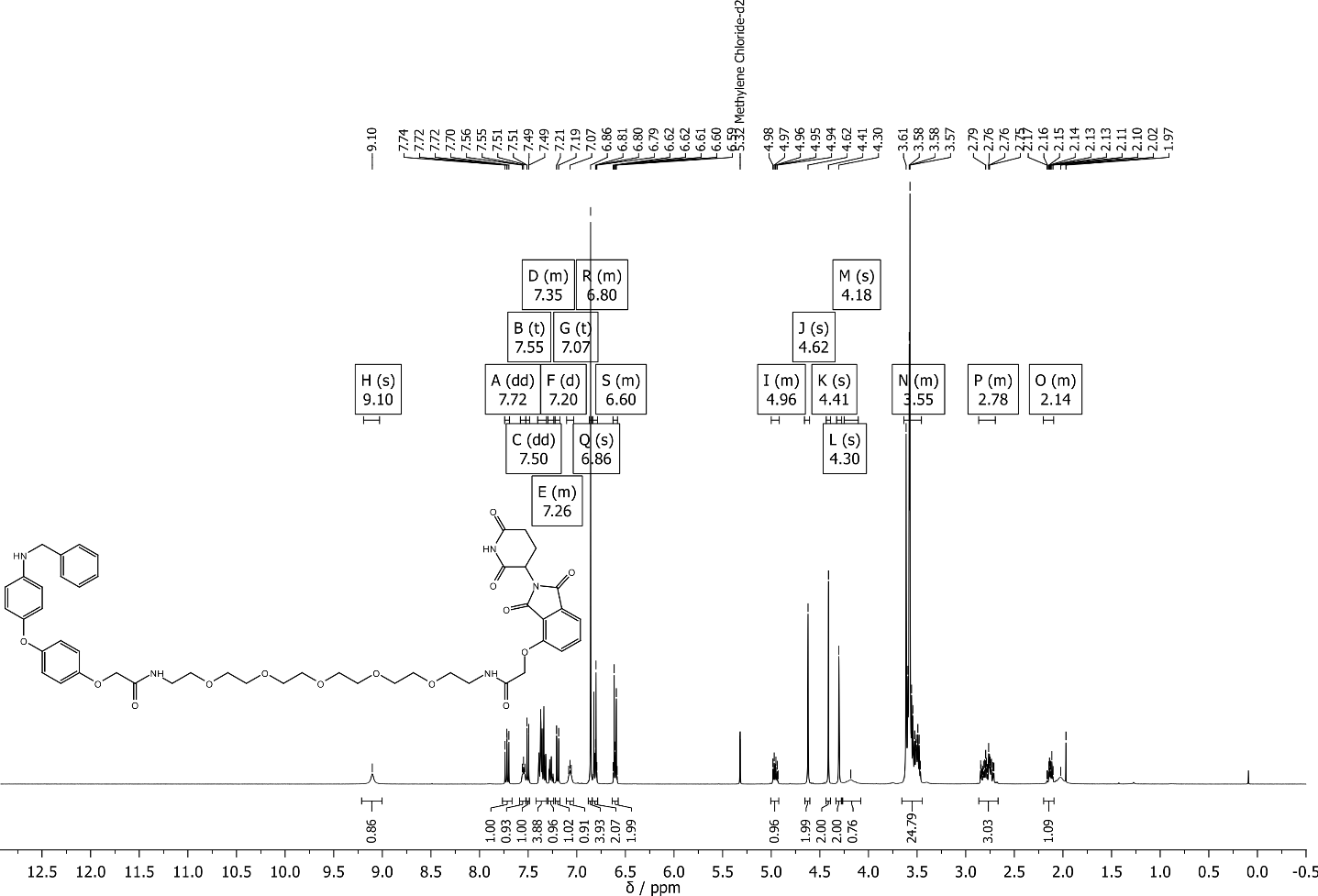

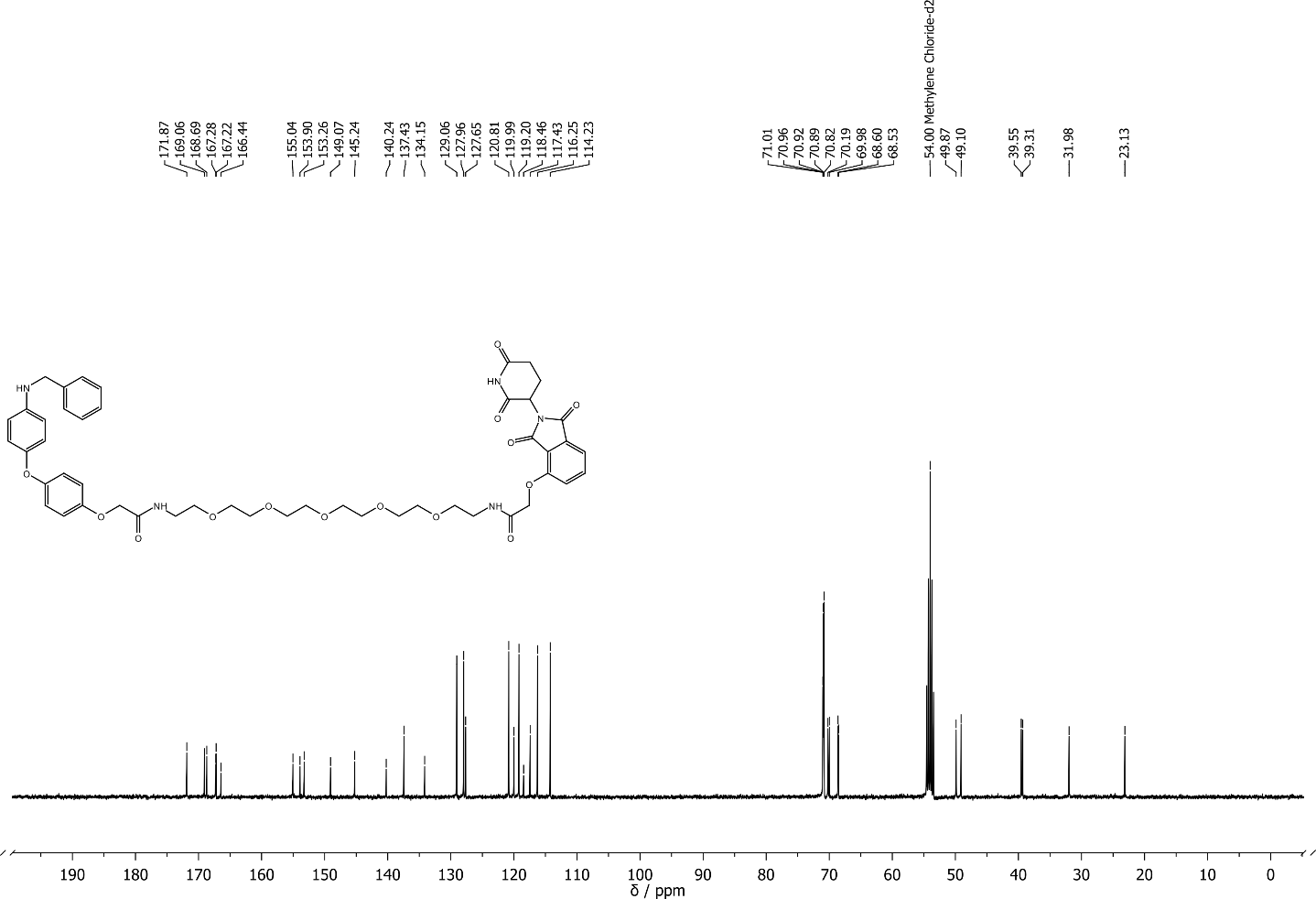

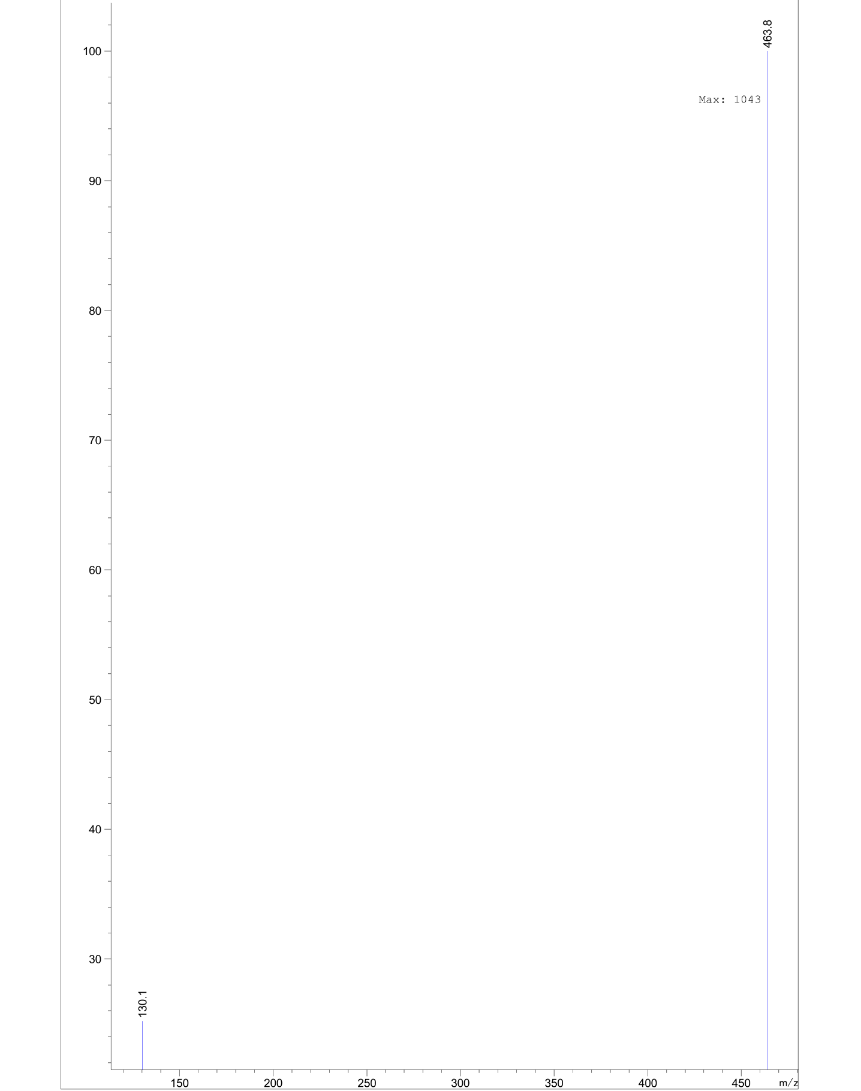

N-benzyl-2-chloro-N-(4-(4-(2-((5-(2-((2-(2,6-dioxopiperidin-3-yl)-1,3-dioxoisoindolin-4-yl)oxy)acetamido)pentyl)amino)-2-oxoethoxy)phenoxy)phenyl)acetamide **1c**

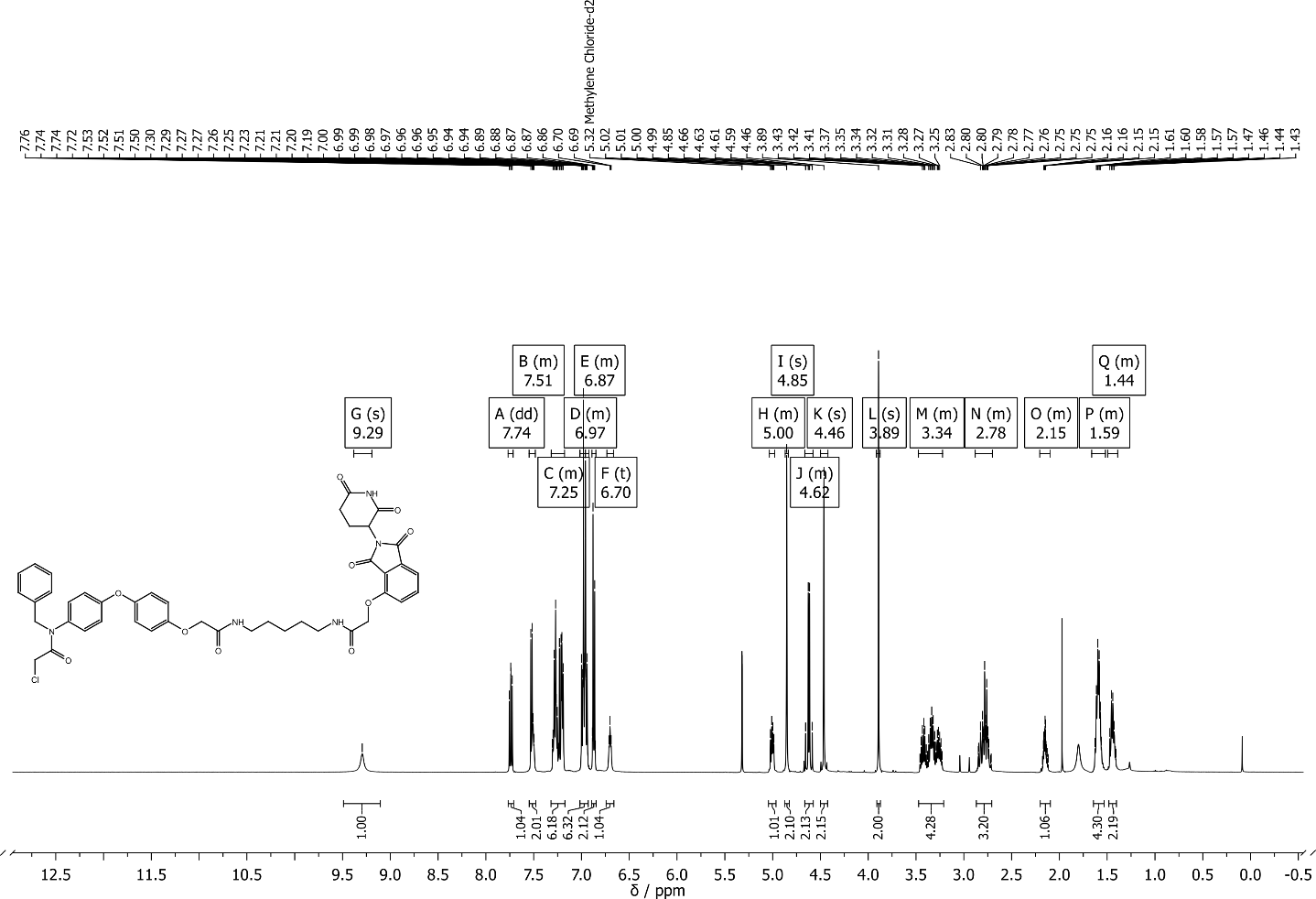

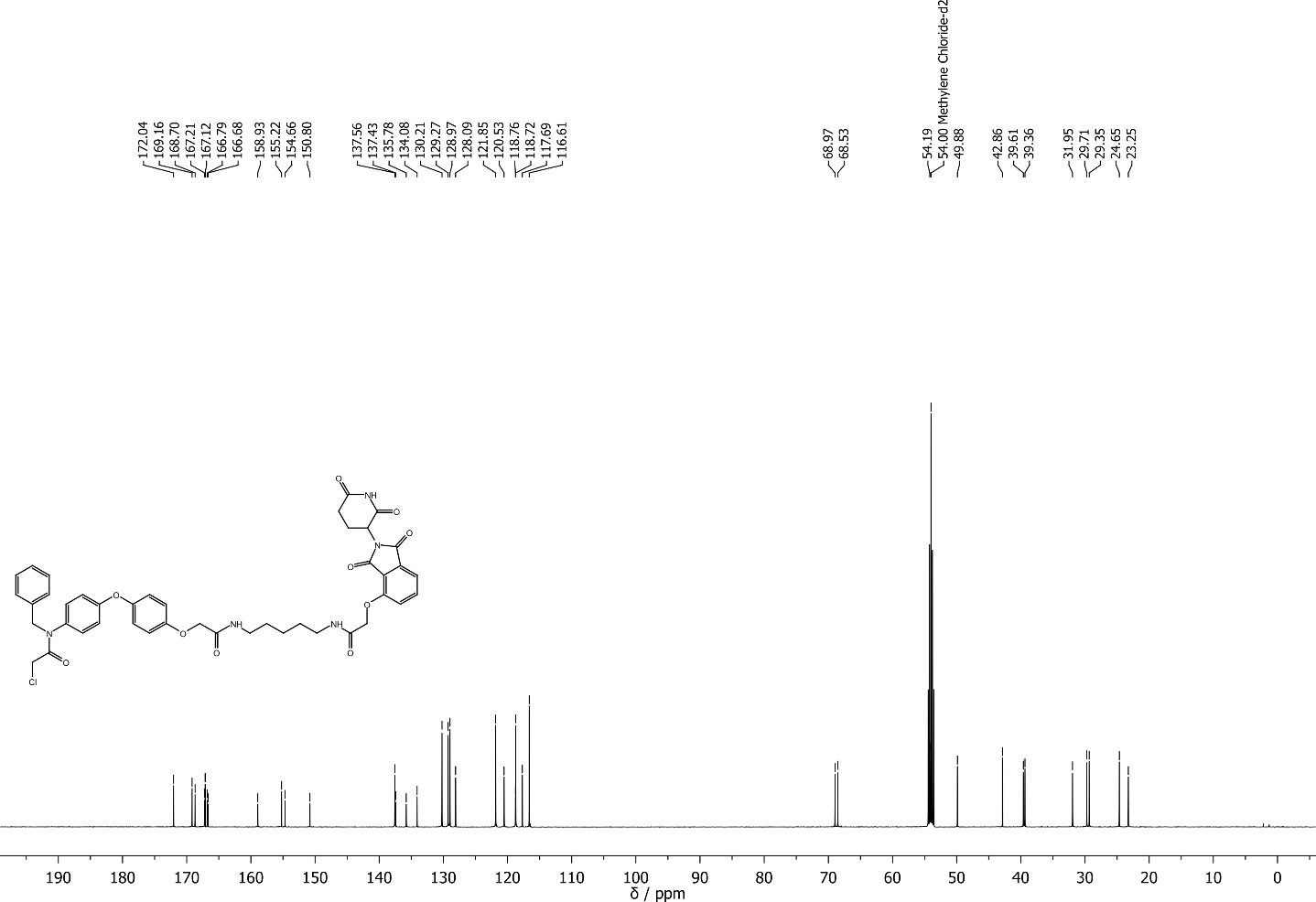

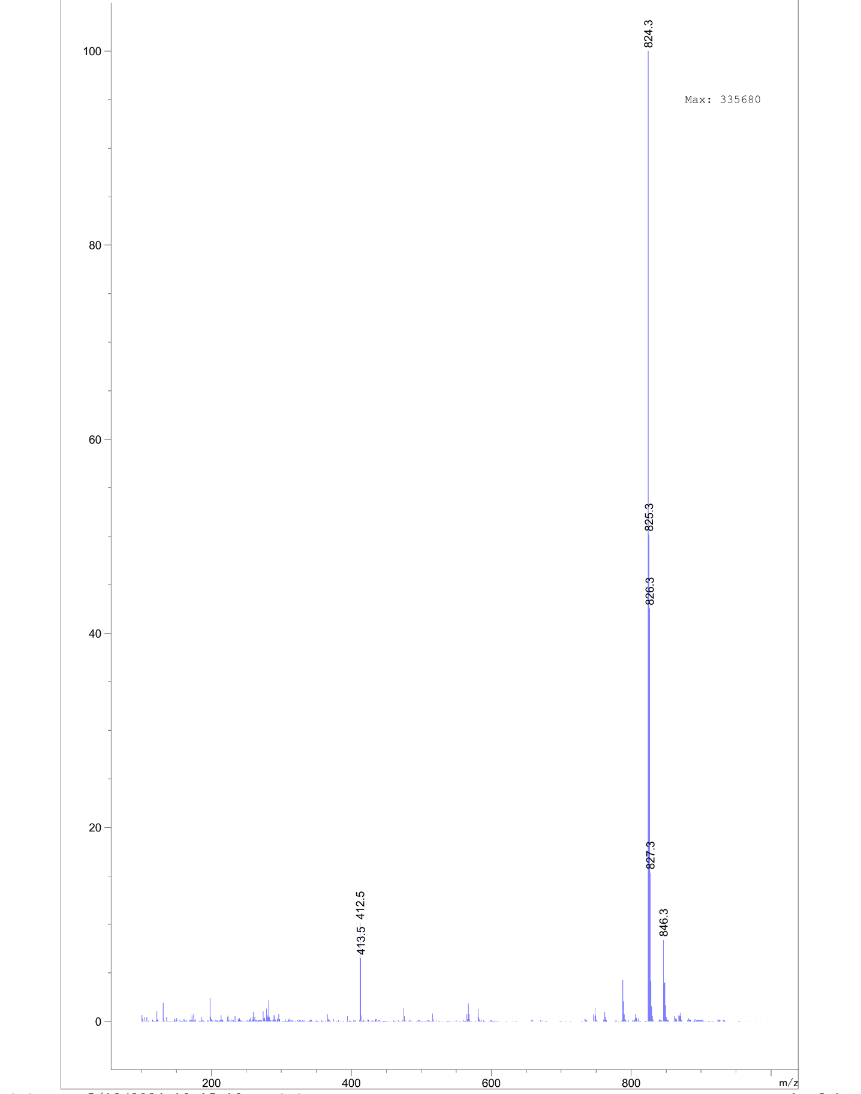

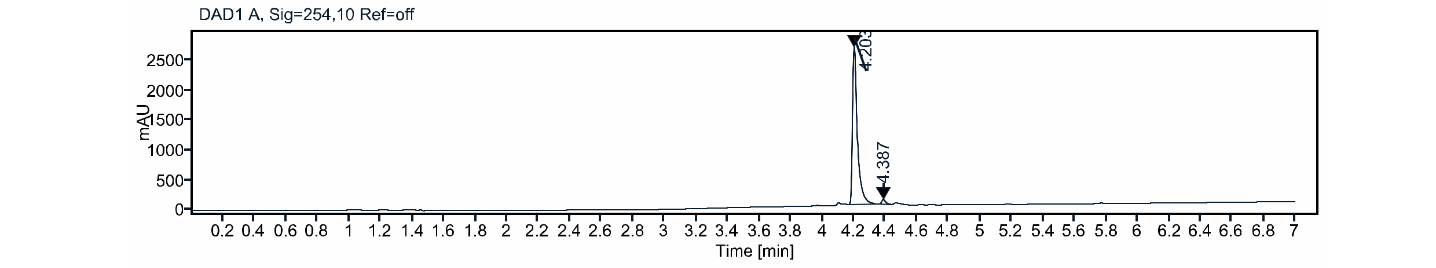

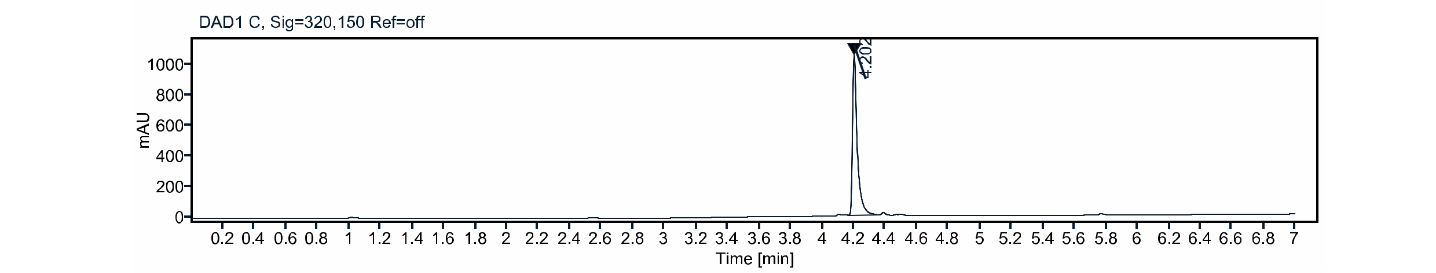

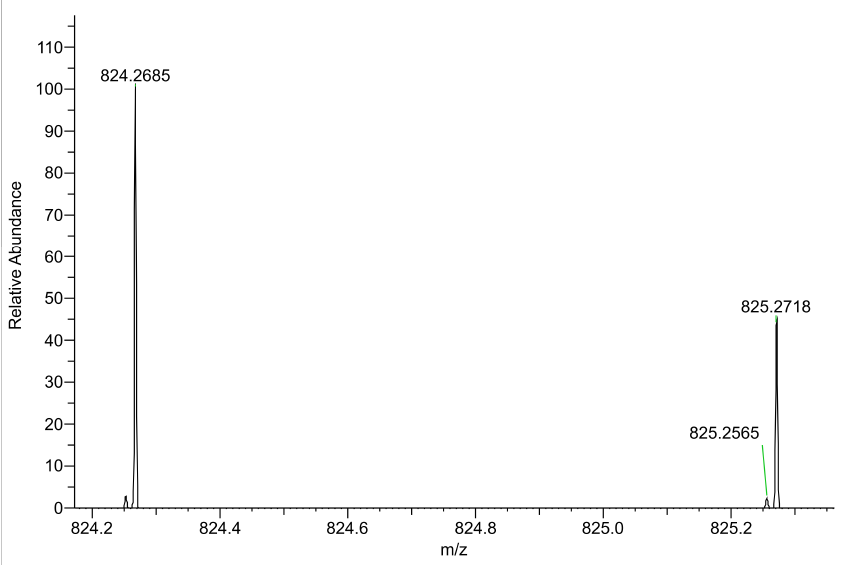

N-benzyl-2-chloro-N-(4-(4-((23-((2-(2,6-dioxopiperidin-3-yl)-1,3-dioxoisoindolin-4-yl)oxy)-2,22-dioxo-6,9,12,15,18-pentaoxa-3,21-diazatricosyl)oxy)phenoxy)phenyl)acetamide **1d**

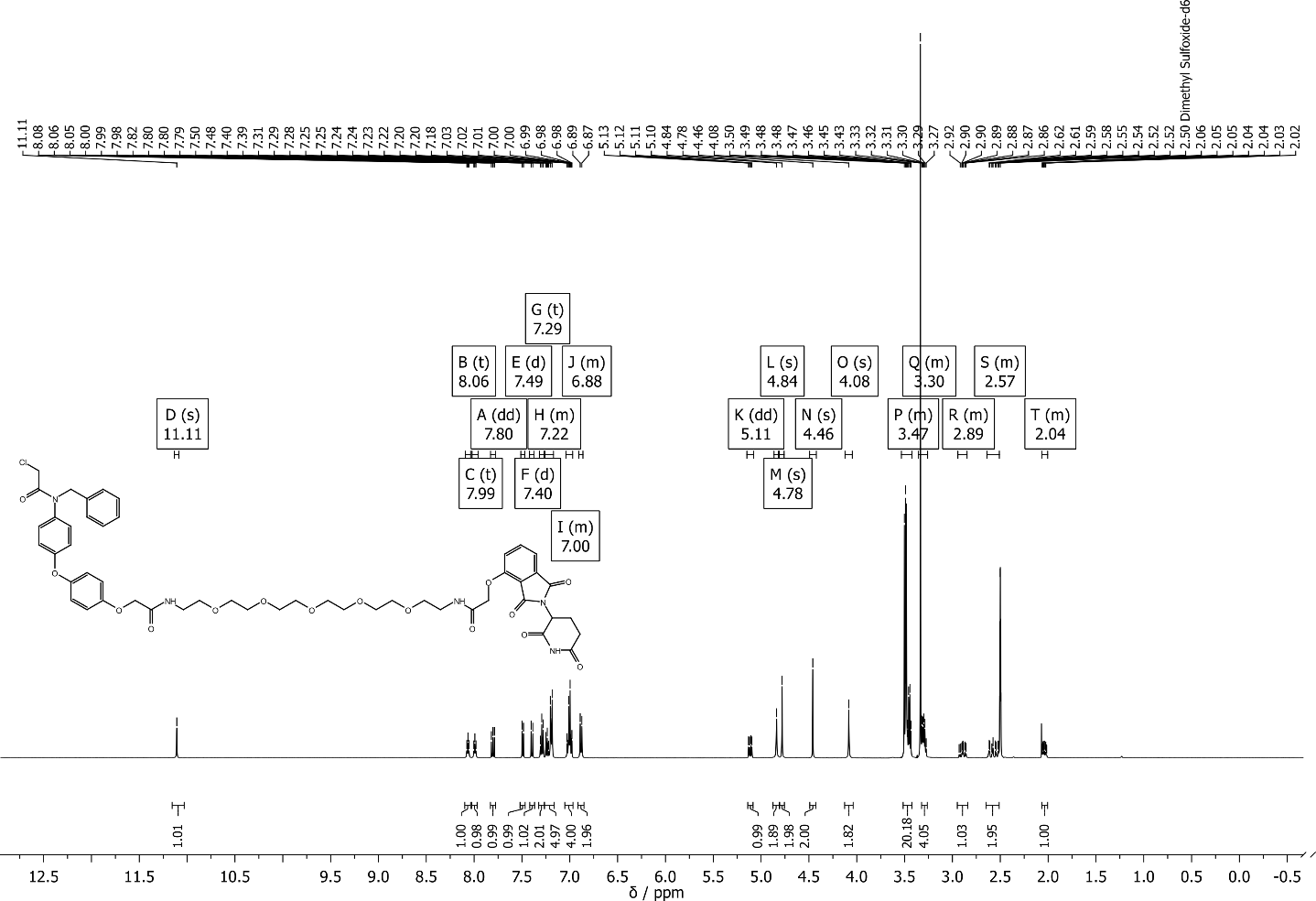

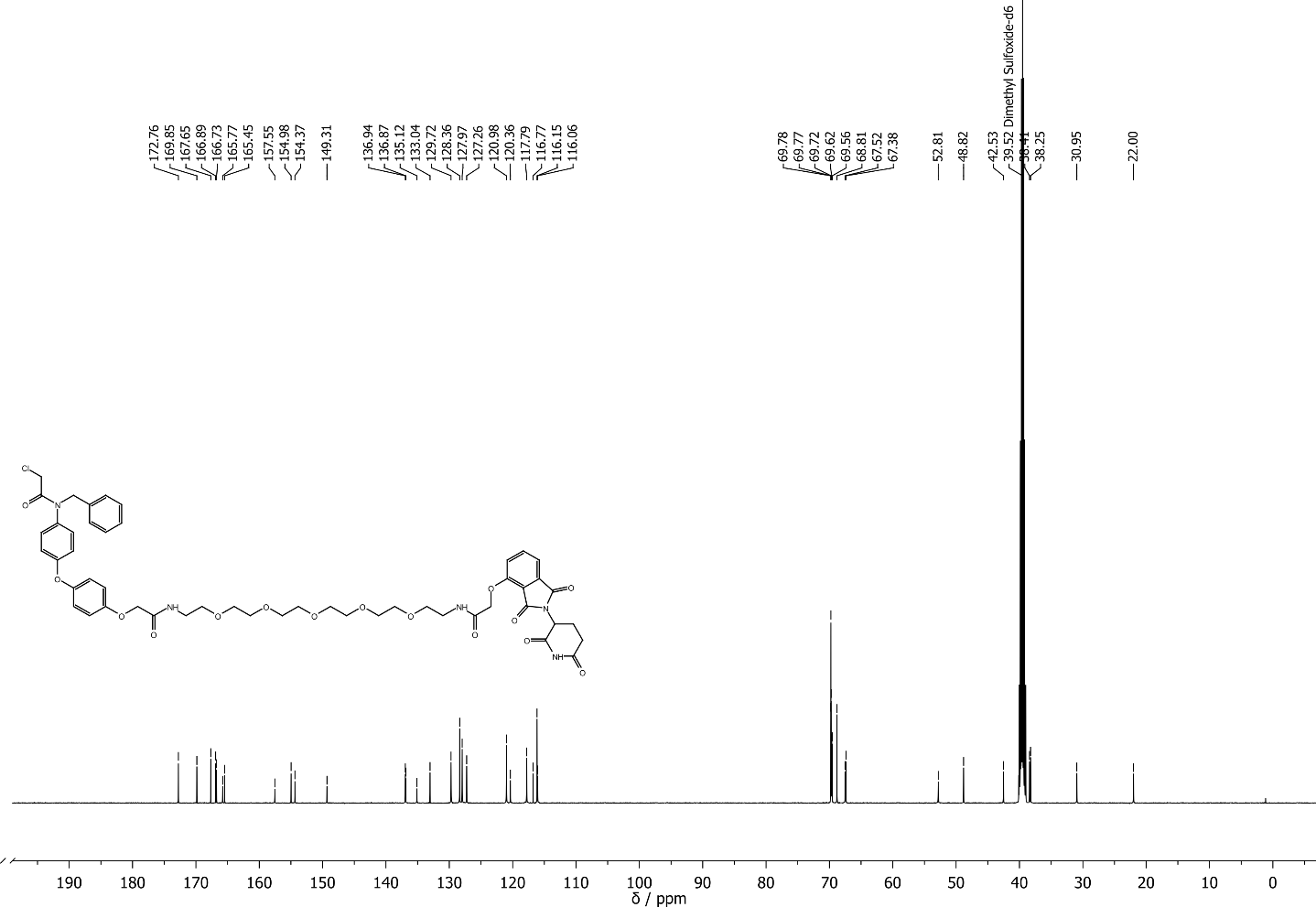

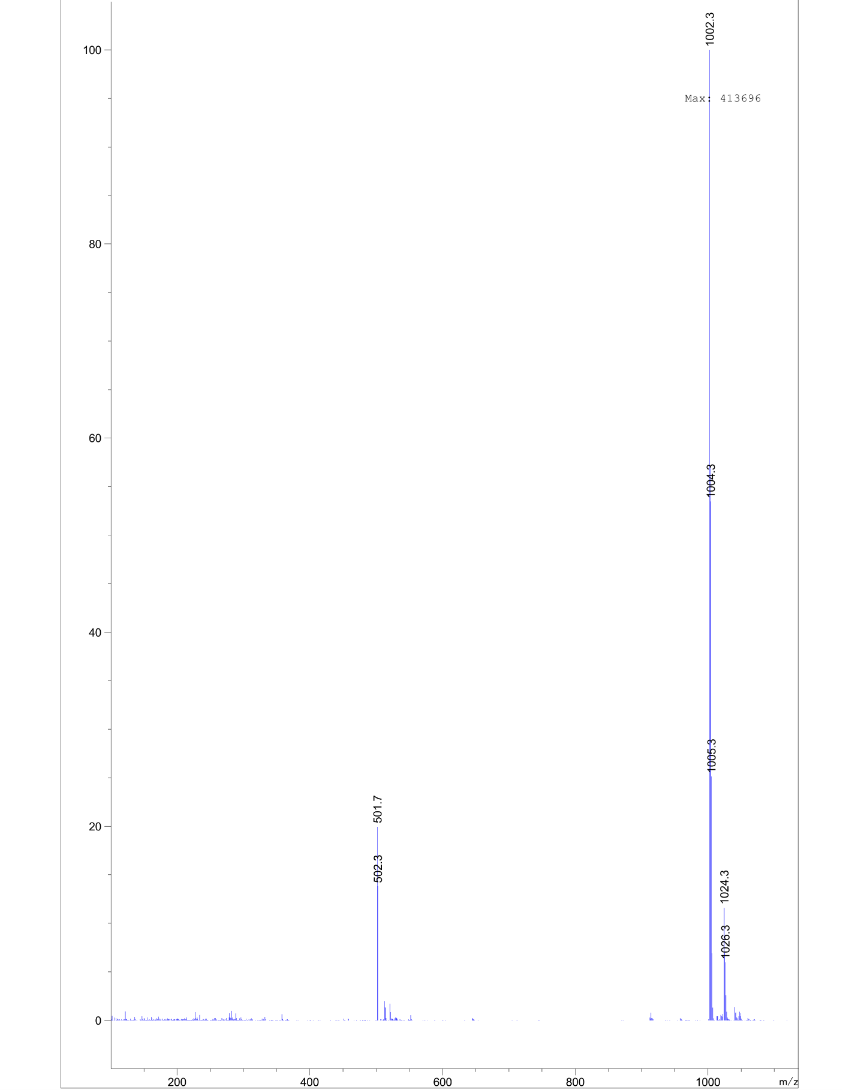

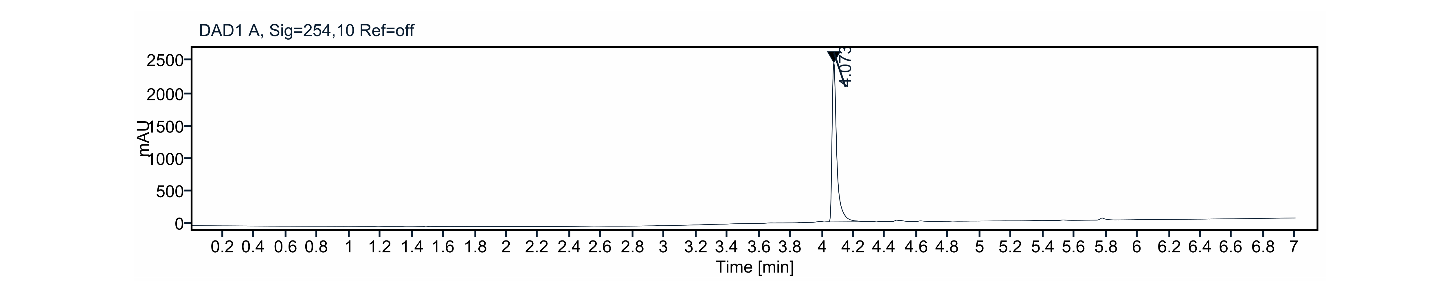

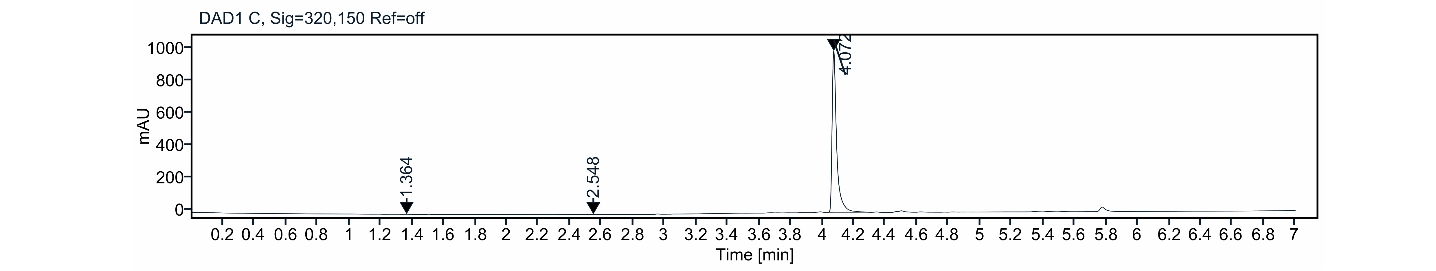

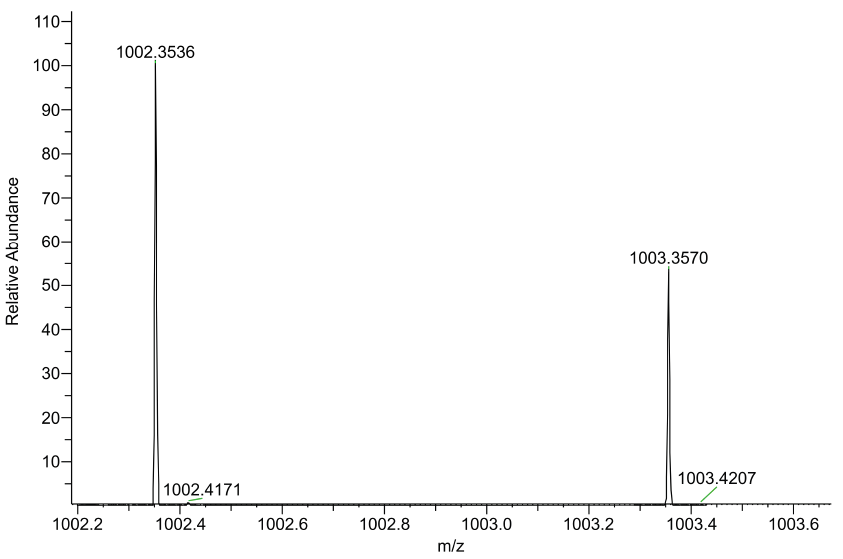

tert-butyl 7-(2-(4-(4-(benzylamino)phenoxy)phenoxy)acetamido)heptanoate **S20**

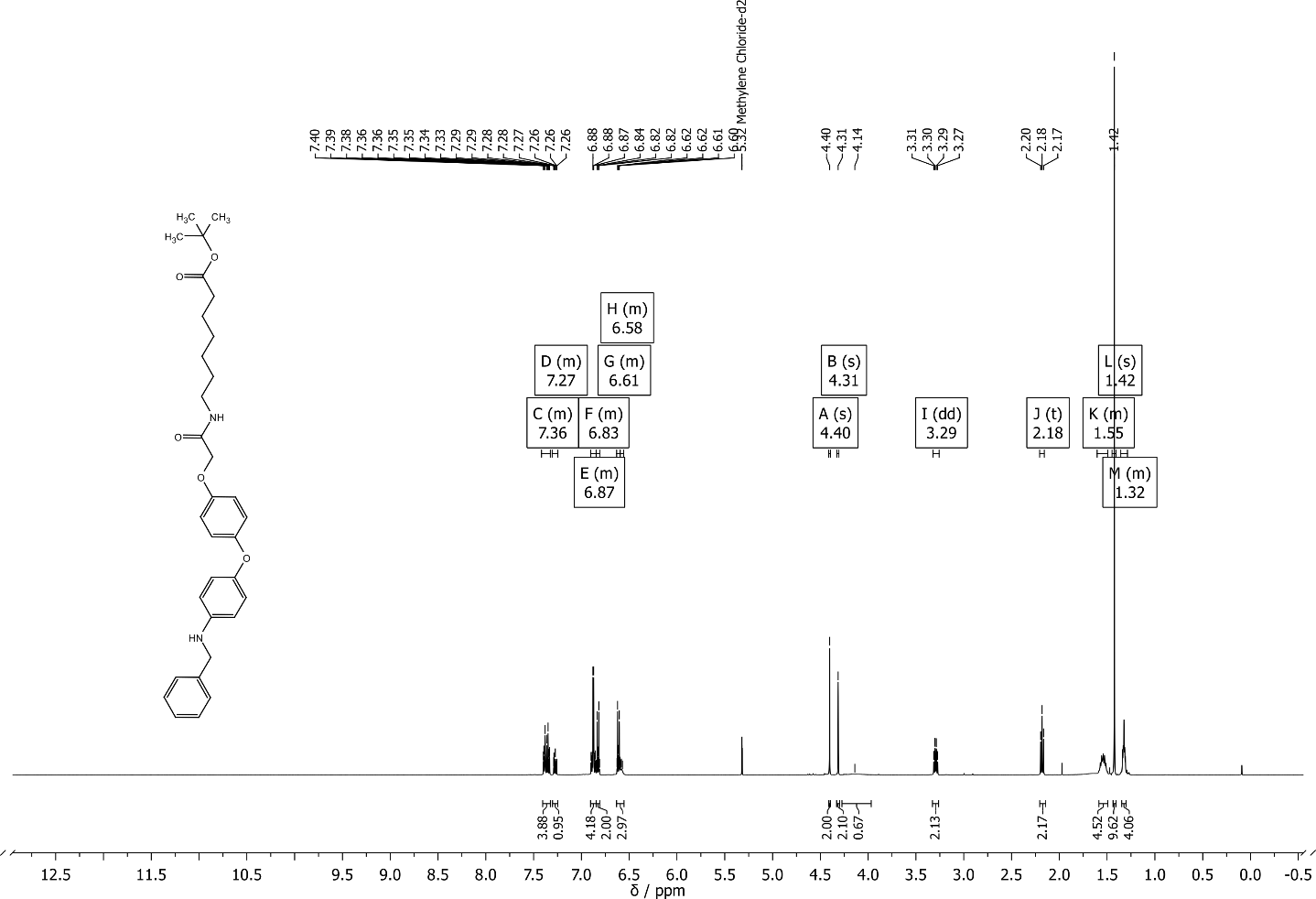

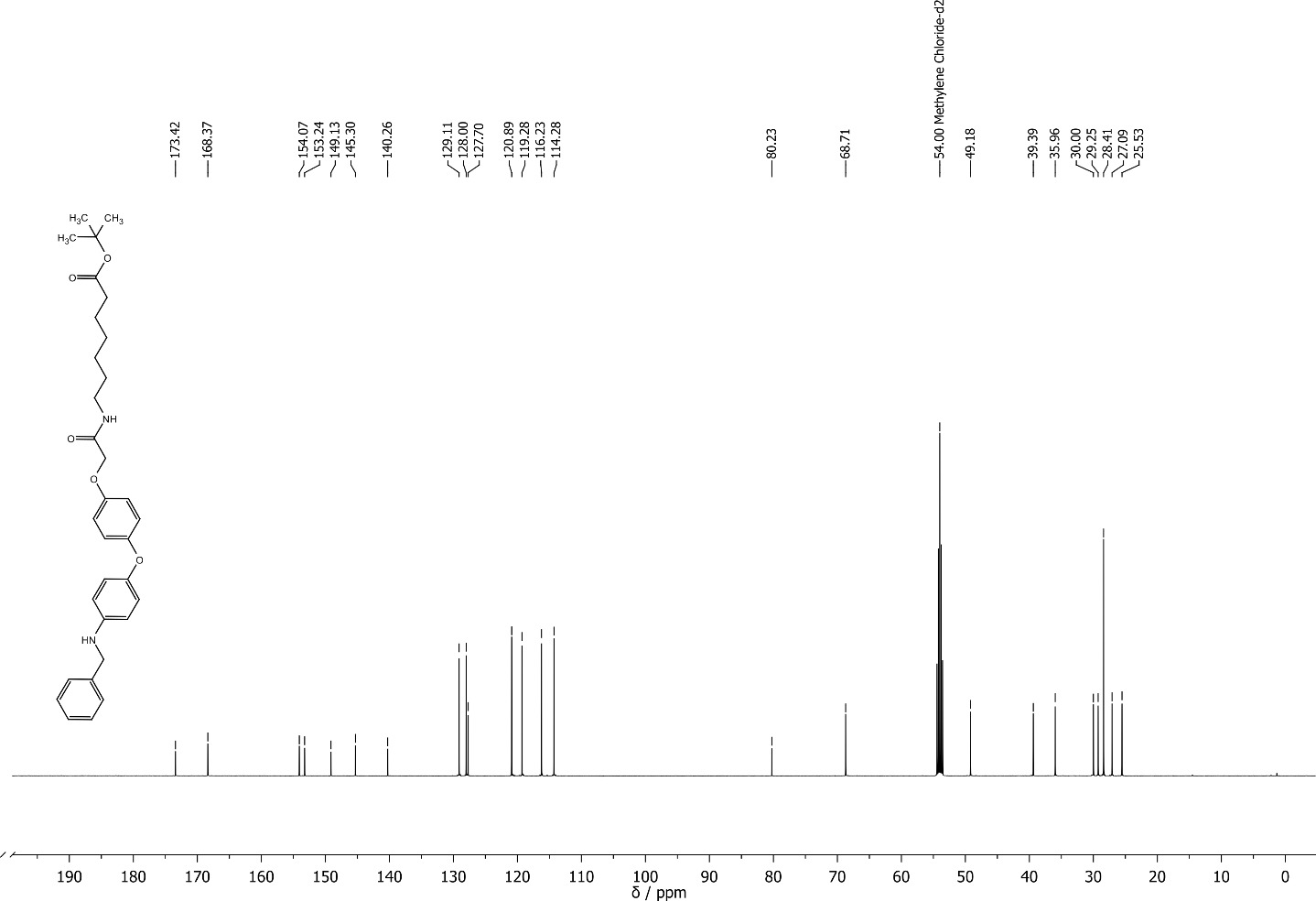

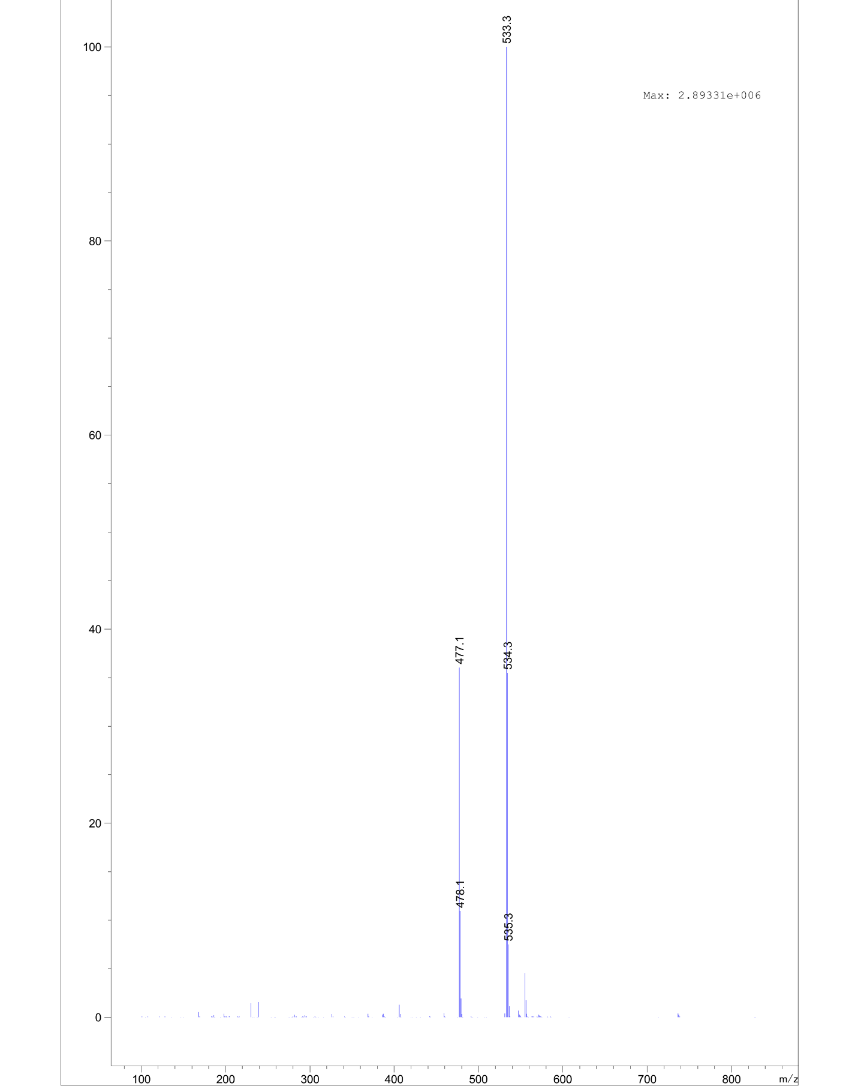

(2S,4R)-1-((S)-2-(7-(2-(4-(4-(benzylamino)phenoxy)phenoxy)acetamido)heptanamido)-3,3-dimethylbutanoyl)-4-hydroxy-N-(4-(4-methylthiazol-5-yl)benzyl)pyrrolidine-2-carboxamide **S21**

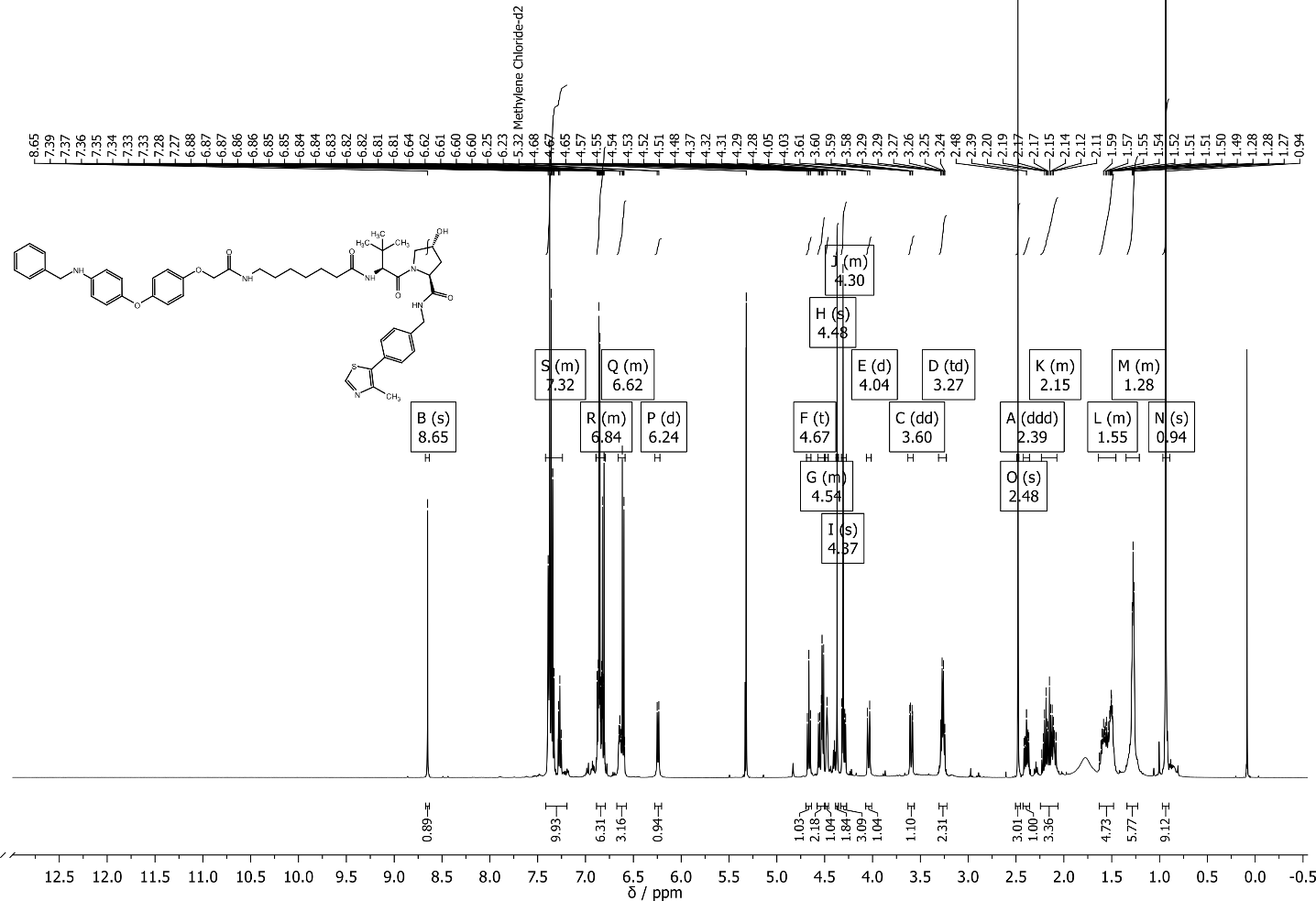

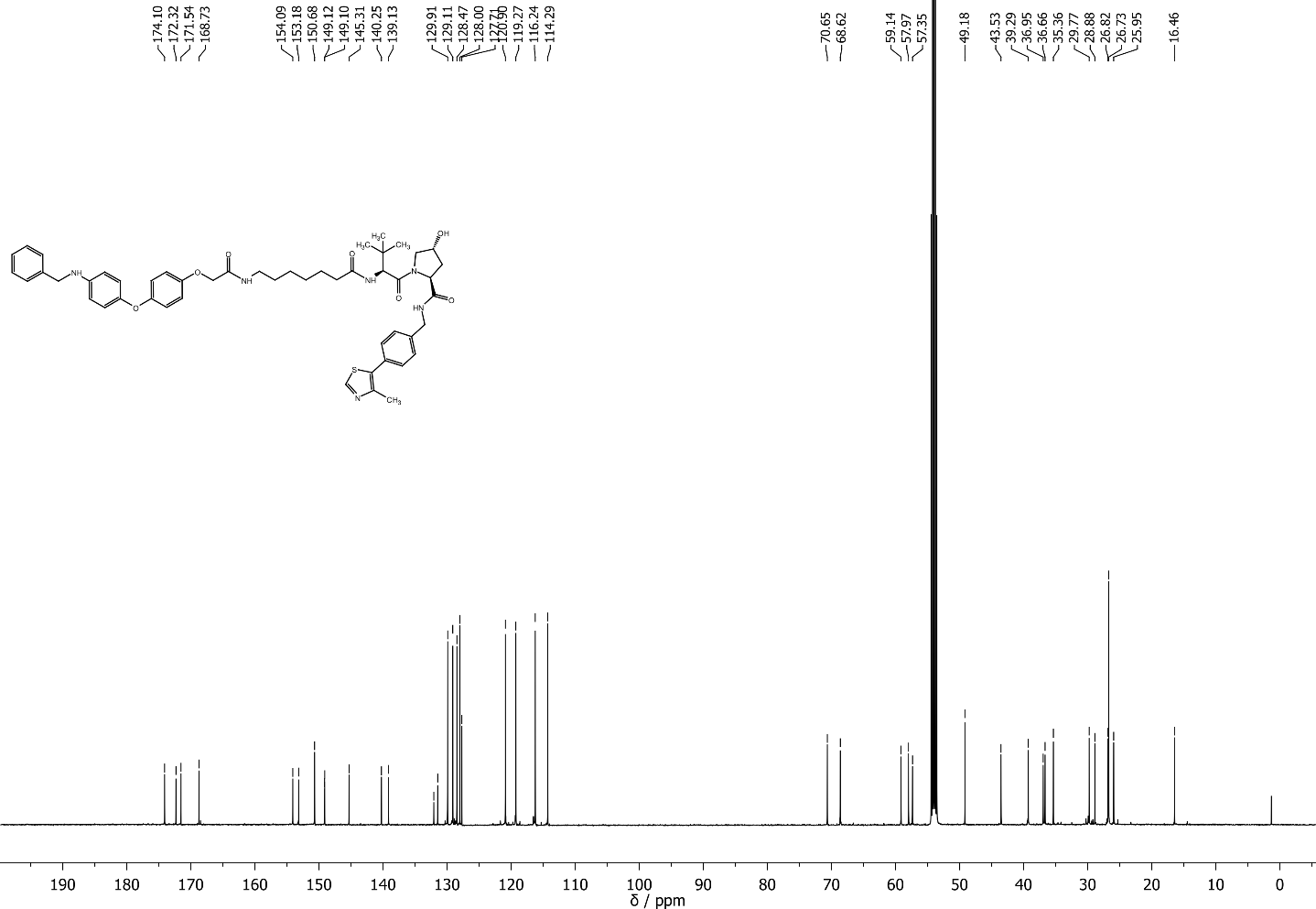

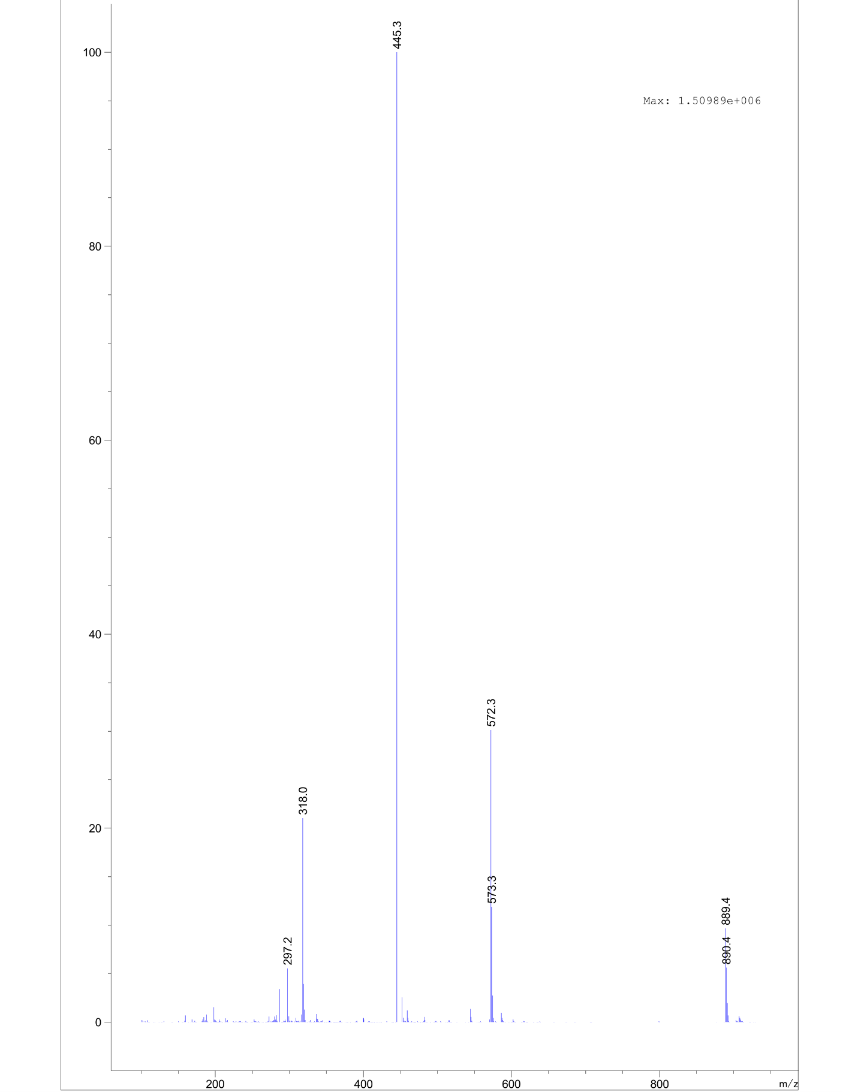

(2S,4R)-1-((S)-2-(7-(2-(4-(4-(N-benzyl-2-chloroacetamido)phenoxy)phenoxy)acetamido)heptanamido)-3,3-dimethylbutanoyl)-4-hydroxy-N-(4-(4-methylthiazol-5-yl)benzyl)pyrrolidine-2-carboxamide **2a**

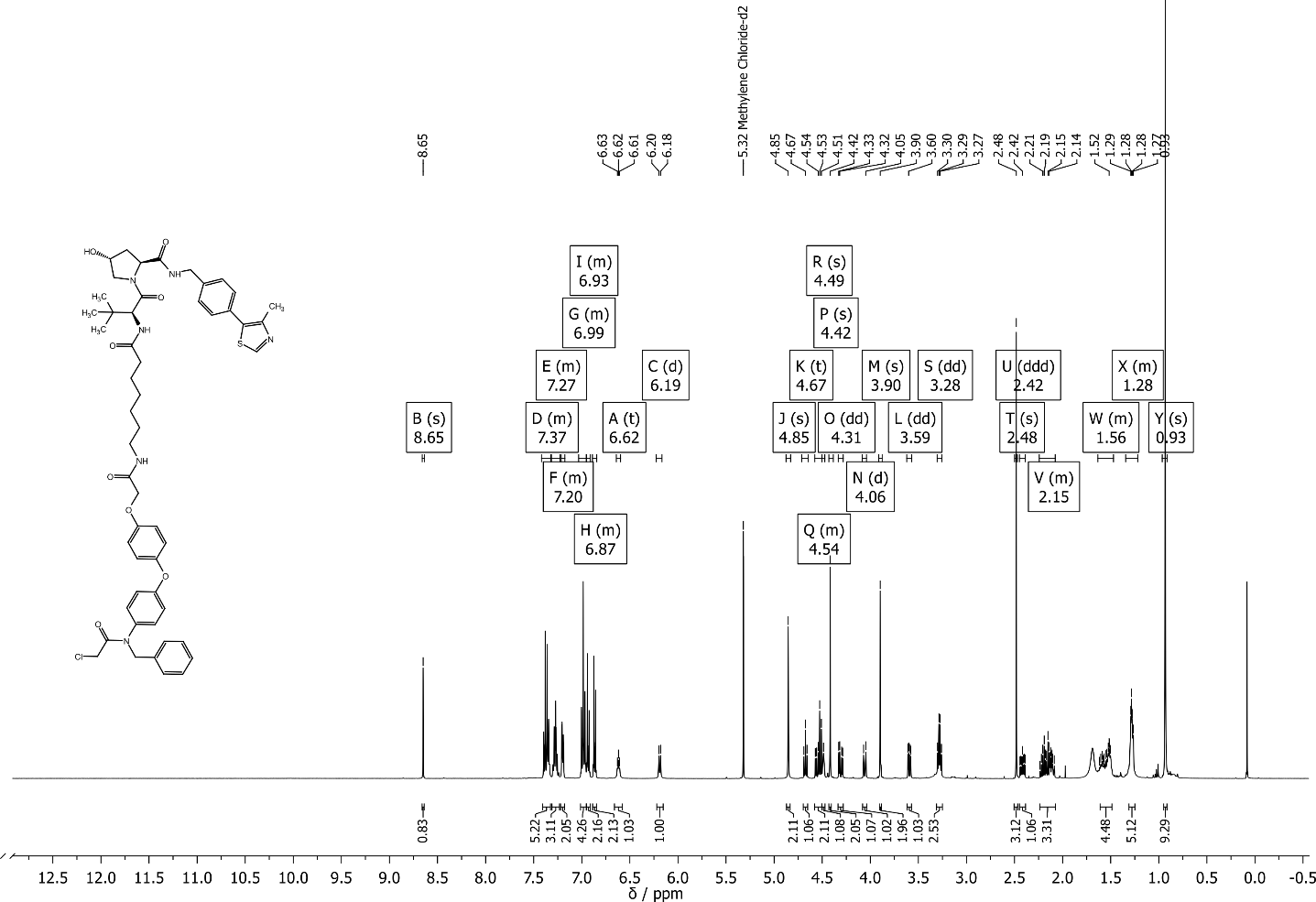

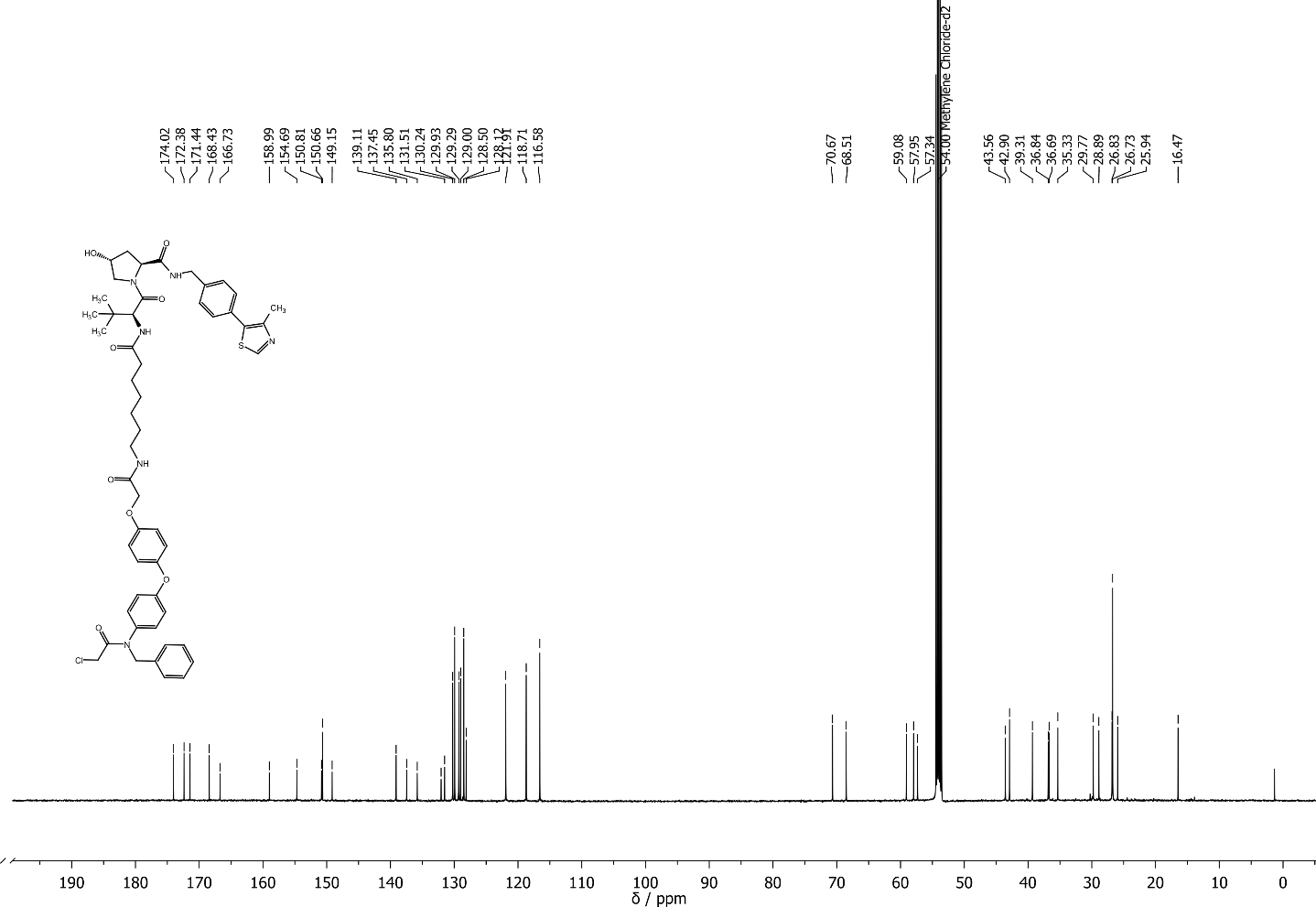

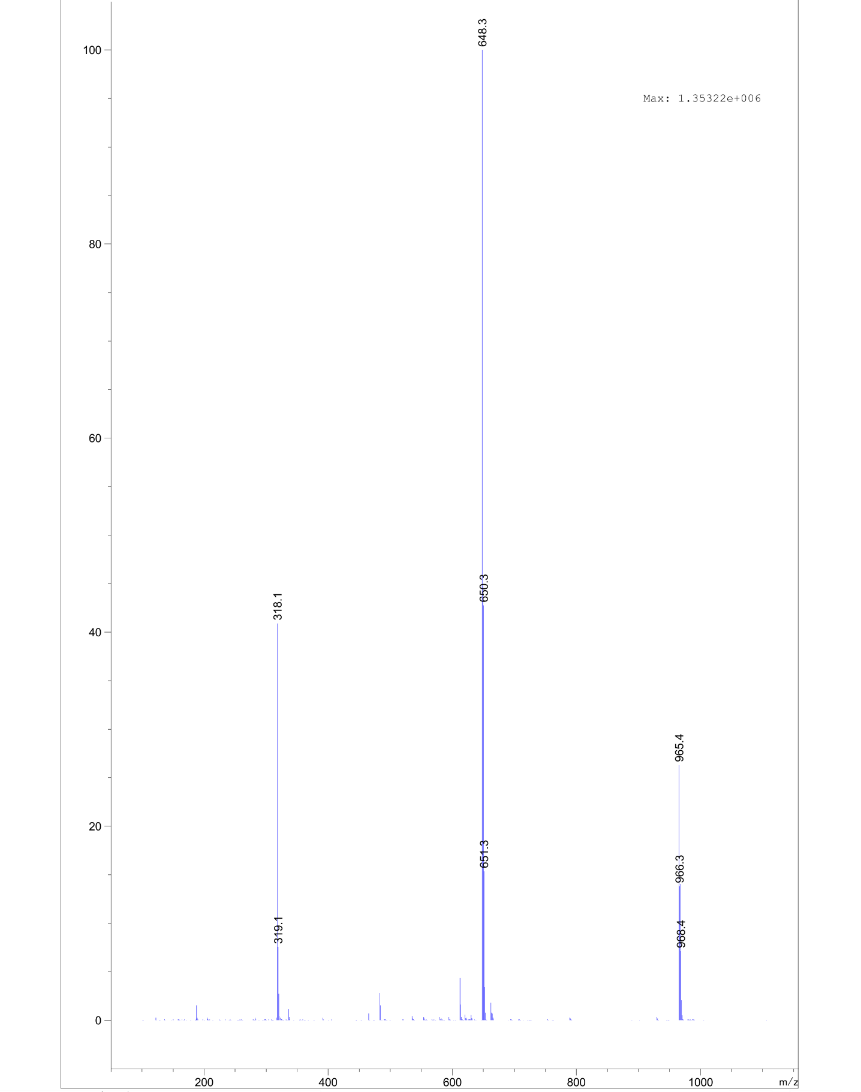

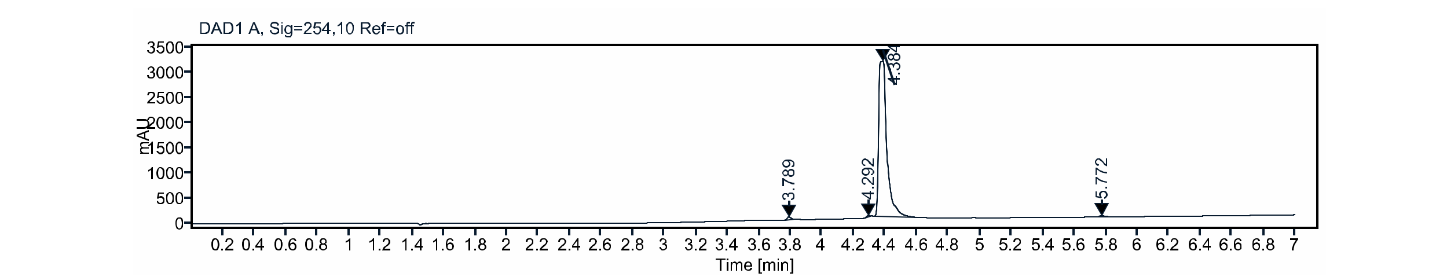

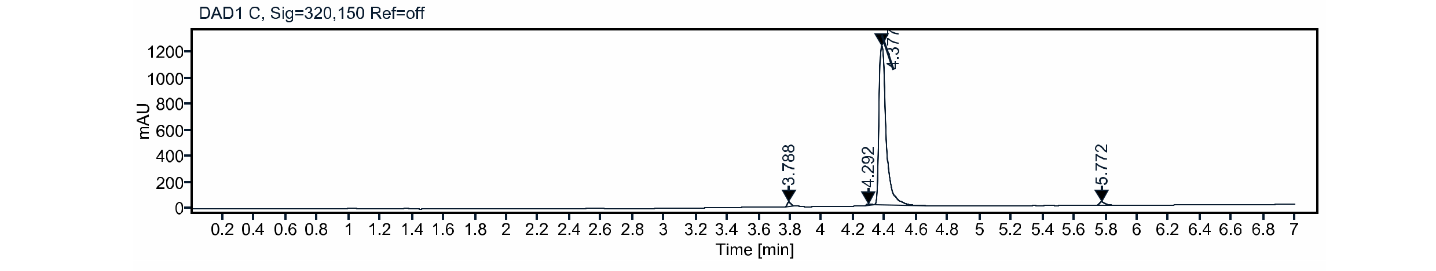

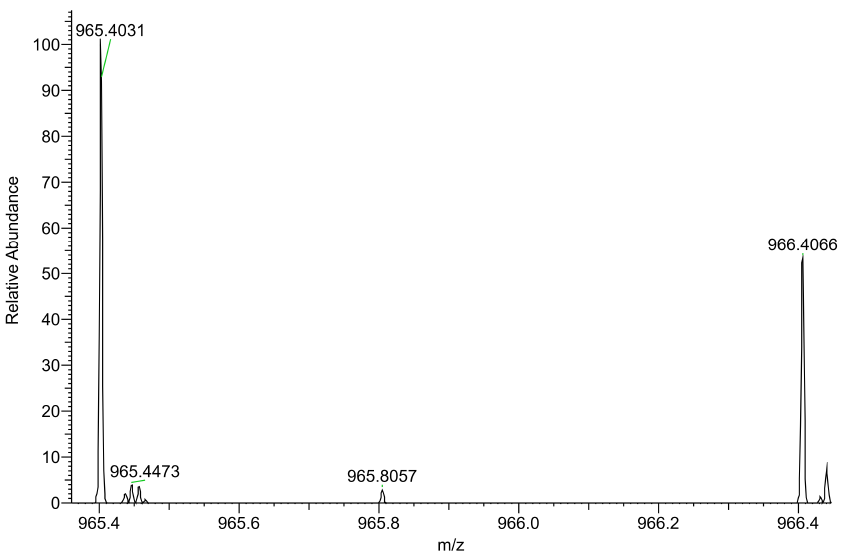

tert-butyl 3-(2-(2-(2-(4-(4-(benzylamino)phenoxy)phenoxy)ethoxy)ethoxy)ethoxy)propanoate **S26**

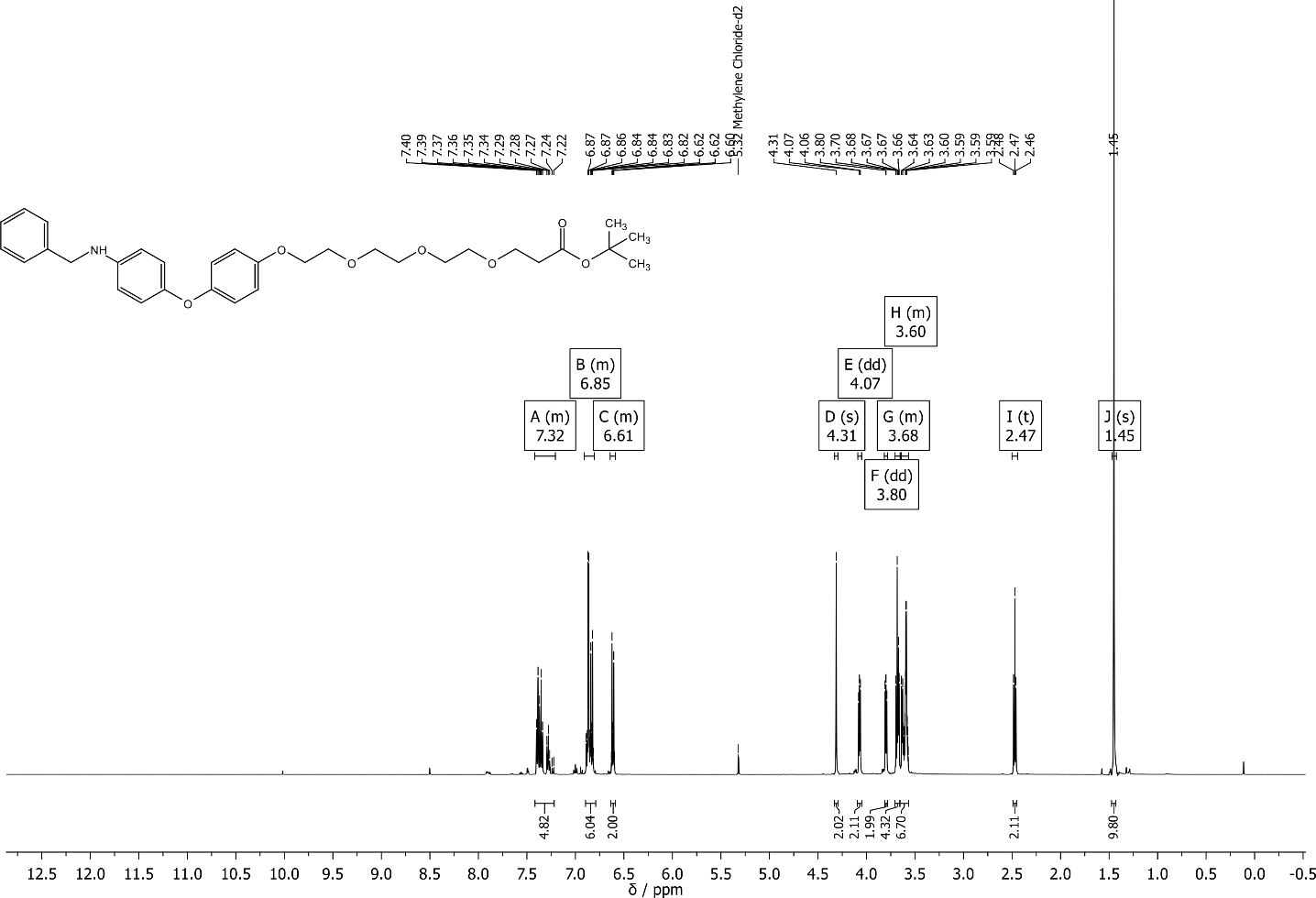

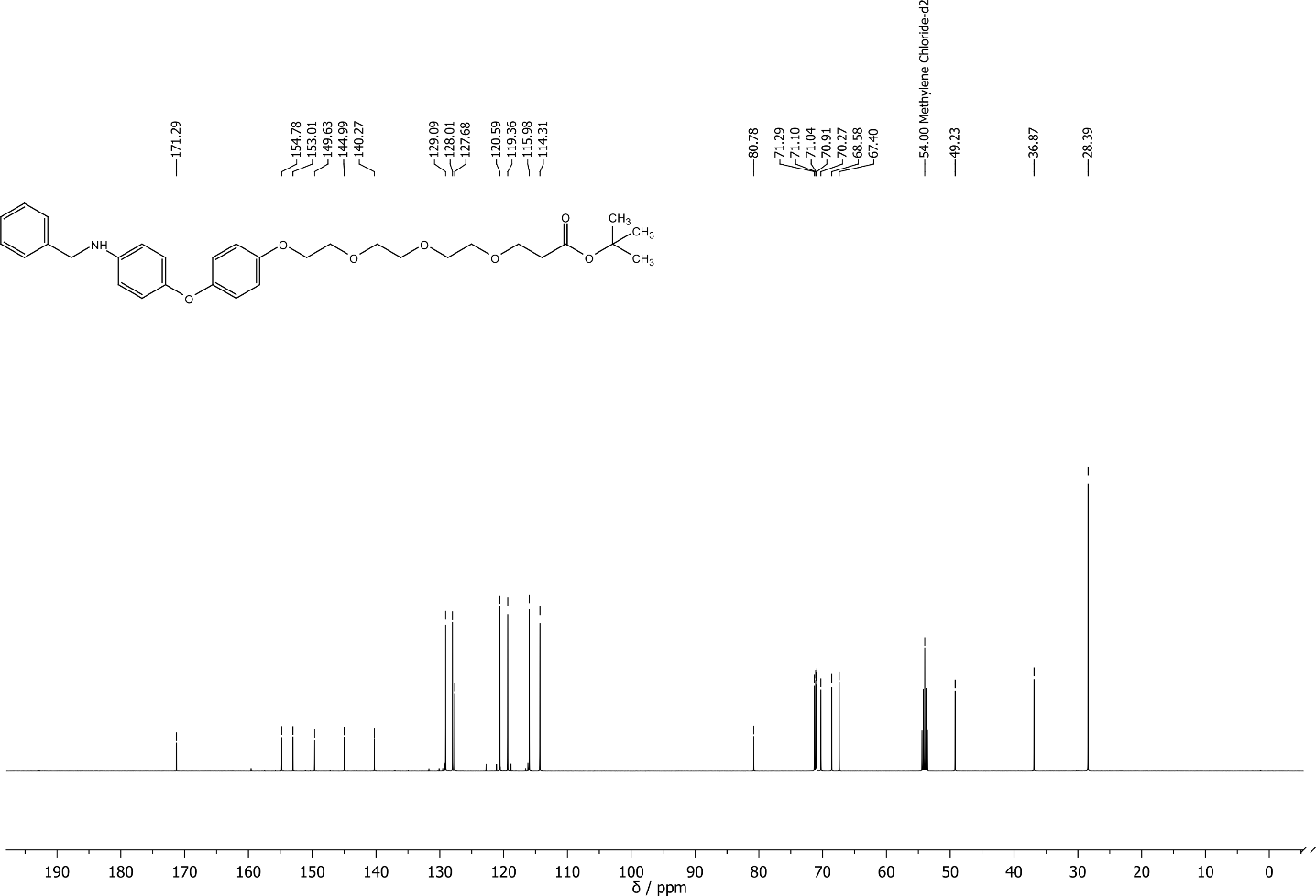

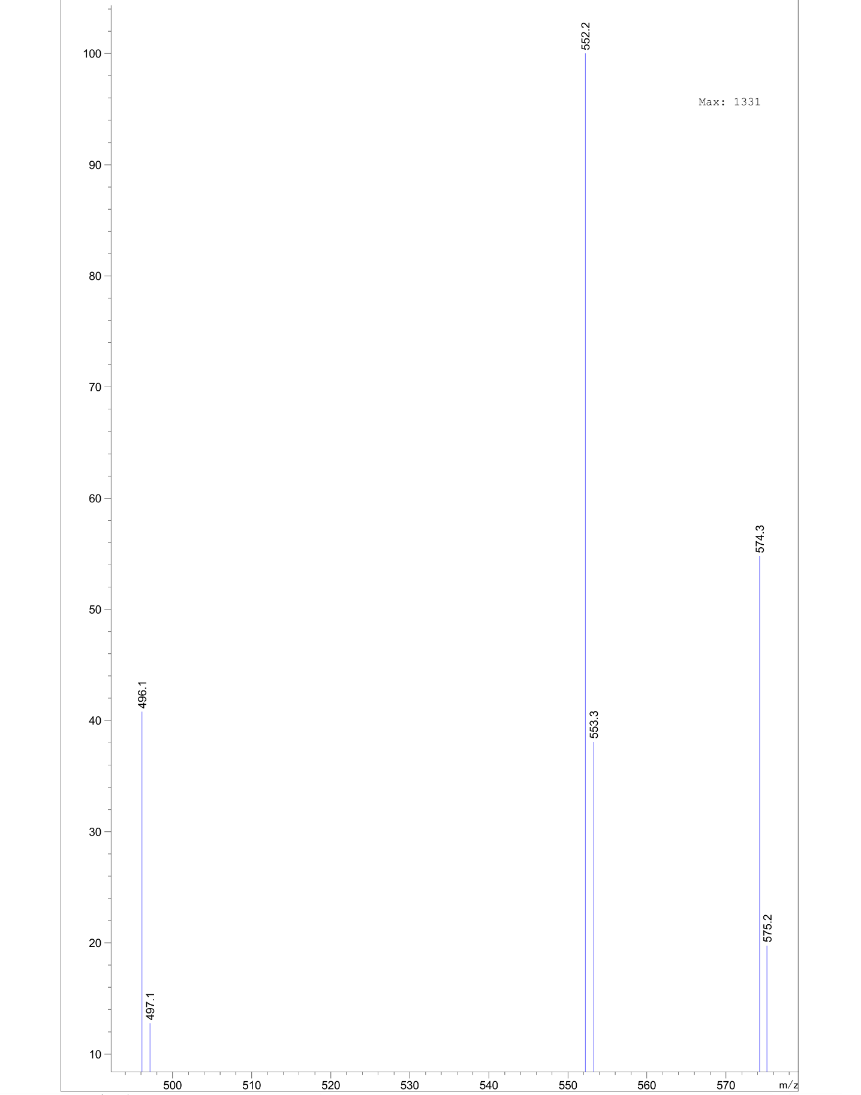

tert-butyl 1-(4-(4-(benzylamino)phenoxy)phenoxy)-3,6,9,12,15-pentaoxaoctadecan-18-oate **S27**

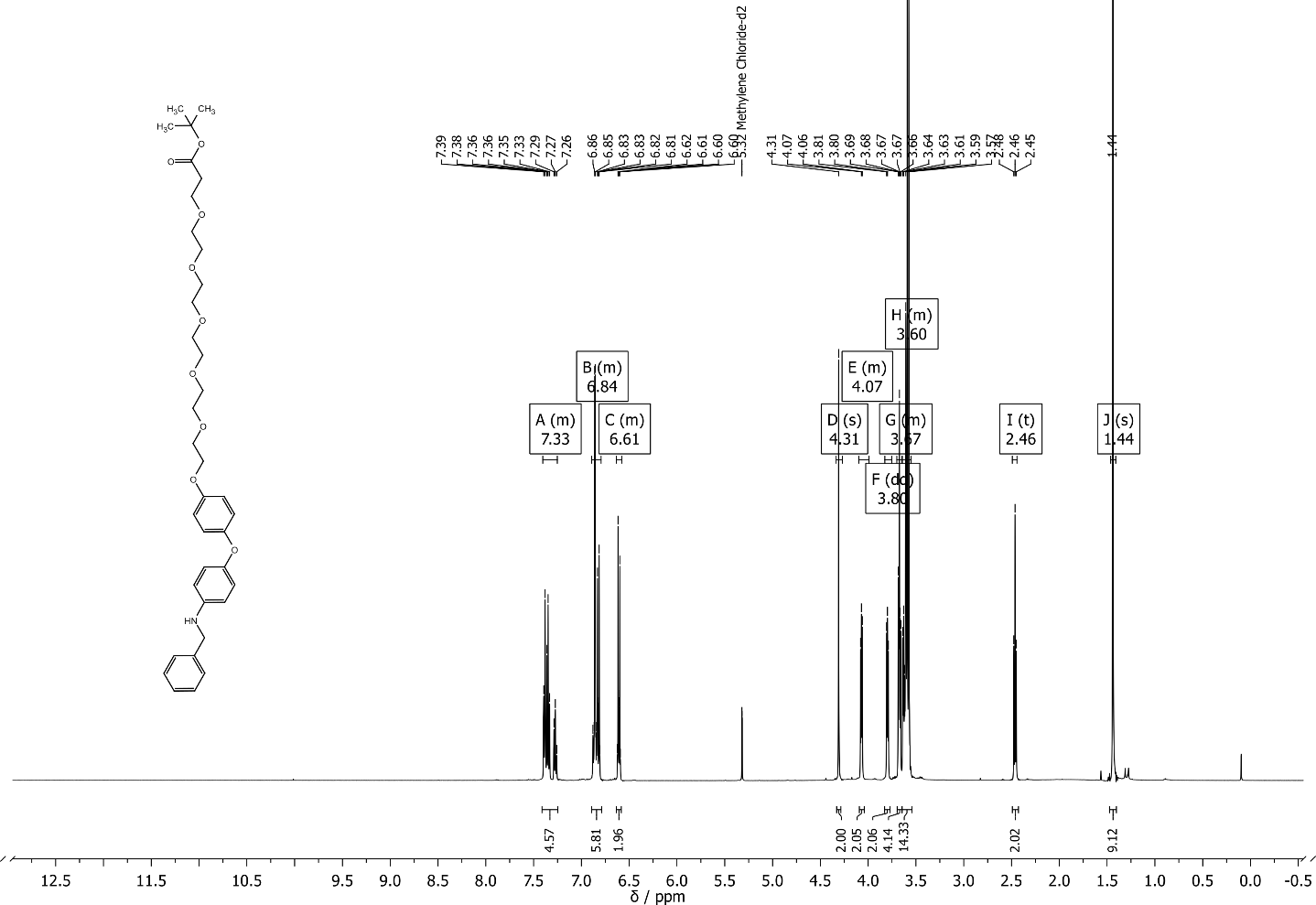

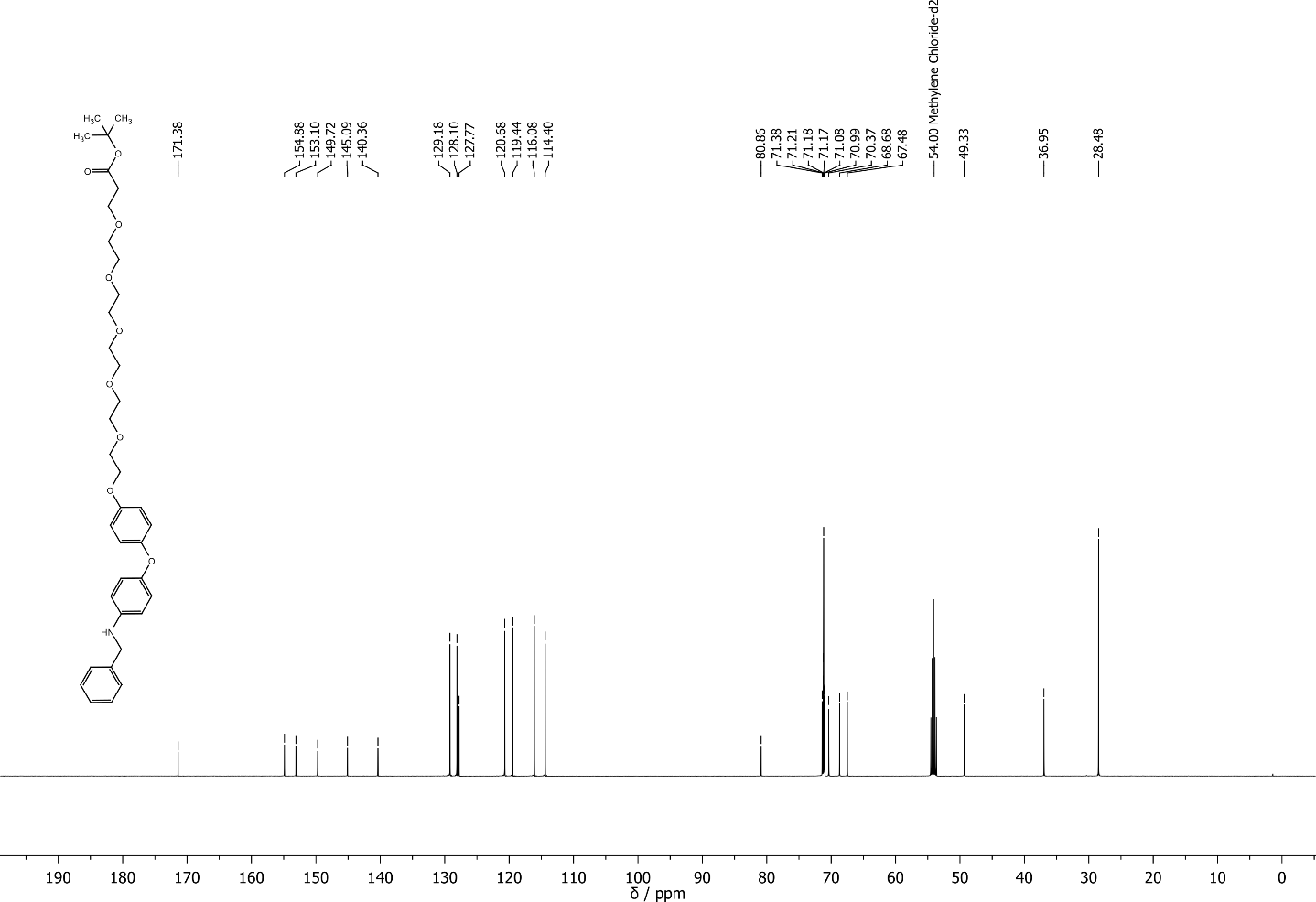

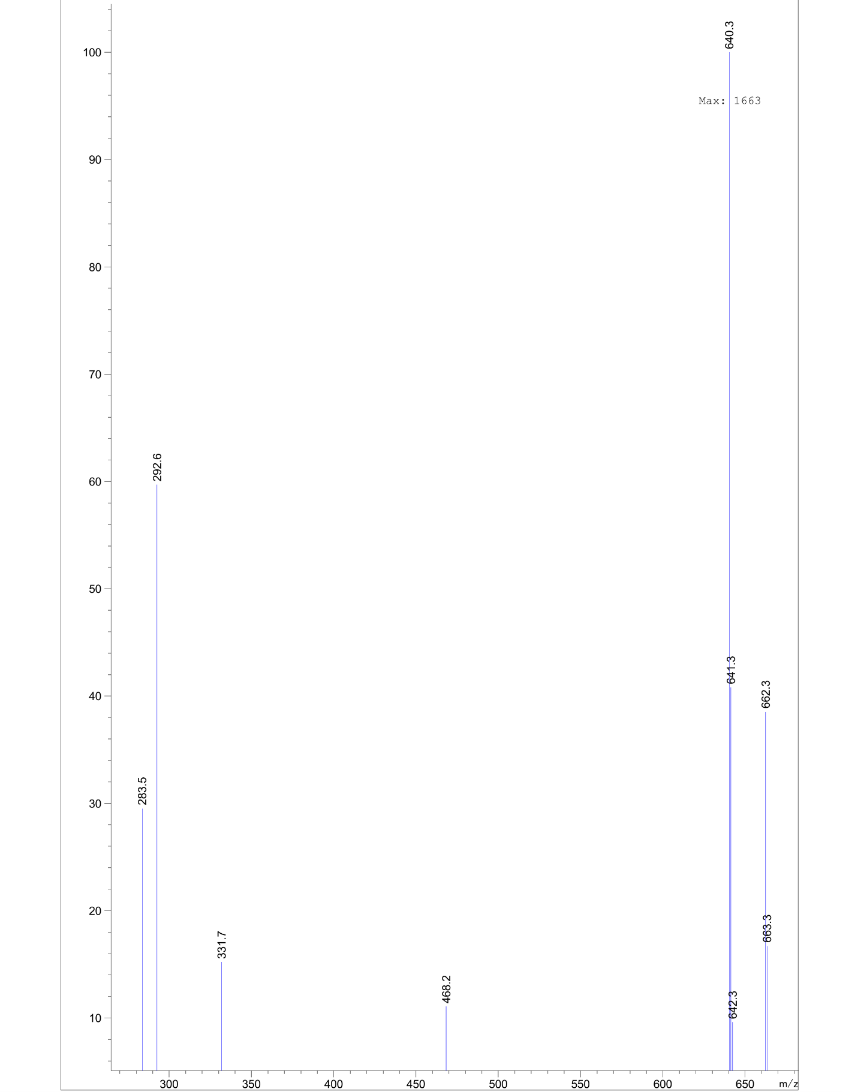

(2S,4R)-1-((S)-1-(4-(4-(benzylamino)phenoxy)phenoxy)-14-(tert-butyl)-12-oxo-3,6,9-trioxa-13-azapentadecan-15-oyl)-4-hydroxy-N-(4-(4-methylthiazol-5-yl)benzyl)pyrrolidine-2-carboxamide **S28**

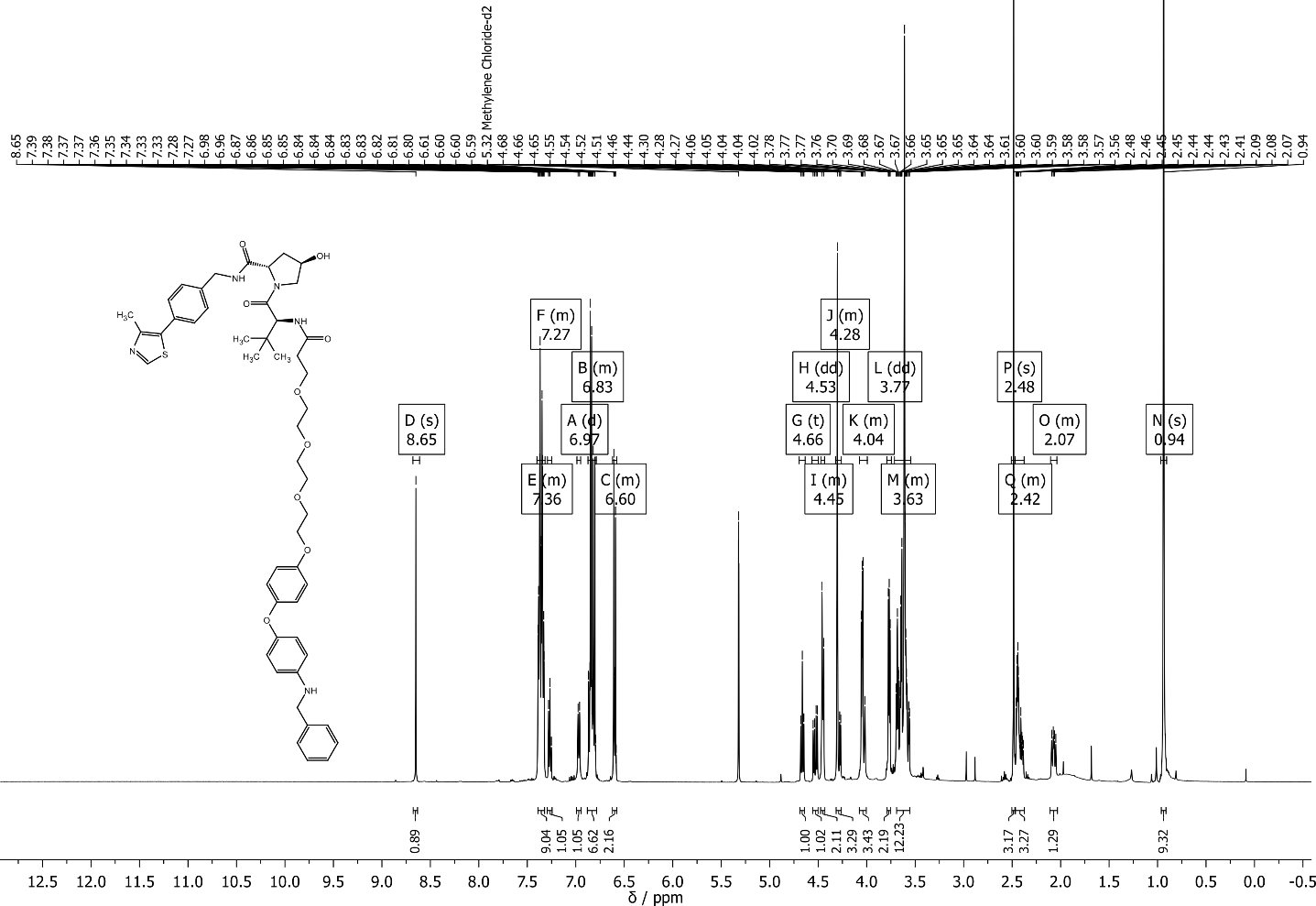

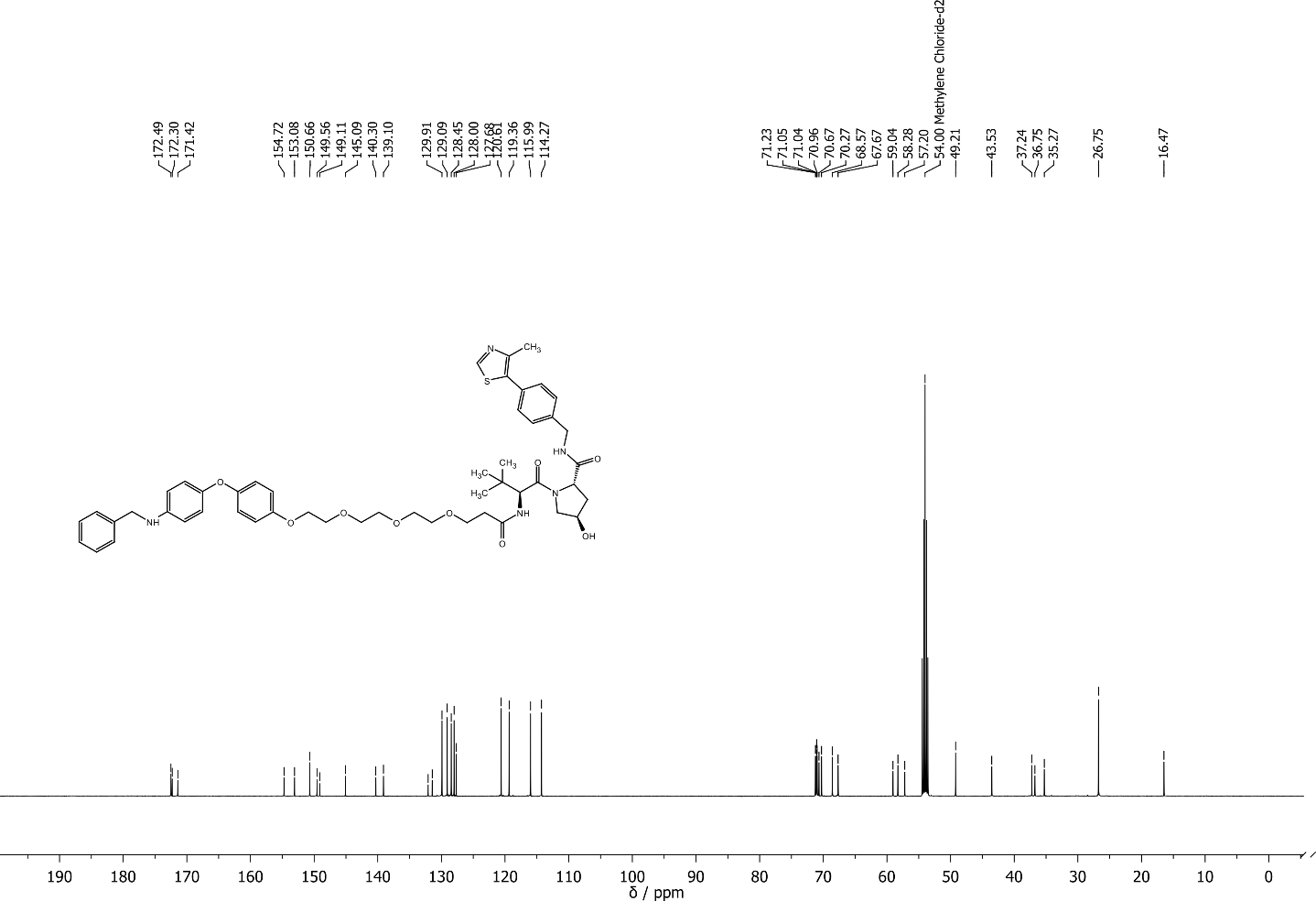

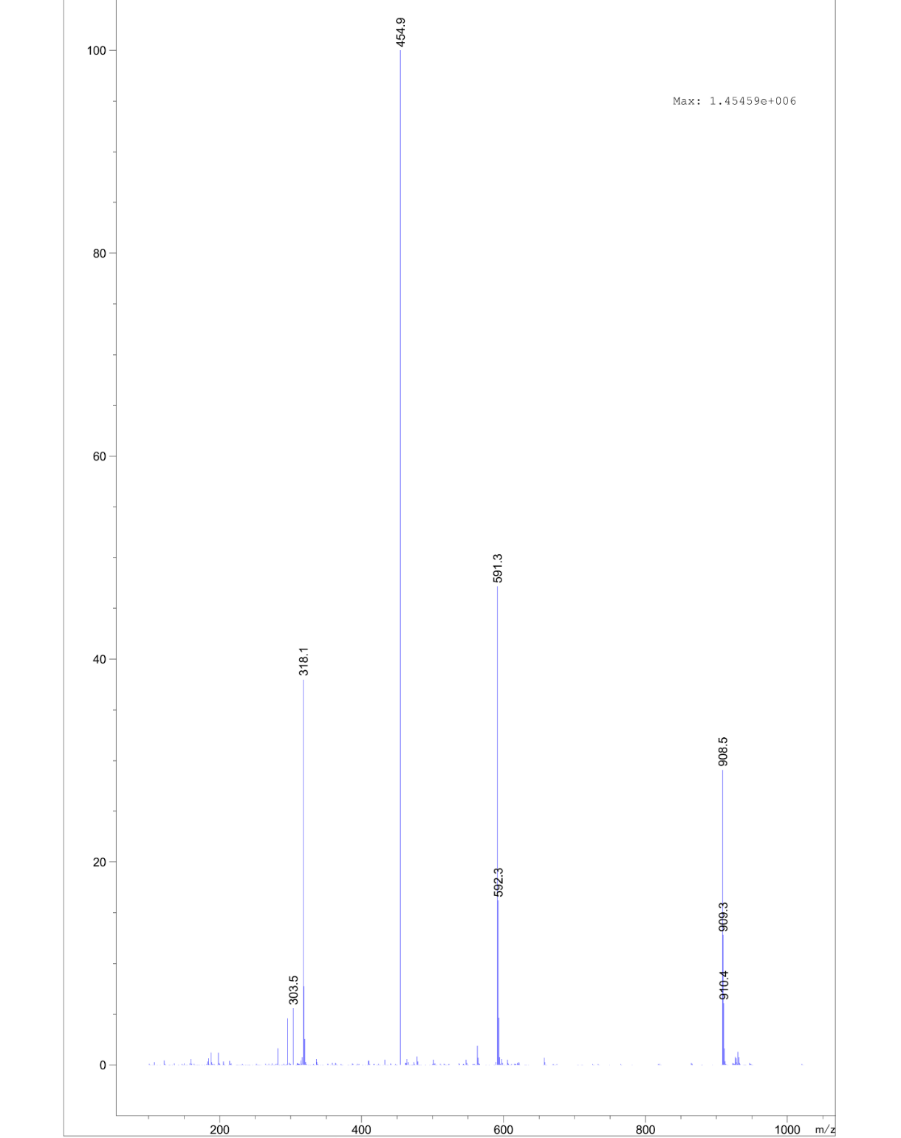

(2S,4R)-1-((S)-1-(4-(4-(benzylamino)phenoxy)phenoxy)-20-(tert-butyl)-18-oxo-3,6,9,12,15-pentaoxa-19-azahenicosan-21-oyl)-4-hydroxy-N-(4-(4-methylthiazol-5-yl)benzyl)pyrrolidine-2-carboxamide **S29**

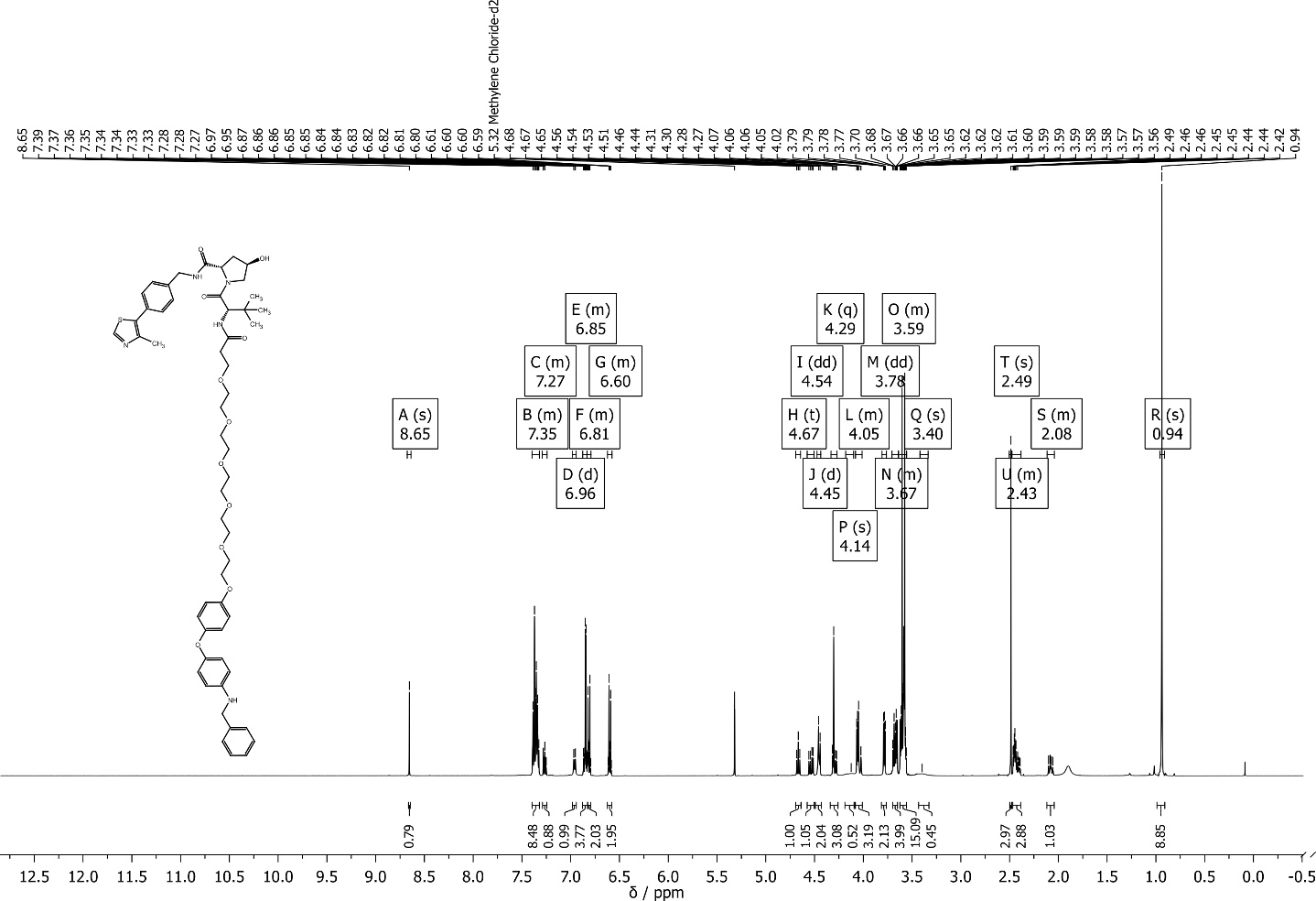

(2S,4R)-1-((S)-1-(4-(4-(N-benzyl-2-chloroacetamido)phenoxy)phenoxy)-14-(tert-butyl)-12-oxo-3,6,9-trioxa-13-azapentadecan-15-oyl)-4-hydroxy-N-(4-(4-methylthiazol-5-yl)benzyl)pyrrolidine-2-carboxamide **2b**

(2S,4R)-1-((S)-1-(4-(4-(N-benzyl-2-chloroacetamido)phenoxy)phenoxy)-20-(tert-butyl)-18-oxo-3,6,9,12,15-pentaoxa-19-azahenicosan-21-oyl)-4-hydroxy-N-(4-(4-methylthiazol-5-yl)benzyl)pyrrolidine-2-carboxamide **2c**

N-(2-(2-(4-(4-(N-benzyl-2-chloroacetamido)phenoxy)phenoxy)ethoxy)ethyl)-5-((3aS,4S,6aR)-2-oxohexahydro-1H-thieno[3,4-d]imidazol-4-yl)pentanamide **Biotin-CCW16**

tert-butyl (4-(4-(4-(benzylamino)phenoxy)phenoxy)butyl)carbamate **S31**

(S)-N-(4-(4-(4-(benzylamino)phenoxy)phenoxy)butyl)-2-(4-(4-chlorophenyl)-2,3,9-trimethyl-6H-thieno[3,2-f][1,2,4]triazolo[4,3-a][1,4]diazepin-6-yl)acetamide **S32**

(S)-N-benzyl-2-chloro-N-(4-(4-(4-(2-(4-(4-chlorophenyl)-2,3,9-trimethyl-6H-thieno[3,2-f][1,2,4]triazolo[4,3-a][1,4]diazepin-6-yl)acetamido)butoxy)phenoxy)phenyl)acetamide **CCW28‑3**
