## Supplemental_Methods_Synthesis for "The cysteine-reactive covalent RNF4 ligand CCW16 induces ferroptosis in AML cells by activation of ROS signaling"

All commercial chemicals and solvents were used without further purification. All reactions were performed in an inert atmosphere (Ar). Product purification was performed on a PuriFlash Flash Column Chromatography System from Interchim using prepacked silica or RP C18 columns.

The synthesized compounds were characterized by ^1^H NMR, ^13^C NMR, and mass spectrometry (ESI). NMR spectra were measured in DMSO*-d_6_* or CD_2_Cl_2_ on a Bruker AV300, AV500 or DPX600 spectrometer. Chemical shifts δ are reported in parts per million (ppm). Determination of the compound purity and mass by HPLC was carried out on an Agilent 1260 Infinity II device with a 1260 DAD HS detector (G7117C; 254 nm, 280 nm, 320 nm) and a LC/MSD device (G6125B). The compounds were analyzed on a Poroshell 120 EC-C18 (Agilent, 3 x 150 mm, 2.7 µm) reversed phase column using 0.1% formic acid in water (A) and 0.1% formic acid in acetonitrile (B) as a mobile phase. The following gradient was used: Method A (ESI pos. 100-1000): 0 min. 5% B - 2 min. 5% B - 7 min. 98% B (flow rate of 0.5 mL/min.). Method B (ESI pos. 100-1300): 0 min. 5% B - 2 min. 5% B - 7 min. 98% B (flow rate of 0.5 mL/min.). UV-detection was performed at 254, 320 nm and all compounds used for further biological characterizations showed >95% purity. HRMS measurements (method1) were executed on a MicrOTOF qII (ESI+) using the Hystar and OTOF control software from Bruker with a Thermo Ultimate 3000 using a flow of 200 µL/min with 50% acetonitrile and 50% water with 0.1% formic acid. Lockmass correction was performed with Bruker Data Analysis software with Carbamezepine (m/z = 237.10224 [M+H]^+^), Flunarizine (m/z = 405.21368 [M+H]^+^) and Reserpine (m/z = 609.28066 [M+H]^+^). HRMS (method2; Orbitrap) measurements were executed on a Exploris 480 Thermo (Bremen, Germany) mass spectrometer equipped with a heated electrospray source (HESI) and coupled to a liquid chromatography system Vanquish VF-P10-A binary pump, VF-A10-A auto sampler which was set to 10 °C and which was equipped with a 25 µL injection syringe and a 100 µL sample loop. Instead of a column a 0.18 mm, 600 mm length capillary was installed within the column compartment VH-C10-A. For automated direct infusion 2.0 µL sample was injected using the flow gradient in Table 1 with 90% pure acetonitrile and 10% water with 0.1% formic acid. The flow was switched according to Table 1 from waste to the MS and back to the waste, to prevent source contamination. For monitoring two full scan modes were selected with the following parameters. Polarity: positive; scan range: 100 to 1500 *m*/*z*; resolution: 480,000; AGC target: “Standard”; maximum IT: “Auto”. General settings: sheath gas flow rate: 20; auxiliary gas flow rate 5; sweep gas flow rate: 1; spray voltage: 3.5 kV; capillary temperature: 325 °C; S-lens RF level: 50; auxiliary gas heater temperature: 125 °C. For negative mode, all values were kept instead of the spray voltage which was set to 2.5 kV.

Table 1

| **time** | **Flow** | **MS Aquisition** | **Waste valve** |
| --- | --- | --- | --- |
| 0 | 0.1 | off | waste |
| 0.5 | 0.1 | on | waste → MS |
| 0.6 | 0.02 | on | MS |
| 4 | 0.02 | on | MS → waste |
| 4.1 | 1 | off | waste |
| 5 | 1 | off | waste |
| 5.1 | 0.2 | off | waste |
| 5.5 | 0.2 | off | waste → MS |
| 5.8 | 0.2 | off | MS → waste |
| 6 | 0.2 | off | waste |

**Synthetic procedures**

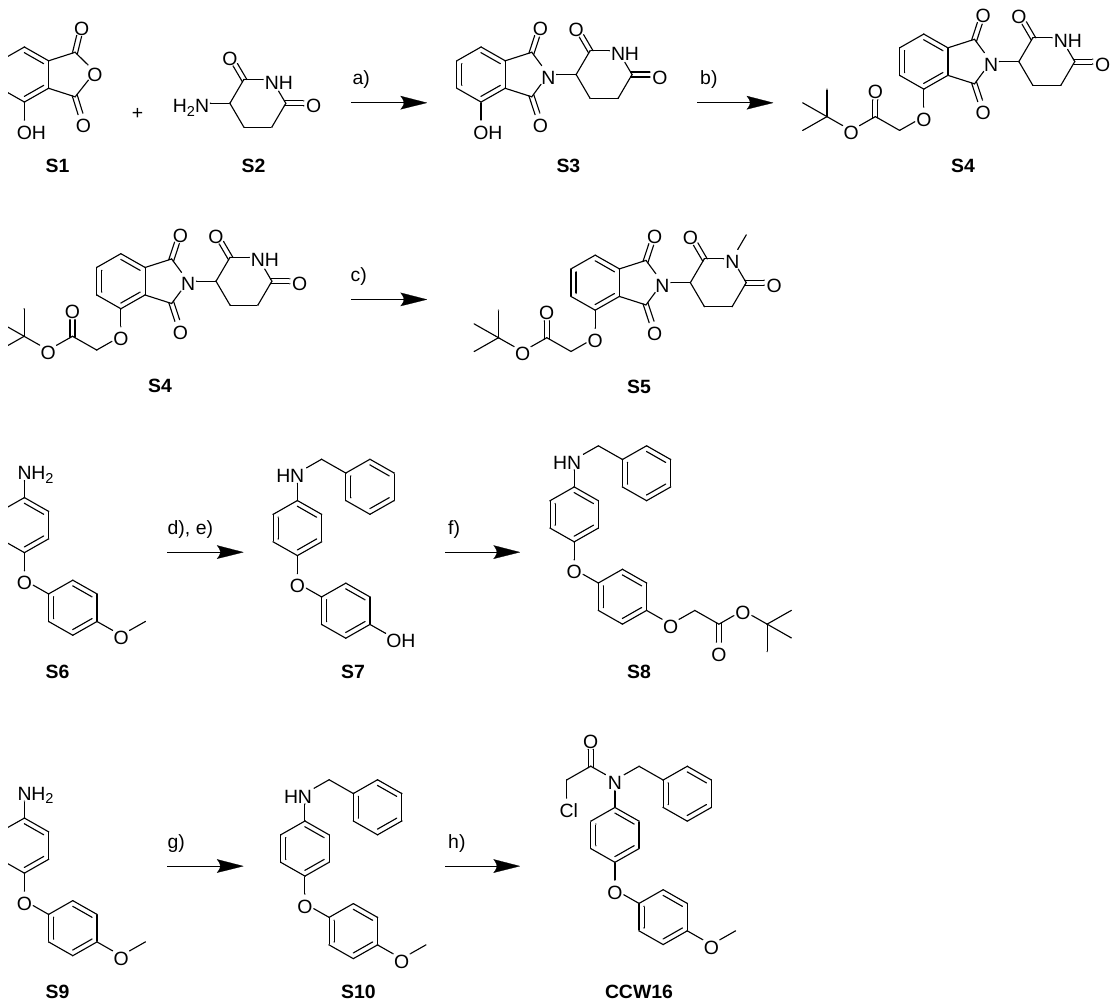

**Figure S##:** Synthesis of CRBN and RNF4 ligands **S4**, **S7** and **CCW16**. a) KOAc, AcOH, reflux, 3 h; b) t*ert*-butylbromoacetate, K_2_CO_3_, DMF, r.t., 4 h; c) K_2_CO_3_, MeI; DMF, r.t., 5 h; d) BBr_3_, DCM, -10 °C, then r.t., 2 h; e) benzaldehyde, AcOH, DCM, .r.t, 30 min, then NaBH(AcO)_3_, r.t., 21 h; f) NaH, THF, 0 °C, 15 min, then *tert*-butyl 2-bromoacetate, r.t., 3 h; g) benzaldehyde, NaBH(AcO)_3_, AcOH, DCM, .r.t, overnight; h) 2-chloroacetyl chloride. TEA, DCM, r.t., overnight.

2-(2,6-dioxopiperidin-3-yl)-4-hydroxyisoindoline-1,3-dione **S3**

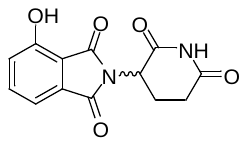

A solution of 4-hydroxyisobenzofuran-1,3-dione **S1** (2.00 g, 12.2 mmol, 1.0eq), 3-aminopiperidine-2,6-dione hydrochloride **S2** (2.21 g, 13.4 mmol, 1.1eq) and potassium acetate (3.71 g, 37.8 mmol, 3.1eq) in acetic acid (40 mL) was refluxed for 3 h. The reaction mixture was poured on water and the precipitate was filtered. After washing with water and hexanes, the title compound was isolated as an off-white solid (2.62 g, 78%).

**^1^H NMR** (500 MHz, DMSO*-d_6_*): δ = 11.16 (s, 1H), 11.07 (s, 1H), 7.65 (dd, *^3^J* = 8.3 Hz, *^3^J* = 7.3 Hz, 1H), 7.32 (d, *^3^J* = 7.1 Hz, 1H), 7.25 (d, *^3^J* = 8.4 Hz, 1H), 5.07 (dd, *^3^J* = 12.8 Hz, *^3^J* = 5.4 Hz, 1H), 2.94-2.82 (m, 1H), 2.65-2.51 (m, 2H), 2.08-1.97 (m, 1H).

**^13^C NMR** (126 MHz, DMSO*-d_6_*): δ = 172.8, 169.97, 166.99, 165.8, 155.5, 136.3, 133.1, 123.6, 114.4, 114.2, 48.6, 30.9, 22.0.

MS (ESI+):

275.0	[(M+H)^+^	calc. 275.07]

297.0	[(M+Na)^+^	calc. 297.05]

338.0	[(M+Na+ACN)^+^	calc. 338.08]

*tert*-butyl 2-((2-(2,6-dioxopiperidin-3-yl)-1,3-dioxoisoindolin-4-yl)oxy)acetate **S4**

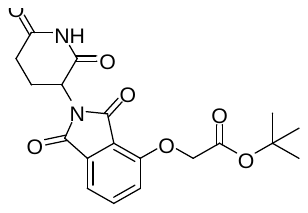

*tert*-Butylbromoacetate (1.1 mL, 7.7 mmol, 1.1eq) was added to a solution of 2-(2,6-dioxopiperidin-3-yl)-4-hydroxyisoindoline-1,3-dione **S3** (2.00 g, 7.29 mmol, 1.0eq)) and potassiumcarbonate (2.02 g, 14.59 mmol, 2.0eq) in DMF (40 mL). The reaction mixture was stirred for 4 h. Ethylacetate and water were added and the layers were separated. The aqueous layer was extracted with ethylacetate (4x) and the combined organic layers were dried with MgSO_4_. The solvent was evaporated under reduced pressure and the residue was purified by flash column chromatography (H/EE). The title compound was isolated as a colorless solid (1.71 g, 60%).

**^1^H NMR** (500 MHz, DMSO*-d_6_*): δ = 11.10 (s, 1H), 7.80 (dd, *^3^J* = 8.5 Hz, *^3^J* = 7.3 Hz, 1H), 7.48 (d, *^3^J* = 7.1 Hz, 1H), 7.38 (d, *^3^J* = 8.5 Hz, 1H), 5.10 (dd, *^3^J* = 12.8 Hz, *^3^J* = 5.5 Hz, 1H), 4.97 (s, 2H), 2.89 (ddd, *^3^J* = 16.9 Hz, *^3^J* = 13.9 Hz, *^4^J* = 5.4 Hz, 1H), 2.65-2.51 (m, 2H), 2.08-2.01 (m, 1H), 1.43 (s, 9H).

**^13^C NMR** (126 MHz, DMSO*-d_6_*): δ = 172.8, 169.9, 167.1, 166.7, 165.1, 155.0, 136.8, 133.3, 119.97, 116.5, 115.9, 81.9, 65.5, 48.8, 30.9, 27.7, 21.95.

MS (ESI+):

333.0	[(M-^t^Bu+2H)^+^	calc. 333.07]

411.0	[(M+Na)^+^	calc. 411.19]

799.3	[(2M+Na)^+^	calc. 799.25]

tert-butyl 2-((2-(1-methyl-2,6-dioxopiperidin-3-yl)-1,3-dioxoisoindolin-4-yl)oxy)acetate **S5**

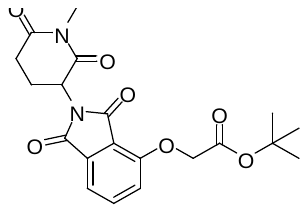

Methyliodide (2M in MTBE, 965 µL, 5.0eq) was added to a solution of *tert*-butyl 2-((2-(2,6-dioxopiperidin-3-yl)-1,3-dioxoisoindolin-4-yl)oxy)acetate **S4** (150 mg, 386 µmol, 1.0eq) and K_2_CO_3_ (107 mg, 772 µmol, 2.0eq) in DMF (2 mL). The reaction mixture was stirred for 5 h. Ethylacetate and water were added and the layers were separated. The aqueous layer was extracted with ethylacetate (4x) and the combined organic layers were dried with Na_2_SO_4_. The solvent was evaporated under reduced pressure and the residue was purified by flash column chromatography (DCM/MeOH). The title compound was isolated as a colorless solid (75 mg, 57%).

**^1^H NMR** (500 MHz, DCM): δ = 7.69 (dd, *^3^J* = 8.5 Hz, *^3^J* = 7.3 Hz, 1H), 7.49 (dd, *^3^J* = 7.3, 0.5 Hz, 1H), 7.11 (d, *^3^J* = 8.2 Hz, 1H), 4.97-4.92 (m, 1H), 4.76 (s, 2H), 3.16 (s, 3H), 2.97-2.88 (m, 1H), 2.81-2.71 (m, 2H), 2.13-2.04 (m, 1H), 1.48 (s, 9H).

**^13^C NMR** (126 MHz, DCM): δ = 171.7, 169.4, 167.5, 167.4, 166.1, 156.0, 136.8, 134.6, 120.0, 118.1, 117.0, 83.4, 67, 50.5, 32.5, 28.3, 27.5, 22.4.

MS (ESI+):

347.0	[(M-^t^Bu+2H)^+^	calc. 347.08]

425.0	[(M+Na)^+^	calc. 425.13]

827.3	[(2M+Na)^+^	calc. 827.27]

4-(4-(benzylamino)phenoxy)phenol **S7** via 4-(4-aminophenoxy)phenol

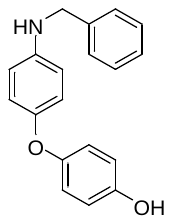

BBr_3_ (34.8 mL, 1 M/DCM, 3.0eq) was added to a solution of 4-(4-methoxyphenoxy)aniline **S6** (2.50 g, 11.6 mmol, 1.0eq) in DCM (40 mL) at -10 °C. The reaction mixture was stirred at r.t. for 2 h. The reaction was quenched with a saturated solution of NaHCO_3_. The layers were separated and the aqueous layer was extracted with ethylacetate (3x). The combined organic layers were dried with MgSO_4_ and the solvent was removed under reduced pressure to yield the title compound as a brown solid (1.49 g, 64%). The crude product was used without further purification in the next step.

Acetic acid (0.950 mL, 16.65 mmol, 1.1eq) and benzaldehyde (0.5578 mL, 5.47 mmol, 1.1eq) were added to a solution of crude 4-(4-aminophenoxy)phenol (1.00 g, 4.97 mmol) in DCM (50 mL). The reaction mixture was stirred for 30 min and sodium triacetoxyborohydride (1.58 g, 7.45 mmol, 1.5eq) was added. The mixture was stirred for 21 h. All volatiles were removed under reduced pressure. Flash column chromatography (H/EE) yielded the title compound (0.933 g, 64%) as colorless oil.

**^1^H NMR** (500 MHz, CD_2_Cl_2_): δ = 7.40-7.25 (m, 5H), 6.84-6.79 (m, 4H), 6.77-6.73 (m, 2H), 6.64-6.59 (m, 2H), 4.31 (s, 2H).

**^13^C NMR** (126 MHz, CD_2_Cl_2_): δ = 152.8, 151.6, 149.9, 144.7, 140.1, 129.1, 128.1, 127.7, 120.5, 119.6, 116.6, 114.5, 49.4.

MS (ESI+):

292.1	[(M+H)^+^	calc. 292.14]

293.0	[(M+2H)^2+^	calc. 293.14]

tert-butyl 2-(4-(4-(benzylamino)phenoxy)phenoxy)acetate **S8**

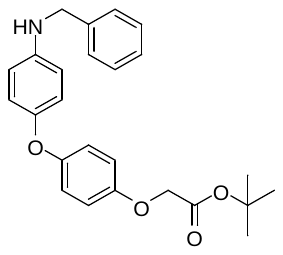

Sodium hydride (0.66 mg, 16.5 mmol, 3.0eq) was added to a solution of 4-(4-(benzylamino)phenoxy)phenol **S7** (1.60 mg, 5.49 mmol, 1.0eq) in THF (50 mL) at 0 °C. The reaction mixture was stirred at for 15 min. *tert*-butyl 2-bromoacetate (0.81 mL, 5.49 mmol, 1.0eq) was added and the reaction was stirred at r.t. for 3 h. All volatiles were removed under reduced pressure and the residue was dissolved with DCM and water. The layers were separated and the aqueous layer was extracted with DCM (2x). The combined organic layers were dried with MgSO_4_ and the crude material was purified using flash column chromatography (CH/EA). The title compound was isolated as yellow oil (1.32 mg, 59%).

**^1^H NMR** (500 MHz, CD_2_Cl_2_): δ = 7.41-7.25 (m, 5H), 6.89-6.78 (m, 6H), 6.64-6.59 (m, 2H), 4.46 (s, 2H), 4.31 (s, 2H), 1.48 (s, 9H).

**^13^C NMR** (126 MHz, CD_2_Cl_2_): δ = 168.6, 153.9, 153.6, 149.4, 145.0, 140.2, 129.1, 128.1, 127.7, 120.8, 119.2, 116.1, 114.4, 82.6, 66.9, 49.3, 28.3.

MS (ESI+):

406.1	[(M+H)^+^	calc. 406.197]

407.0	[(M+2H)^2+^	calc. 407.20]

428.1	[(M+Na)^+^	calc. 428.19]

N-benzyl-4-(4-methoxyphenoxy)aniline **S9**

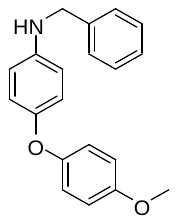

NaBH(OAc)_3_ (2.88 g, 13.6 mmol, 1.5eq) was added to a solution of 4-(4-methoxyphenoxy)aniline **S6** (1.95 g, 9.06 mmol, 1.0eq), benzaldehyde ( 1.1 mL, 10.9 mmol, 1.2eq) and acetic acid (622 µL, 10.9 mmol, 1.2eq) in DCM (80 mL). The reaction mixture was stirred overnight. The solvent removed under reduced pressure and the residue was dissolved in ethyl acetate and a sat. NaHCO_3_ solution. The layers were separated and the aqueous layer was extracted with ethyl acetate (2x). The combined organic layers were washed with water and dried using MgSO_4_. The solvent was removed under reduced pressure and the crude material was purified using flash column chromatography (CH/EA). The title compound was isolated as a yellow oil and was used directly for the next step.

N-benzyl-2-chloro-N-(4-(4-methoxyphenoxy)phenyl)acetamide **CCW16**

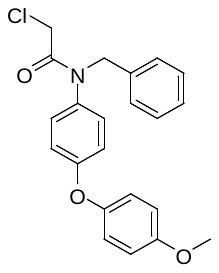

2-chloroacetyl chloride (238 µL, 2.99 mmol, 4.0eq) and TEA (416 µL, 2.99 mmol, 4.0eq) were added to a solution of N-benzyl-4-(4-methoxyphenoxy)aniline **S10** (288 mg, 0.747 mmol, 1.0eq) in DCM (20 mL). The reaction mixture was stirred overnight. The reaction was quenched with water and the layers were separated. The aqueous layer was extracted with DCM (2x) and the combined organic layers were dried with MgSO_4_. Reversed phase flash column chromatography (ACN/water) yielded the title compound as a yellow solid (120 mg, 42%).

**^1^H NMR** (500 MHz, CD_2_Cl_2_): δ = 7.32-7.19 (m, 5H), 7.01-6.94 (m, 4H), 6.93-6.89 (m, 2H), 6.89-6.85 (m, 2H), 4.86 (s, 2H), 3.90 (s, 2H), 3.80 (s, 3H).

**^13^C NMR** (126 MHz, CD_2_Cl_2_): δ = 166.7, 159.4, 157.1, 149.6, 137.5, 135.5, 130.1, 129.3, 128.98, 128.1, 121.8, 118.4, 115.5, 56.2, 54.1, 42.9.

HPLC: R_t_ = 5.18 min (method A): Purity: >98% (254 nm); >98% (320 nm)

MS (ESI+):

382.0	[(M+H)^+^	calc. 382.12]

384.0	[(M+3H)^3+^	calc. 384.13]

HRMS (ESI+):

382.1199	[(M+H)^+^	calc. 382.1205]

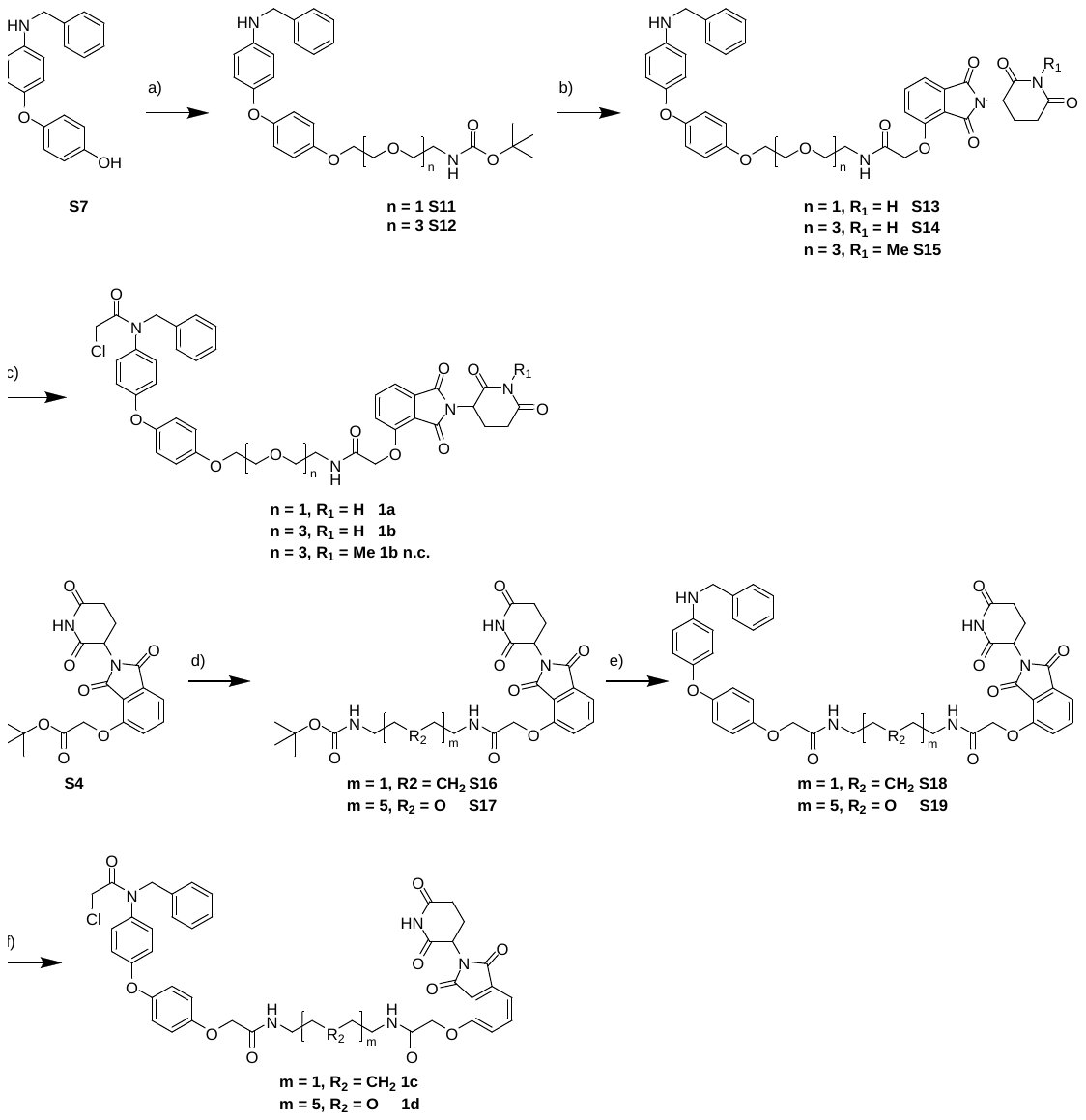

**Figure S##:** Synthesis of CRBN based PROTACs **1a-d** and negative control **1b n.c.**: a) appropriate linker bromide, K_2_CO_3_, Acetone, reflux, overnight; b) 1. intermediate **S4** or **S5**, TFA, DCM, r.t., 2-3 h; 2. DIPEA, HATU, DMF, r.t., 18 h; c) 2-chloroacetyl chloride, TEA, r.t., 16-18 h; d) 1. TFA, DCM, r.t., 2 h; 2. DIPEA, HATU, DMF, r.t., 18 h; e) 1. intermediate **S8**, TFA, DCM, r.t., 2-3 h; 2. DIPEA, HATU, DMF, r.t., 12-18 h; f) 2-chloroacetyl chloride, TEA, r.t., 17-20 h.

tert-butyl (2-(2-(4-(4-(benzylamino)phenoxy)phenoxy)ethoxy)ethyl)carbamate **S11**

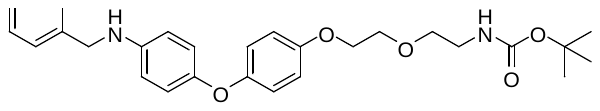

A solution of 4-(4-(benzylamino)phenoxy)phenol **S7** (36 mg, 124 µmol, 1.0eq), tert-butyl (2-(2-bromoethoxy)ethyl)carbamate (50 mg, 186 µmol, 1.5eq) and potassium carbonate (52 mg, 373 µmol, 3.0eq) in Acetone (5 mL) was refluxed overnight. The solvent was removed under reduced pressure. The residue was dissolved with DCM and water. The layers were separated and the aqueous layer was extracted with DCM twice. The combined organic layers were dried with MgSO_4_ and the solvent was removed under reduced pressure. Reversed phase column chromatography (ACN/water) yielded the title compound as a yellow oil (26 mg, 44%).

**^1^H NMR** (500 MHz, CD_2_Cl_2_): δ = 7.41-7.25 (m, 5H), 6.90-6.80 (m, 6H), 6.64-6.58 (m, 2H), 5.01 (s, 1H), 4.31 (s, 2H), 4.10 (s, 1H), 4.08-4.04 (m, 2H), 3.81-3.74 (m, 2H), 3.58 (t, *^3^J* = 5.3 Hz, 2H), 3.30 (dd, *^3^J* = 10.7 Hz, *^3^J* = 5.4 Hz, 2H), 1.43 (s, 9H).

**^13^C NMR** (126 MHz, CD_2_Cl_2_): δ = 156.4, 154.7, 153.1, 149.6, 145.2, 140.3, 129.1, 128.0, 127.7, 120.6, 119.4, 116.0, 114.3, 70.9, 70.1, 68.6, 55.7, 49.2, 40.96, 28.7.

MS (ESI+):

379.2	[(M-Boc+2H)^+^	calc. 379.20]

423.2	[(M-Boc+2Na)^3+^	calc. 423.17]

501.3	[(M+Na)^+^	calc. 501.24]

tert-butyl (2-(2-(2-(2-(4-(4-(benzylamino)phenoxy)phenoxy)ethoxy)ethoxy)ethoxy)ethyl)carbamate **S12**

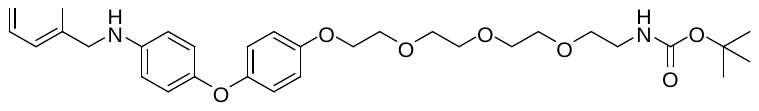

A solution of 4-(4-(benzylamino)phenoxy)phenol **S7** (61 mg, 211 µmol, 1.0eq), tert-butyl (2-(2-(2-(2-bromoethoxy)ethoxy)ethoxy)ethyl)carbamate (75 mg, 211 µmol, 1.0eq) and potassium carbonate (87 mg, 632 µmol, 3.0eq) in Acetone (10 mL) was refluxed overnight. The solvent was removed under reduced pressure. The residue was dissolved with DCM and water. The layers were separated and the aqueous layer was extracted with DCM twice. The combined organic layers were dried with MgSO_4_ and the solvent was removed under reduced pressure. Reversed phase column chromatography (ACN/water) yielded the title compound as a colorless oil (88 mg, 74%).

**^1^H NMR** (500 MHz, CD_2_Cl_2_): δ = 7.41-7.33 (m, 4H), 7.30-7.26 (m, 1H), 6.90-6.80 (m, 6H), 6.64-6.59 (m, 2H), 5.16 (s, 1H), 4.31 (s, 2H), 4.13 (s, 1H), 4.07 (dd, *^3^J* = 5.4 Hz, *^3^J* = 4.0 Hz, 2H), 3.80 (dd, *^3^J* = 5.3 Hz, *^3^J* = 4.1 Hz, 2H), 3.69 (dd, *^3^J* = 6.1 Hz, *^3^J* = 3.2 Hz, 2H), 3.67-3.57 (m, 6H), 3.51 (t, *^3^J* = 5.2 Hz, 2H), 3.27 (dd, *^3^J* = 10.3 Hz, *^3^J* = 5.1 Hz, 2H), 1.43 (s, 9H).

**^13^C NMR** (126 MHz, CD_2_Cl_2_): δ = 156.4, 154.7, 153.1, 149.6, 145.1, 140.3, 129.1, 127.99, 127.7, 120.6, 119.3, 115.99, 114.3, 79.3, 71.3, 71.1, 71.0, 70.8, 70.7, 70.3, 68.6, 49.2, 40.99, 28.7.

MS (ESI+):

467.1	[(M-Boc+2H)^+^	calc. 467.25]

511.2	[(M-Boc+2Na)^+^	calc. 511.23]

567.3	[(M+H)^+^	calc. 567.31]

N-(2-(2-(4-(4-(benzylamino)phenoxy)phenoxy)ethoxy)ethyl)-2-((2-(2,6-dioxopiperidin-3-yl)-1,3-dioxoisoindolin-4-yl)oxy)acetamide **S13**

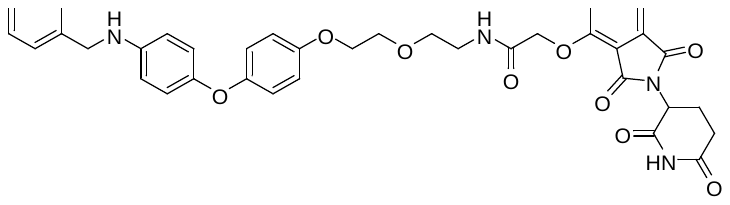

A solution of tert-butyl (2-(2-(4-(4-(benzylamino)phenoxy)phenoxy)ethoxy)ethyl)carbamate **S11** (52 mg, 109 µmol, 1.1eq) in DCM/TFA (4 mL, 1/1) was stirred for 2 h. All volatiles were removed under reduced pressure. DCM was added and the solvent was removed under reduced pressure. This procedure was repeated twice.

A solution of tert-butyl 2-((2-(2,6-dioxopiperidin-3-yl)-1,3-dioxoisoindolin-4-yl)oxy)acetate **S4** (38 mg, 99 µmol, 1.0eq) in DCM/TFA (4 mL, 1/1) was stirred for 2 h. All volatiles were removed under reduced pressure. DCM was added and the solvent was removed under reduced pressure. This procedure was repeated twice.

A solution of the crude amine, DIPEA (43 µL, 247 µmol, 2.5eq), the crude acid and HATU (45 mg, 119 µmol, 1.2eq) in DMF (2 mL) was stirred for 18 h. DCM, water and a sat. solution of NaHCO_3_ were added and the layers were separated. The aqueous layer was extracted with DCM (3x) and the combined organic layers were dried with MgSO_4_. The solvent was evaporated under reduced pressure and the residue was purified by reversed flash column chromatography to yield the title compound as a yellow solid (40 mg, 58%).

**^1^H NMR** (500 MHz, CD_2_Cl_2_): δ = 8.45 (s, 1H), 7.71 (dd, *^3^J* = 8.4 Hz, *^3^J* = 7.4 Hz, 1H), 7.54 (t, *^3^J* = 5.3 Hz, 1H), 7.50 (dd, *^3^J* = 7.3 Hz, *^4^J* = 0.4 Hz, 1H), 7.41-7.32 (m, 4H), 7.29-7.24 (m, 1H), 7.19 (d, *^3^J* = 8.4 Hz, 1H), 6.82-6.77 (m, 6H), 6.62-6.58 (m, 2H), 4.90 (dd, *^3^J* = 12.4 Hz, *^3^J* = 5.4 Hz, 1H), 4.62 (s, 2H), 4.30 (s, 2H), 4.12 (s, 1H), 4.07 (dd, *^3^J* = 6.0 Hz, *^3^J* = 3.9 Hz, 2H), 3.84-3.78 (m, 2H), 3.68 (q, *^3^J* = 5.4 Hz, 2H), 3.59-3.52 (m, 2H), 2.82-2.60 (m, 3H), 2.11-2.04 (m, 1H).

**^13^C NMR** (126 MHz, CD_2_Cl_2_): δ = 171.6, 168.9, 167.3, 167.2, 166.4, 155.0, 154.6, 153.1, 149.4, 145.1, 140.3, 137.5, 134.1, 129.1, 128.0, 127.7, 120.7, 120.1, 119.2, 118.5, 117.5, 115.95, 114.3, 70.2, 70.0, 68.6, 68.5, 49.8, 49.2, 39.5, 31.9, 23.1.

MS (ESI+):

347.1	[(M/2+H)^+^	calc. 347.13]

693.3	[(M+H)^+^	calc. 693.26]

N-(2-(2-(2-(2-(4-(4-(benzylamino)phenoxy)phenoxy)ethoxy)ethoxy)ethoxy)ethyl)-2-((2-(2,6-dioxopiperidin-3-yl)-1,3-dioxoisoindolin-4-yl)oxy)acetamide **S14**

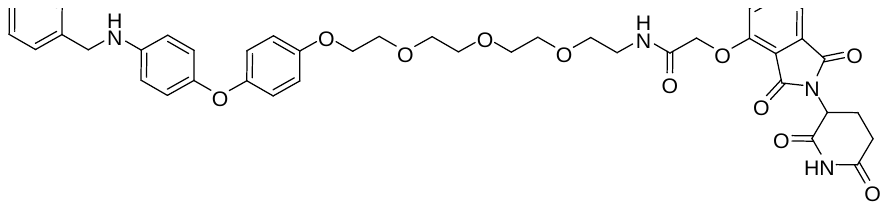

A solution of tert-butyl (2-(2-(2-(2-(4-(4-(benzylamino)phenoxy)phenoxy)ethoxy)ethoxy)ethoxy)ethyl)carbamate **S12** (58 mg, 102 µmol, 1.1eq) in DCM/TFA (4 mL, 1/1) was stirred for 2 h. All volatiles were removed under reduced pressure. DCM was added and the solvent was removed under reduced pressure. This procedure was repeated twice.

A solution of tert-butyl 2-((2-(2,6-dioxopiperidin-3-yl)-1,3-dioxoisoindolin-4-yl)oxy)acetate **S4** (93 mg, 36 µmol, 1.0eq) in DCM/TFA (4 mL, 1/1) was stirred for 3 h. All volatiles were removed under reduced pressure. DCM was added and the solvent was removed under reduced pressure. This procedure was repeated twice.

A solution of the crude amine, DIPEA (41 µL, 233 µmol, 2.5eq), the crude acid and HATU (42 mg, 112 µmol, 1.2eq) in DMF (2 mL) was stirred for 18 h. DCM, water and a sat. solution of NaHCO_3_ were added and the layers were separated. The aqueous layer was extracted with DCM (3x) and the combined organic layers were dried with MgSO_4_. The solvent was evaporated under reduced pressure and the residue was purified by reverse flash column chromatography to yield the title compound as a yellow oil (21 mg, 29%).

**^1^H NMR** (500 MHz, DMSO*-d_6_*): δ = 8.76 (s, 1H), 7.73 (dd, *^3^J* = 8.4 Hz, *^3^J* = 7.4 Hz, 1H), 7.56 (t, *^3^J* = 4.9 Hz, 1H), 7.52 (d, *^3^J* = 7.3 Hz, 1H), 7.41-7.24 (m, 5H), 7.21 (d, *^3^J* = 8.4 Hz, 1H), 6.87-6.79 (m, 6H), 6.63-6.58 (m, 2H), 4.90 (dd, *^3^J* = 12.4 Hz, *^3^J* = 5.4 Hz, 1H), 4.63 (s, 2H), 4.30 (s, 2H), 4.08-4.04 (m, 2H), 3.81-3.77 (m, 2H), 3.69-3.59 (m, 10H), 3.58-3.46 (m, 2H), 2.86-2.60 (m, 3H), 2.14-2.06 (m, 1H).

**^13^C NMR** (126 MHz, DMSO*-d_6_*): δ = 171.7, 168.9, 167.3, 167.2, 166.5, 155.1, 154.7, 153.1, 149.6, 145.1, 140.3, 137.5, 134.2, 129.1, 128.0, 127.7, 120.6, 120.1, 119.4, 118.5, 117.5, 115.96, 114.3, 71.3, 70.96, 70.83, 70.75, 70.2, 70.0, 68.6, 68.5, 49.8, 49.2, 39.6, 31.9, 23.2.

MS (ESI+):

391.3	[(M/2+H)^+^	calc. 391.16]

781.3	[(M+H)^+^	calc. 781.30]

N-(2-(2-(2-(2-(4-(4-(benzylamino)phenoxy)phenoxy)ethoxy)ethoxy)ethoxy)ethyl)-2-((2-(1-methyl-2,6-dioxopiperidin-3-yl)-1,3-dioxoisoindolin-4-yl)oxy)acetamide **S15**

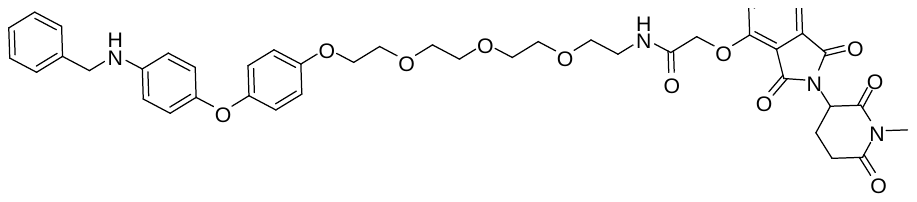

A solution of tert-butyl (2-(2-(2-(2-(4-(4-(benzylamino)phenoxy)phenoxy)ethoxy)ethoxy)ethoxy)ethyl)carbamate **S12** (22 mg, 39 µmol, 1.1eq) in DCM/TFA (4 mL, 1/1) was stirred for 3 h. All volatiles were removed under reduced pressure. DCM was added and the solvent was removed under reduced pressure. This procedure was repeated twice.

A solution of tert-butyl 2-((2-(1-methyl-2,6-dioxopiperidin-3-yl)-1,3-dioxoisoindolin-4-yl)oxy)acetate **S5** (14 mg, 35 µmol, 1.0eq) in DCM/TFA (4 mL, 1/1) was stirred for 3 h. All volatiles were removed under reduced pressure. DCM was added and the solvent was removed under reduced pressure. This procedure was repeated twice.

A solution of the crude amine, DIPEA (25 µL, 141 µmol, 4.0eq), the crude acid and HATU (16 mg, 42 µmol, 1.2eq) in DMF (2 mL) was stirred for 18 h. DCM, water and a sat. solution of NH_4_Cl were added and the layers were separated. The aqueous layer was extracted with DCM (3x) and the combined organic layers were dried with MgSO_4_. The solvent was evaporated under reduced pressure and the residue was purified by reverse flash column chromatography to yield the title compound as a yellow oil (21 mg, 75%).

**^1^H NMR** (500 MHz, CD_2_Cl_2_): δ = 7.75-7.71 (m, 1H), 7.53-7.47 (m, 2H), 7.40-7.32 (m, 4H), 7.27 (t, *^3^J* = 7.1 Hz, 1H), 7.21 (d, *^3^J* = 8.4 Hz, 1H), 6.88-6.78 (m, 6H), 6.66-6.61 (m, 2H), 4.99-4.93 (m, 1H), 4.64 (s, 2H), 4.31 (s, 2H), 4.07-4.02 (m, 2H), 3.80-3.74 (m, 2H), 3.66-3.58 (m, 10H), 3.52 (dd, *^3^J* = 10.7 Hz, *^3^J* = 5.4 Hz, 2H), 3.16 (s, 3H), 3.00-2.87 (m, 1H), 2.85-2.68 (m, 2H), 2.14-2.07 (m, 1H).

**^13^C NMR** (126 MHz, CD_2_Cl_2_): δ = 171.6, 169.4, 167.4, 167.3, 166.6, 155.1, 154.8, 152.9, 150.1, 144.3, 139.8, 137.5, 134.2, 129.1, 128.2, 127.8, 120.5, 120.2, 119.5, 118.6, 117.5, 115.99, 114.9, 71.3, 71.1, 71.0, 70.9, 70.3, 70.0, 68.7, 68.6, 50.6, 49.6, 39.6, 32.4, 27.5, 22.4.

MS (ESI+):

398.3	[(M/2+H)^+^	calc. 398.17]

795.3	[(M+H)^+^	calc. 795.33]

N-benzyl-2-chloro-N-(4-(4-(2-(2-(2-((2-(2,6-dioxopiperidin-3-yl)-1,3-dioxoisoindolin-4-yl)oxy)acetamido)ethoxy)ethoxy)phenoxy)phenyl)acetamide **1a**

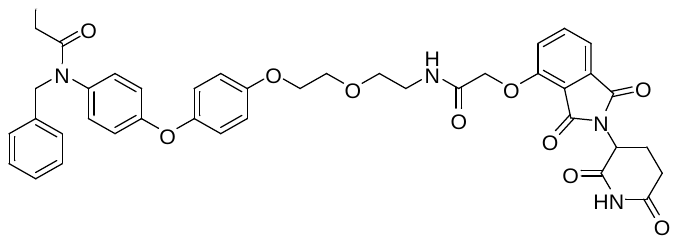

2-chloroacetyl chloride (7 µl, 92 µmol, 1.3eq) and TEA (29 µl, 211 µmol, 3.0eq) were added to a solution of N-(2-(2-(4-(4-(benzylamino)phenoxy)phenoxy)ethoxy)ethyl)-2-((2-(2,6-dioxopiperidin-3-yl)-1,3-dioxoisoindolin-4-yl)oxy)acetamide **S13** (49 mg, 70 µmol, 1.0eq) in DCM (10 mL) at 0 °C. The reaction mixture was stirred for 20 h at r.t. The reaction was quenched with water and the layers were separated. The aqueous layer was extracted with DCM (3x) and the combined organic layers were dried with MgSO_4_. Reversed phase flash column chromatography (ACN/water) yielded the title compound as a colorless oil (40 mg, 74%).

**^1^H NMR** (500 MHz, CD_2_Cl_2_): δ = 8.67-8.48 (m, 1H), 7.75-7.70 (m, 1H), 7.54 (t, *^3^J* = 5.0 Hz, 1H), 7.51 (d, *^3^J* = 7.3 Hz, 1H), 7.31-7.17 (m, 6H), 6.98-6.94 (m, 2H), 6.94-6.87 (m, 4H), 6.87-6.83 (m, 2H), 4.92 (dd, *^3^J* = 12.2 Hz, *^3^J* = 5.3 Hz, 1H), 4.85 (s, 2H), 4.63 (s, 2H), 4.13-4.09 (m, 2H), 3.89 (s, 2H), 3.83 (dd, *^3^J* = 5.5 Hz, *^3^J* = 4.2 Hz, 2H), 3.70 (t, *^3^J* = 5.4 Hz, 2H), 3.62-3.51 (m, 2H), 2.84-2.62 (m, 3H), 2.13-2.06 (m, 1H).

**^13^C NMR** (126 MHz, CD_2_Cl_2_): δ = 171.6, 168.9, 167.4, 167.3, 167.2, 166.7, 166.5, 159.2, 156.2, 155.1, 149.8, 137.5, 137.5, 135.6, 134.1, 130.1, 129.3, 128.96, 128.1, 121.7, 120.1, 118.5, 118.5, 117.6, 116.3, 70.3, 69.97, 68.6, 68.5, 49.8, 42.9, 39.5, 31.9, 23.1.

HPLC: R_t_ = 4.31 min (method A): Purity: >95% (254 nm); >96% (320 nm)

MS (ESI+):

385.1	[(M/2+H)^+^	calc. 385.12]

769.3	[(M+H)^+^	calc. 769.23]

HRMS (ESI+):

769.2264	[(M+H)^+^	calc. 769.2271]

N-benzyl-2-chloro-N-(4-(4-((1-((2-(2,6-dioxopiperidin-3-yl)-1,3-dioxoisoindolin-4-yl)oxy)-2-oxo-6,9,12-trioxa-3-azatetradecan-14-yl)oxy)phenoxy)phenyl)acetamide **1b**

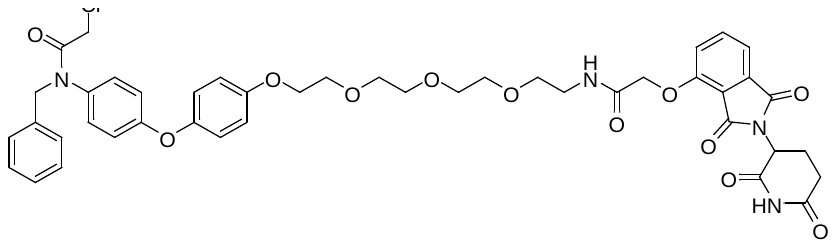

2-chloroacetyl chloride (3 µl, 35 µmol, 1.3eq) and TEA (11 µl, 81 µmol, 3.0eq) were added to a solution of N-(2-(2-(2-(2-(4-(4-(benzylamino)phenoxy)phenoxy)ethoxy)ethoxy)ethoxy)ethyl)-2-((2-(2,6-dioxopiperidin-3-yl)-1,3-dioxoisoindolin-4-yl)oxy)acetamide **S14** (21 mg, 21 µmol, 1.0eq) in DCM (10 mL) at 0 °C. The reaction mixture was stirred for 20 h at r.t. The reaction was quenched with water and the layers were separated. The aqueous layer was extracted with DCM (3x) and the combined organic layers were dried with MgSO_4_. Reversed phase flash column chromatography (ACN/water) yielded the title compound as a colorless oil (16 mg, 69%).

**^1^H NMR** (500 MHz, CD_2_Cl_2_): δ = 8.73 (s, 1H), 7.74 (dd, *^3^J* = 8.3 Hz, *^3^J* = 7.4 Hz, 1H), 7.59 (s, 1H), 7.52 (d, *^3^J* = 7.3 Hz, 1H), 7.31-7.18 (m, 6H), 6.99-6.94 (m, 4H), 6.93-6.89 (m, 2H), 6.88-6.85 (m, 2H), 4.90 (dd, *^3^J* = 12.3 Hz, *^3^J* = 5.4 Hz, 1H), 4.85 (s, 2H), 4.64 (s, 2H), 4.13-4.08 (m, 2H), 3.89 (s, 2H), 3.82 (dd, *^3^J* = 5.4 Hz, *^3^J* = 4.0 Hz, 2H), 3.71-3.59 (m, 10H), 3.59-3.47 (m, 2H), 2.85-2.61 (m, 3H), 2.15-2.07 (m, 1H).

**^13^C NMR** (126 MHz, CD_2_Cl_2_): δ = 171.6, 168.9, 167.4, 167.2, 166.8, 166.5, 159.3, 156.2, 155.1, 149.9, 137.5, 137.5, 135.6, 134.2, 130.2, 129.3, 128.98, 128.1, 121.8, 120.1, 118.6, 118.5, 117.6, 116.3, 71.3, 70.9, 70.8, 70.7, 70.2, 70.1, 68.6, 68.5, 53.98, 49.9, 42.9, 39.6, 31.9, 23.2.

HPLC: R_t_ = 4.36 min (method A): Purity: >99% (254 nm); >99% (320 nm)

MS (ESI+):

429.2	[(M/2+H)^+^	calc. 429.14]

857.3	[(M+H)^+^	calc. 857.28]

HRMS (ESI+):

857.2789	[(M+H)^+^	calc. 857.2795]

N-benzyl-2-chloro-N-(4-(4-((1-((2-(1-methyl-2,6-dioxopiperidin-3-yl)-1,3-dioxoisoindolin-4-yl)oxy)-2-oxo-6,9,12-trioxa-3-azatetradecan-14-yl)oxy)phenoxy)phenyl)acetamide **1b n.c.**

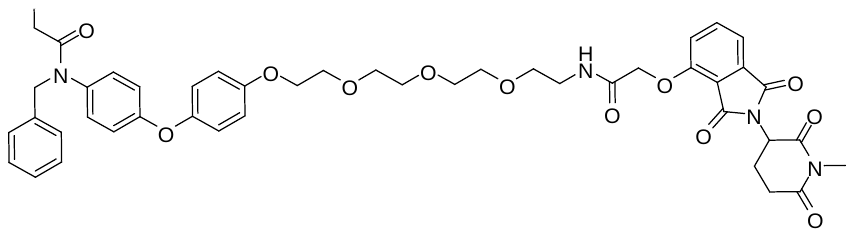

2-chloroacetyl chloride (3 µl, 34 µmol, 1.3eq) and TEA (23 µl, 132 µmol, 5.0eq) were added to a solution of N-(2-(2-(2-(2-(4-(4-(benzylamino)phenoxy)phenoxy)ethoxy)ethoxy)ethoxy)ethyl)-2-((2-(1-methyl-2,6-dioxopiperidin-3-yl)-1,3-dioxoisoindolin-4-yl)oxy)acetamide **S15** (21 mg, 26 µmol, 1.0eq) in DCM (3 mL) at 0 °C. The reaction mixture was stirred for 16 h at r.t. The reaction was quenched with water and the layers were separated. The aqueous layer was extracted with DCM (3x) and the combined organic layers were dried with MgSO_4_. Reversed phase flash column chromatography (ACN/water) yielded the title compound as colorless solid (20 mg, 87%).

**^1^H NMR** (600 MHz, CD_2_Cl_2_): δ = 7.76-7.72 (m, 1H), 7.52 (d, *^3^J* = 7.2 Hz, 1H), 7.48 (t, *^3^J* = 5.1 Hz, 1H), 7.31-7.18 (m, 6H), 6.99-6.93 (m, 4H), 6.93-6.89 (m, 2H), 6.88-6.84 (m, 2H), 4.96 (ddd, *^3^J* = 8.9 Hz, *^3^J* = 5.4 Hz, *^4^J* = 2.3 Hz, 1H), 4.85 (s, 2H), 4.64 (s, 2H), 4.12-4.05 (m, 2H), 3.89 (s, 2H), 3.81-3.77 (m, 2H), 3.66-3.63 (m, 2H), 3.63-3.59 (m, 8H), 3.52 (dd, *^3^J* = 10.8 Hz, *^3^J* = 5.3 Hz, 2H), 2.98-2.89 (m, 1H), 2.83-2.71 (m, 2H), 2.15-2.06 (m, 1H).

**^13^C NMR** (151 MHz, CD_2_Cl_2_): δ = 171.6, 169.4, 167.4, 167.3, 166.7, 166.6, 159.3, 156.3, 155.2, 149.8, 137.5, 137.5, 135.6, 134.3, 130.2, 129.3, 128.99, 128.1, 121.8, 120.2, 118.7, 118.5, 117.6, 116.3, 71.3, 71.1, 71.0, 70.9, 70.2, 70.1, 68.8, 68.6, 54.4, 54.2, 50.7, 42.9, 39.6, 32.5, 27.5, 22.4.

HPLC: R_t_ = 4.51 min (method A): Purity: >99% (254 nm); >97% (320 nm)

MS (ESI+):

436.2	[(M/2+H)^+^	calc. 436.15]

871.3	[(M+H)^+^	calc. 871.30]

HRMS (ESI+):

871.2943	[(M+H)^+^	calc. 871.2952]

tert-butyl (5-(2-((2-(2,6-dioxopiperidin-3-yl)-1,3-dioxoisoindolin-4-yl)oxy)acetamido)pentyl)carbamate **S17**

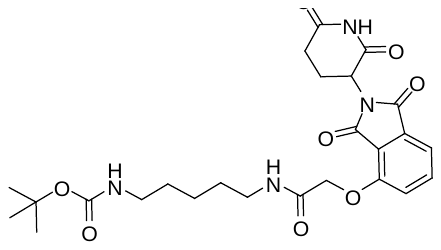

A solution of tert-butyl 2-((2-(2,6-dioxopiperidin-3-yl)-1,3-dioxoisoindolin-4-yl)oxy)acetate **S4** (44 mg, 113 µmol, 1.0eq) in DCM/TFA (4 mL, 1/1) was stirred for 2 h. All volatiles were removed under reduced pressure. DCM was added and the solvent was removed under reduced pressure. This procedure was repeated twice.

A solution of tert-butyl (5-aminopentyl)carbamate (25 mg, 125 µmol, 1.1eq), DIPEA (49 µL, 283 µmol, 2.5eq), the crude acid and HATU (52 mg, 136 µmol, 1.2eq) in DMF (2 mL) was stirred for 17 h. Ethylacetate, water and a sat. solution of NaHCO_3_ were added and the layers were separated. The aqueous layer was extracted with ethylacetate (3x) and the combined organic layers were dried with MgSO_4_. The solvent was evaporated under reduced pressure and the residue was purified by reversed flash column chromatography (ACN /water) to yield the title compound as a colorless solid (48 mg, 82%).

**^1^H NMR** (500 MHz, CD_2_Cl_2_): δ = 9.53-9.09 (m, 1H), 7.75 (dd, *^3^J* = 8.4 Hz, *^3^J* = 7.4 Hz, 1H), 7.52 (d, *^3^J* = 7.0 Hz, 1H), 7.49 (s, 1H), 7.24 (d, *^3^J* = 8.3 Hz, 1H), 5.06-4.97 (m, 1H), 4.83-4.73 (m, 1H), 4.68-4.60 (m, 2H), 3.39 (dd, *^3^J* = 12.4 Hz, *^3^J* = 6.0 Hz, 1H), 3.28 (td, *^3^J* = 12.5 Hz, *^3^J* = 6.4 Hz, 1H), 3.08 (s, 2H), 2.87-2.70 (m, 3H), 2.51 (s, 1H), 2.20-2.09 (m, 1H), 1.63-1.54 (m, 2H), 1.54-1.45 (m, 2H), 1.45-1.34 (m, 11H).

**^13^C NMR** (126 MHz, CD_2_Cl_2_): δ = 172.0, 169.2, 167.3, 167.3, 166.8, 156.6, 155.2, 137.6, 134.1, 120.5, 118.7, 117.7, 79.4, 68.9, 49.9, 41.0, 39.5, 31.95, 30.2, 29.4, 28.7, 24.5, 23.2.

MS (ESI+):

417.1	[(M-Boc+2H)^+^	calc. 417.16]

418.1	[(M-Boc+3H)^2+^	calc. 418.17]

tert-butyl (1-((2-(2,6-dioxopiperidin-3-yl)-1,3-dioxoisoindolin-4-yl)oxy)-2-oxo-6,9,12,15,18-pentaoxa-3-azaicosan-20-yl)carbamate **S17**

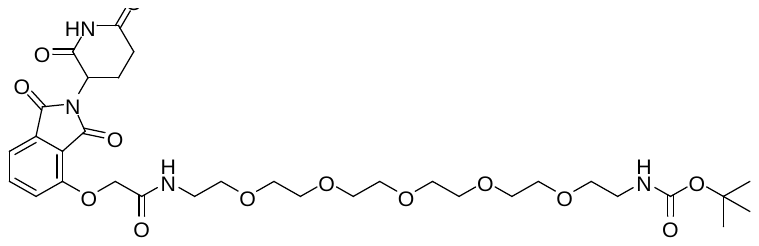

A solution of tert-butyl 2-((2-(2,6-dioxopiperidin-3-yl)-1,3-dioxoisoindolin-4-yl)oxy)acetate **S4** (42 mg, 108 µmol, 1.0eq) in DCM/TFA (4 mL, 1/1) was stirred for 2 h. All volatiles were removed under reduced pressure. DCM was added and the solvent was removed under reduced pressure. This procedure was repeated twice.

A solution of tert-butyl (17-amino-3,6,9,12,15-pentaoxaheptadecyl)carbamate (45 mg, 119 µmol, 1.1eq), DIPEA (47 µL, 270 µmol, 2.5eq), the crude acid and HATU (49 mg, 130 µmol, 1.2eq) in DMF (2 mL) was stirred for 12 h. DCM, water and a sat. solution of NaHCO_3_ were added and the layers were separated. The aqueous layer was extracted with DCM (3x) and the combined organic layers were dried with MgSO_4_. The solvent was evaporated under reduced pressure and the residue was purified by reversed flash column chromatography (ACN /water) to yield the title compound as a yellow oil (56 mg, 75%).

**^1^H NMR** (600 MHz, CD_2_Cl_2_): δ = 8.90 (s, 1H), 7.75 (dd, *^3^J* = 8.4 Hz, *^3^J* = 7.4 Hz, 1H), 7.61 (t, *^3^J* = 5.5 Hz, 1H), 7.53 (dd, *^3^J* = 7.3 Hz, *^4^J* = 0.5 Hz, 1H), 7.24 (dd, *^3^J* = 8.5 Hz, *^3^J* = 0.4 Hz, 1H), 5.19 (s, 1H), 4.97 (dd, *^3^J* = 12.2 Hz, *^3^J* = 5.6 Hz, 1H), 4.65 (s, 2H), 3.64-3.57 (m, 18H), 3.55-3.48 (m, 4H), 3.27-3.23 (m, 2H), 2.88-2.70 (m, 3H), 2.18-2.11 (m, 1H), 1.41 (s, 9H).

**^13^C NMR** (151 MHz, CD_2_Cl_2_): δ = 171.8, 168.97, 167.3, 167.3, 166.5, 156.5, 155.2, 137.5, 134.3, 120.1, 118.6, 117.6, 71.1, 70.99, 70.96, 70.95, 70.90, 70.85, 70.83, 70.76, 70.74, 70.1, 68.7, 49.9, 40.97, 39.6, 32.0, 28.7, 23.2.

MS (ESI+):

595.3	[(M-Boc+2H)^+^	calc. 595.26]

717.3	[(M+Na)^+^	calc. 717.299]

2-(4-(4-(benzylamino)phenoxy)phenoxy)-N-(5-(2-((2-(2,6-dioxopiperidin-3-yl)-1,3-dioxoisoindolin-4-yl)oxy)acetamido)pentyl)acetamide **S18**

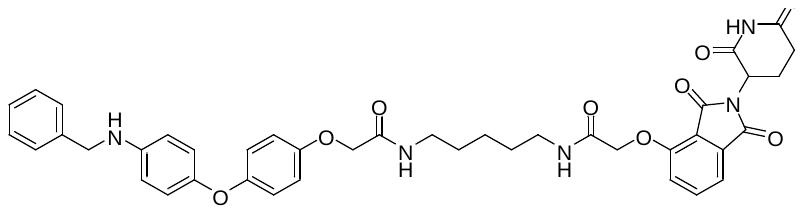

A solution of tert-butyl 2-(4-(4-(benzylamino)phenoxy)phenoxy)acetate **S8** (34 mg, 84 µmol, 1.0eq) in DCM/TFA (4 mL, 1/1) was stirred for 3 h. All volatiles were removed under reduced pressure. DCM was added and the solvent was removed under reduced pressure. This procedure was repeated twice.

A solution of tert-butyl (5-(2-((2-(2,6-dioxopiperidin-3-yl)-1,3-dioxoisoindolin-4-yl)oxy)acetamido)pentyl)carbamate **S16** (48 mg, 93 µmol, 1.1eq) in DCM/TFA (4 mL, 1/1) was stirred for 3 h. All volatiles were removed under reduced pressure. DCM was added and the solvent was removed under reduced pressure. This procedure was repeated twice.

A solution of the crude amine, DIPEA (37 µL, 211 µmol, 2.5eq), the crude acid and HATU (39 mg, 101 µmol, 1.2eq) in DMF (2 mL) was stirred for 18 h. DCM, water and a sat. solution of NaHCO_3_ were added and the layers were separated. The aqueous layer was extracted with DCM (3x) and the combined organic layers were dried with MgSO_4_. The solvent was evaporated under reduced pressure and the residue was purified by reversed flash column chromatography (ACN /water) to yield the title compound as a colorless solid (38 mg, 60%).

**^1^H NMR** (500 MHz, CD_2_Cl_2_): δ = 9.25 (s, 1H), 7.73 (dd, *^3^J* = 8.3 Hz, *^3^J* = 7.4 Hz, 1H), 7.53-7.48 (m, 2H), 7.40-7.25 (m, 5H), 7.21 (d, *^3^J* = 8.3 Hz, 1H), 6.86 (s, 4H), 6.84-6.79 (m, 2H), 6.69 (t, *^3^J* = 5.3 Hz, 1H), 6.63-6.59 (m, 2H), 5.00 (dd, *^3^J* = 12.0 Hz, *^3^J* = 5.8 Hz, 1H), 4.66-4.57 (m, 2H), 4.42 (s, 2H), 4.31 (s, 2H), 3.48-3.20 (m, 4H), 2.86-2.69 (m, 3H), 2.19-2.10 (m, 1H), 1.65-1.52 (m, 4H), 1.50-1.38 (m, 2H).

**^13^C NMR** (126 MHz, CD_2_Cl_2_): δ = 172.0, 169.1, 168.97, 167.2, 167.1, 166.8, 155.2, 154.1, 153.2, 149.1, 145.3, 140.3, 137.6, 134.1, 129.1, 127.99, 127.7, 120.9, 120.5, 119.3, 118.8, 117.7, 116.3, 114.3, 68.95, 68.7, 49.9, 49.2, 39.6, 39.4, 31.95, 29.7, 29.4, 24.7, 23.3.

MS (ESI+):

374.7	[(M/2+H)^+^	calc. 374.65]

748.3	[(M+H)^+^	calc. 748.297]

2-(4-(4-(benzylamino)phenoxy)phenoxy)-N-(1-((2-(2,6-dioxopiperidin-3-yl)-1,3-dioxoisoindolin-4-yl)oxy)-2-oxo-6,9,12,15,18-pentaoxa-3-azaicosan-20-yl)acetamide **S19**

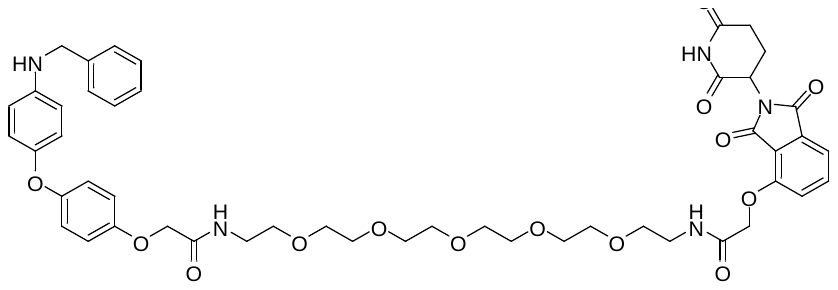

A solution of tert-butyl 2-(4-(4-(benzylamino)phenoxy)phenoxy)acetate **S8** (33 mg, 81 µmol, 1.0eq) in DCM/TFA (4 mL, 1/1) was stirred for 2 h. All volatiles were removed under reduced pressure. DCM was added and the solvent was removed under reduced pressure. This procedure was repeated twice.

A solution of tert-butyl (1-((2-(2,6-dioxopiperidin-3-yl)-1,3-dioxoisoindolin-4-yl)oxy)-2-oxo-6,9,12,15,18-pentaoxa-3-azaicosan-20-yl)carbamate **S17** (56 mg, 81 µmol, 1.0eq) in DCM/TFA (4 mL, 1/1) was stirred for 2 h. All volatiles were removed under reduced pressure. DCM was added and the solvent was removed under reduced pressure. This procedure was repeated twice.

A solution of the crude amine, DIPEA (35 µL, 202 µmol, 2.5eq), the crude acid and HATU (37 mg, 97 µmol, 1.2eq) in DMF (2 mL) was stirred for 12 h. DCM, water and a sat. solution of NaHCO_3_ were added and the layers were separated. The aqueous layer was extracted with DCM (3x) and the combined organic layers were dried with MgSO_4_. The solvent was evaporated under reduced pressure and the residue was purified by reversed flash column chromatography (ACN /water) to yield the title compound as a yellow oil (41 mg, 55%).

**^1^H NMR** (400 MHz, CD_2_Cl_2_): δ = 9.10 (s, 1H), 7.72 (dd, *^3^J* = 8.4 Hz, *^3^J* = 7.4 Hz, 1H), 7.55 (t, *^3^J* = 5.1 Hz, 1H), 7.50 (dd, *^3^J* = 7.3 Hz, *^4^J* = 0.5 Hz, 1H), 7.40-7.31 (m, 4H), 7.30-7.24 (m, 1H), 7.20 (d, *^3^J* = 8.4 Hz, 1H), 7.07 (t, *^3^J* = 5.4 Hz, 1H), 6.86 (s, 4H), 6.84-6.79 (m, 2H), 6.63-6.58 (m, 2H), 5.00-4.92 (m, 1H), 4.62 (s, 2H), 4.41 (s, 2H), 4.30 (s, 2H), 4.18 (s, 1H), 3.64-3.46 (m, 24H), 2.87-2.70 (m, 3H), 2.20-2.10 (m, 1H).

**^13^C NMR** (101 MHz, CD_2_Cl_2_): δ = 171.9, 169.1, 168.7, 167.3, 167.2, 166.4, 155.0, 153.9, 153.3, 149.1, 145.2, 140.2, 137.4, 134.2, 129.1, 127.96, 127.7, 120.8, 119.99, 119.2, 118.5, 117.4, 116.3, 114.2, 71.0, 70.96, 70.9, 70.9, 70.8, 70.2, 69.98, 68.6, 68.5, 49.9, 49.1, 39.6, 39.3, 31.98, 23.1.

MS (ESI+):

463.8	[(M/2+H)^+^	calc. 463.69]

N-benzyl-2-chloro-N-(4-(4-(2-((5-(2-((2-(2,6-dioxopiperidin-3-yl)-1,3-dioxoisoindolin-4-yl)oxy)acetamido)pentyl)amino)-2-oxoethoxy)phenoxy)phenyl)acetamide **1c**

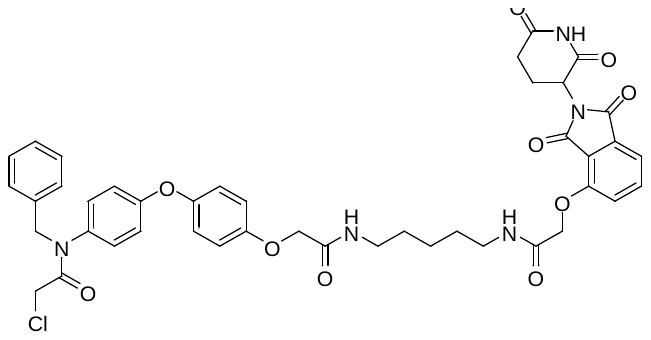

2-chloroacetyl chloride (5 µl, 66 µmol, 1.3eq) and TEA (21 µl, 152 µmol, 5.0eq) were added to a solution of 2-(4-(4-(benzylamino)phenoxy)phenoxy)-N-(5-(2-((2-(2,6-dioxopiperidin-3-yl)-1,3-dioxoisoindolin-4-yl)oxy)acetamido)pentyl)acetamide **S18** (38 mg, 51 µmol, 1.0eq) in DCM (3 mL) at 0 °C. The reaction mixture was stirred for 17 h at r.t. The reaction was quenched with water and the layers were separated. The aqueous layer was extracted with DCM (3x) and the combined organic layers were dried with MgSO_4_. Reversed phase flash column chromatography (ACN/water) yielded the title compound as colorless solid (31 mg, 74%).

**^1^H NMR** (500 MHz, CD_2_Cl_2_): δ = 9.29 (s, 1H), 7.74 (dd, *^3^J* = 8.3 Hz, *^3^J* = 7.4 Hz, 1H), 7.55-7.48 (m, 2H), 7.32-7.18 (m, 6H), 7.01-6.92 (m, 6H), 6.89-6.85 (m, 2H), 6.70 (t, *^3^J* = 5.5 Hz, 1H), 5.03-4.98 (m, 1H), 4.85 (s, 2H), 4.66-4.58 (m, 2H), 4.46 (s, 2H), 3.89 (s, 2H), 3.47-3.22 (m, 4H), 2.88-2.70 (m, 3H), 2.21-2.10 (m, 1H), 1.66-1.52 (m, 4H), 1.49-1.39 (m, 2H).

**^13^C NMR** (126 MHz, CD_2_Cl_2_): δ = 172.0, 169.2, 168.7, 167.2, 167.1, 166.8, 166.7, 158.9, 155.2, 154.7, 150.8, 137.6, 137.4, 135.8, 134.1, 130.2, 129.3, 128.97, 128.1, 121.9, 120.5, 118.8, 118.7, 117.7, 116.6, 68.97, 68.5, 54.2, 49.9, 42.9, 39.6, 39.4, 31.95, 29.7, 29.4, 24.7, 23.3.

HPLC: R_t_ = 4.20 min (method A): Purity: >97% (254 nm); >99% (320 nm)

MS (ESI+):

412.5	[(M/2+H)^+^	calc. 412.60]

824.3	[(M+H)^+^	calc. 824.27]

HRMS (ESI+):

824.2685	[(M+H)^+^	calc. 824.2693]

N-benzyl-2-chloro-N-(4-(4-((23-((2-(2,6-dioxopiperidin-3-yl)-1,3-dioxoisoindolin-4-yl)oxy)-2,22-dioxo-6,9,12,15,18-pentaoxa-3,21-diazatricosyl)oxy)phenoxy)phenyl)acetamide **1d**

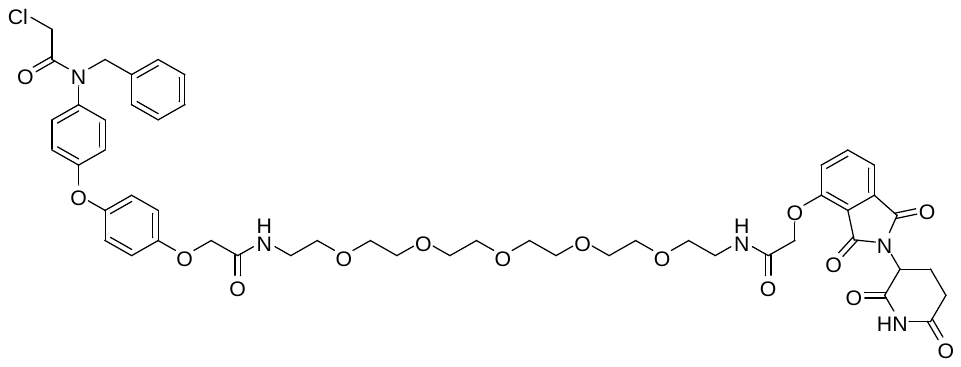

2-chloroacetyl chloride (5 µl, 69 µmol, 1.3eq) and TEA (22 µl, 158 µmol, 3.0eq) were added to a solution of 2-(4-(4-(benzylamino)phenoxy)phenoxy)-N-(1-((2-(2,6-dioxopiperidin-3-yl)-1,3-dioxoisoindolin-4-yl)oxy)-2-oxo-6,9,12,15,18-pentaoxa-3-azaicosan-20-yl)acetamide **S19** (49 mg, 53 µmol, 1.0eq) in DCM (10 mL) at 0 °C. The reaction mixture was stirred for 20 h at r.t. The reaction was quenched with water and the layers were separated. The aqueous layer was extracted with DCM (3x) and the combined organic layers were dried with MgSO_4_. Reversed phase flash column chromatography (ACN/water) yielded the title compound as a yellow oil (31 mg, 59%).

**^1^H NMR** (500 MHz, DMSO*-d_6_*): δ = 11.11 (s, 1H), 8.06 (t, *^3^J* = 5.7 Hz, 1H), 7.99 (t, *^3^J* = 5.6 Hz, 1H), 7.80 (dd, *^3^J* = 8.5 Hz, *^3^J* = 7.3 Hz, 1H), 7.49 (d, *^3^J* = 7.2 Hz, 1H), 7.40 (d, *^3^J* = 8.5 Hz, 1H), 7.29 (t, *^3^J* = 7.2 Hz, 2H), 7.26-7.17 (m, 5H), 7.04-6.97 (m, 4H), 6.91-6.86 (m, 2H), 5.11 (dd, *^3^J* = 12.7 Hz, *^3^J* = 5.5 Hz, 1H), 4.84 (s, 2H), 4.78 (s, 2H), 4.46 (s, 2H), 4.08 (s, 2H), 3.53-3.42 (m, 20H), 3.36-3.26 (m, 4H), 2.94-2.85 (m, 1H), 2.64-2.51 (m, 2H), 2.06-2.00 (m, 1H).

**^13^C NMR** (126 MHz, DMSO*-d_6_*): δ = 172.8, 169.9, 167.7, 166.9, 166.7, 165.8, 165.5, 157.6, 154.98, 154.4, 149.3, 136.9, 136.9, 135.1, 133.0, 129.7, 128.4, 127.97, 127.3, 120.98, 120.4, 117.8, 116.8, 116.2, 116.1, 69.78, 69.77, 69.7, 69.6, 69.6, 68.8, 67.5, 67.4, 52.8, 48.8, 42.5, 39.5, 38.4, 38.3, 30.95, 22.0.

HPLC: R_t_ = 4.07 min (method B): Purity: >99% (254 nm); >98% (320 nm)

MS (ESI+):

501.7	[(M/2+H)^+^	calc. 501.68]

1002.3	[(M+H)^+^	calc. 1002.35]

HRMS (ESI+):

1002.3536	[(M+H)^+^	calc. 1002.3534]

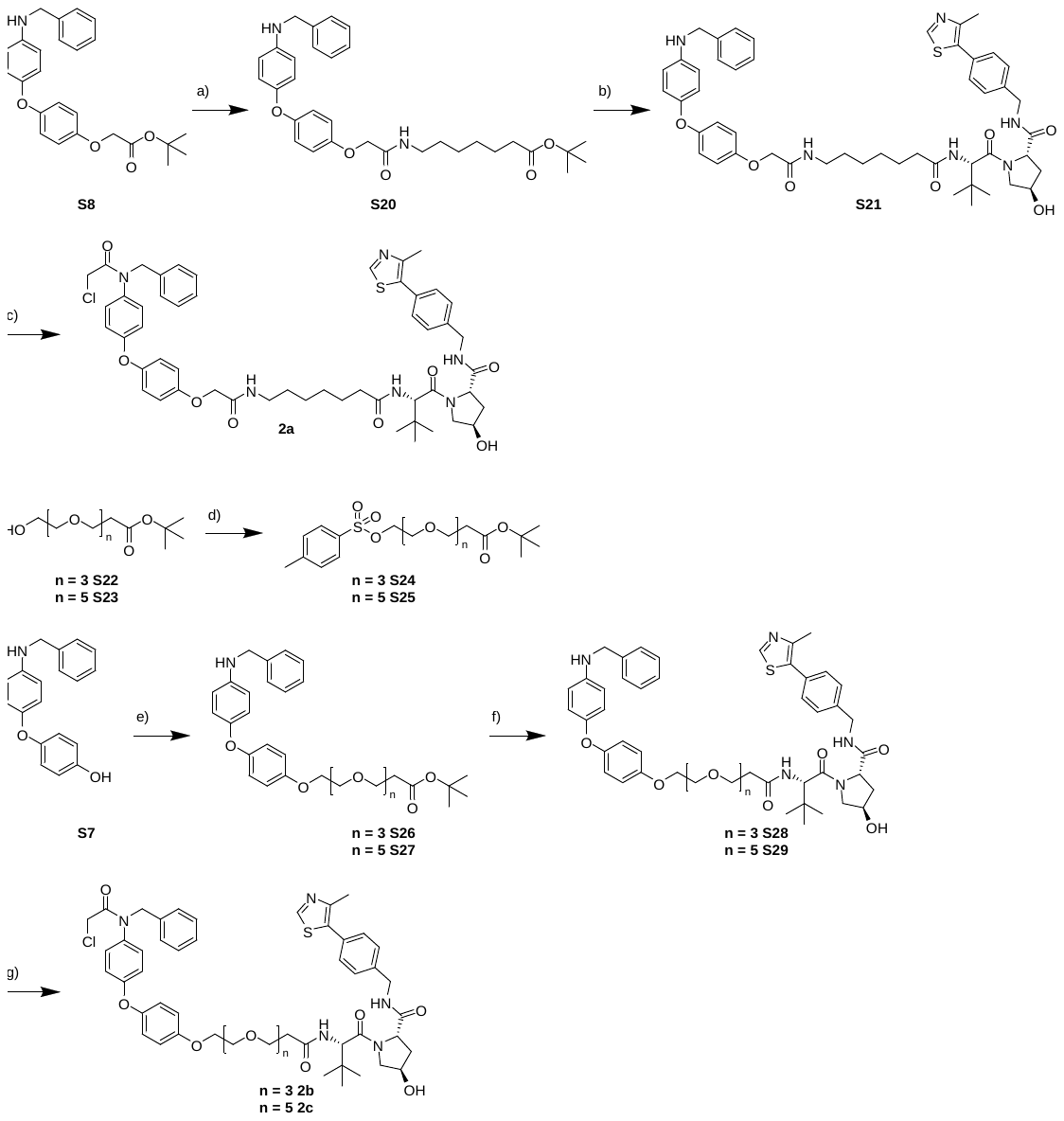

**Figure S##:** Synthesis of VHL based PROTACs **2a-c**: a) 1. TFA, DCM, r.t., 2.5 h; 2. DIPEA, linker-amine, HATU, DMF, r.t., 24.5 h; b) 1. TFA, DCM, r.t., 2.5 h; 2. DIPEA, VHL032-amine, HATU, DMF, r.t., 24.5 h; c) 2-chloroacetyl chloride, TEA, 0 °C, then r.t., 20 h; d) TsCl, 4-DMAP, TEA, DCM, -10 °C, then r.t., 24-38 h; e) intermediate **S22** or **S23**, K_2_CO_3_, DMF, 90 °C, overnight; f) 1. TFA, DCM, r.t., 4.5 h; 2. DIPEA, VHL032-amine, HATU, DMF, r.t., 17 h; g) 2-chloroacetyl chloride, TEA, 0 °C, then r.t., 22 h;

tert-butyl 7-(2-(4-(4-(benzylamino)phenoxy)phenoxy)acetamido)heptanoate **S20**

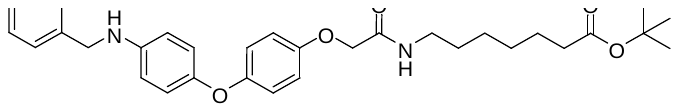

A solution of tert-butyl 2-(4-(4-(benzylamino)phenoxy)phenoxy)acetate **S8** (59 mg, 146 µmol, 1.0eq) in DCM/TFA (4 mL, 1/1) was stirred for 2.5 h. All volatiles were removed under reduced pressure. DCM was added and the solvent was removed under reduced pressure. This procedure was repeated twice.

DIPEA (51 µL, 291 µmol, 2.0eq) and HATU (66 mg, 175 µmol, 1.2eq) were added to a solution of *tert*-butyl 7-aminoheptanoate (35 mg, 175 µmol, 1.2eq) and the crude acid in DMF (1 mL) The reaction mixture was stirred for 24.5 h. Ethylacetate, water and brine were added and the layers were separated. The aqueous layer was extracted with ethylacetate (3x) and the combined organic layers were dried with MgSO_4_. The solvent was removed under reduced pressure and the residue was purified by reversed phase flash column chromatography to yield the title compound as a yellow oil (25 mg, 32%).

**^1^H NMR** (500 MHz, CD_2_Cl_2_): δ = 7.42-7.33 (m, 4H), 7.31-7.25 (m, 1H), 6.90 – 6.85 (m, 4H), 6.84-6.81 (m, 2H), 6.63-6.59 (m, 2H), 6.59-6.56 (m, 1H), 4.40 (s, 2H), 4.31 (s, 2H), 3.29 (dd, *^3^J* = 13.5 Hz, *^3^J* = 6.8 Hz, 2H), 2.18 (t, *^3^J* = 7.5 Hz, 2H), 1.60-1.49 (m, 4H), 1.42 (s, 9H), 1.36-1.29 (m, 4H).

**^13^C NMR** (126 MHz, CD_2_Cl_2_): δ = 173.4, 168.4, 154.1, 153.2, 149.1, 145.3, 140.3, 129.1, 128.0, 127.7, 120.9, 119.3, 116.2, 114.3, 80.2, 68.7, 49.2, 39.4, 35.96, 30.0, 29.3, 28.4, 27.1, 25.5.

MS (ESI+):

477.1	[(M-^t^Bu+2H)^+^	calc. 477.23]

533.3	[(M+H)^+^	calc. 533.297]

(2S,4R)-1-((S)-2-(7-(2-(4-(4-(benzylamino)phenoxy)phenoxy)acetamido)heptanamido)-3,3-dimethylbutanoyl)-4-hydroxy-N-(4-(4-methylthiazol-5-yl)benzyl)pyrrolidine-2-carboxamide **S21**

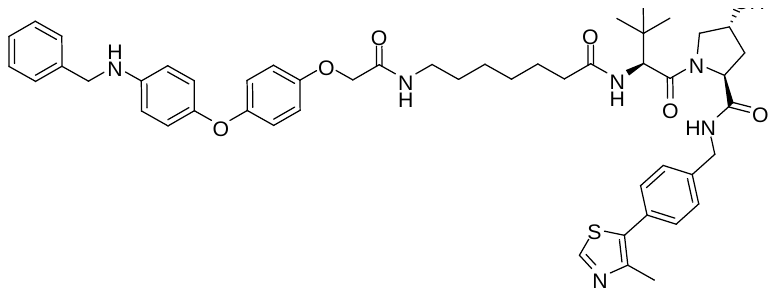

A solution of tert-butyl 7-(2-(4-(4-(benzylamino)phenoxy)phenoxy)acetamido)heptanoate **S20** (25 mg, 471 µmol, 1.0eq) in DCM/TFA (4 mL, 1/1) was stirred for 2 h. All volatiles were removed under reduced pressure. DCM was added and the solvent was removed under reduced pressure. This procedure was repeated twice.

DIPEA (16 µL, 94 µmol, 2.0eq) and HATU (21 mg, 56 µmol, 1.2eq) were added to a solution of the (2S,4R)-1-((S)-2-amino-3,3-dimethylbutanoyl)-4-hydroxy-N-(4-(4-methylthiazol-5-yl)benzyl)pyrrolidine-2-carboxamide hydrochloride (24 mg, 52 µmol, 1.1eq) and the crude acid in DMF (1.0 mL). The reaction mixture was stirred for 15 h. Ethylacetate, water and brine were added and the layers were separated. The aqueous layer was extracted with ethylacetate (3x) and the combined organic layers were dried with MgSO_4_. The solvent was evaporated under reduced pressure and the residue was purified by reverse flash column chromatography to yield the title compound as yellow oil (13 mg, 31%).

**^1^H NMR** (500 MHz, CD_2_Cl_2_): δ = 8.65 (s, 1H), 7.42-7.24 (m, 10H), 6.89-6.79 (m, 6H), 6.66-6.59 (m, 3H), 6.24 (d, *^3^J* = 8.8 Hz, 1H), 4.67 (t, *^3^J* = 8.1 Hz, 1H), 4.57-4.50 (m, 2H), 4.48 (s, 1H), 4.37 (s, 2H), 4.33-4.27 (m, 3H), 4.04 (d, *^3^J* = 11.4 Hz, 1H), 3.60 (dd, *^3^J* = 11.3 Hz, *^3^J* = 3.6 Hz, 1H), 3.27 (td, *^3^J* = 7.4 Hz, *^4^J* = 1.3 Hz, 2H), 2.48 (s, 3H), 2.39 (ddd, *^2^J* = 13.0 Hz, *^3^J* = 8.2 Hz, *^3^J* = 4.5 Hz, 1H), 2.24-2.07 (m, 3H), 1.64-1.46 (m, 4H), 1.35-1.21 (m, 5H), 0.94 (s, 9H).

**^13^C NMR** (126 MHz, CD_2_Cl_2_): δ = 174.1, 172.3, 171.5, 168.7, 154.1, 153.2, 150.7, 149.1, 149.1, 145.3, 140.3, 139.1, 132.1, 131.5, 129.9, 129.1, 128.5, 128.0, 127.7, 120.9, 119.3, 116.2, 114.3, 70.7, 68.6, 59.1, 57.97, 57.4, 49.2, 43.5, 39.3, 36.95, 36.7, 35.4, 29.8, 28.9, 26.8, 26.7, 25.95, 16.5.

MS (ESI+):

318.0	[(fragment+2H)^+^	calc. 318.12]

445.3	[(M/2+H)^+^	calc. 445.22]

572.3	[(fragment)^+^	calc. 572.31]

889.4	[(M+H)^+^	calc. 889.43]

(2S,4R)-1-((S)-2-(7-(2-(4-(4-(N-benzyl-2-chloroacetamido)phenoxy)phenoxy)acetamido)heptanamido)-3,3-dimethylbutanoyl)-4-hydroxy-N-(4-(4-methylthiazol-5-yl)benzyl)pyrrolidine-2-carboxamide **2a**

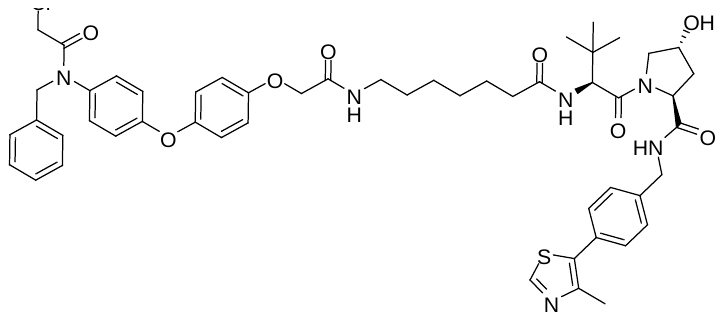

2-chloroacetyl chloride (1 µl, 15 µmol, 1.3eq) and TEA (5 µl, 34 µmol, 3.0eq) were added to a solution of (2S,4R)-1-((S)-2-(7-(2-(4-(4-(benzylamino)phenoxy)phenoxy)acetamido)heptanamido)-3,3-dimethylbutanoyl)-4-hydroxy-N-(4-(4-methylthiazol-5-yl)benzyl)pyrrolidine-2-carboxamide **S21** (10 mg, 11 µmol, 1.0eq) in DCM (5 mL) at 0 °C. The reaction mixture was stirred for 20 h at r.t. The reaction was quenched with water and the layers were separated. The aqueous layer was extracted with DCM (3x) and the combined organic layers were dried with Na_2_SO_4_. Reverse phase flash column chromatography yielded the title compound as beige solid (8 mg, 74%).

**^1^H NMR** (500 MHz, CD_2_Cl_2_): δ = 8.65 (s, 1H), 7.42-7.32 (m, 5H), 7.32-7.23 (m, 3H), 7.21-7.18 (m, 2H), 7.03-6.95 (m, 4H), 6.95-6.91 (m, 2H), 6.89-6.84 (m, 2H), 6.62 (t, *^3^J* = 5.7 Hz, 1H), 6.19 (d, *^3^J* = 8.7 Hz, 1H), 4.85 (s, 2H), 4.67 (t, *^3^J* = 8.1 Hz, 1H), 4.58-4.50 (m, 2H), 4.49 (s, 1H), 4.42 (s, 2H), 4.31 (dd, *^2^J* = 15.1 Hz, *^3^J* = 5.3 Hz, 1H), 4.06 (d, *^3^J* = 11.4 Hz, 1H), 3.90 (s, 2H), 3.59 (dd, *^3^J* = 11.3 Hz, *^3^J* = 3.6 Hz, 1H), 3.28 (dd, *^2^J* = 13.3 Hz, *^3^J* = 7.1 Hz, 2H), 2.48 (s, 3H), 2.42 (ddd, *^2^J* = 12.9 Hz, *^3^J* = 8.1 Hz, *^4^J* = 4.5 Hz, 1H), 2.24-2.08 (m, 3H), 1.63-1.47 (m, 4H), 1.34-1.22 (m, 5H), 0.93 (s, 9H).

**^13^C NMR** (126 MHz, CD_2_Cl_2_): δ = 174.0, 172.4, 171.4, 168.4, 166.7, 158.99, 154.7, 150.8, 150.7, 149.2, 139.1, 137.5, 135.8, 132.1, 131.5, 130.2, 129.9, 129.3, 129.0, 128.5, 128.1, 121.9, 118.7, 116.6, 70.7, 68.5, 59.1, 57.95, 57.3, 54.02, 43.6, 42.9, 39.3, 36.8, 36.7, 35.3, 29.8, 28.9, 26.8, 26.7, 25.9, 16.5.

HPLC: R_t_ = 4.53 min (method B): Purity: >98% (254 nm); >96% (320 nm)

MS (ESI+):

318.1	[(fragment+2H)^+^	calc. 318.12]

648.3	[(fragment)^+^	calc. 648.28]

965.4	[(M+H)^+^	calc. 965.40]

HRMS (ESI+):

965.4031	[(M+H)^+^	calc. 965.4033]

tert-butyl 3-(2-(2-(2-(4-(4-(benzylamino)phenoxy)phenoxy)ethoxy)ethoxy)ethoxy)propanoate **S26** via *tert*-butyl 3-(2-(2-(2-(tosyloxy)ethoxy)ethoxy)ethoxy)propanoate **S24**

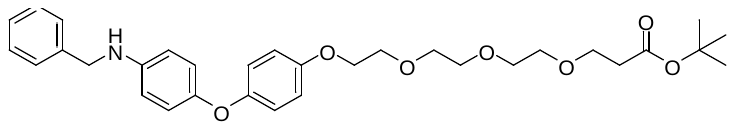

TsCl (81 mg, 0.43 mmol, 1.4eq) was added portion wise (3x) to a solution of the tert-butyl 3-(2-(2-(2-hydroxyethoxy)ethoxy)ethoxy)propanoate **S22** (84 mg, 0.30 mmol, 1.0eq), 4-DMAP (9 mg, 0.07 μmol, 0.2eq) and Et_3_N (0.055 mL, 0.40 mmol, 1.3eq) in DCM (5 mL) at -10 °C. The reaction mixture was allowed to warm to room temperature and stirred for 37.5 h. The reaction was quenched by adding 4 mL of a saturated solution of NH_4_Cl in water. The organic layer was separated and the remaining aqueous layer was extracted with DCM (3x). The combined organic layers were dried with Na_2_SO_4_ and the solvent was removed under reduced pressure. Purification by column chromatography (DCM/MeOH 99/1) yielded the intermediate product (105 mg, 80%) as a colorless oil.

A solution of the 4-(4-(benzylamino)phenoxy)phenol **S7** (83 mg, 283 µmol, 1.0eq), the tosylate **S24** (147 mg, 340 µmol, 1.2eq) and potassium carbonate (118 mg, 850 µmol, 3.0eq) in DMF (4 mL) was stirred at 90 °C overnight. The reaction was quenched with water and extracted with ethylacetate (3x). The combined organic layers were dried with MgSO_4_ and the solvent was removed under reduced pressure. Reversed phase column chromatography (ACN/water) yielded the title compound as a colorless oil (78 mg, 50%).

**^1^H NMR** (500 MHz, CD_2_Cl_2_): δ = 7.42-7.21 (m, 5H), 6.91-6.81 (m, 6H), 6.64-6.59 (m, 2H), 4.31 (s, 2H), 4.07 (dd, *^3^J* = 5.4 Hz, *^3^J* = 4.1 Hz, 2H), 3.80 (dd, *^3^J* = 5.4 Hz, *^3^J* = 4.1 Hz, 2H), 3.70-3.65 (m, 4H), 3.64-3.57 (m, 6H), 2.47 (t, *^3^J* = 6.5 Hz, 2H), 1.45 (s, 9H).

**^13^C NMR** (126 MHz, CD_2_Cl_2_): δ = 171.3, 154.8, 153.0, 149.6, 144.99, 140.3, 129.1, 128.0, 127.7, 120.6, 119.4, 115.98, 114.3, 80.8, 71.3, 71.1, 71.0, 70.9, 70.3, 68.6, 67.4, 49.2, 36.9, 28.4.

MS (ESI+):

496.1	[(M-^t^Bu+2H)^+^	calc. 496.23]

552.2	[(M+H)^+^	calc. 552.298]

553.3	[(M+2H)^2+^	calc. 553.30]

574.33	[(M+Na)^+^	calc. 574.279]

tert-butyl 1-(4-(4-(benzylamino)phenoxy)phenoxy)-3,6,9,12,15-pentaoxaoctadecan-18-oate **S27** via tert-butyl 1-(tosyloxy)-3,6,9,12,15-pentaoxaoctadecan-18-oate **S25**

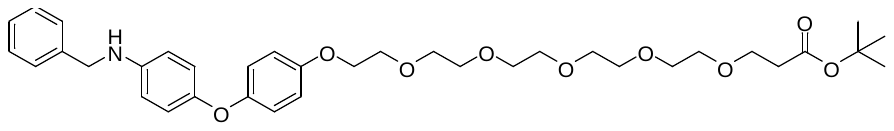

TsCl (80 mg, 0.42 mmol, 1.4eq) was added to a solution of the tert-butyl 1-hydroxy-3,6,9,12,15-pentaoxaoctadecan-18-oate **S23** (107 mg, 0.292 mmol, 1.0eq), 4-DMAP (9 mg, 0.07 mmol, 0.2eq) and Et_3_N (0.050 mL, 0.36 mmol, 1.2eq) in DCM (5 mL) at -10 °C. The reaction mixture was stirred for 30 min, allowed to warm to room temperature and stirred for another 24 h. The reaction was quenched by adding a saturated solution of NH_4_Cl in water. The organic layer was separated and the remaining aqueous layer was extracted with DCM (3x). The combined organic layers were washed with brine, dried with Na_2_SO_4_ and the solvent was removed under reduced pressure. Purification by column chromatography (DCM/MeOH 99/1) yielded the intermediate product (117 mg, 77%) as a colorless oil.

A solution of the 4-(4-(benzylamino)phenoxy)phenol **S7** (54 mg, 186 µmol, 1.0eq), the tosylate **S25** (116 mg, 223  µmol, 1.2eq) and potassium carbonate (78 mg, 557  µmol, 3.0eq) in DMF (3 mL) was stirred at 90 °C overnight. The reaction was quenched with water and extracted with ethylacetate (3x). The combined organic layers were dried with MgSO_4_ and the solvent was removed under reduced pressure. Reversed phase column chromatography (ACN/water) yielded the title compound as a colorless oil (77 mg, 65%).

**^1^H NMR** (500 MHz, CD_2_Cl_2_): δ = 7.40 – 7.25 (m, 5H), 6.89-6.79 (m, 6H), 6.63-6.58 (m, 2H), 4.31 (s, 2H), 4.10-3.99 (m, 2H), 3.80 (dd, *^3^J* = 11.1 Hz, *^3^J* = 6.3 Hz, 2H), 3.70-3.64 (m, 4H), 3.65-3.55 (m, 14H), 2.46 (t, *^3^J* = 6.5 Hz, 2H), 1.44 (s, 9H).

**^13^C NMR** (126 MHz, CD_2_Cl_2_): δ = 171.4, 154.9, 153.1, 149.7, 145.1, 140.4, 129.2, 128.1, 127.8, 120.7, 119.4, 116.1, 114.4, 80.9, 71.4, 71.21, 71.18, 71.17, 71.1, 70.99, 70.4, 68.7, 67.5, 49.3, 36.95, 28.5.

MS (ESI+):

331.7	[(M/2+H)^+^	calc. 331.69]

640.3	[(M+H)^+^	calc. 640.35]

662.3	[(M+Na)^+^	calc. 662.33]

(2S,4R)-1-((S)-1-(4-(4-(benzylamino)phenoxy)phenoxy)-14-(tert-butyl)-12-oxo-3,6,9-trioxa-13-azapentadecan-15-oyl)-4-hydroxy-N-(4-(4-methylthiazol-5-yl)benzyl)pyrrolidine-2-carboxamide **S28**

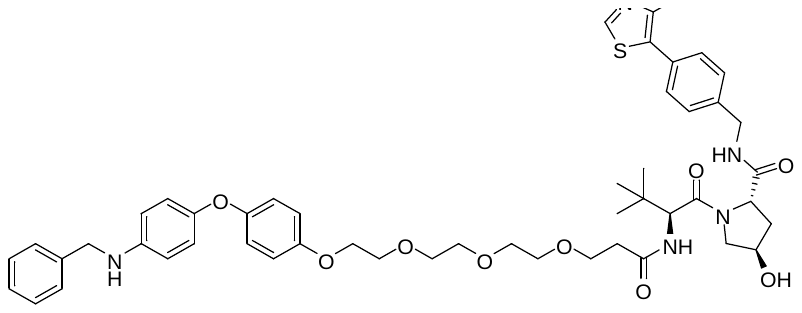

A solution of tert-butyl 3-(2-(2-(2-(4-(4-(benzylamino)phenoxy)phenoxy)ethoxy)ethoxy)ethoxy)propanoate **S26** (26 mg, 47 µmol, 1.0eq) in DCM/TFA (4 mL, 1/1) was stirred for 4.5 h. All volatiles were removed under reduced pressure. DCM was added and the solvent was removed under reduced pressure. This procedure was repeated once.

DIPEA (16 µL, 52 µmol, 2.0eq) and HATU (22 mg, 57 µmol, 1.2eq) were added to a solution of (2S,4R)-1-((S)-2-amino-3,3-dimethylbutanoyl)-4-hydroxy-N-(4-(4-methylthiazol-5-yl)benzyl)pyrrolidine-2-carboxamide hydrochloride (24 mg, 52 µmol, 1.1eq) and the crude acid in DMF (1.5 mL). The reaction mixture was stirred for 17 h. Ethylacetate, water and brine were added and the layers were separated. The aqueous layer was extracted with ethylacetate (3x) and the combined organic layers were dried with MgSO_4_. The solvent was evaporated under reduced pressure and the residue was purified by reverse flash column chromatography to yield the title compound as a yellow oil (26 mg, 61%).

**^1^H NMR** (500 MHz, CD_2_Cl_2_): δ = 8.65 (s, 1H), 7.40-7.31 (m, 9H), 7.29-7.24 (m, 1H), 6.97 (d, *^3^J* = 8.15 Hz, 1H), 6.88-6.79 (m, 6H), 6.62-6.58 (m, 2H), 4.66 (t, *^3^J* = 7.91 Hz, 1H), 4.53 (dd,*^2^J* = 15.05 Hz, *^3^J* = 6.57 Hz, 1H), 4.47-4.43 (m, 2H), 4.32-4.26 (m, 3H), 4.08-4.00 (m, 3H), 3.81-3.55 (m, 13H), 2.48 (s, 3H), 2.47-2.38 (m, 3H), 2.12-2.04 (m, 1H), 0.94 (s, 9H).

**^13^C NMR** (126 MHz, CD_2_Cl_2_): δ = 172.5, 172.3, 171.4, 154.7, 153.1, 150.7, 149.6, 149.1, 145.1, 140.3, 139.1, 132.1, 131.4, 129.9, 129.1, 128.5, 128.0, 127.7, 120.6, 119.4, 115.99, 114.3, 71.2, 71.1, 71.0, 70.96, 70.7, 70.3, 68.6, 67.7, 59.0, 58.3, 57.2, 49.2, 43.5, 37.2, 36.8, 35.3, 26.8, 16.5.

MS (ESI+):

318.1	[(fragment+2H)^+^	calc. 318.12]

454.9	[(M/2+H)^+^	calc. 454.72]

591.3	[(fragment)^+^	calc. 591.31]

908.5	[(M+H)^+^	calc. 908.43]

(2S,4R)-1-((S)-1-(4-(4-(benzylamino)phenoxy)phenoxy)-20-(tert-butyl)-18-oxo-3,6,9,12,15-pentaoxa-19-azahenicosan-21-oyl)-4-hydroxy-N-(4-(4-methylthiazol-5-yl)benzyl)pyrrolidine-2-carboxamide **S29**

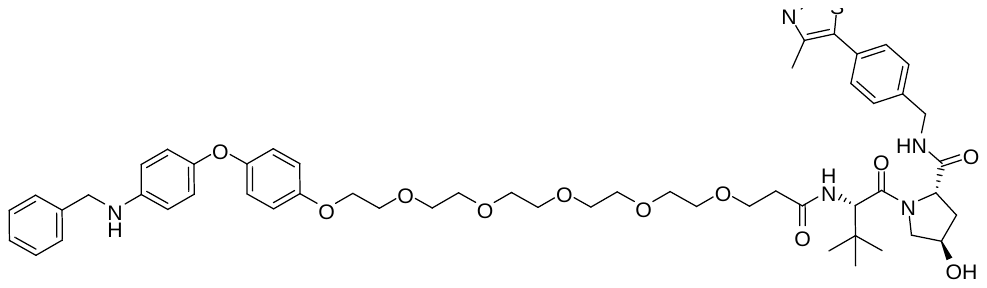

A solution of tert-butyl 1-(4-(4-(benzylamino)phenoxy)phenoxy)-3,6,9,12,15-pentaoxaoctadecan-18-oate **S27** (31 mg, 48 µmol, 1.0eq) in DCM/TFA (4 mL, 1/1) was stirred for 4.5 h. All volatiles were removed under reduced pressure. DCM was added and the solvent was removed under reduced pressure. This procedure was repeated once.

DIPEA (17 µL, 97 µmol, 2.0eq) and HATU (22 mg, 58 µmol, 1.2eq) were added to a solution of (2S,4R)-1-((S)-2-amino-3,3-dimethylbutanoyl)-4-hydroxy-N-(4-(4-methylthiazol-5-yl)benzyl)pyrrolidine-2-carboxamide hydrochloride (25 mg, 53 µmol, 1.1eq) and the crude acid in DMF (1.5 mL) The reaction mixture was stirred for 17 h. Ethylacetate, water and brine were added and the layers were separated. The aqueous layer was extracted with ethylacetate (3x) and the combined organic layers were dried with MgSO_4_. The solvent was removed under reduced pressure and the crude material was purified using reversed phase flash column chromatography. The title compound was isolated as a yellow oil (28 mg, 58%).

**^1^H NMR** (500 MHz, CD_2_Cl_2_): δ = 8.65 (s, 1H), 7.41-7.32 (m, 8H), 7.28-7.24 (m, 1H), 6.96 (d, *^3^J* = 8.39 Hz), 6.88-6.79 (m, 5H), 6.62-6.59 (m, 2H), 4.67 (t, *^3^J* = 8.08 Hz, 1H), 4.54 (dd, *^2^J* = 15.16 Hz, *^3^J* = 6.71 Hz, 1H), 4.48-4.43 (m, 2H), 4.33-4.27 (m, 3H), 4.12 (s, 1H), 4.08-4.01 (m, 3H), 3.81-3.77 (m, 2H), 3.72-3.55 (m, 19H), 3.40 (s, 1H), 2.49 (s, 3H), 2.47-2.38 (m, 3H). 2.12-2.04 (m, 1H), 0.94 (s, 9H).

**^13^C NMR** (126 MHz, CD_2_Cl_2_): δ = 172.5, 172.3, 171.4, 154.8, 153.0, 150.7, 149.6, 149.1, 145.1, 140.3, 139.1, 132.1, 131.4, 129.9, 129.1, 128.5, 128.0, 127.7, 120.6, 119.4, 115.99, 114.3, 71.3, 71.08, 71.07, 71.05, 71.03, 71.00, 70.9, 70.7, 70.3, 68.6, 67.7, 59.0, 58.3, 57.2, 49.2, 43.5, 37.3, 36.7, 35.3, 26.8, 16.5.

MS (ESI+):

498.8	[(M/2+H)^+^	calc. 498.74]

679.3	[(fragment)^+^	calc. 679.36]

996.5	[(M+H)^+^	calc. 996.50]

(2S,4R)-1-((S)-1-(4-(4-(N-benzyl-2-chloroacetamido)phenoxy)phenoxy)-14-(tert-butyl)-12-oxo-3,6,9-trioxa-13-azapentadecan-15-oyl)-4-hydroxy-N-(4-(4-methylthiazol-5-yl)benzyl)pyrrolidine-2-carboxamide **2b**

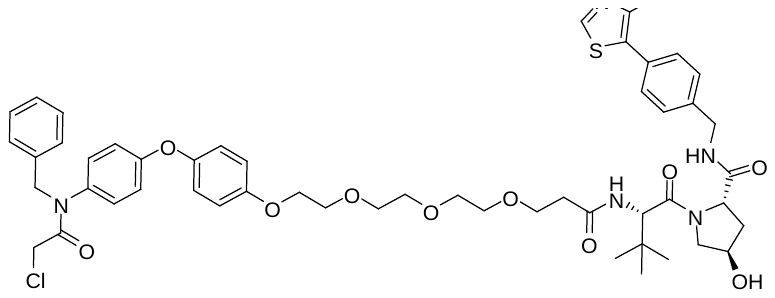

2-chloroacetyl chloride (3 µl, 37 µmol, 1.3eq) and TEA (12 µl, 86 µmol, 3.0eq) were added to a solution of (2S,4R)-1-((S)-1-(4-(4-(benzylamino)phenoxy)phenoxy)-14-(tert-butyl)-12-oxo-3,6,9-trioxa-13-azapentadecan-15-oyl)-4-hydroxy-N-(4-(4-methylthiazol-5-yl)benzyl)pyrrolidine-2-carboxamide **S28** (26 mg, 29 µmol, 1.0eq) in 2 mL DCM at 0 °C. The reaction mixture was stirred for 22 h at r.t. The reaction was quenched with water and the layers were separated. The aqueous layer was extracted with DCM (3x) and the combined organic layers were dried with MgSO_4_. Reverse phase flash column chromatography (ACN/water) yielded the title compound (8 mg, 28%) as yellow oil.

**^1^H NMR** (500 MHz, CD_2_Cl_2_): δ = 8.65 (s, 1H), 7.40-7.36 (m, 2H), 7.35-7.23 (m, 6H), 7.21-7.18 (m, 2H), 6.99-6.89 (m, 7H), 6.88-6.84 (m, 2H), 4.85 (s, 2H), 4.68 (t, *^3^J* = 8.0 Hz, 1H), 4.54 (dd, *^2^J* = 15.1 Hz, *^3^J* = 6.8 Hz, 1H), 4.47 (d, *^3^J* = 9.0 Hz, 1H), 4.42 (d, *^3^J* = 8.2 Hz, 1H), 4.29 (dd, *^2^J* = 15.1 Hz, 5.2 Hz, 1H), 4.11-4.03 (m, 3H), 3.89 (s, 2H), 3.81-3.77 (m, 2H), 3.72-3.59 (m, 11H), 3.56 (dd, *^3^J* = 11.4 Hz, *^3^J* = 3.6 Hz, 1H), 2.50-2.43 (m, 6H), 2.07 (ddt, *^2^J* = 13.3 Hz, *^3^J* = 8.1 Hz, *^4^J* = 1.9 Hz, 1H), 0.93 (s, 9H).

**^13^C NMR** (126 MHz, CD_2_Cl_2_): δ = 172.5, 172.4, 171.2, 166.7, 159.3, 156.2, 150.6, 149.8, 149.1, 139.1, 137.5, 135.5, 132.1, 131.5, 130.2, 129.9, 129.3, 128.98, 128.5, 128.1, 121.8, 118.5, 116.3, 71.3, 71.1, 71.0, 70.97, 70.7, 70.2, 68.5, 67.7, 58.9, 58.3, 57.1, 54.0, 43.6, 42.9, 37.2, 36.5, 35.1, 26.7, 16.5.

HPLC: R_t_ = 4.55 min (method B): Purity: >97% (254 nm); >95% (320 nm)

MS (ESI+):

318.1	[(fragment+2H)^+^	calc. 318.12]

667.3	[(M/2+H)^+^	calc. 667.28]

HRMS (ESI+):

984.3976	[(M+H)^+^	calc. 984.3979]

(2S,4R)-1-((S)-1-(4-(4-(N-benzyl-2-chloroacetamido)phenoxy)phenoxy)-20-(tert-butyl)-18-oxo-3,6,9,12,15-pentaoxa-19-azahenicosan-21-oyl)-4-hydroxy-N-(4-(4-methylthiazol-5-yl)benzyl)pyrrolidine-2-carboxamide **2c**

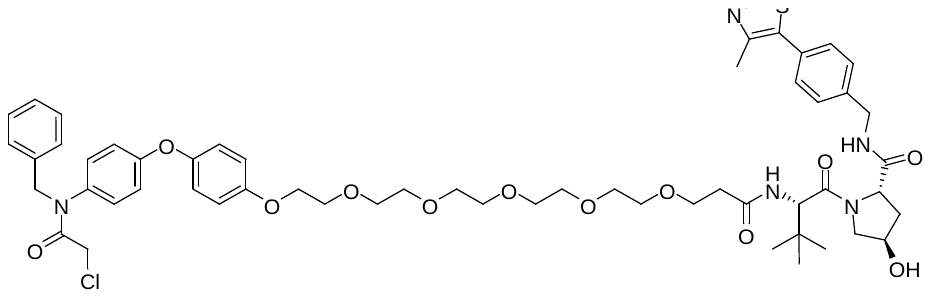

2-chloroacetyl chloride (3 µl, 39 µmol, 1.3eq) and TEA (12 µl, 84 µmol, 3.0eq) were added to a solution (2S,4R)-1-((S)-1-(4-(4-(benzylamino)phenoxy)phenoxy)-20-(tert-butyl)-18-oxo-3,6,9,12,15-pentaoxa-19-azahenicosan-21-oyl)-4-hydroxy-N-(4-(4-methylthiazol-5-yl)benzyl)pyrrolidine-2-carboxamide **S29** (28 mg, 28 µmol, 1.0eq) in 2 mL DCM at 0 °C. The reaction mixture was stirred for 22 h at r.t. The reaction was quenched with water and the layers were separated. The aqueous layer was extracted with DCM (3x) and the combined organic layers were dried with MgSO_4_. Reverse phase flash column chromatography yielded (ACN/water) the title compound as a yellow oil (14 mg, 46%).

**^1^H NMR** (500 MHz, CD_2_Cl_2_): δ = 8.66 (s, 1H), 7.40-7.31 (m, 5H), 7.31-7.23 (m, 3H), 7.23-7.17 (m, 2H), 6.99-6.90 (m, 7H), 6.88-6.84 (m, 2H), 4.85 (s, 2H), 4.68 (d, *^3^J* = 8.0 Hz, 1H), 4.55 (dd, *^2^J* = 15.1 Hz, *^3^J* = 6.8 Hz, 1H), 4.46 (s, 1H), 4.44 (d, *^3^J* = 8.2 Hz, 1H), 4.30 (dd, *^2^J* = 15.1 Hz, *^3^J* = 5.3 Hz, 1H), 4.10 (dd, *^3^J* = 5.4 Hz, *^3^J* = 4.0 Hz, 2H), 4.06 (d, *^2^J* = 11.5 Hz, 1H), 3.89 (s, 2H), 3.82-3.79 (m, 2H), 3.71-3.66 (m, 4H), 3.64-3.55 (m, 14H), 2.50-2.48 (m, 3H), 2.47-2.41 (m, 3H), 2.08 (ddt, *^2^J* = 13.3 Hz, *^3^J* = 8.1 Hz, *^4^J* = 1.9 Hz, 1H), 0.93 (s, 9H).

**^13^C NMR** (126 MHz, CD_2_Cl_2_): δ = 172.6, 172.4, 171.3, 166.7, 159.3, 156.3, 150.6, 149.8, 149.1, 139.1, 137.5, 135.5, 132.1, 131.5, 130.1, 129.9, 129.3, 128.97, 128.5, 128.1, 121.8, 118.5, 116.3, 71.3, 71.08, 71.06, 71.04, 71.02, 70.99, 70.91, 70.7, 70.2, 68.5, 67.7, 58.9, 58.3, 57.1, 43.5, 42.9, 37.2, 36.6, 35.2, 26.7, 16.5.

HPLC: R_t_ = 4.53 min (method B): Purity: >98% (254 nm); >97% (320 nm)

MS (ESI+):

318.1	[(fragment+2H)^+^	calc. 318.12]

536.9	[(M/2+H)^+^	calc. 536.73]

755.4	[(fragment)^+^	calc. 755.33]

HRMS (ESI+):

1072.4507	[(M+H)^+^	calc. 1072.4503]

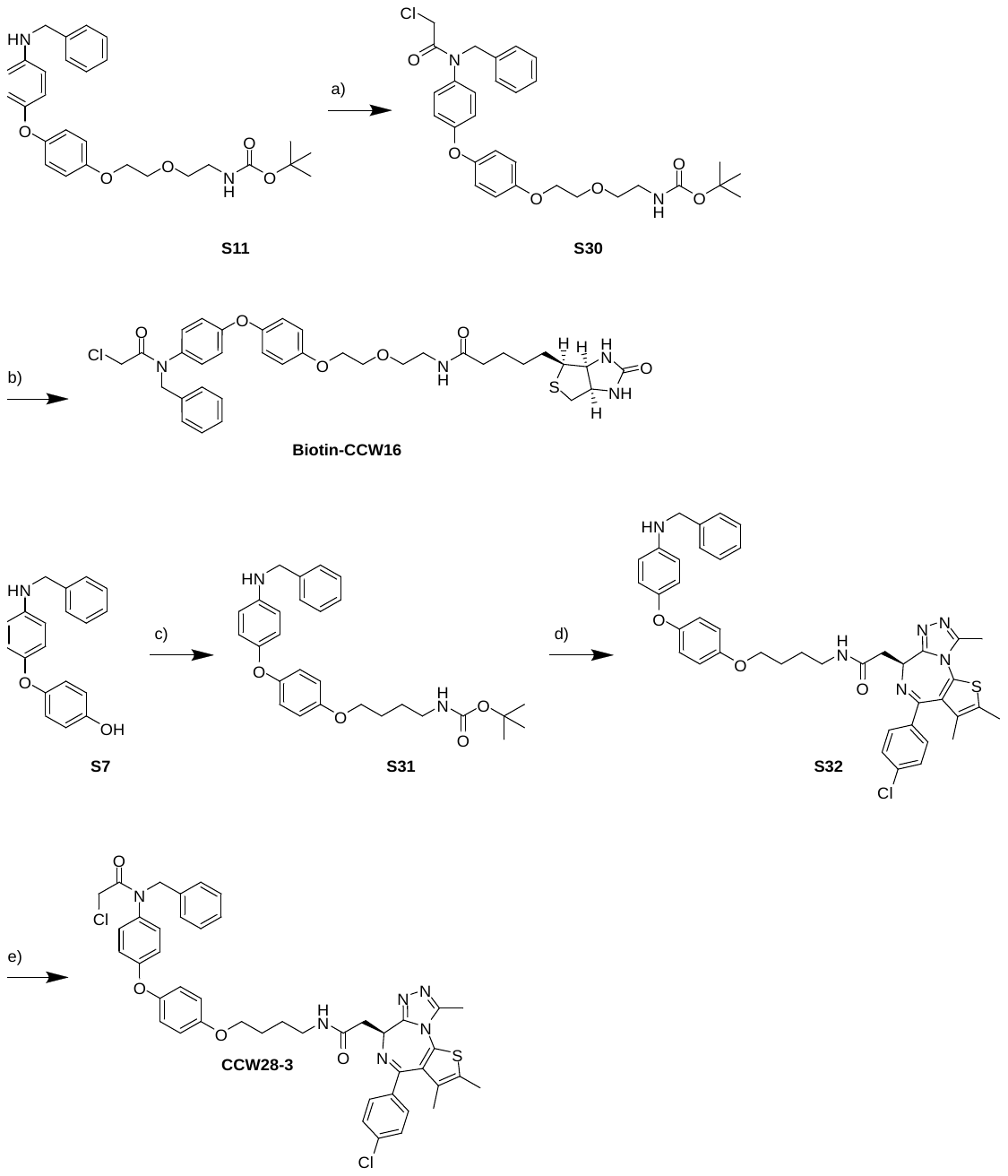

**Figure S##:** Synthesis of **Biotin-CCW16** and BRD4 PROTACs **CCW28-3**: a) 2-chloroacetyl chloride, TEA, 0 °C, then r.t., 23 h; b) 1. TFA, DCM, r.t., 2 h; 2. DIPEA, Biotin-NHS ester, DMF, r.t., 2 h; c) K_2_CO_3_, linker-bromide, Acetone, reflux, overnight; d) 1. TFA, DCM, r.t., 3 h; 2. DIPEA, JQ1-acid, HATU, DMF, r.t., 16 h; e) 2-chloroacetyl chloride, TEA, 0 °C, then r.t., 23 h.

N-(2-(2-(4-(4-(N-benzyl-2-chloroacetamido)phenoxy)phenoxy)ethoxy)ethyl)-5-((3aS,4S,6aR)-2-oxohexahydro-1H-thieno[3,4-d]imidazol-4-yl)pentanamide **Biotin-CCW16** via tert-butyl (2-(2-(4-(4-(N-benzyl-2-chloroacetamido)phenoxy)phenoxy)ethoxy)ethyl)carbamate **S30**

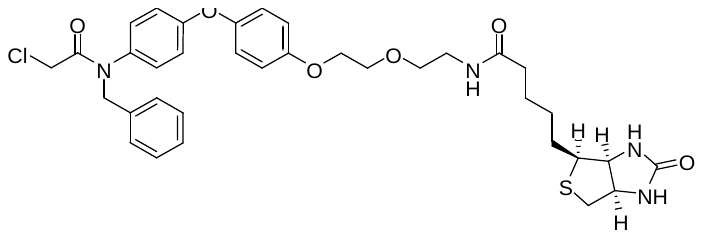

2-chloroacetyl chloride (32 µL, 401 µmol, 4.0eq) and TEA (56 µl, 401 µmol, 4.0eq) were added to a solution of tert-butyl (2-(2-(4-(4-(benzylamino)phenoxy)phenoxy)ethoxy)ethyl)carbamate **S11** (48 mg, 100 µmol, 1.0eq) in 2 mL DCM at 0 °C. The reaction mixture was stirred for 23 h at r.t. The reaction was quenched with water and the layers were separated. The aqueous layer was extracted with DCM (3x) and the combined organic layers were dried with MgSO_4_. The solvent was removed under reduced pressure and the crude product was used without further purification for the next step (54 mg, 97%).

A solution of tert-butyl (2-(2-(4-(4-(N-benzyl-2-chloroacetamido)phenoxy)phenoxy)ethoxy)ethyl)carbamate **S30** (56 mg, 101 µmol, 1.0eq) in TFA/DCM (4 mL, 1/1) was stirred for 2 h at r.t. all volatiles were removed under reduced pressure, DCM was added and the all volatiles were removed under reduced pressure. This process was repeated two additional times.

A solution of the crude amine, DIPEA (100 µL, 575 µmol, 5.7eq) and 2,5-dioxopyrrolidin-1-yl 5-((3aS,4S,6aR)-2-oxohexahydro-1H-thieno[3,4-d]imidazol-4-yl)pentanoate (34 mg, 101 µmol, 1.0eq) in DMF (2 mL) was stirred for 2 h at r.t. Water and DCM were added and the layers were separated. The aqueous layer was extracted with DCM (2x) and the combined organic layers were dried with MgSO_4_. The solvent was removed under reduced pressure and the crude product was purified using reversed phase column chromatography (ACN/water). The title compound was isolated as a beige solid (4 mg, 6%).

**^1^H NMR** (500 MHz, CD_2_Cl_2_): δ = 7.31-7.24 (m, 3H), 7.22-7.18 (m, 2H), 7.01-6.91 (m, 6H), 6.89-6.85 (m, 2H), 6.25 (t, *^3^J* = 4.9 Hz, 1H), 5.83 (s, 1H), 5.03 (s, 1H), 4.85 (s, 2H), 4.47-4.43 (m, 1H), 4.30-4.25 (m, 1H), 4.12-4.09 (m, 2H), 3.82-3.80 (m, 2H), 3.61 (t,*^3^J* = 5.2 Hz, 2H), 3.45-3.40 (m, 2H), 3.13 (td, *^3^J* = 7.4 Hz, *^3^J* = 4.6 Hz, 1H), 2.88 (dd, *^2^J* = 12.8 Hz, *^3^J* = 5.0 Hz, 1H), 2.68 (d, *^2^J* = 12.8 Hz, 1H), 2.23-2.10 (m, 2H), 1.75-1.57 (m, 4H), 1.46-1.38 (m, 2H).

**^13^C NMR** (126 MHz, CD_2_Cl_2_): δ = 173.4, 166.7, 163.9, 159.3, 156.2, 149.9, 137.5, 135.6, 130.2, 129.3, 128.99, 128.1, 121.9, 118.5, 116.3, 70.6, 70.0, 68.5, 62.2, 60.7, 55.9, 54.1 42.9, 41.2, 39.7, 36.3, 28.6, 28.5, 26.1.

HPLC: R_t_ = 4.03 min (method A):	Purity: >96% (254 nm); >95% (320 nm)

MS (ESI+):

341.0	[(M/2+H)^+^	calc. 341.13]

681.3	[(M+H)^+^	calc. 681.25]

HRMS (ESI+):

681.2536	[M+H)^+^	calc. 681.2508]

tert-butyl (4-(4-(4-(benzylamino)phenoxy)phenoxy)butyl)carbamate **S31**

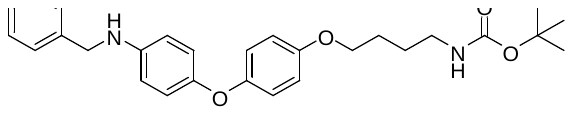

A solution of the 4-(4-(benzylamino)phenqq3457oxy)phenol **S7** (72 mg, 247 µmol, 1.0eq), tert-butyl (4-bromobutyl)carbamate (93 mg, 371 µmol, 1.5eq) and potassium carbonate (102 mg, 741 µmol, 3.0eq) in Acetone (10 mL) was refluxed overnight. The solvent was removed under reduced pressure. Reverse phase column chromatography (ACN/water) yielded the title compound as a colorless oil (89 mg, 78%).

**^1^H NMR** (400 MHz, CD_2_Cl_2_): δ = 7.42-7.24 (m, 4H), 6.90-6.78 (m, 6H), 6.63-6.57 (m, 2H), 4.31 (s, 2H), 4.08 (s, 2H), 3.93 (t, *^3^J* = 6.3 Hz, 2H), 3.15 (q, *^3^J* = 6.7 Hz, 2H), 1.81-1.73 (m, 2H), 1.69-1.56 (m, 2H), 1.42 (s, 9H).

**^13^C NMR** (101 MHz, CD_2_Cl_2_): δ = 156.4, 155.0, 152.8, 149.7, 145.0, 140.3, 129.1, 128.0, 127.7, 120.5, 119.4, 115.9, 114.3, 79.2, 68.7, 49.2, 40.8, 28.7, 27.4, 27.2.

MS (ESI+):

407.1	[(-^t^Bu+2H)^+^	calc. 407.19]

485.3	[(M+Na)^+^	calc. 485.24]

(S)-N-(4-(4-(4-(benzylamino)phenoxy)phenoxy)butyl)-2-(4-(4-chlorophenyl)-2,3,9-trimethyl-6H-thieno[3,2-f][1,2,4]triazolo[4,3-a][1,4]diazepin-6-yl)acetamide **S32**

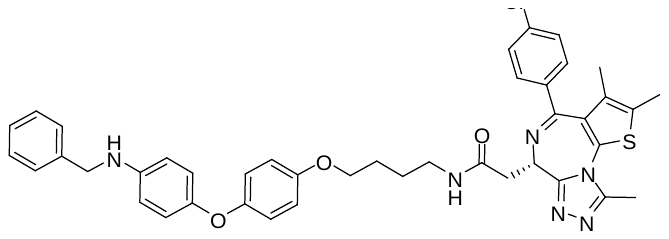

A solution of tert-butyl (4-(4-(4-(benzylamino)phenoxy)phenoxy)butyl)carbamate **S31** (30 mg, 65 µmol, 1.1eq) in 4 mL DCM/TFA (1/1) was stirred for 3 h. All volatiles were removed under reduced pressure. DCM was added and the solvent was removed under reduced pressure. This procedure was repeated once.

A solution of DIPEA (21 µL, 118 µmol, 2.0eq), (S)-2-(4-(4-chlorophenyl)-2,3,9-trimethyl-6H-thieno[3,2-f][1,2,4]triazolo[4,3-a][1,4]diazepin-6-yl)acetic acid (24 mg, 59 µmol, 1.0eq) and HATU (27 mg, 71 µmol, 1.2eq) in DMF (1.5 mL) was stirred for 10 min. The crude amine was added and the reaction mixture was stirred for 16 h. DCM, water and brine were added and the layers were separated. The aqueous layer was extracted with DCM (3x) and the combined organic layers were dried with MgSO_4_. The solvent was evaporated under reduced pressure and the residue was purified by reverse flash column chromatography to yield the title compound as a colorless solid (34 mg, 77%).

**^1^H NMR** (500 MHz, DMSO*-d_6_*): δ = 8.22 (t, *^3^J* = 5.6 Hz, 1H), 7.44 (d, *^3^J* = 8.8 Hz, 2H), 7.41 (d, *^3^J* = 8.7 Hz, 2H), 7.36 (d, *^3^J* = 7.4 Hz, 2H), 7.32 (t, *^3^J* = 7.6 Hz, 2H), 7.22 (t, *^3^J* = 7.2 Hz, 1H), 6.87-6.82 (m, 2H), 6.81-6.77 (m, 2H), 6.75-6.70 (m, 2H), 6.59-6.54 (m, 2H), 6.09 (t, *^3^J* = 6.0 Hz, 1H), 4.51 (dd, *^3^J* = 8.3 Hz, *^3^J* = 5.8 Hz, 1H), 4.23 (d, *^3^J* = 5.9 Hz, 2H), 3.91 (t, *^3^J* = 6.4 Hz, 2H), 3.29-3.09 (m, 4H), 2.59 (s, 3H), 2.40 (s, 3H), 1.77-1.69 (m, 2H), 1.63-1.55 (m, 5H).

**^13^C NMR** (126 MHz, DMSO*-d_6_*): δ = 169.4, 162.98, 155.1, 153.8, 151.9, 149.8, 147.2, 145.1, 140.3, 136.7, 135.2, 132.2, 130.7, 130.1, 129.8, 129.6, 128.4, 128.2, 127.2, 126.6, 119.8, 118.3, 115.3, 113.1, 67.6, 53.9, 46.98, 39.5, 38.2, 37.7, 26.2, 25.9, 14.0, 12.7, 11.3.

MS (ESI+):

373.1	[(M/2+H)^+^	calc. 373.14]

655.2	[(M-Bn+2H)^+^	calc. 655.22]

745.3	[(M+H)^+^	calc. 745.27]

(S)-N-benzyl-2-chloro-N-(4-(4-(4-(2-(4-(4-chlorophenyl)-2,3,9-trimethyl-6H-thieno[3,2-f][1,2,4]triazolo[4,3-a][1,4]diazepin-6-yl)acetamido)butoxy)phenoxy)phenyl)acetamide **CCW28‑3**

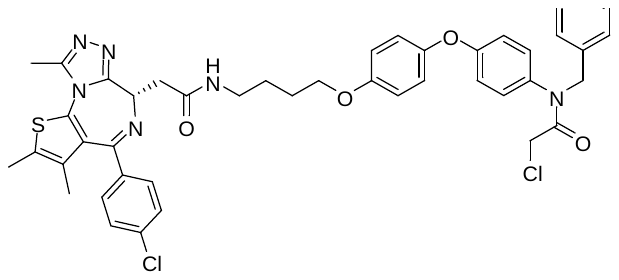

2-chloroacetyl chloride (4 µL, 52 µmol, 1.3eq) and TEA (28 µl, 201 µmol, 5.0eq) were added to a solution of (S)-N-(4-(4-(4-(benzylamino)phenoxy)phenoxy)butyl)-2-(4-(4-chlorophenyl)-2,3,9-trimethyl-6H-thieno[3,2-f][1,2,4]triazolo[4,3-a][1,4]diazepin-6-yl)acetamide **S32** (30 mg, 40 µmol, 1.0eq) in 2 mL DCM at 0 °C. The reaction mixture was stirred for 23 h at r.t. The reaction was quenched with water and the layers were separated. The aqueous layer was extracted with DCM (3x) and the combined organic layers were dried with MgSO_4_. The solvent was removed under reduced pressure and the crude material was purified using reversed phase flash column chromatography (H_2_O/ACN). The title compound was isolated as a colorless solid (22 mg, 67%).

**^1^H NMR** (500 MHz, CD_2_Cl_2_): δ = 7.46-7.40 (m, 2H), 7.33 (d, *^3^J* = 8.8 Hz, 2H), 7.31-7.23 (m, 3H), 7.22-7.18 (m, 2H), 6.99-6.93 (m, 4H), 6.91-6.84 (m, 4H), 6.63 (t, *^3^J* = 5.6 Hz, 1H), 4.85 (s, 2H), 4.59 (t, *^3^J* = 6.9 Hz, 1H), 3.95 (t, *^3^J* = 6.3 Hz, 2H), 3.89 (s, 2H), 3.45 (dd, *^3^J* = 14.3 Hz, *^3^J* = 7.3 Hz, 1H), 3.38 (td, *^3^J* = 13.2 Hz, *^3^J* = 6.9 Hz, 1H), 3.33-3.25 (m, 2H), 2.63 (s, 3H), 2.41-2.36 (m, 3H), 1.85-1.78 (m, 2H), 1.75-1.68 (m, 3H), 1.67 (s, 3H).

**^13^C NMR** (126 MHz, CD_2_Cl_2_): δ = 170.7, 166.7, 164.4, 159.4, 156.5, 156.3, 150.6, 149.6, 137.5, 137.4, 137.1, 135.5, 132.9, 131.6, 131.3, 130.9, 130.5, 130.1, 129.3, 129.1, 128.98, 128.1, 121.8, 118.4, 116.2, 68.6, 55.1, 42.9, 39.9, 39.6, 27.2, 26.9, 14.7, 13.4, 12.2.

HPLC: R_t_ = 5.36 min (method A):	Purity: >99% (254 nm); >99% (320 nm)

MS (ESI+):

411.2	[(M/2+H)^+^	calc. 411.13]

821.3	[(M+H)^+^	calc. 821.24]

HRMS (ESI+):

821.2434	[(M+H)^+^	calc. 821.2438]
